# Immediate perturbations of DNA methylation and transcriptome upon acute prenatal alcohol exposure in the mouse developing brain cortex

**DOI:** 10.1101/2022.03.09.481186

**Authors:** Agathe Duchateau, Carole Chaput, Lea Livramento, Névé Auperin, Federico Miozzo, Olivier Kirsh, Anne Le Mouel, Myriame Mohamed, Sascha Ott, Délara Saberan-Djoneidi, Valérie Mezger

## Abstract

The reshaping of the DNA methylome landscape after prenatal alcohol exposure (PAE) has been well-documented in the adult brain, therefore a long time after the end of the exposure. However, the question of the immediate deposition or loss of DNA methylation marks in the prenatal neocortex, just after the end of PAE has not yet been directly addressed, genome widely.

Using a binge-drinking-like model of PAE and capture of the DNA methylome, we have identified differentially methylated regions (DMRs) that are established immediately, within two hours after the end of PAE. Remarkably, these DMRs are prominently and statistically associated with: (i) enhancers that are active in the brain, associated with GO terms of importance for neurogenesis, neurodevelopment, and neuronal differentiation; (ii) genes that, in physiological conditions show dynamic gain in chromatin accessibility and/or upregulation of their expression in the time-window of exposure; (iii) imprinted genes and members of protocadherin genes clusters, two gene families playing key roles in neurodevelopment, whose mono-allelically expression is regulated by DNA methylation and impaired upon PAE. We observed that DMR-containing mono-allelically expressed genes, as well as other genes important for neurodevelopment, are also immediately upregulated upon PAE, suggesting that these early DNA methylation perturbations are thus highly susceptible to rapidly alter gene expression after PAE. DMRs in imprinted and protocadherin genes have been previously identified, both in the adult rodent brain prior-exposed to alcohol prenatally, and in cohorts of children diagnosed with fetal alcohol spectrum disorders (FASD). Our study thus strongly suggests that the DNA methylation profiles of key regulatory regions of these gene families are very quickly disturbed after the PAE and that these immediate altered regions could be persistently affected long after the stress. This strongly reinforces their potential as future biomarkers of PAE. In addition, binge-drinking-like PAE provokes immediate rewiring of the transcriptome with potentially dual consequences: 1) beneficial impacts that could support the recovery of cells from alcohol exposure, through slowing down of protein synthesis and energy-consuming respiratory pathways; 2) detrimental effects, through the inappropriate activation of critical pathways may directly perturb neurodevelopment.

**Ethical issues:** The breeding and treatments of wild type C57BL/6N mice, used for the experimental protocols described in this study have been approved by the Institutional Animal Care and Use Ethical Committee of the Paris Cité University (registration number CEEA-40). The project has been recorded under the following reference by the Ministère de l’Enseignement Supérieur et de la Recherche (#2016040414515579). All efforts were made to reduce stress and pain to animals.

## INTRODUCTION

The prenatal developing brain is particularly vulnerable to detrimental events that generate neurodevelopmental defects and potentially have long-term consequences in the adulthood (Bale et al., 2010; Schang et al., 2018a). Among a diversity of adverse *in utero* stresses, prenatal alcohol exposure (PAE) is a leading cause of non-genetic mental retardation in the Western world (Popova et al., 2012, 2016). Depending on many parameters, such as the timing of exposure, the drinking pattern (chronic or acute) and the amount of alcohol consumed, PAE gives rise to a wide range of neurodevelopmental defects, referred to as part of the clinical picture of fetal alcohol spectrum disorders (FASD), whose prevalence is estimated around 9 for 1000 live births (Burd et al., 2003; Jones and Smith, 1973; Kleiber et al., 2013; Lemoine et al., 1968; Mattson et al., 2011; Popova et al., 2012, 2016). These defects lead to impairment in cognition, behavior, executive function, attention (linked or not to hyperactivity), learning, judgment and social adaptation (Gibbard et al., 2003). In addition to these primary defects, individuals diagnosed with FASD are at high risk for neuropsychiatric disorders, including anxiety disorders, depression, and addiction in their adulthood (Gibbard et al., 2003; reviewed in Kodituwakku, 2007 and O’Connor and Paley, 2009, in line with the DOHaD concept (Developmental Origins of Health and Disease; Schang et al., 2018b). The most severe form of FASD, called fetal alcohol syndrome (FAS), is defined by characteristic facial dysmorphology, growth retardation and severe macroscopic structural abnormalities of the central nervous system (CNS; Jones and Smith, 1973). Early diagnosis of PAE is necessary to enable efficient intervention (Burd et al., 2003; Paley and O’Connor, 2011). Visible and well-characterized phenotypic defects observed for FAS newborns facilitated the early diagnosis of these individuals. In contrast, diagnosis of FASD children is more challenging, because the history of exposure is often unknown and clinical phenotypes are more cryptic, yet crucial since the functioning of the central nervous system damages is compromised (Gibbard et al., 2003; Popova et al., 2015). Indeed, PAE affects neurodevelopment at any stage, therefore impacting all neurodevelopmental processes, such as proliferation of neural progenitors, migration of young post-mitotic neurons and their differentiation, neuronal survival, synaptogenesis, neurotransmission and neuronal plasticity (Kleiber et al., 2013; El Fatimy et al., 2014; Guerri et al., 2009; Hashimoto-Torii et al., 2014; Ishii et al., 2017).

Although defects caused by prenatal alcohol exposure are well identified, the exact molecular mechanisms underlying these alterations and their persistence are still unclear. However, PAE is known to long-lasting impact on gene expression levels. These transcriptional disturbances, observed in the adult brain, occur in a manner dependent on the developmental stage at which the exposure has occurred. Indeed, transcriptional changes affect genes falling into distinct GO (Gene Ontology) categories, depending on the time-window of exposure in preclinical models, (equivalent to the first to third trimester of pregnancy in women) (Kleiber et al., 2013, 2014). These long-lasting disturbances therefore affect processes governed by gene networks that are active during the gestation, at the time of exposure. This “memory” of the exposure could occur *via* Epigenetic mechanisms.

Remarkably, brain development is tightly controlled by epigenetic mechanisms, as underlined by the impact of mutations or variants in genes encoding epigenetic actors in the emergence of neurodevelopmental and neuropsychiatric disorders, such as Rett syndrome, Rubinstein-Taybi syndrome and autism spectrum disorders (Bourgeron, 2015; Gräff et al., 2011; LaSalle et al., 2013). The long-lasting and stage-specific transcriptomic alterations, described above and observed in the PAE-exposed brain, might be underlined by modifications in the epigenetic landscape, including perturbation of DNA methylation profile. Indeed, starting from seminal works on rodent models (*e.g.* Haycock and Ramsay, 2009; Kaminen-Ahola et al., 2010), a number of studies have identified disturbances in DNA methylation on candidate genes or *loci*, or in a genome-wide manner, in the mouse adult brain that was exposed to alcohol prenatally (Kleiber et al., 2013; Laufer et al., 2013; reviewed in Lussier et al., 2017). Alterations in DNA methylation profile observed in mouse brain in response to PAE has been corroborated in peripheral tissues (cheek swabs) in cohorts of FASD children (Laufer et al., 2015; Lussier et al., 2018). PAE during the preimplantation period has also been associated to DNA methylation perturbations observed more than one week after and associated to neurodevelopmental defects (Legault et al., 2021).

Strikingly, the short-term impacts of PAE on the genome-wide architecture of DNA methylation in the developing brain have been understudied and there is a need to investigate whether the aberrant DNA methylation events are the direct and immediate results of alcohol exposure. First, this is of importance for the field, because DNA methylation disturbances observed after birth or in the adulthood could (i) result from immediate PAE-dependent DNA methylation changes that would persist throughout life or (ii) be due to PAE-induced brain dysfunction. Indeed, in the last case, since neuronal activity can reshape DNA methylation throughout lifetime (Guo et al., 2011), the remodeling of the DNA methylome could represent secondary effects of PAE, resulting from the PAE-induced alteration of neuronal activity in the infant or adult brain. Second, this is also a question of medical relevance. In many cases, the history of exposure to alcohol is unknown and FASD children are often diagnosed late, which compromised early intervention (Gibbard et al., 2003). The search for accurate and relevant molecular biomarkers, especially for biomarkers of exposure, is thus necessary. Attempts to identify biomarkers of exposure linked to DNA methylation perturbations have been given special attention in the field. Ideally, such DNA methylation changes should occur quickly after exposure. This has remained to be determined.

We thus asked whether immediate changes in DNA methylation could be detected across the genome, in the developing mouse cerebral cortex, using a tailor-made methylome capture approach. To capture immediate events of DNA methylation alteration and since binge drinking has become an increasing mode of alcohol consumption in young women (Dwyer-Lindgren et al., 2015; Tan et al., 2015; Popova et al., 2018), we chose a binge-drinking-like mouse model of PAE. We observe hundreds of differentially methylated regions (DMRs), as early as two hours after the last alcohol injection. We show that DMRs are significatively associated to genes or loci which, in physiological conditions, are dynamically regulated at the time of exposure, in terms of chromatin accessibility or expression, and which are linked to biological processes of importance for brain development and functions. Moreover, DMRs are overrepresented in two families of mono-allelically expressed genes of importance for neurodevelopment — imprinted genes and genes of clustered protocadherins — and correlate with immediate modifications in their transcript levels. Our results therefore show that some genomic regions are the target of differential methylation events, very rapidly after exposure, that are associated with concomitant changes in gene expression. Interestingly, according to published results, DMRs associated with imprinted genes and protocadherins have been observed at temporal distance of PAE, in the mouse and/or human postnatal and adult brain, this suggest that these *loci* might represent immediate targets of PAE with long-lived DNA methylation perturbation and provide valuable and meaningful biomarkers of exposure in the future.

## RESULTS

### Working model: a methylome capture in a murine binge drinking model

To identify DNA methylation changes that would occur immediately after an acute prenatal alcohol exposure, we used a murine model of binge drinking at a developmental stage equivalent to the second trimester of pregnancy in human: mouse embryonic cortices were exposed to PAE, *in utero*, at embryonic days E15 and E16 (**Figure 1A**, see **Materials and Methods**). We chose this binge-drinking model to be as close as possible to a human mode of acute consumption. The quantity of injected ethanol corresponded to a dose known to induce brain defects that mimic FAS, in rodent pups (Carloni et al., 2004; Ikonomidou et al., 2000; Olney et al., 2002). We searched for methylation modifications immediately occurring after PAE, using a capture of the DNA methylome (*EpiCapture* technology; see **Materials and Methods**), which allowed us to study, with a good resolution, a repertoire of 58,611 selected DNA regions corresponding to about 81.3 Mb. In particular, the capture design was composed of more than 75% of mouse promoters and all the enhancers that are active in the brain (**Figure 1B**, **Supplementary Figure 1A**; see **Materials and Methods** for details).

**Figure 1.**
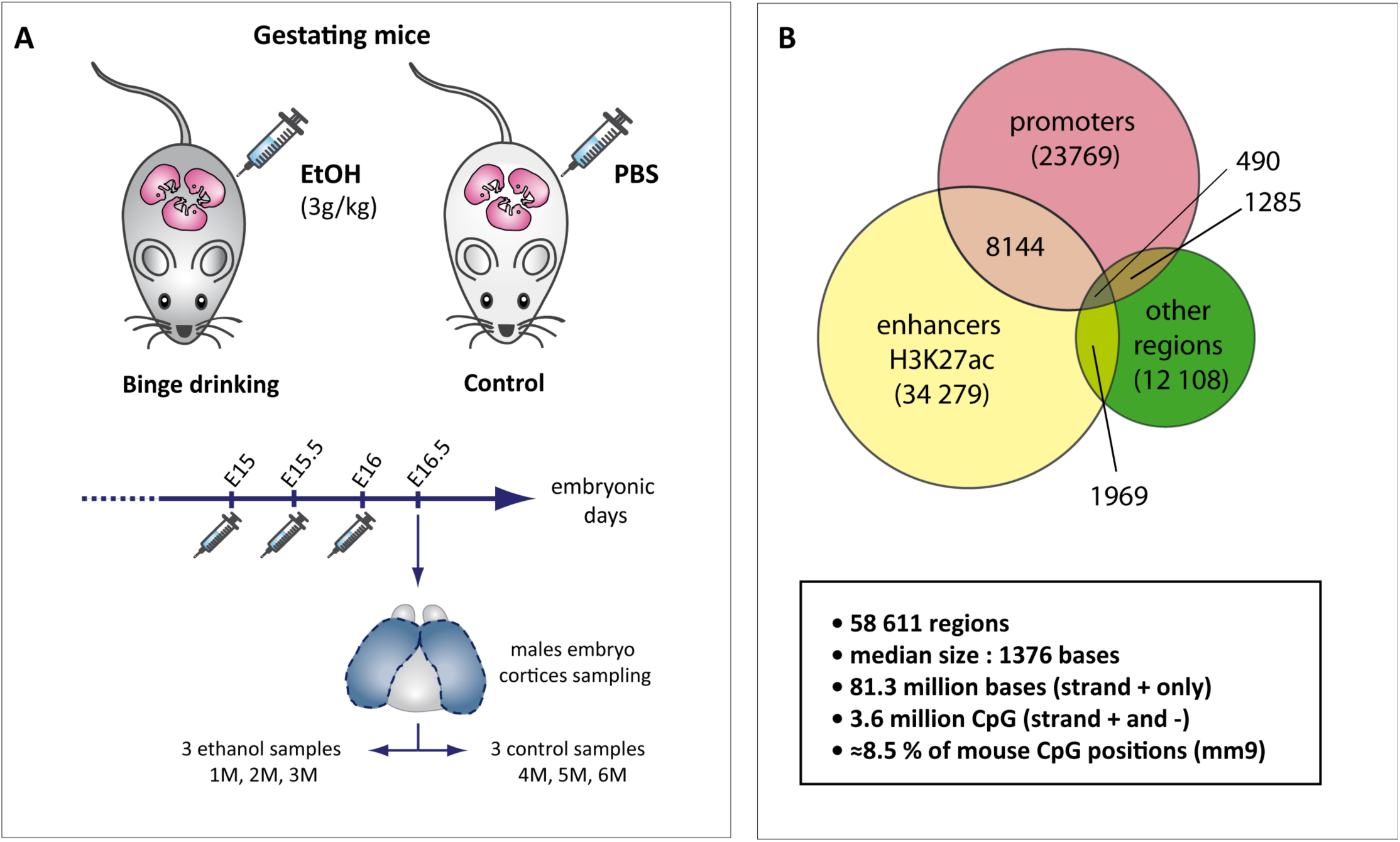
Binge drinking model and DNA methylome capture design. **(A) *Binge drinking model***. C57BL/6N pregnant mice received 1 intraperitoneal injection (IP)/day of ethanol (3g/kg) at embryonic days E15, E15.5 and E16. Control animals were similarly treated with IP of PBS. Fetal cortices were collected 2 hours after the last IP (at E16). For DNA methylome analysis, only male samples were studied. Three samples per group were generated (4M, 5M and 6M for the control group; 1M, 2M and 3M for the EtOH-treated one). To reduce variability between samples, each replicate was composed of 4 hemi-cortices, from 4 embryos of distinct litters (**Supplementary Figure 1B**). **(B) *Design of the DNA methylome capture*.** The capture is composed of 58,611 chosen genomic regions, based on ENCODE available data (H3K27ac histone mark, a hallmark of enhancer regions and promoter regions) and on our previous results (for *other regions*). For more details about capture composition, see **Supplementary Figure 1A** and **Materials and methods**.

### Quality control of the methylome capture

This methylome capture approach used sodium bisulfite (BS) conversion, to distinguish between methylated and unmethylated cytosines. First, we verified that the BS conversion had correctly occurred for all samples (Clark et al., 2006; Krueger et al., 2012). Abnormal *per-base-sequence-content* plot obtained with FASTQC was observed for each sample, as expected for DNA sequences converted with BS (**Supplementary Figure 2A,B**). Moreover, BS conversion rate of all samples was quantified, using spike-in DNA. Conversion rate was high (>97,8%) for all the samples, which means that no conversion biases were present in our data analyses (**Supplementary Table 1**).

To minimize inter-individual variability in the same group, we mixed half-cortices from 4 embryos from 4 different litters (2 left and 2 right half-cortices; **Supplementary Figure 1B**). In line with our sampling choice, we observed that intra-group variability was limited, according to hierarchical clustering analysis and Pearson correlation between samples (**Supplementary Figure 2C,D**). However, triplicates from the EtOH-treated group showed more differences between each other than did triplicates from control group (PCA analysis and hierarchical clustering analysis, **Supplementary Figure 2C,E**), suggesting that, even if samples were globally affected in the same manner by *in utero* alcohol exposure, some subtle differences in term of CpG methylation level could be observed. In addition, CpG base Pearson correlation values indicated that inter-group variability was also limited. This suggests that PAE does not majorly reshape the DNA methylome.

Using the bioinformatics workflow described in **Supplementary Figure 2A**, to compare control and alcohol-treated groups, we only identified three hyper- and one hypomethylated isolated (individual) CpGs, all having a high methylation differential percent (> 35%; **Figure 2A**; *MethylKit* analysis on normalised and filtered datasets, meth. diff < 5%, qval. < 0.05). Two of these differentially methylated cytosines (DMC) are located in intergenic regions (on chromosomes 5 and 10). One hypermethylated CpG is in the *Kalrn* gene, encoding the Kalirin RhoGEF kinase that is involved in various mechanisms, such as neuronal shape regulation, growth and plasticity. The third hypermethylated CpG is associated to *Tiam2* gene, encoding a RAC1-specific guanine nucleotide exchange factor.

**Figure 2.**
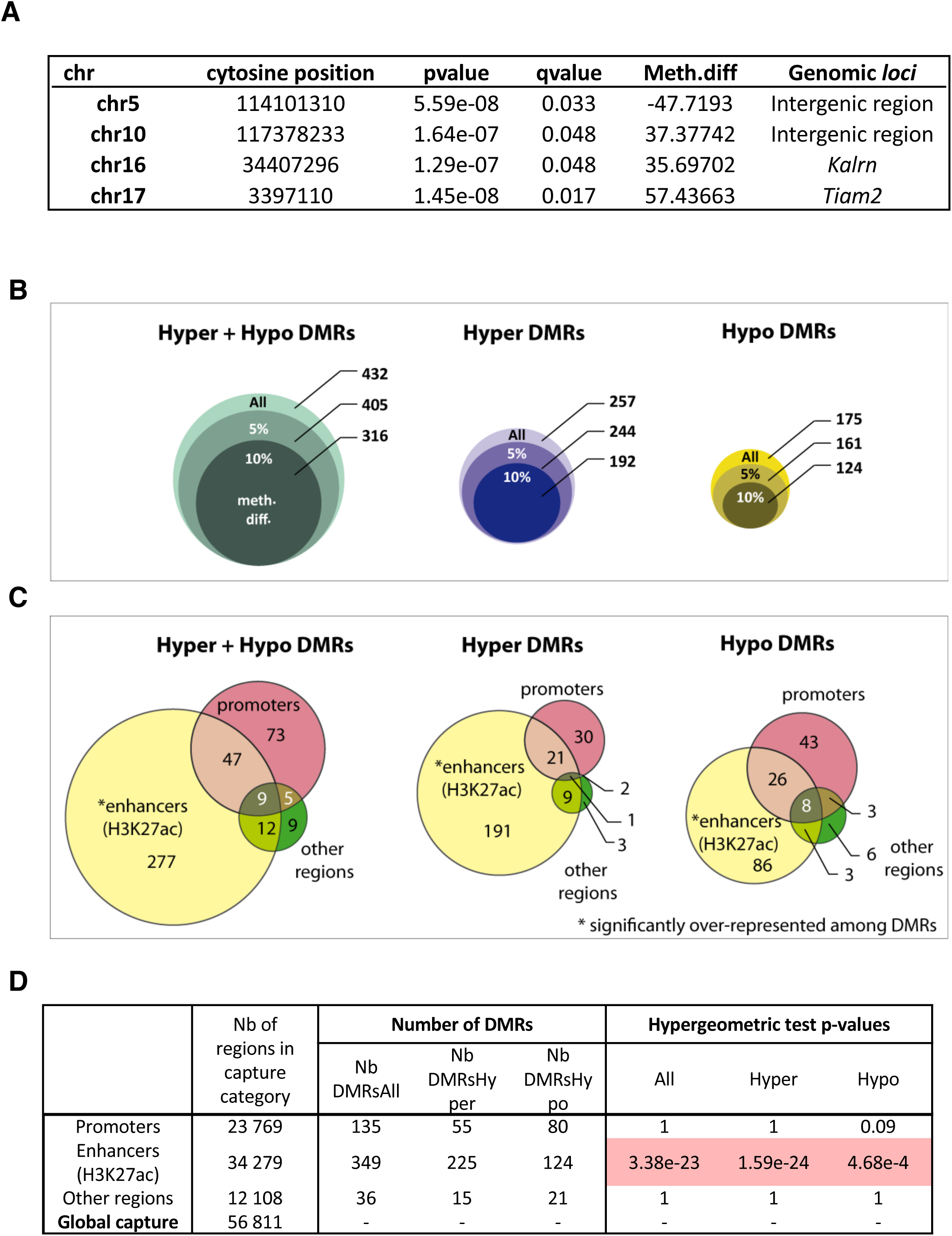
Immediate modifications of DNA methylation are observed in fetal cortices upon PAE. **(A) Differentially methylated CpG (DMC) identified upon PAE.** Only cytosines in CpG context that are covered in all samples were investigated. DMC were identified using *MethylKit* (see details in **Materials and Methods**). *Meth.diff:* differential methylation rate (%) observed between control and EtOH-treated groups. A negative percentage indicates a loss of methylation, compared to control. q-val. < 0.05; a minimum of 5% of Meth.diff is required for a CpG to be considered a DMC. **(B) Number of differentially methylated regions (DMRs) immediately observed upon PAE.** Number of DMRs filtered by methylation differential rate (indicated as percent) observed between EtOH-treated and control groups are also indicated. Regions with gain (Hyper-DMRs) or loss (Hypo-DMRs) of methylation; (Hyper+Hypo), regions with either gain or loss of methylation. All: all DMRs. **(C) Number of DMRs in each category of capture regions.** A DMR is considered to be associated to a category of capture regions, if it overlaps to or if it is closed to a capture region (distance less than 500 bp). Asterisks indicate categories that are significantly over-represented among DMRs, in a statistical manner, according to hypergeometric tests (see (**D**)). **(D) DMRs immediately detected upon PAE are predominantly located in brain active enhancers**. Results of hypergeometric tests realized to determine whether specific categories of capture regions nare over-represented among DMRs. Significant values are highlighted in red. For this analysis, both DMRs that overlap or are located closed to capture region categories) were considered (*i.e.* distance less than 500bp between the regions. The number (Nb) of DMRs considered in each analysis is indicated. They differ very slightly from the numbers indicated in (**C**), because some DMRs overlap several capture regions, and are, thus, counted more than one time.

### Definition of DNA methylated regions for capture analysis

Although the methylation of an individual CpG can affect gene expression (Xu et al., 2007), DMRs are considered as more relevant in terms of impact on gene expression. In line with this consideration, studies of differentially methylated regions (DMRs) have been more frequently performed (Bock, 2012). We therefore pursued our analyses on the identification of DMRs.

Bioinformatic analysis of this methylome capture required special attention, especially to define potential differentially methylated regions (DMRs) between control and alcohol-treated groups. Indeed, to define such genomic regions, existing bioinformatics tools were often design for analyses considering the whole genome (or well-characterised arrays), as a reference. This kind of approach is not adapted to a customized capture based on a repertoire of selected regions. For this reason, we developed our own *R* function to define DMRs between control and alcohol-treated groups, which combined neighbouring CpGs that share similar differential methylation states, either all showing loss of methylation (hypo CpGs) or all showing gain of methylation (hyper CpGs). More precisely, we defined DMRs on the following basis (see **Materials and Methods**; **Supplementary Figure 3)**: the association of at least 5 CpG having a same methylation state, in a region of maximum 2000 bases, with a maximum distance of 100 bases between two successive selected CpG, according to a p-value threshold (**Supplementary Table 2**). For this, a pre-selection of *relevant CpGs* was done, according to their p-value (CpGs with p-value < 0.07). This p-value threshold was determined using comparisons of real and random datasets (see below, **Supplementary Figure 4**, **Supplementary Table 2**).

### Immediate alterations in DNA methylation are detected in brain development upon PAE

Using this captured-specific bioinformatic workflow (**Supplementary Figure 2A**; **Supplementary Table 3**, see **Materials and Methods** and **Supplementary Data notebook 1**) we identified 432 regions that were differentially methylated in embryonic cortices, immediately after PAE, among the 58,611 regions included in the capture (**Figure 2B**, **Supplementary Dataset 1**). Among these, DMRs showing gain of methylation were predominant, compared to DMRs corresponding to loss of methylation (257 “hyper-DMRs”; 175 “hypo-DMRs”; **Figure 2B**). In either hyper- or hypo-DMRs, the median differential levels of DNA methylation in the majority of DMRs reached more than 10% (**Figure 2B**).

Even if there was a limited number of regions altered in their DNA-methylation levels, at least in the repertoire of the captured *loci*, our results suggest that DNA methylation is rapidly and locally redistributed in the cortices exposed to PAE. We performed hypergeometric tests either on all DMRs, or by separating hypo-DMRs and hyper-DMRs and observed that genomic regions were not randomly altered by PAE in their DNA methylation status. Indeed, brain active enhancers, characterized by the H3K27ac histone mark, were significantly over-represented among the DMRs identified in the capture, (345 on 432 DMRs *i.e*., 79.86% of DMRs; hypergeometric test, p-val < 0.05. **Figure 2C,D**; **Figure 3**). In contrast, promoters and other regions of interest, which represent respectively 134 and 35 DMRs, were not significantly affected by the binge drinking stress (**Figure 2B,C**).

**Figure 3.**
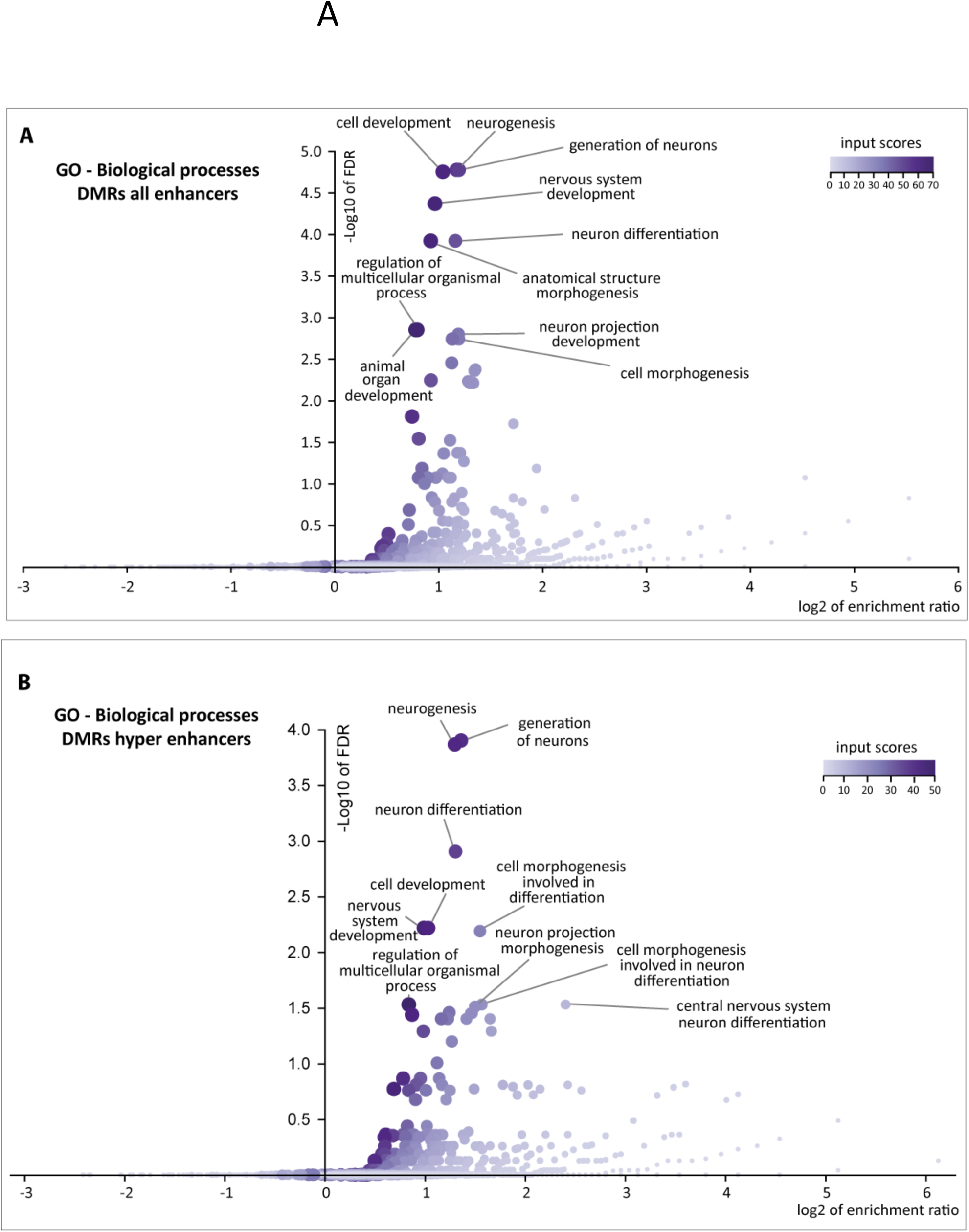
Gene ontology (GO) analyses performed on genes associated to DMRs that are observed into (or close to) active enhancers. **Volcano plots showing statistically relevant GO biological processes** for genes that are associated to **(A)** all DMRs (corresponding to either gain or loss of methylation) **(B)** to DMRs with gain of methylation, only, and observed into or close to active enhancers. The distance between DMRs and active enhancers regions is less than 500 bp. The size and share of color of each dot is proportional to the number of overlap with one given region category (Liao et al., 2019). The genes associated to the ten most relevant GO Biological Processes of these analyses are described in **Table 1** (for GO performed on all DMRs observed in active enhancers, related to Figure 2D) and **Table 2** (for GO performed only on regions that gain methylation upon PAE in active enhancers). These data were obtained using *Web-based GEne SeT AnaLysis Toolkit* (WebGestalt; Liao et al., 2019).

To identify potential binding sites of transcription factors within the DNA sequences of enhancers located in DMRs, we explored the presence of DNA sequence motifs using *findMotifsGenome* (HOMER). We could not observe any over-representation of specific motifs. This result suggests that there are no sub-categories of enhancers (*i.e.* characterized by specific transcription factors), that would be more severely affected by PAE than others. Since these enhancers belong to an active repertoire in the mouse brain, we then performed a Gene Ontology (GO) analysis on all the enhancers containing DMRs, in order to unveil specific Biological Processes potentially rapidly altered by PAE. We observed that a significant number of genes associated to these enhancers played a role in neurogenesis, generation of neurons, cell development, nervous system development, and neuron differentiation (**Figure 3A**; **Table 1**). When considered separately, enhancers with a decrease in methylation levels upon PAE were not significantly associated to a specific Biological Process. In contrast, hypermethylated enhancers were predominantly associated with genes that are involved in the above-cited Biological Processes (**Figure 3B**, **Table 2**).

**Table 1.**
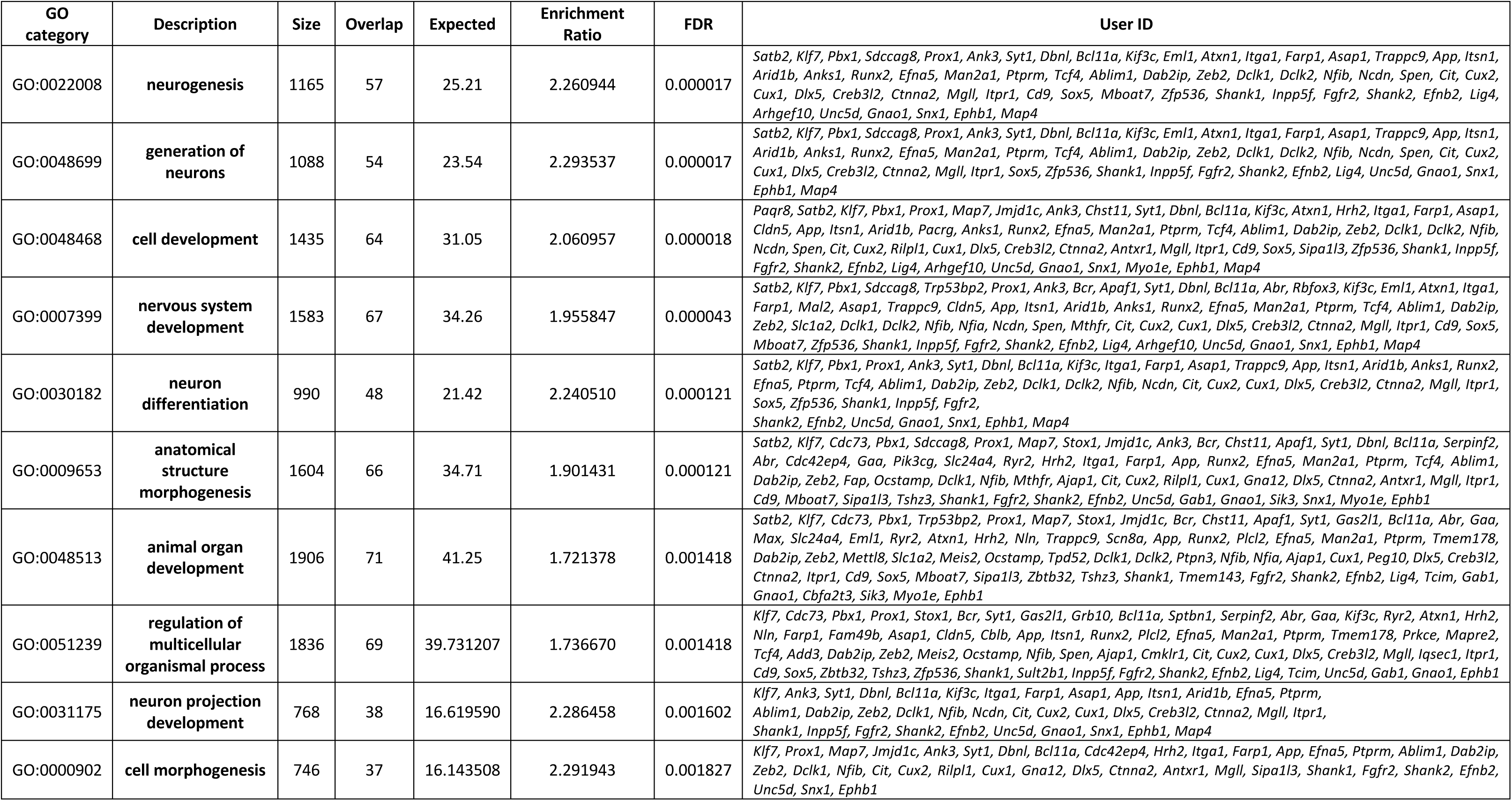
GO results for genes associated to all DMRs (Hypo and Hypermethylation) that are localized in active enhancers (H3K27ac)

**Table 2.**
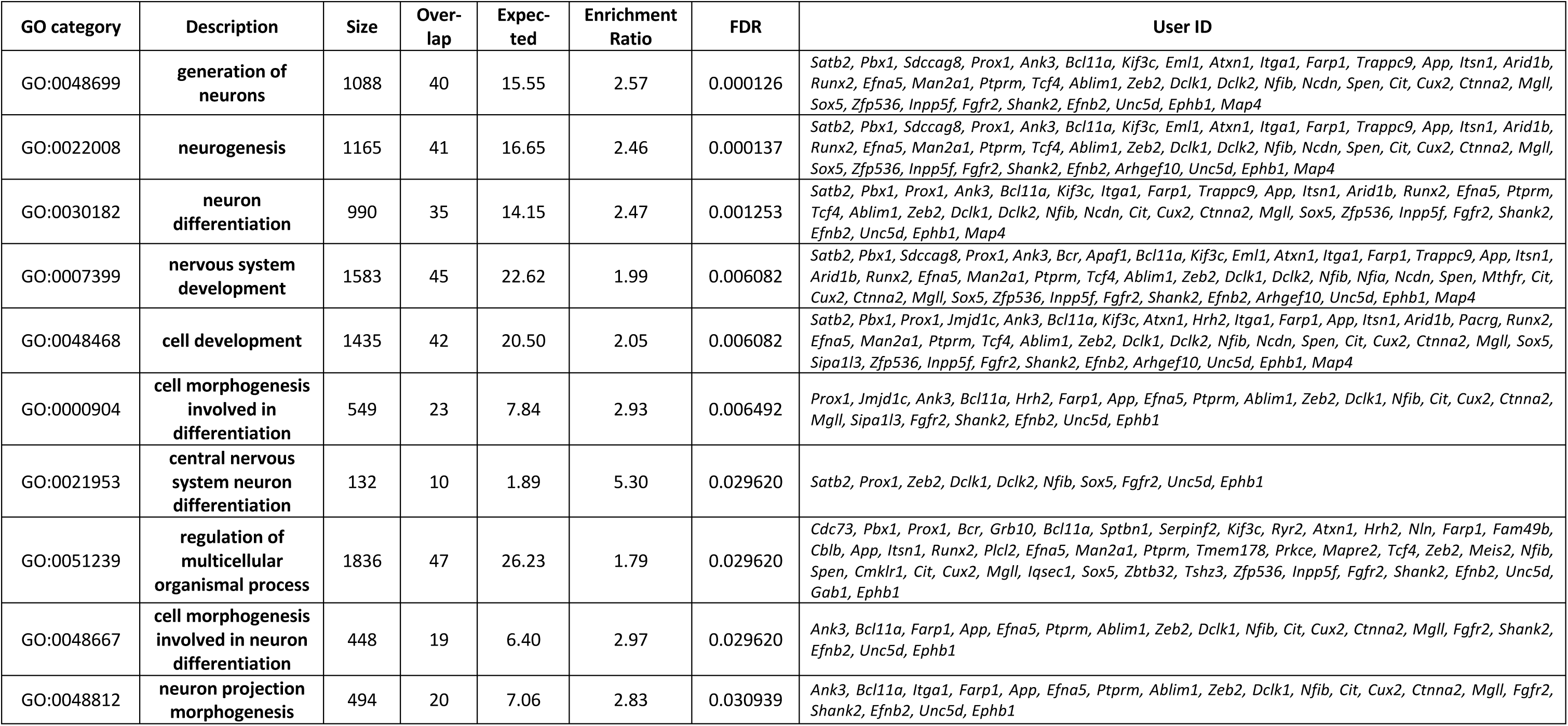
GO results for genes associated to Hypermethylated regions, observed upon EPA, that are localized in active enhancers (H3K27ac)

### Regions showing chromatin remodelling during physiological brain development are significantly associated with PAE-induced DNA methylation events

The striking enrichment of DMRs in enhancer regions, which were specifically active in the brain, let us think that these DNA methylation perturbations could occur in regions whose chromatin accessibility was undergoing changes at cortical developmental stages around the PAE exposure. We thus both investigated the changes of global chromatin accessibility and gene expression dynamics throughout the physiological mouse forebrain development around E14.5 to E16.5. For this, we used a bioinformatics workflow, which allows for pairwise comparison of successive developmental stages, to analyse public *ENCODE* ATAC-seq and RNA-seq datasets (**Supplementary Figure 2A**, and **Materials and Methods**, **Supplementary Data notebook 2** and **3**, **Supplementary Tables 4 and 5**). Our results are described in **Supplementary Dataset 1**. The number of identified regions or genes in each dataset is indicated in **Supplementary Figure 5A** (for quality control of these data, see **Supplementary Figure 5B,C**).

We observed DMR enrichment in regions whose chromatin accessibility was changing around the PAE time-window, which concerned regions that gained chromatin accessibility between E14.5 and E15.5, in physiological conditions (**Figure 4A,B**; **Supplementary Data notebook 4**, **Supplementary. Dataset 2**). This represented 5 regions out of 313 regions of the capture repertoire, among 1,442 total regions in the differentially opened regions (DOR) of the original ENCODE dataset (please see the legend of **Figure 5** for normalization details taking into account the constraints of the capture approach). Interestingly, these 5 regions corresponded to active enhancers and were associated with with GO terms of importance for neurodevelopment and brain function: for example, cell motility and cell migration (*Elmo1*), synaptic signalling (*Plcl2*), regulation of multicellular organization process and cytoskeleton protein (Mapre2), and cell development, cell biogenesis, plasma membrane part, abnormal mouse morphology (*Antxr*). In addition, our analysis of the RNA-seq ENCODE dataset (**Figure 4**) revealed that DMRs were significantly associated with genes that were upregulated between E14.5 and E15.5, and between E15.5 and E16.6 under physiological conditions (**Figure 4D**; for quality control of these data, see **Supplementary Figure 6A,B**).

**Figure 4.**
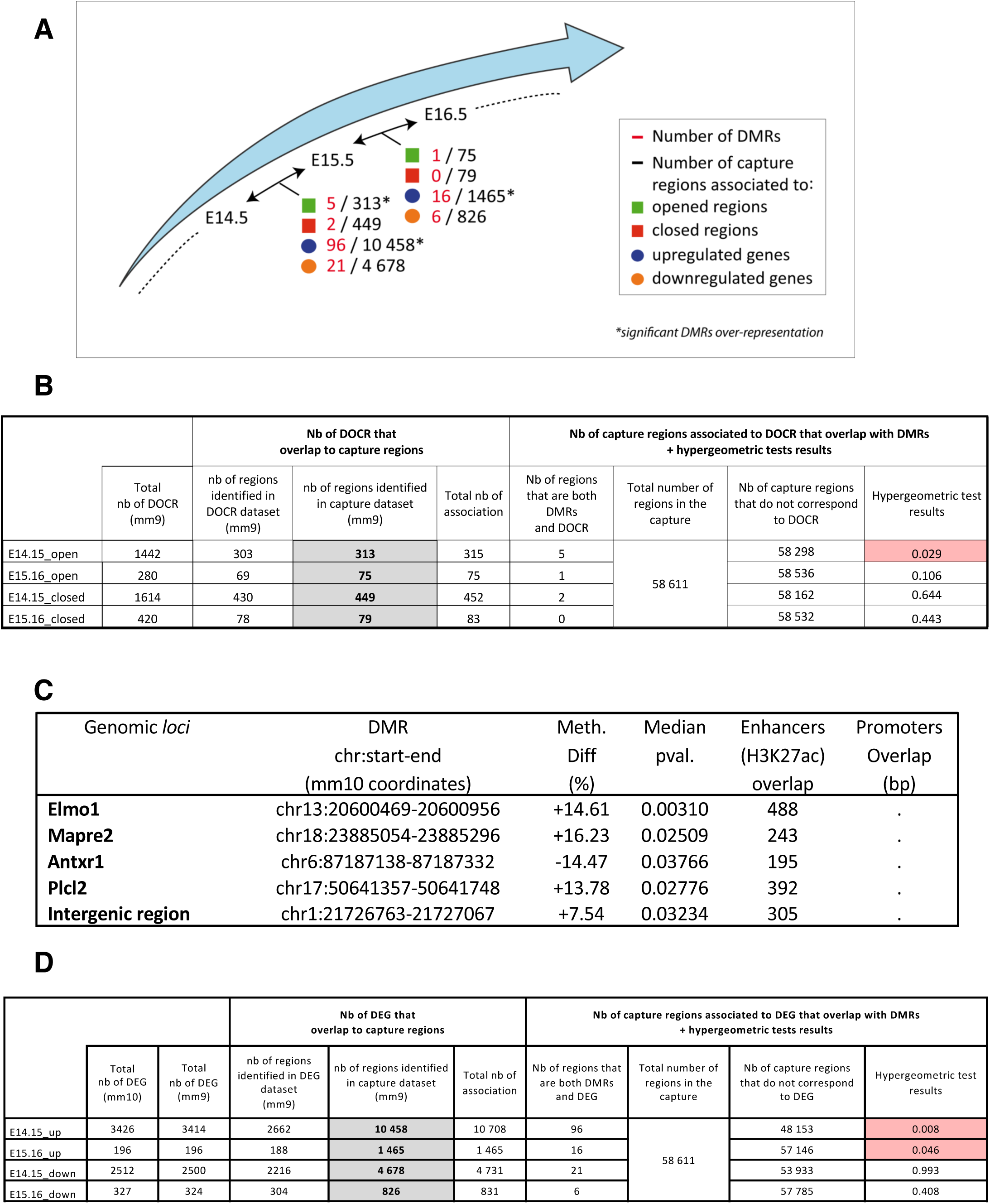
Immediate DMRs significantly fall into genomic regions that are remodeled in terms of chromatin accessibility or gene expression, during physiological brain development. **(A) *Number of capture regions that are associated to differentially opened or closed regions (DOCRs) or differentially expressed genes (DEGs) observed in the developing brain, under physiological conditions.*** DOCRs and DEGs are obtained in a pairwise-comparison manner of two successive developmental stages. Only DOCRs and promoters of DEGs strictly overlapping capture regions are taken into account (numbers indicated in black). Among these, DMRs strictly overlapping DOCRs/DEGs are considered for the analyses and their number of these regions is indicated in red. Asterisks indicated statistically significant results, *i.e.* when DMRs are found significantly over-represented (hypergeometric tests, see (**B**)). Total numbers of DOCRs and DEGs found during physiological brain development are indicated in **Supplementary Figure 5A** (*i.e.* independently of their presence in capture regions). Number of DOCR / DEG in **Supplementary Figure 5A** could be lower than the number indicated here, because a given DOCR or DEG could be represented by several capture regions. DOCR and DEG were identified using *edgeR* (using no logFC threshold but Benjamini-Hochberg p-value adjustment was performed and level of controlled false-positive rate was set to 0.05). **(B) *Hypergeometric test data showing the identification of over-represented DMRs among differentially opened or closed regions (DOCRs)*.** Values highlighted in red are significant. Tests compare opened regions between two successive developmental stages: E14.15 *versus* E15.5 and E15.5 *versus* E16. The younger developmental stage is taken as a reference; *i.e* : “E14.15_open” define regions that are more open at E15.5 compared to E14.5). Hypergeometric tests were performed using the number of methylome capture regions associated to DOCRs (values highlighted in grey). When comparisons between datasets regions are done, only regions with an overlap are taken into account. **(C) *DMRs significantly associated to genomic* regions *that are more opened between E14.5 and E15.5, under physiological conditions***. *Meth.diff:* differential methylation (%) observed between control (PBS-treated) and EtOH-treated groups. A negative percentage indicates a loss of methylation. *Median p-val.*: median of p-values observed for each relevant isolated CpG located in the DMRs (p-value < 0.07, see **Materials and Methods**). **(D) *Hypergeometric test data showing the identification of over-represented DMRs associated to differentially expressed genes (DEGs) observed during brain development, under physiological conditions***. As in (B).

**Figure 5.**
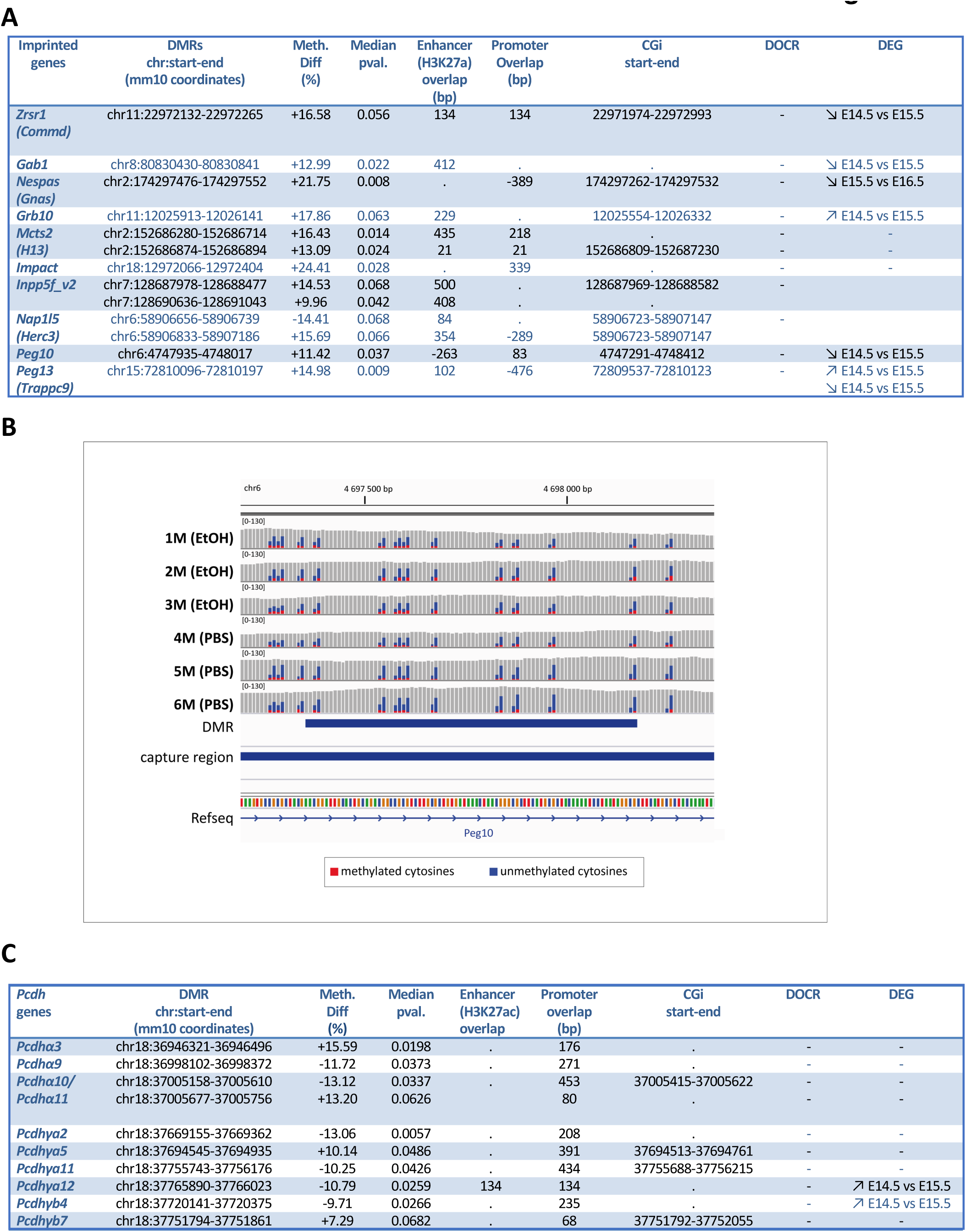
Mono-allelically expressed genes presenting DMR immediately after PAE. **(A) *Imprinted genes associated to immediate DMRs upon PAE.*** Overlap between “PAE-affected” imprinted genes and capture regions corresponds to either strict overlap or to close vicinity (distance < 500bp). Negative values correspond to DMRs located close to capture regions with no overlap. *Median pval.*: median of p-values for each relevant CpG (with pval < 0.07) that composes the DMRs. **(B) *Illustration of a DMR observed in* Peg10 *imprinted gene.*** Visualization of deduplicated read coverage of EtOH- and PBS-treated samples. Genomic region containing a DMR with gain of methylation, located into the *Peg10* gene is shown (thin blue bar at the bottom). Methylated cytosines in CpG context are in red, unmethylated ones in blue. Capture region is also indicated (thick blue bar). This screenshot was obtained using IGV. Scale is the same for all samples. **(C) *Protocadherin genes that are associated to immediate DMRs upon PAE*.** As in (A).

### Immediate DNA-methylation and concomitant gene expression changes affect mono-allelically expressed gene upon PAE

First, when exploring CpG islands (CGis), which are relevant targets of DNA methylation and its potential perturbations (Saxonov et al., 2006), we found that only 36 out of the 432 identified DMRs were located into CGis. However, a quarter of these DMRs (9 out of 36; **Figure 5A,B**), significantly, corresponded to imprinted genes, which are tightly regulated by DNA methylation (Perez et al., 2016). As a whole, 13 DMRs were associated to 10 distinct imprinted genes in our analysis (9 DMRs in CGis, 4 DMRs not associated with CGis; **Figure 5A,B**). Moreover, these DMRs were located into the imprinting control regions (ICRs), which govern the expression of the imprinted locus (**Table 3**). This statistically significant over-representation DMR associated to imprinted genes among DMRs was confirmed by a hypergeometric test (see details in **Materials and Methods**). Interestingly, a majority of DMRs associated to imprinted genes were located within the promoter regions (8/13) and/or active enhancers in the developing brain (11/13, **Figure 5A**).

**Table 3.**
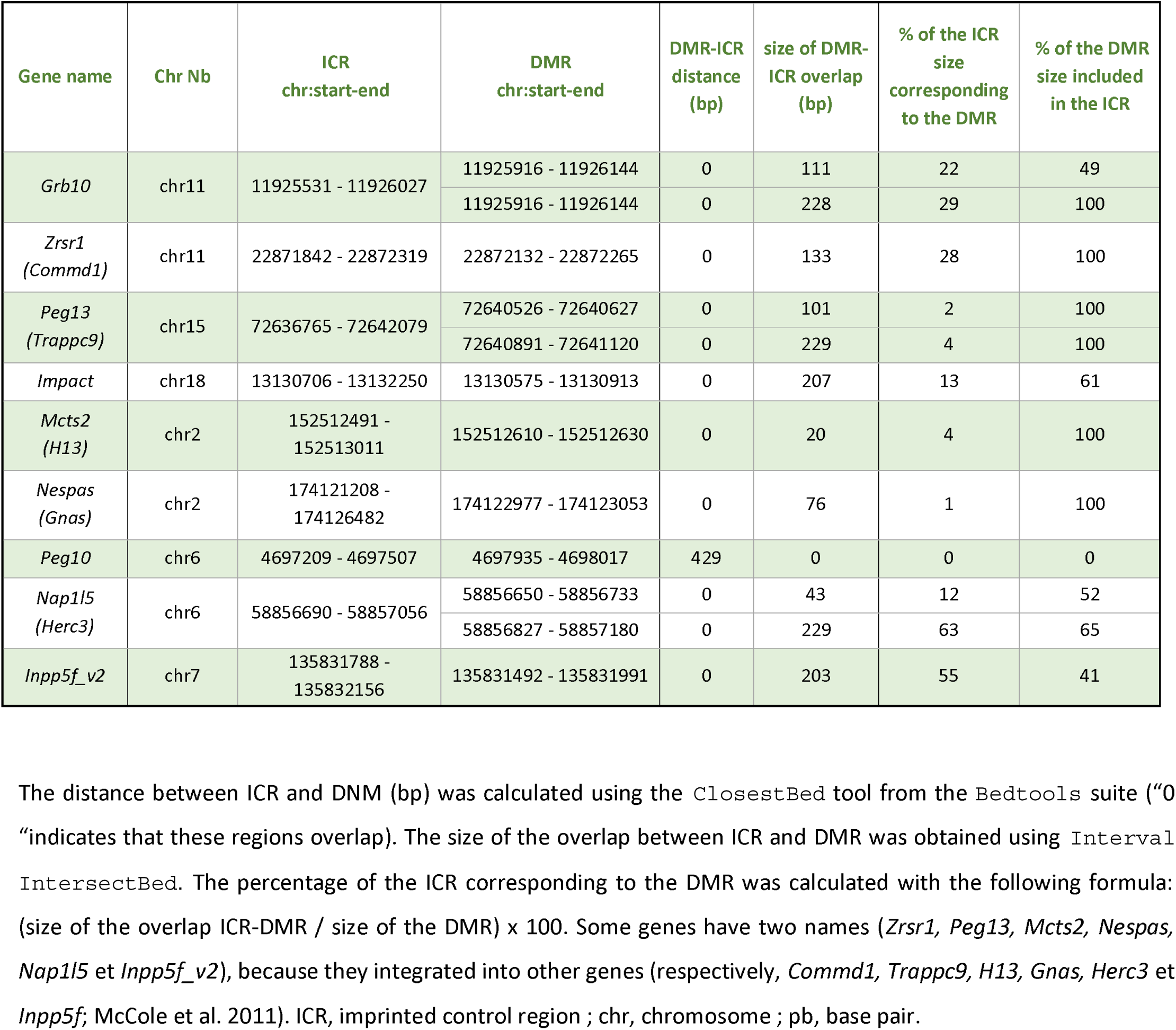
Imprinting control region (ICR) containing DMR in response to PAE.

| Gene name | Chr Nb | ICR<br>chr:start-end | DMR<br>chr:start-end | DMR-ICR<br>distance<br>(bp) | size of DMR-ICR overlap<br>(bp) | % of the ICR<br>size<br>corresponding<br>to the DMR | % of the DMR<br>size included<br>in the ICR |
| --- | --- | --- | --- | --- | --- | --- | --- |
| *Grb10* | chr11 | 11925531 - 11926027 | 11925916 - 11926144 | 0 | 111 | 22 | 49 |
|  |  |  | 11925916 - 11926144 | 0 | 228 | 29 | 100 |
| *Zrsr1*<br>(*Commd1*) | chr11 | 22871842 - 22872319 | 22872132 - 22872265 | 0 | 133 | 28 | 100 |
| *Peg13*<br>(*Trappc9*) | chr15 | 72636765 - 72642079 | 72640526 - 72640627 | 0 | 101 | 2 | 100 |
|  |  |  | 72640891 - 72641120 | 0 | 229 | 4 | 100 |
| *Impact* | chr18 | 13130706 - 13132250 | 13130575 - 13130913 | 0 | 207 | 13 | 61 |
| *Mcts2*<br>(*H13*) | chr2 | 152512491 -<br>152513011 | 152512610 - 152512630 | 0 | 20 | 4 | 100 |
| *Nespas*<br>(*Gnas*) | chr2 | 174121208 -<br>174126482 | 174122977 - 174123053 | 0 | 76 | 1 | 100 |
| *Peg10* | chr6 | 4697209 - 4697507 | 4697935 - 4698017 | 429 | 0 | 0 | 0 |
| *Nap1l5*<br>(*Herc3*) | chr6 | 58856690 - 58857056 | 58856650 - 58856733 | 0 | 43 | 12 | 52 |
|  |  |  | 58856827 - 58857180 | 0 | 229 | 63 | 65 |
| *Inpp5f_v2* | chr7 | 135831788 -<br>135832156 | 135831492 - 135831991 | 0 | 203 | 55 | 41 |

The distance between ICR and DNM (bp) was calculated using the `ClosestBed` tool from the `Bedtools` suite (“0” indicates that these regions overlap). The size of the overlap between ICR and DMR was obtained using `Interval IntersectBed`. The percentage of the ICR corresponding to the DMR was calculated with the following formula: (size of the overlap ICR-DMR / size of the DMR) x 100. Some genes have two names (*Zrsr1*, *Peg13*, *Mcts2*, *Nespas*, *Nap1l5* et *Inpp5f\_v2*), because they integrated into other genes (respectively, *Commd1*, *Trappc9*, *H13*, *Gnas*, *Herc3* et *Inpp5f*; McCole et al. 2011). ICR, imprinted control region ; chr, chromosome ; pb, base pair.

In addition, we found that a significative number of early DMRs were associated to clustered protocadherin genes (*Pcdhs*). Clustered *Pcdhs* are also mono-allelically expressed, but in a random manner, and are individually regulated by DNA methylation (Phillips et al., 2017). Namely, we identified 10 DMRs in 9 distinct *Pcdh*, from both α- and γ-protocadherins clusters (**Figure 5C**). For 4 *Pcdhs*, DMRs observed immediately upon PAE are also located in a CpGi (**Figure 5C**).

The tight regulation of the monoallelic expression of imprinted genes and *protocadherins* by DNA methylation is crucial for neurodevelopment and normal brain functions (Davies et al., 2008). For example, clustered *Pcdhs* play important roles in the modulation of dendrites arborisation and synaptogenesis or limitation of autapse formation through cell-cell recognition (Light and Jontes, 2017; Phillips et al., 2017; reviewed in El Hajj et al., 2016; Matsunaga et al., 2017; Molumby et al., 2017; Yamagata et al., 2018). The early alteration of their methylation levels by PAE could thus potentially affect their expression with potential defects typical of FASD. We thus investigated whether the establishment of DMRs in these genes were concomitant with changes in their expression levels. Using RT-qPCR, we examined the expression of four imprinted genes (*Inpp5f_v2, Nap1l5, Peg13,* and *Zrsr1*), three genes of protocadherin clusters (*Pcdhα3, Pcdhα9,* and *Pcdhγa2*). We also included three genes in our analysis, whose mutations in human lead to neurodevelopment disorders and which were also associated with DMRs with similar percentages of methylation differences (**Supplementary Figure 7A**): *Mid1*, associated to the Opitz BBB/G syndrome reviewed in (Winter et al., 2016) and *Shank2* and *Auts2* both linked to autism spectrum disorders (ASD; reviewed in Bourgeron, 2015 and Pang et al., 2021), with which FASD share overlapping traits (Lussier et al., 2021; Popova et al., 2016). We showed that these genes which were associated with immediate DMRs exhibited concomitant disturbances of their expression in the prenatal cortex (**Figure 6**).

**Figure 6.**
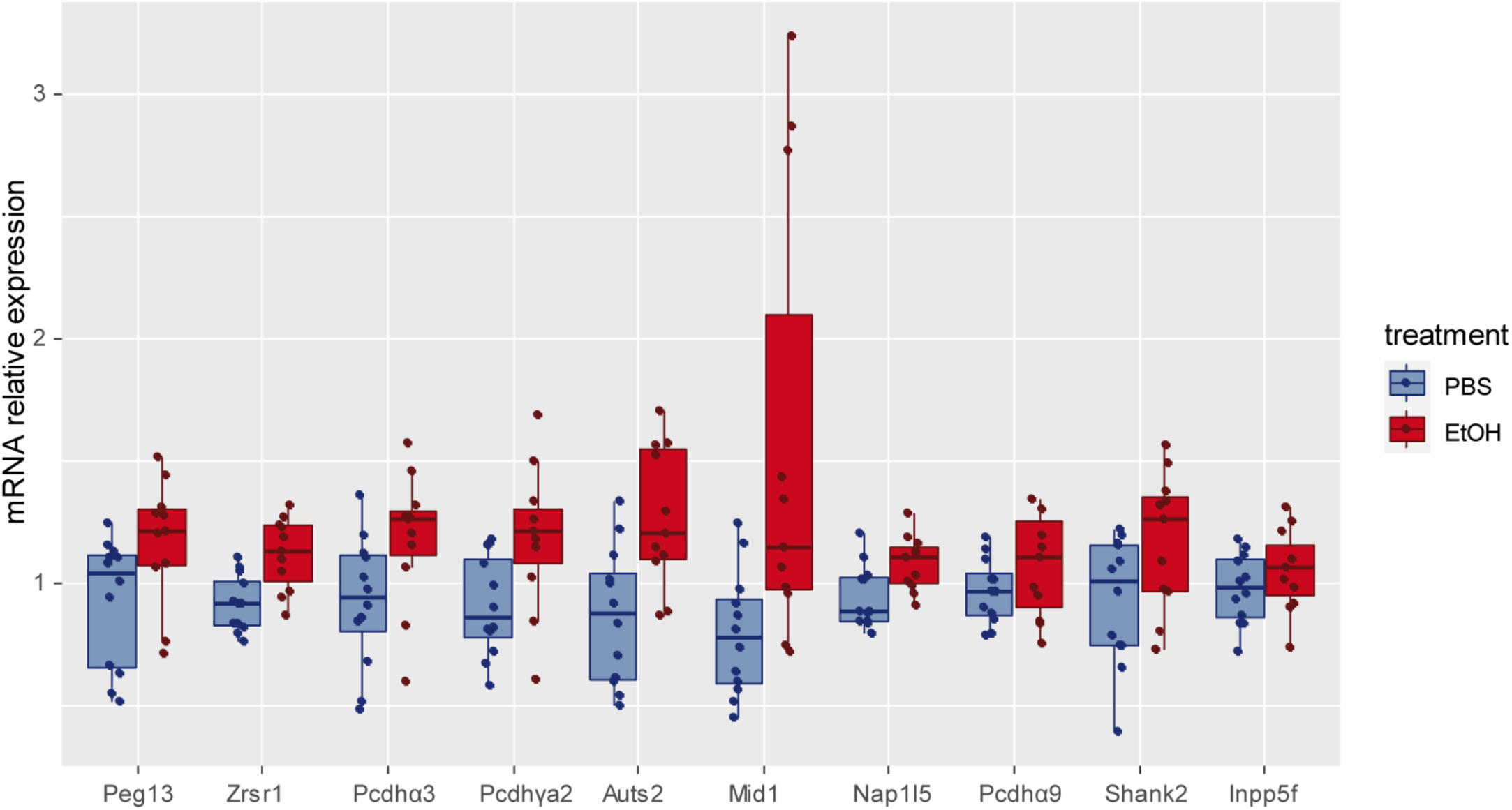
PAE-induced alterations of gene expression are concomitant with immediate perturbations of DNA methylation. ***Quantitative RT-PCR of 10 genes associated to immediate DMRs.*** Boxplots showing relative mRNA levels from E16.5 embryonic cortices upon PBS- or EtOH-treatment. Each dot represents calibrated and normalized relative gene expression (CNRQ) value of each sample. mRNA levels are normalized to the expression of *Tpt1* and *Hprt1* genes. Results are generated using n=12 PBS samples and n=11 EtOH samples, from 3 distinct litters in each condition. Both males and females are considered together, since no sexual dimorphism is detected in terms of the expression of the studied genes (**Supplementary Figure 7B,C**). Significance differences between PBS- and EtOH-treated samples were determined using non-parametric Mann-Whitney test. To take into account multiple tests, p-values were corrected into adjusted p-values (padj.), using Benjamini-Hochberg correction. Asterisks (*) denote significant differences between PBS- and EtOH-treated samples (when padj. < 0.05). ns, statistically non-significant results.

### Immediate impacts of PAE on the transcriptome

To get a broader picture of the immediate transcriptomic changes induced by PAE in the prenatal cortex, and putative impact on cortical development, we performed RNA-Seq analysis of control or PAE fetal cortices (8 for each condition; **Supplementary Figures 2A** and **Statistical report of project Sartools-edgeR-RNAseq-PBSvsEtOH_report_note.html**). Multidimensional scaling analysis indicated that control (PBS) samples were grouped together, and, thus, more similar between each other, compared to the EtOH samples (**Figure 7A**). In contrast, EtOH-exposed samples were more distant one from each other: EtOH3 and EtOH4 samples were similar to the control ones, whereas other EtOH samples were distant from the cluster of control samples (**Figure 7A**). This likely reflects the variability of the response to PAE among individual fetal cortices, knowing that embryos from the same litter do not systematically cluster (**Figure 7A**). We found that a total of 4808 genes were significantly dysregulated in their expression (adjusted p-values < 0.05), among which 2598 were up- and 2210 downregulated (see **Sartools-edgeR-RNAseq-PBSvsEtOH_report_note.html** and **Supplementary Table 6**; these differentially expressed genes (DEGs) were hereafter termed UP DEGs and DOWN DEGs. Volcano and MA plot visualization of the data indicated that a majority of genes exhibited limited expression alterations in terms of fold-change intensity (**Figure 7B,C**) and that only 11 DOWN DEGs and UP DEGS were found to have a log2 FC > or < 1, respectively (highlighted **in Supplementary Table 6**). A large majority of DEGs (4732) were not found associated with a DMR. We found 40 DEGs that were both associated with a DMR and either localized in a region which, in unstressed cortices, gain chromatin accessibility between E14.5 and 515.5 or 15.5 and EE16.5 (see **Figure 4**) or whose expression increased during one of these time-windows. Five imprinted genes were identified in our RNA-seq dataset: three that were upregulated and in which a DMR was located in the ICR (*Gnas*; *Herc3*, and *Impact*), one that was downregulated and whose ICR contained a DMR (*Peg13*(*TrappC9*), and one downregulated (Commd1) but which was not associated with a DMR (**Supplementary Table 7**). GO-term analysis indicated that DOWN DEGs fell into biological processes and cellular components linked to: i) ribonucleoprotein complex and ribosome biogenesis, rRNA, nRNA, and mRNA maturation and processing, including splicing, and into categories associated with mitochondrial translation, mitochondrial protein complexes, respiration, and oxidative phosphorylation, vesicle exocytosis and transport, as well as synaptic vesicles (p-values comprised between 2.98e-62 and 2.00e-16) (**Figure 8A**; **Supplementary Table 8**). Whereas the numbers of DOWN and UP genes were similar, DOWN DEGs were enriched in GO-terms with lowest adjusted p-values than UP DEGs, which suggested that PAE immediate impacts on the transcriptome are majorly mediated by downregulating gene expression (**Figures 8A** and **9A**). UP DEGs were associated to (i) the Wnt pathway, and SMAD binding, regionalization and forebrain development, transmembrane transporters, (ii) repression of transcription, polycomb-group proteins, Polycomb Repressive Complex 1 (PRC1), and (iii) hallmarks of stress responses, like cytoplasmic ribonucleoprotein granules, P-bodies, stress granules, and centriolar satellites (**Figures 9A**). In addition, among the DOWN DEGs identified in PAE-exposed samples, gene-set enrichment analysis (GSEA) confirmed that a majority of the genes present in a given GO-term was significantly impacted in terms of dysregulation of expression. This was the case for cellular and mitochondrial protein synthesis and for respiration: notably, “ligase activity, forming carbon-oxygen bonds” (i.e. amino-acid tRNA ligase), “tRNA-binding”, “ribosome biogenesis”, “rRNA processing”, and mitochondrial translation, and “NADH Dehydrogenase complex assembly” (**Figure 8B,C** and **Supplementary Figure 8A,B**; **Supplementary Table 9** and **10**). Likewise, UP DEGs globally fell into a more restricted number of GSEA categories: genes involved in the negative regulation of BMP signaling (for biological processes), in transforming growth factor beta (TGF)-binding (for molecular functions), and PRC1 (for cellular components) (**Figure 9B,C and Supplementary Table 9** & **10**).

**Figure 7:**
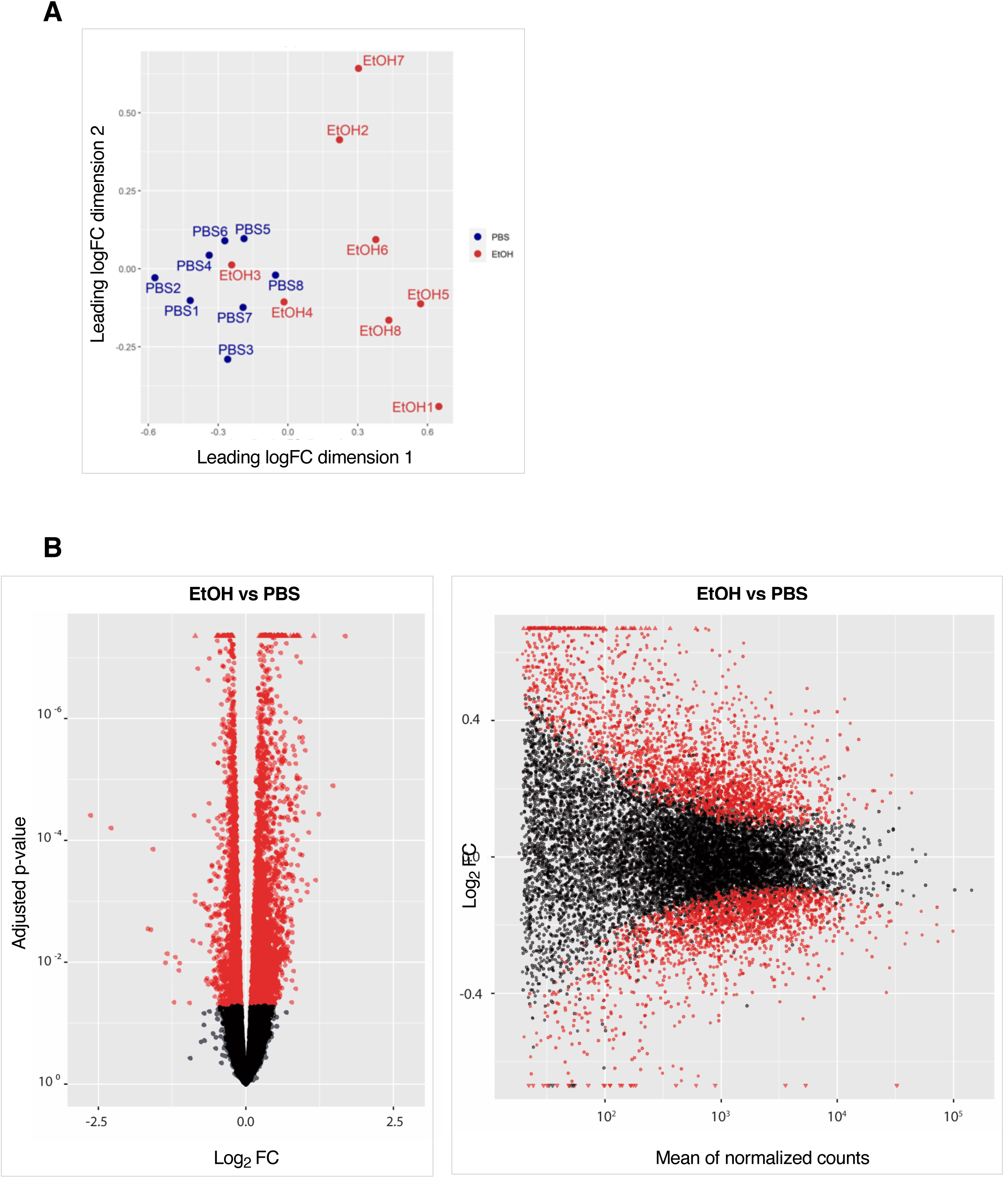
Immediate impacts of PAE on the transcriptome. **(A) *Multidimensional scaling plot (MDS) of the samples based on RNA-seq analysis*.** In blue control sample (PBS1 to 8). In red Ethanol treated samples (EtOH 1 to 8). The 1er dimension separates PBS from EtOH sample except samples EtOH 3 and 4 that are clustered with the control ones. While Control samples are quite similar, Ethanol treated samples are more variable **(B) Volcano plots showing all detected mRNAs** Left panel: Each dot represents a single gene with its differential gene expression level plotted on the x axis and statistical significance (pvaladj) on the y axis. Red dots represent significantly differentially expressed features (pvaladj < 0.05). 11 upregulated genes upon ethanol (right side of the plot) have a logFC>1. 11 downregulated genes upon ethanol (left side of the plot) have a logFC>1 Right panel: MA-plot representing the log ratio of differential expression as a function of the mean intensity for each feature. Red dots represent significantly differentially expressed features (pvaladj < 0.05). FC, fold change. Triangles correspond to features having a too low/high log2(FC) to be displayed on the plot.

**Figure 8:**
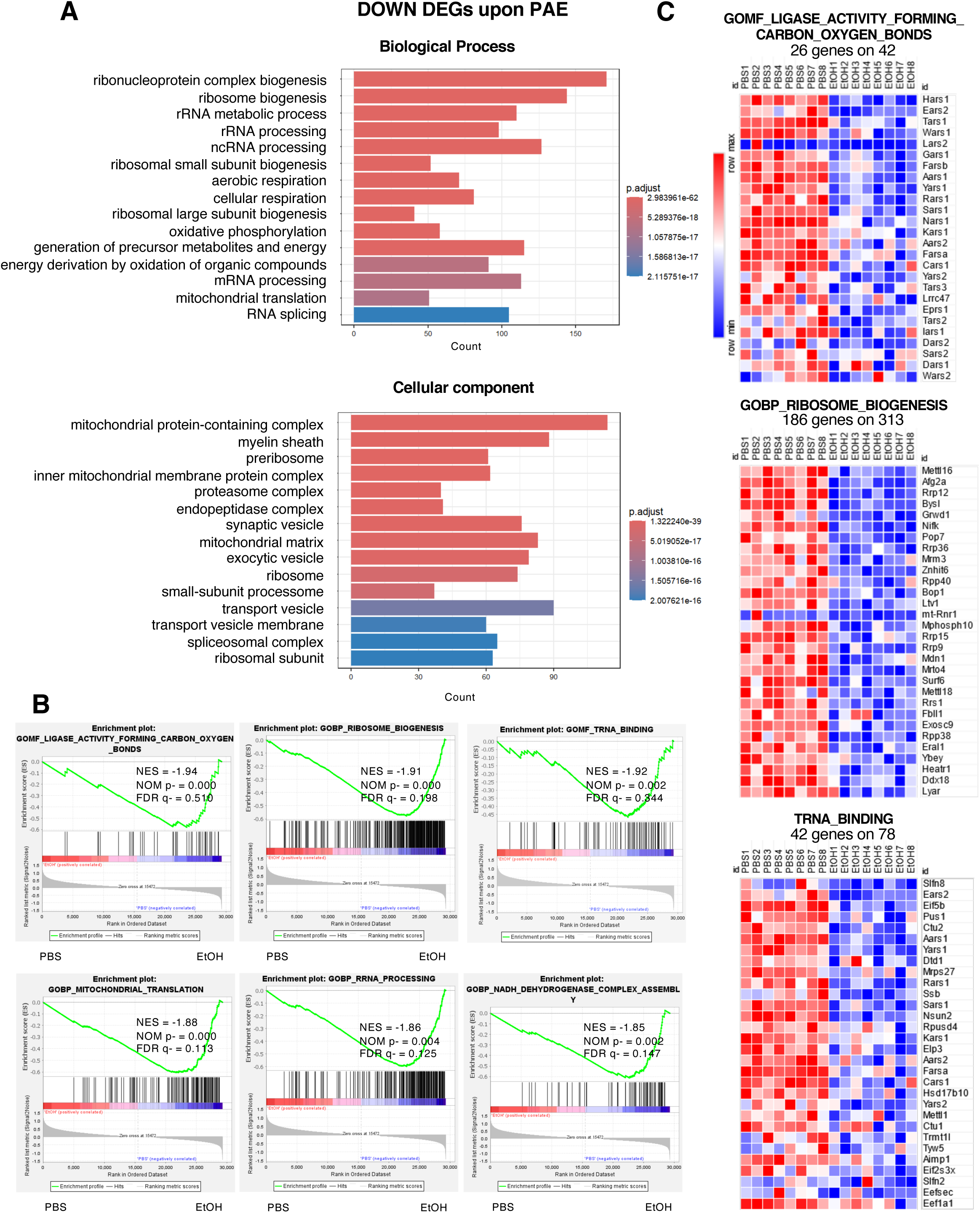
Gene ontology (GO) analysis of down regulated genes upon Ethanol treatment: **(A)** GO terms enriched in Biological process (BP) and cellular component (CC) categories in genes significantly downregulated in PAE samples. **(B)** GO enrichment plot of six selected GO-terms gene sets (GSEA, gene set enrichment analysis). The top portion of plots show the enrichment scores for each gene, and the bottom portion shows the ranked genes. Y-axis: ranking metric, X-axis: individual ranks for all genes. The normalized enrichment score (NES), the false discovery rates (FDR) and the nominal p-value (NOM p) are indicated for each gene set. Each bar at the bottom of plot represents a member gene of the respective gene set. **(C)** Heatmaps representing the level of expression in each sample of the top30 downregulated genes of selected GO gene sets. The number of DEGs involved and the number of genes in the selected gene set are indicated.

**Figure 9:**
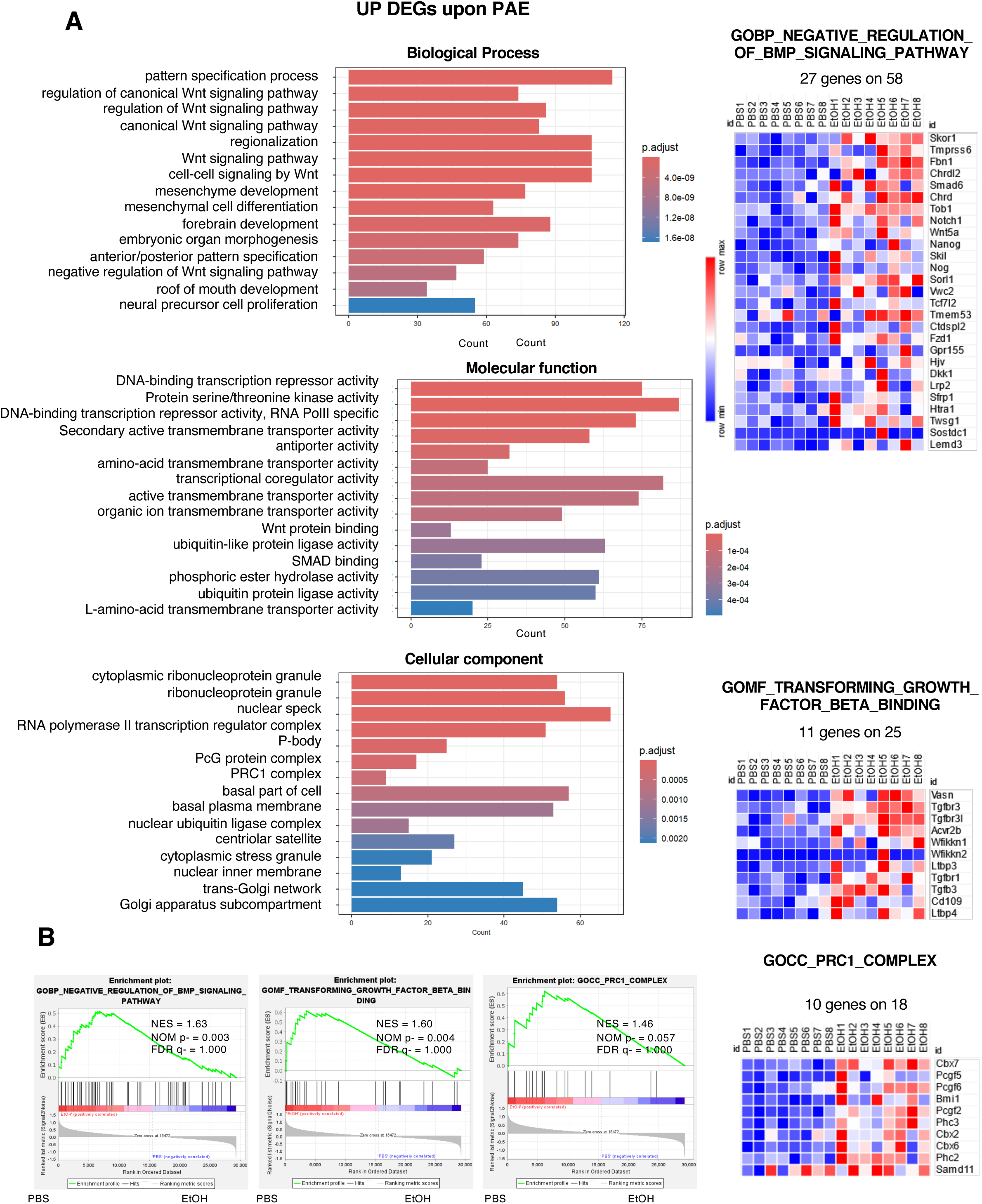
Gene ontology (GO) analysis of down regulated genes upon Ethanol treatment: **(A)** GO terms enriched in Biological process (BP) category in significantly upregulated genes in in PAE samples. **(B)** GO enrichment plot of three selected GO-terms gene sets (GSEA, gene set enrichment analysis). The top portion of plots show the enrichment scores for each gene, and the bottom portion shows the ranked genes. Y-axis: ranking metric, X-axis: individual ranks for all genes. The normalized enrichment score (NES), the false discovery rates (FDR) and the nominal p-value (NOM p) are indicated for each gene set. Each bar at the bottom of plot represents a member gene of the respective gene set. **(C)** Heatmaps representing the level of expression in each sample of the top30 upregulated genes of selected GO gene sets. The number of DEGs involved and the number of genes in the selected gene set are indicated.

Modest transcriptional changes, as those observed here (**Figure 7B**), that affect a wide number of genes involved in the same pathways are susceptible to impact neurodevelopment by their coordinated modulation. To gain deeper insight into the potential coordinated impact of proteins encoded by DEGs and their potential involvement in protein–protein interaction networks, we conducted network analyses based in protein-protein interactions (STRING, v1.7.0 https://string-db.org/) and performed visualization by using Cytoscape (v. 3.9.1) (Figure 10 and 11; Supplementary Table 12). When DOWN DEGs were considered, we identified 7 clusters of protein-protein interactions. Among the fourth containing the highest numbers of genes, Cluster 1 corresponded to “ribonucleoprotein complex” and “ribosome biogenesis”, and “RNA metabolism and processing”, in particular. Cluster 2 was linked to “spliceosome” and “splicing”, Cluster 3 gathered “respiratory electron transport/respirasome”, and “inner mitochondrial membrane protein complex”, “electron transport chain”, and Cluster 4 was associated to “mitochondrial translation” (**Figure 10**). Based on the population of UP DEGs, we detected 21 clusters of protein-protein interactions (**Figure 11**). In particular, Cluster 1 contains co-upregulated genes belonging to PRC1. Cluster 2 is centered around the transcriptional co-activator p300. Cluster 3 correspond to the TGF - SMAD - BMP signaling and Cluster 4 to the NF-kB pathways.

**Figure 10:**
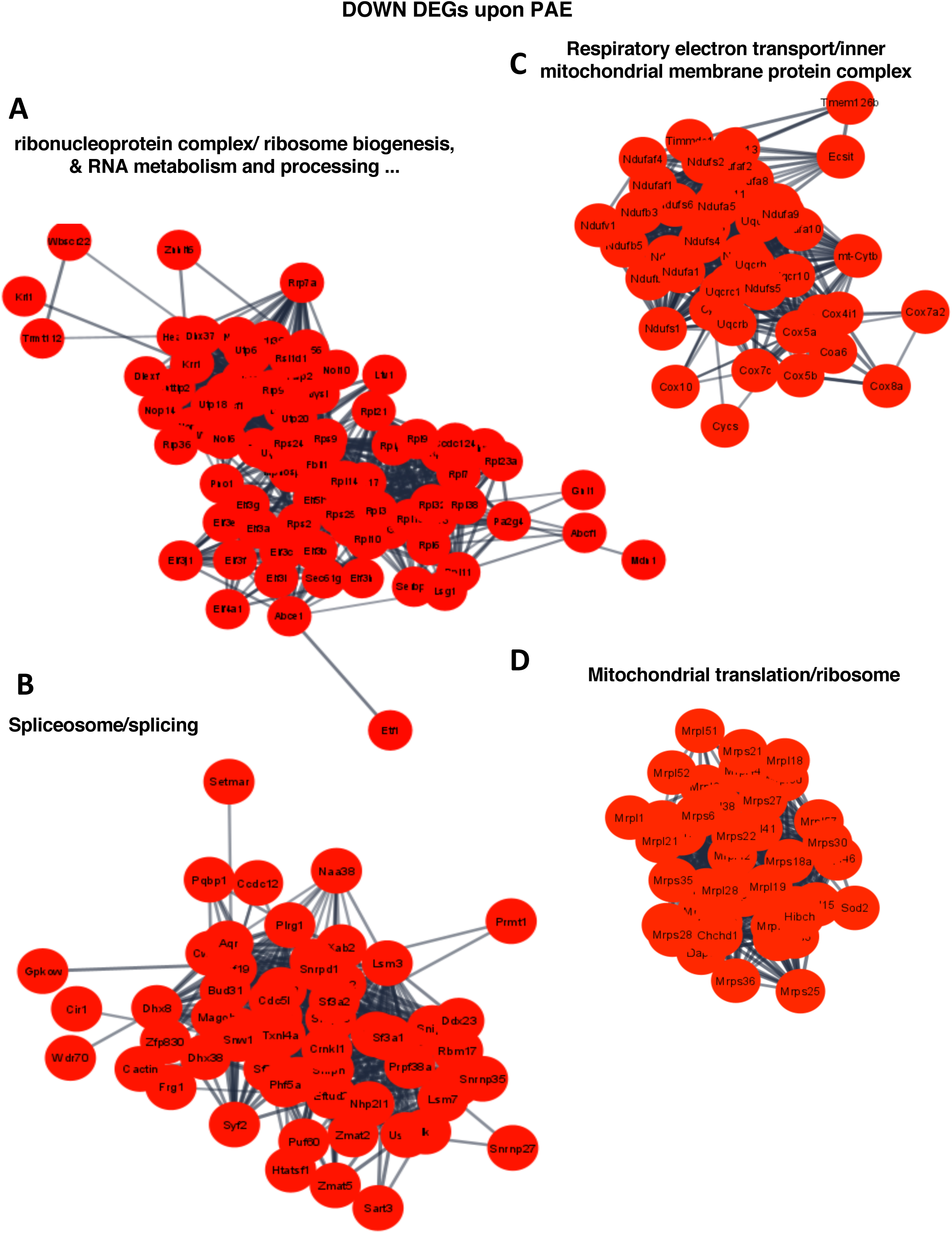
Functional protein interaction analysis using STRING software for proteins encoded by the downregulated genes identified upon Ethanol treatment. The four most significant clusters are shown. **(A)** Cluster 1: Ribonucleoprotein complex/ribosome biogenesis and RNA metabolism and processing involving 84 proteins presenting 1014 potential interactions between them. **(B)** Cluster 2: Spliceosome/slicing complex involving 58 proteins presenting 527 potential interactions between them. **(C)** Cluster 3: Respiratory electron transport/inner mitochondrial membrane protein complex involving 44 proteins presenting 543 potential interactions between them. **(D)** Cluster 4: Mitochondrial translation/ribosome involving 41 proteins presenting 528 potential interactions between them.

**Figure 11:**
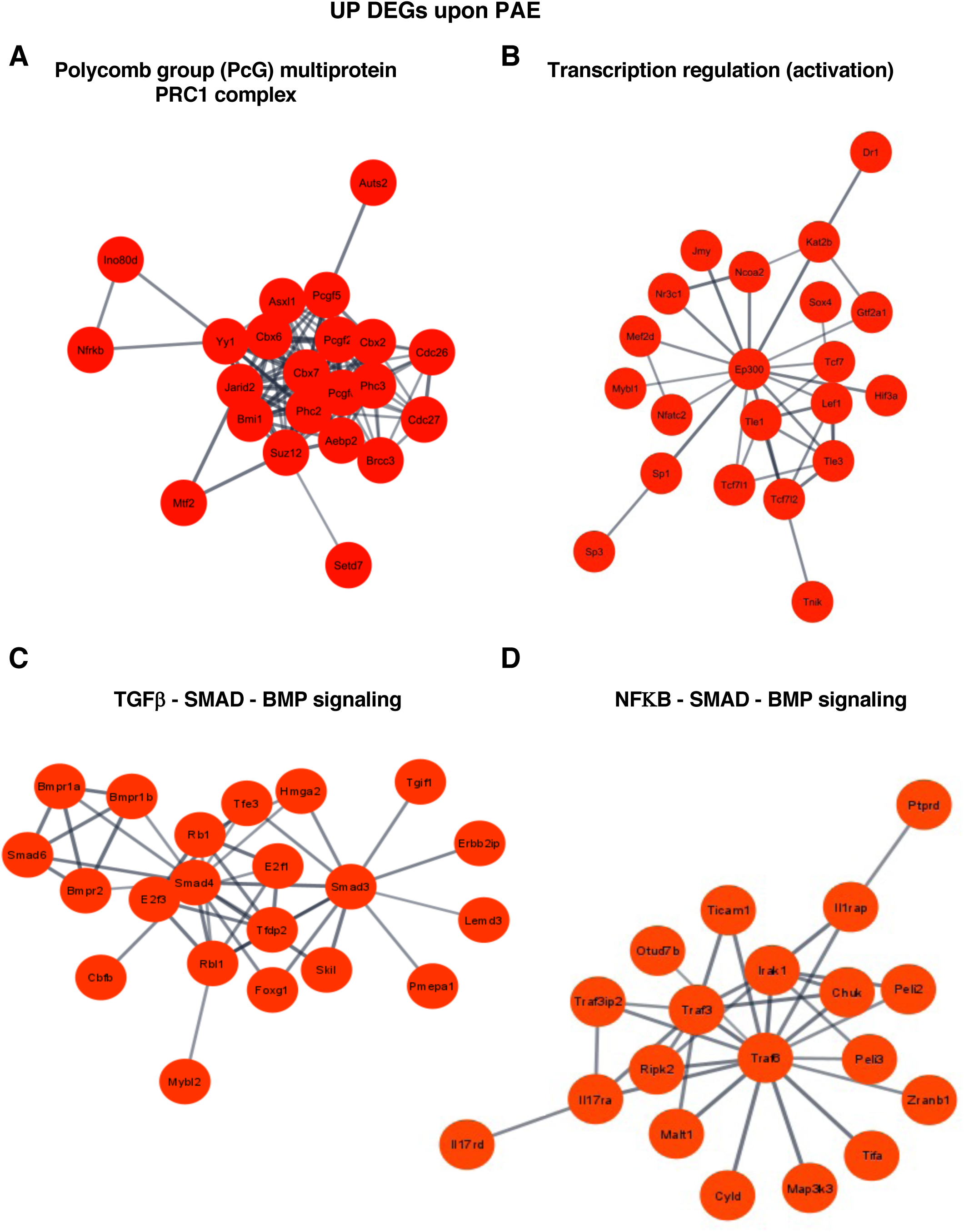
Functional protein interaction analysis using STRING software for proteins encoded by the upregulated genes identified upon Ethanol treatment. The four most significant clusters are shown. **(A)** Cluster 1: Polycomb group (PcG) multiprotein PRC1 complex involving 22 proteins presenting 119 potential interactions between them. **(B)** Cluster 2: TGFβ-BMP signaling complex involving 21 proteins presenting 33 potential interactions between them. **(C)** Cluster 3: Transcription regulation (activation) complex involving 21 proteins presenting 41 potential interactions between them. **(D)** Cluster 4: NFκB-SMAD-BMP signaling complex involving 19 proteins presenting 31 potential interactions between them.

Altogether, our GO, GSEA, and STRING analyses pointed to major downregulation of genes of importance for cytoplasmic and mitochondrial ribosome functions, mitochondrial translation, and respiration. Similarly, both GSEA and STRING analyses indicated that genes that are upregulated in a coordinated manner by PAE encode proteins that belong to an interaction network governing the TGF⍰- SMAD - BMP signaling pathway.

## DISCUSSION

The impact of PAE on the DNA methylome in the brain has been almost exclusively interrogated in the postnatal and adult brain, at long temporal distance from the time-window of exposure (Laufer et al., 2017) and the question whether perturbations of DNA methylation occur quickly after exposure has remained elusive. A recent study also investigated the effect of prenatal alcohol exposure during the preimplantation stages of the mouse embryo and, 10 days later, identified DMRs that interestingly affected genes important for neurodevelopment (Legault et al., 2021). To the best of our knowledge, our study represents the first analysis of the effect of PAE on the DNA methylome in the developing neocortex, addressing the question of the immediate deposition of aberrant DNA methylation marks, at very short temporal distance from the exposure.

### Statistical intra- and inter-group variability

We observed that the intra-group variability was limited (**Supplementary Figure 2D**). This is likely due to our sampling mode which aimed at minimizing inter-individual variability in the same group, by mixing half-cortices from four embryos from four different litters in the same NGS sample (**Supplementary Figure 1B**). However, we also observe that the triplicates of the EtOH-treated group show more differences between each other than do triplicates from the control group. Interestingly, this is also observed in another paradigm prenatal stress due to neuroinflammation between samples within the exposed group (Schang et al., 2018a), and more generally the exacerbated cell-to-cell variability provoked by PAE (Ishii et al., 2017). Indeed, this might reflect the immediate impact of cellular stress caused by ethanol exposure, and the disequilibrium, with some stochastic aspects, it imposes to cellular functions and molecular pathways, including the regulation of DNA methylation, before the system reaches a new equilibrium.

Alcohol exposure has multiple potential effects on DNA methylation in terms of availability of precursors of the methyl group (Hutson et al., 2012; Kleiber et al., 2014; Chater-Diehl et al., 2017), elevation of DNMT levels (Miozzo et al., 2018), or potential redistribution of DNMTs in the genome, which would explain the fact that gain and loss of methylation can be observed depending on the genomic region considered. Our statistical analyses suggested that inter-group variability (control versus PAE-treated samples) is limited, and that PAE therefore might not majorly reshape the DNA methylome – at least in the repertoire of captured regions (**Supplementary Figure 2D**). Three observations could explain these results: i) it could result from the stochasticity of PAE effects on DNA methylation, and consequently, due to the fact that we capture only the most frequent and robust alterations in DNA methylation; this would not mean that the other perturbations of DNA methylation in one individual couldn’t be meaningful in terms of clinical phenotypic consequences (Cobben et al., 2019). ii) Another possibility is that, in spite of pleiotropic alcohol impacts, the DNA methylome is globally unchanged after such a binge-drinking-like mode of PAE due to rapid protective mechanisms, which would be interesting to unravel. iii) Alternatively, this apparent robustness of the DNA methylome might only reflect the fact that other perturbations of DNA methylation could occur later in life, for example due to abnormal neuronal plasticity due to PAE, that could reshape the DNA methylome profile in the postnatal and adult brain (Guo et al., 2011). Nevertheless, studies that explored the DNA methylation patterns at temporal distances in animal models or FASD cohorts, using capture-like or genome-wide approaches also identified a limited number of DMRs (a few hundred in most studies, around 6600 promoter regions in one study; Laufer et al., 2013, 2016; Portales-Casamar et al., 2016; Lussier et al., 2018; Cobben et al., 2019). Whether these DMRs originate from early or late alterations of DNA methylation patterns, this means that the DNA methylome is altered in a limited number of regions, which does not mean that this could be detrimental for the brain.

### Detection of individual differentially methylated cytosines, shortly after PAE

We detect few individual cytosines showing statistically differential methylation after PAE. Two of them, hypermethylated, reside into genes, *KALRN* and *Tiam2*, that are involved in a pathway important for neurodevelopment: guanine nucleotide exchange factors. These genes participate to neuronal shape and polarity, axon growth and/or neuronal plasticity (Cahill et al., 2009; Honda et al., 2017). *KALRN* misexpression has been linked to neuropsychiatric disorders like schizophrenia and addiction, and this gene has been found, in genome-wide association studies, to be related to ADHD, and schizophrenia, all pathologies being relevant for FASD (reviewed in Remmers et al., 2014). Because the methylation of an individual CpG can affect gene expression (Xu et al., 2007), these immediate DMCs could have deleterious effects on neuronal function.

### Immediate DMRs are mostly found in brain active enhancers upon PAE

We have mainly focused on the identification and exploration of DMRs, which are, in general, more robustly associated to impacts on gene expression. We identified 432 DMRs, showing that DNA methylation is rapidly redistributed in the genome, shortly after PAE. We found that these immediate DMRs are not randomly distributed among the repertoire of capture regions: indeed, in contrast to promoters and other genomic regions, which are unaffected, enhancers that are known to be active in the adult brain are significantly over-represented, since more than 75 % of DMRs identified in our capture approach correspond to these genomic features. Interestingly, DMRs in these enhancers, that are particularly important for brain functions, if they would persist overtime in the adult, could disturb their activities, associated gene expression and explain part of the brain defects associated to FASD. The fact that we could identified statistically robust immediate DMRs is suggestive that, in the perturbed, somewhat chaotic, environment, PAE triggers immediate DNA methylation alterations that are common to and could explained shared phenotypic traits among individuals diagnosed for FADS. Conversely, other immediate changes that would occur more randomly amongst individuals might also have phenotypic consequences and be linked to the clinical variability observed in FASD patients (Cobben et al., 2019).

### Immediate DMRs are linked to genes showing dynamic, physiological changes in chromatin accessibility and gene expression

We found that DNA methylation changes occur in specific regions that are, under no stress conditions, physiologically modified in terms of chromatin accessibility and gene expression, at the stage at which PAE was inflicted. First, DMRs are particularly enriched in regions whose chromatin gains accessibility between E14.5 and E15.5 in physiological conditions, and the corresponding enhancers are associated with genes whose GO terms underline importance for brain development. Any perturbation of their methylation landscape by PAE might thus impair the physiological expression of these genes. Second, DMRs are also significantly associated with genes exhibiting upregulation of their expression, either between E14.5 and E15.5 or between E15.5 and E16.5. Notably, *Satb2*, one of the genes that undergoes upregulation during these two time-windows, in physiological conditions, is associated in our analysis with a PAE-induced hypermethylated DMR located in an enhancer (DMR in intron 2 on 10). This gene was previously identified by Hashimoto-Torii et al (2011) as the most downregulated gene in a similar paradigm of PAE (ethanol injections between E14 and E16). They also found that *SATB2* expression was dysregulated in human cortices isolated from gestational week 15-18 fetuses, which were *ex vivo* exposed to ethanol. In their mouse model, the perturbation of *Satb2* gene expression persists after birth (postnatal day 14, Hashimoto-Torii et al., 2011), suggesting that the presence of such DMRs could potentially and persistently alter gene expression at temporal distance from the last ethanol injection. Similarly, the DMR-associated *Mapre2* gene, which encodes a microtubule-associated protein, shows physiological increased chromatin accessibility between E14.5 and E15.5, and upregulated expression between E15.5 and E16.5. DNA methylation changes in *Mapre2* DMR could therefore been followed by alteration in its expression. Interestingly, *Tiam2*, which contains one statistically significant DMC also belongs to the group of genes upregulated between E15.5 and 16.5.

In addition, we verified that three genes, *Mid1, Auts2 and Shank2*, whose mutations are associated with neurodevelopmental disorders including ASD for *Auts2* and *Shank2,* are associated with DMRs and show correlative disturbances of expression, immediately after PAE.

### Immediate DNA-methylation changes affect mono-allelically genes and correlates with misexpression upon PAE

Immediate DMRs are over-represented in imprinted genes in our PAE paradigm. They are mainly located within the promoter regions and active enhancers. We find that most of imprinted genes are ranked in the top of DMR-containing genes in terms of percentage of differential methylation and that these DMRs mostly overlap with the ICR of these genes, which is tightly controlled by DNA methylation and essential for parental monoallelic expression of these genes. The other family of genes prominently showing immediate alteration of DNA methylation patterns is the *clustered protocadherin* genes, which are individually regulated by DNA methylation. Interestingly, these two family of genes, which are key for neurodevelopment and neuronal function, have been already identified as associated to DMRs as in the adult brain in mice or FASD cohorts (Laufer et al., 2013; Lussier et al., 2017) and in buccal swabs in cohorts of FASD children (Cobben et al., 2019; Laufer et al., 2015) therefore at temporal distance from the cessation of alcohol exposure.

Historically, alteration of DNA methylation patterns in imprinted genes has been also identified at the *H19/Igf2* control region in mouse preimplantation embryos exposed *in utero* to alcohol at temporal distance of exposure, since material (whole embryos and placentas) was collected at E10.5 (Haycock, 2009). Early alteration in the DNA methylation status of imprinted genes has been suggested by Downing et al. (2011), who observed subtle decrease in DNA methylation and gene expression at the mouse *Igf2* locus at E9, four hours after prenatal alcohol exposure in whole embryos and placentas, which were ameliorated by diet supplementation with methyl-group precursors Downing et al., 2011). Other studies have also identified differential methylation status in imprinted genes after *ex vivo* ethanol exposure of cultured mouse embryos at the early neurulation stage (after 44 hours of exposure, Liu et al., 2009), as well as in human embryonic stem cells exposed to ethanol for 24 or 48 hours (Khalid et al., 2014). These two *ex vivo* studies were suggestive that at least some imprinted genes could show early modifications in their DNA methylation profiles and thus, in line with our unbiased *in vivo* findings, using a methylome capture in the neocortex and two hours after PAE, only.

Among the genes whose expression is physiologically upregulated between E14.5 and E15.5 and that are associated with DMRs in our PAE paradigm, we identify *Grb10,* whose specific expression from the paternally inherited allele is involved in adult behaviour (Dent and Isles, 2014; Perez et al., 2016). Interestingly, *Grb10* has already been identified as differentially methylated in adult mice, after PAE by Laufer et al. (Laufer et al., 2013) and by Liu et al. (2009), as well as deregulated expression in response to very brief alcohol exposure in astrocytes (Pignataro et al., 2009). Interestingly, some DMRs are located in promoter region of several protocadherin genes, including *Pcdhgb4* and *Pcdhga12*. These observations suggest that expression of these particular genes could be affected both immediately and later after the PAE.

PAE therefore quickly disturbs DNA methylation in these mono-allelically expressed genes, which are key for neurodevelopment. Moreover, we show that, at least for some of them, the presence of immediate DMRs in the ICR correlates with concomitant perturbation of their expression, which could potentially have rapid detrimental consequences on neurodevelopment. Note that the expression of the genes is disturbed in a relatively modest manner. However, even a 20% difference in gene expression upon PAE correlates with neurodevelopmental defects (El Fatimy et al., 2014).

Moreover, as quoted above, at long temporal distances from PAE, DMRs were detected in several mono-allelically expressed genes of these two families, in the adult brain in mice (Laufer et al., 2013; Lussier et al., 2017; Legault et al., 2021) and in buccal swabs in cohorts of FASD children (Cobben et al., 2019; Laufer et al., 2015). Our data thus suggest that these immediate alterations in DNA methylation may persist throughout life. In addition, the fact that we demonstrate that they occur so early upon PAE reinforces their potential as biomarkers of exposure. Notably, DMR persistence might not consist in a perfect conservation of the percentage of methylation for each CpG. Indeed, DNA methylation overtime might be modified by many factors, including brain activity itself (Guo et al., 2011). Rather, such persistence might be more qualitative than quantitative in the sense that a given DMR may globally exhibit differential methylation, both at short and long temporal distance from the end of exposure, but with different percentage of individual cytosine methylation at an early, compared to a late time-window.

To the best of our knowledge, our data demonstrate for the first time in an unbiased manner, that DNA methylation alterations in the developing cortex are detected immediately after the cessation of PAE across the genome. Although the number of DMRs that are formed right after PAE is limited, these alterations do not occur randomly, but target enhancer regions that are active at the time of alcohol exposure. Moreover, regions that undergo changes in chromatin accessibility and gene expression seem to be particularly vulnerable to perturbation of DNA methylation patterns and are associated with genes of importance for neurodevelopment. In addition, we show that genes of two gene families, whose monoallelic expression is tightly controlled by DNA methylation and crucial for brain development and function, carry immediate DMRs associated to synchronous dysregulation of their expression. These gene families have been previously identified as carrying or associated to DMRs in the adult brain, in human children or mouse models after PAE. Although they were suspected to be long-lasting epigenetic marks of PAE, there was no direct evidence that that they were established early in response to PAE. Our observations demonstrate that abnormalities in DNA methylation profiles are established immediately upon PAE and strongly suggest that they can persist at very long temporal distance from the end of exposure, and this could be considered as “scars” of exposure to prenatal stress. This makes of these two gene families strong candidates as biomarkers of exposure, which are so urgently needed to better and earlier diagnose FASD children and accelerate and ameliorate follow-up and intervention.

Our characterization of immediate effects of a binge-drinking-like PAE first points to higher variability in EtOH-exposed cortical samples, compared to control ones (**Figure 7**). This observation corroborates previous studies that showed that PAE led to exacerbated cell-to-cell variability in both participant-derived cells, associated with schizophrenia, or in a PAE-mouse model. Notably, these publications are based on a protocol resembling to ours (Hashimoto-Torii et al., 2011; Ishii et al., 2017). The fact that cortices from embryos of the same litter behave differently is also in line with these reports. Nevertheless, we identify genes associated with statistically significant changes in the levels of mRNAs. A large majority of DEGs were not found associated with a DMR. This could be biased by the limits of the DMR capture design: in order to detect subtle methylation differences, only regions extended [-1 kb; +1kb] around a promoter or enhancer were selected for the capture, which excluded a large number of regulatory regions. Nevertheless, our RNA-seq data analysis highlights the dysregulation of the expression of imprinted genes in agreement with the presence of a DMR in their ICR, and with our RT-qPCR results. Combined to the fact that a tight epigenetic control of the expression of some these genes (*Zrsr1*, *Peg13*, *Nap1l5* et *Inpp5f*) is necessary for brain development and function, the finding of their dysregulation encourages future investigations to evaluate their potential as biomarkers of PAE. Indeed, their deregulation is observed very shortly upon PAE, which suggests that it could be directly linked to alcohol exposure and not due to long-term indirect process. The immediate changes in gene expression elicited by this binge-drinking-like PAE are modest in terms of level of fold change. This could be due to the fact that we chose to observed transcriptomic changes only 2 hours upon PAE, in our attempt to identify immediate effects of PAE. However, this timepoint could correspond to the start of transcriptional programs, which could occur slowly, at least for UP DEG. Strikingly, based on GO-term, GSEA, and STRING analyses, we describe a wide downregulation of families of genes involved in ribosome biogenesis, RNA metabolism and processing, and cytoplasmic and mitochondrial translation. We also observe decreased expression in genes involved in splicing. Correlatively, we find the UP DEGs are involved in the formation of diverse cellular granules. For instance, nuclear speckles into which splicing factors accumulate and host genomic regions of shared regulation self-organize in a three-dimensional (3D) manner, to optimize the efficiency and co-transcriptional occurrence of splicing (reviewed in Bhat et al., 2024). We also identified gene networks of importance for cytoplasmic stress granules and p-bodies. They are membrane-less cellular compartments, composed of ribonucleoproteins, which assemble in response to stress and, respectively, contain translation initiation factors and factors related to mRNA degradation and decay. They also represent storage spaces for mRNAs, which altogether support their roles as new players of translational control (reviewed by Riggs et al., 2020). In addition, our analyses pointed to centriolar satellites that participates in the formation of centrosomes and primary cilia structures that are essential for neurodevelopment (reviewed in Odabasi et al., 2020 and Devi, 2021) possibly through supporting translation in these particles of key proteins for these structures (Pachinger et al., 2024). Collectively, our data suggest that shortly after PAE, brain cortical cells adopt a “frozen” state by abruptly slowing down splicing, translation, and respiration, and by increasing the expression of genes that support the formation of diverse cellular bodies, involved in the hosting or sequestration of RNAs and important molecular players that preserve these functions. Importantly, both the slow-down of translation, which has long been shown to be a primary consequence of the cellular response to heat shock (HSR; Lindquist 1981; Desroches Altamirano et al., 2024) and the stalling of ribosomes (ribosome collision) induced by stress, which triggers ribosome quality control (RQC), maintain proteostasis (homeostasis of the proteome; reviewed in Kim and Zaher, 2022). Our findings suggests that PAE quickly triggers a program that by reducing translation, ribosome biogenesis, and respiration and by allowing storing RNAs and RNA-binding proteins in specialized cytoplasmic or nuclear domains, contributes to proteostasis, and thereby protects the brain cortical cells. In the same line of freezing process that could lead to proteome damage under stress, and activating those that could contribute to maintain the cell homeostasis, GO-terms analyses also suggest that genes important for vesicle exocytosis and transport, as well as synaptic vesicles, are downregulated, whereas genes encoding transmembrane transporters are upregulated, evoking a need to secure the maintenance of a correct intracellular environment in the cells of the developing cortex exposed to alcohol.

In addition, our data indicate that PAE rapidly triggers the upregulation of the Wnt pathway that governs cell fate, cell migration, cell polarity, and neural patterning (Rim et al., 2022) and the TGFb - SMAD - BMP signaling pathway, which is necessary for the differentiation of neural stem cells and neural progenitor cells, neurite outgrowth, and dendritic development, and axon growth, retrograde transport along the axon (likely through cytoskeleton dynamics), and synapse formation (Kashima and Hata, 2017).

In conclusion, PAE provokes immediate rewiring of the transcriptome. This potentially has dual consequences: 1) this could lead to beneficial impacts on the recovery of cells from alcohol exposure, through slowing down protein synthesis and energy-consuming respiratory pathways; 2) it could be detrimental through the inappropriate activation of critical gene expression programs that are susceptible to rapidly and directly perturb neurodevelopment.

## MATERIALS AND METHODS

### Mice mating and alcohol exposure

C57Bl/6N murine females of 2- to 4-months-old were time-mated and assessed for mating based on the presence of a vaginal plug. The noon of vaginal plug day was considered as embryonic day 0.5 (E0.5). For binge drinking stress, pregnant females received intra-peritoneal injections of ethanol (3g/kg, diluted in a final volume of 500 µL of PBS), at embryonic days E15, E15.5 and E16 (**Figure 1A**). Control group received similarly PBS alone. Embryonic cortices were collected 2 hours after the last injection.

### Tissue collection and genotyping

Embryonic cortices were harvested on ice, in cold L-15 medium (Leibovitz Gibco #11415-049). Only male cortices were used for methylome analysis, whereas both male and female cortices were used for gene expression analysis. For methylome analysis, three replicates were generated per group.1M, 2M and 3M are ethanol-treated samples, while 4M, 5M and 6M design the PBS-treated ones (control). In order to reduce inter-litter variability, each replicate was composed of 2 right hemi-cortices and 2 left-hemi cortices, from embryos originated from 4 distinct litters, see **Supplementary Figure 1B**.

Since sexual dimorphism was reported for behaviors of individuals exposed *in utero* to alcohol (Hellemans et al., 2010), we determined the sex of each embryo. Genomic DNA was extracted from animal tails incubation (95°C for 1h) in the extraction buffer (25 mM NaOH, 0.2 mM EDTA) then neutralized in Tris-HCl 40mM, pH5 (vol/vol). Sex of the embryos were identified by PCR genotyping using Ube1R (5’-CACCTGCACGTTGCCCTT-3’) and Ube1F (5’-TGGATGGTGTGGCCAATG-3’) primers that target *Ube1X* and *Ube1Y* genes (Sugimoto and Abe, 2007). These primers allow to amplify a 252bp amplicon from *Ube1X* and 334bp one from *Ube1Y*.

### ATAC-seq and RNA-seq ENCODE data

Data mining from ATAC-seq and RNA-seq datasets was performed, using available public ENCODE data. These ATAC-seq and RNA-seq experiments were done using embryonic and newborn forebrains of C57BL/6N mice (experiments from Bing Ren laboratory, UCSD for the ATAC-seq and experiments of B. Wold, from Caltech laboratory for the RNA-seq).

ENCODE accession number of the data analysed

|  |  | E13.5 | E14.5 | E15.5 | E16.5 | P0 |
| --- | --- | --- | --- | --- | --- | --- |
| ATAC-Seq<br>ENCODE<br>accession<br>number | Sample<br>#1 | ENCFF401VUV (R1)<br>ENCFF898NRO (R2) | ENCFF048MTG (R1)<br>ENCFF890LGM (R2) | ENCFF248PXW (R1)<br>ENCFF825UHO (R2) | ENCFF058IAE (R1)<br>ENCFF765HUX (R2) | ENCFF197GTC (R1)<br>ENCFF209GGJ (R2) |
|  | Sample<br>#2 | ENCFF721LGJ (R1)<br>ENCFF777UK (R2) | ENCFF633MTW (R1)<br>ENCFF666DRJ (R2) | ENCFF906VXU (R1)<br>ENCFF500SXI (R2) | ENCFF776GDQ (R1)<br>ENCFF588XZG (R2) | ENCFF296GZG (R1)<br>ENCFF664RZO (R2) |
| RNA-seq<br>ENCODE<br>accession<br>number | Sample<br>#1 | ENCFF235DNM (R1)<br>ENCFF959PSX (R2) | ENCFF270GKY (R1)<br>ENCFF460TCF (R2) | ENCFF179JEC (R1),<br>ENCFF891HIX (R2) | ENCFF931IVO (R1)<br>ENCFF114DRT (R2) | ENCFF037JQC (R1)<br>ENCFF358MFI (R2) |
|  | Sample<br>#2 |  | ENCFF126IRS (R1)<br>ENCFF748SRJ (R2) |  |  | ENCFF447EXU (R1)<br>ENCFF458NWF (R2) |

### Methylome capture and high-throughput sequencing

To analyze the methylation of genomic regions that we selected (see below details about the design of regions, **Figure 1B** and **Supplementary Figure 1A**), a methylome capture was performed, using *SeqCapEpi Developer Medium Enrichment kit* (*NimbleGen*, *Roche*), following User’s Guide and manufacturer’s instructions. Briefly, gDNA samples were fragmented using Covaris technologies and non-directional libraries were prepared using KAPA Biosystems DNA Library Preparation Kits with *NimbleGen SeqCap* Adapter Kits. Then, bisulfite conversion of these DNA samples libraries was performed using the *Zymo EZ DNA Methylation-Lightning* kit, before a DNA amplification using *KAPA HiFi HotStart Uracil* and *ReadyMix*. Capture of these bisulfite-converted sample libraries was done by hybridization, using *SeqCap Developer M Enrichment*, *SeqC*ap *Epi Accessory*, and *SeqCap HE-Oligo* Kits. In order to reduce variability between samples, the six libraries (one per sample) were multiplexed and captured all together on beads. This multiplexing during the ‘capture step’ has already been successfully tested with similar capture approaches (Allum et al., 2015; van der Werf et al., 2015). DNA was then amplified again using the *KAPA HiFi HotStart ReadyMix*.

Captured and amplified bisulfite-converted gDNA fragments were sequenced using an *Illumina* sequencing instrument (Paired-end, 150bp, NextSeq 500 High Throughput). All these steps were performed by *Institut du Cerveau et de la Moelle épinière* (ICM) sequencing platform.

### Methylome capture composition

The capture is composed of 58,611 chosen genomic regions (**Supplementary dataset 3**), based on ENCODE available data or based on previous lab results (**Figure 1B** and **Supplementary Figure 1A**). Majority of captured regions corresponds to active enhancers in adult (8 weeks-old) mouse cortex (characterized by the H3K27ac histone mark, regions of +/- 500 bases from the middle of the peaks, ENCODE data). Capture also includes mouse promoter regions (+/- 500 bases from the transcription start site, ENCODE data). Due to repetitive sequences that prevent the design of specific probes of some regions, and also because of annotation database information differences, only 75% of known promoter regions could be included in the capture. “Other regions” category is composed of genomic regions of interest based on previous lab results. It corresponds to regions potentially involved in stress response: (**i**) HSF2 binding sites found in unstressed mice cortices of E16.5 embryos (ChIP-seq data, VM lab); and to dynamic regions upon stress : (**ii**) differentially opened or closed regions (DOCR) identified in isolated oligodendrocyte precursors (O4+ cells) from cortices of 5 days-old mice, after an inflammatory stress from postnatal day P1 to P5, mimicking a neuroinflammation in the third trimester of pregnancy in human, (Schang et al., 2018, GSE197563); (**iii**) enhancers, characterized by the H3K4me1 histone mark (+/- 100 bases from the middle of the peaks) of adult (8 weeks-old) mouse cortices among genes (and at +/-20 kb from genes ends) that were differentially expressed in response to another prenatal stress (neuroinflammation; Krishnan et al., 2017, GSE197563). Detail number on these sub-category “other regions” are indicated **Supplementary Figure 1A**. List of mm9 coordinates of all the genomic regions contained in the capture are available in **Supplementary Dataset 3**.

### Bioinformatic analysis of methylome capture

Bioinformatic workflow used to analyze methylome capture is described in details in **Supplementary data Notebook 1** and key figures of the analysis are shown in **Supplementary Table 2**. Key steps of the analysis and associated tools are shown in **Supplementary Figure 2A**.

#### Trimming, mapping, methylation count table, and visualization

Briefly, reads quality was verified using *FASTQC* (http://www.bioinformatics.babraham.ac.uk/projects/fastqc). To decrease methylation call errors from poor quality data, a trimming was carried out with *Trimmomatic* (Bolger et al., 2014) and *Cutadapt* (Martin, 2011). Trimmed reads were then paired-end mapped on the whole *Mus musculus* reference genome mm9, using *bismark* (Krueger and Andrews, 2011) which is adapted for bisulfite converted reads. Libraries were considered as non-directional for the mapping. Deduplication of reads was performed using *deduplicate_bismark* function (from *bismark* suite)

Only cytosines on CpG context were studied. For each sample, number of methylated and unmethylated cytosines that covered a given cytosine site was obtained using *bismark_methylation_extractor* (BME) function (from *bismark suite*), with *no_overlap* option to prevent double counting of paired-end results. Then, to select only information that corresponds to capture regions, *bedtools intersect* (Quinlan and Hall, 2010) was used to intersect capture regions with mapping files, or with CpG count tables. To avoid removal of potential interesting sequences, at the border of capture regions, each region was extended up and downstream with 150 bases, before the intersections, and were named *enlarged capture*). Mapping files restricted to the *enlarged capture* regions were sorted and indexed using *samtools suite* (Li et al., 2009) before their visualization on *IGV* (Robinson et al., 2011).

#### Detection of differentially methylated isolated CpG (DMCs)

Data from BME were formatted on *R* (The R Core Team, 2018) in order to use *MethylKit* R package (Akalin et al., 2012), for detection of significant differentially methylated isolated CpG between control and alcohol-treated samples. For this statistical analysis, formatted data were filtered: cytosine positions displaying abnormal high read coverage (> 99.9^th^ percentile) or low read coverage (number of reads < 10) were excluded from the analysis, to avoid bias (Krueger et al., 2012). Read coverage was also normalized before statistical analysis, using median to calculate scaling factor. Only methylation state of CpG sites that were covered in all samples was investigated for the analysis (corresponding to 1,259,111 positions). DMCs were considered as significant when difference of methylation state between the two groups was higher than 5%, for a qvalue < 0.05.

#### Detection of differentially methylated regions (DMRs)

We generated an *R* function (called *get_close_loci()*) to define DMRs (**Supplementary Figure 3,A**-D). This function combines neighbouring CpGs that share similar differential methylation state, (all hypo- or all hypermethylated CpGs, **Supplementary Figure 3A,C**). Because *MethylKit* provides relevant statistical information about individual CpG, we used *MethylKit* statistical output as an input of *get_close_loci()* function (*i.e.* position of the CpG site, percent of methylation difference between groups for a given site, p-value and q-value). Data from BME were thus formatted on *R* (The R Core Team, 2018) before using *MethylKit* (Akalin et al., 2012). Formatted data were filtered and normalized, with the same parameters than those used for DMC detection. Only methylation state of CpG sites that were covered in all samples was investigated for the analysis (corresponding to 1,259,111 positions). To define DMRs with *get_close_loci()*, only CpG sites having a reliable methylation state must be used, since one non relevant CpG is sufficient to affect DMR detection (**Supplementary Figure 3C**). Thus, a pre-selection of *relevant CpGs* was done, according to their p-value (CpGs with pvalue < 0.07), whose threshold was determined using comparisons of real and random datasets (see below, **Supplementary Figure S4**, **Supplementary Table 2**). To define DMRs, parameters were defined as follow: association of at least 5 CpG having a same methylation state, in a region of maximum 2000 bases, with a maximum distance of 100 bases between two successive selected CpG, according to p-value threshold (pval < 0.07, **Supplementary Table 2**).

#### Randomization to define appropriate parameters for DMR detection

Random datasets (n=2) were generated and compared to real dataset, to define parameters of *get_close_loci()* function that were appropriated for DMRs detection in the real data. To obtain these random data, methylation states from real samples, were randomly switched, CpG position by CpG position, between samples and replicates using our own *R* script (**Supplementary Figure S4**, see details in **Supplementary data notebook 1**). Each methylation state is thus reassigned to a given sample, but methylation states between distinct CpG positions are not shuffled. This randomization assigns biologically plausible methylation values, actually observed at a given cytosine position (this is not the case if randomization was global, *i.e* redistribution of CpG methylation state, regardless of the CpG position, because there would be a risk of assigning, for example, high DNA methylation rates to a cytosine position that is always detected as non-methylated, regardless the conditions, PBS or EtOH-treatment). Randomization was done on filtered but unnormalized real dataset. Read coverage normalization was performed after the randomization, using median to calculate scaling factor. To define appropriate parameters for DMR detection, different parameters were modulated and the number of DMRs found in the two datasets (real versus random) was compared (**Supplementary Table 2**). Similar results were obtained by using the two random datasets (**Supplementary Table 2**).

#### Annotation

To be able to compare DMRs with other bioinformatic analysis (ATAC-seq and RNA-seq results), data obtained with mm9 coordinates were then converted with mm10 coordinates using *LiftOver* software (https://genome.ucsc.edu/cgi-bin/hgLiftOver), available on UCSC, with default parameters. Annotation file, based on mm10 coordinates was obtained using *BiomaRt R* package (Durinck et al., 2009; Huang et al., 2009). DMRs were annotated in *R* according to this file information (see details in **Supplementary data notebook 1**). A file combining all information (DMRs coordinates, annotation and statistical information) were generated (**Supplementary Dataset 1**). To estimate if DMRs are particularly located in a given capture categories (enhancers, promoters…), hypergeometric tests were done on *R* (using *phyper()* function), taking into account the number of each capture category that is represented in the methylome capture. Threshold chosen: pvalue < 0.05.

#### Motif enrichment

Transcription factors binding site enrichment within the DMRs located in active enhancer regions (H3K27ac mark), were explored using *findMotifsGenome* (*HOMER* suite, see details in **Supplementary data notebook 1**; Heinz et al., 2010).

#### Over-representation of Imprinted genes among DMRs

To estimate whether DMRs are particularly located in imprinted genes (IG), hypergeometric tests were done on *R* (using *phyper()* function). Threshold chosen: pvalue < 0.05. The exact number of capture regions that corresponds to imprinted genes are unknown, but we estimated that about 150 capture regions could be attributed to imprinted gene regions (there are almost 150 mouse IG, according to http://www.geneimprint.com/site/genes-by-species.Mus+musculus website, and we potentially have, at least one promoter for each of this gene in the capture). Using this approximation, results of *phyper* test (phyper(11,150,58611-150,432)) is equal to 1.51e-09, which is significant. Even with an over-estimation of capture regions that correspond to imprinted gene regions, results of *phyper* test is significant (*e.g.* if we considered that 300 capture regions corresponds to imprinted genes regions (*i.e.* two regions per IG), phyher(11,300,58611-300,432) = 2.89e-06).

#### Estimation of sodium bisulfite conversion efficiency

To ensure that sodium bisulfite treatment correctly converted unmethylated cytosines, we evaluated the conversion efficiency by looking at conversion rate of a spike-in DNA (*i.e.* unmethylated known sequences of non-mammalian DNA), which was added in each sample preparation. This spike-in DNA, corresponding to sequences of lambda phage, was mapped on an appropriate reference genome, using *bismark*. The efficiency of sodium bisulfite conversion is obtained using deduplicated spike-in DNA mapped reads, by calculating the ratio of converted reads (containing thymines) to the total number of reads covering these cytosines (**Supplementary Table S1**).

### Data mining of ENCODE data: Chromatin accessibility profile of the developing brain: ATAC-seq data time-course analysis

Bioinformatic workflow used to analyze ATAC-seq data is described in details in **Supplementary data notebook 2**. Key steps of the analysis and associated tools are showed in **Supplementary Figure S2A** and key features of the analysis are shown in **Supplementary Table S4**. Quality of sequenced reads was verified for each sample using *FASTQC* (http://www.bioinformatics.babraham.ac.uk/projects/fastqc). Reads having bad sequencing quality were trimmed using *Trimmomatic* (Bolger et al., 2014). Trimmed reads were then paired-end mapped on *Mus musculus* reference genome mm9, using *bowtie2* (Langmead and Salzberg, 2012). Then, reads were deduplicated using *samtools rmdup* (Li et al., 2009). Reads that mapped to mitochondrial chromosome were removed using tools on *Galaxy* platform (Afgan et al., 2016, default parameters), by splitting the mapped reads per chromosome using *bam-splitter tool* (Barnett et al., 2011). Then all the files were merged except the reads mapped on mitochondrial chromosome, using *merge-bam tool*. *MACS2* (Zhang et al., 2008) was then used to identify peaks corresponding to opened genomic regions, in each sample. For that, mapped reads from duplicates were merged, before this Peak calling. Then, table containing all opened regions, detected in at least one sample, was generated, using *bedtools multiinter* and *bedtools merge* (Quinlan and Hall, 2010). To obtain reads count at these regions, in each sample, *htseq-count* (Anders et al., 2015) was runned. This tool prevents double counting of paired-end results since files containing mapped reads were sorted by read name. To identify regions where chromatin accessibility significantly changes during development, pairwise comparisons of successive developmental stages were performed using *edgeR* (Robinson et al., 2010), an *R* software (The R Core Team, 2018) and *Bioconductor* package (Gentleman et al., 2004). We adapted a script template called *template_script_edgeR.r* from ***Sartools*** *R* package (Varet et al., 2016), which implements some ***edgeR*** functions (see details in **Supplementary Data notebook 2**). To compare ATAC-seq results with other bioinformatic analyses (methylome capture and RNA-seq results), data obtained with mm9 coordinates were then converted with mm10 coordinates using *LiftOver* software (https://genome.ucsc.edu/cgi-bin/hgLiftOver), with default parameters. Annotation file, based on mm10 coordinates was obtained using *BiomaRt R* package (Durinck et al., 2009; Huang et al., 2009). Differentially open and closed regions (DOCR) observed during brain development were annotated in *R* according to this file information (see details in **Supplementary Data notebook 2**). For each pairwise comparison, a file combining all information (DOCR coordinates, annotation and statistical information) were generated (**Supplementary Dataset 1**). To visualize data on *IGV* (Robinson et al., 2011), files containing mapped reads were sorted and indexed using *samtools* suite (Li et al., 2009).

### Data mining of ENCODE data: Transcriptome profile of the developing brain: RNA-seq data time-course analysis

Bioinformatic workflow used to analyze RNA-seq data is described in detail in **Supplementary data notebook 3**. Key steps of the analysis and associated tools are showed in **Supplementary Figure 2A** and key features of the analysis are shown in Supplementary Table 5. For all samples, sequenced read quality was verified for each sample using *FASTQC* (http://www.bioinformatics.babraham.ac.uk/projects/fastqc). Adapter sequences and reads having bad quality were trimmed using *TrimGalore* (https://github.com/FelixKrueger/TrimGalore). Trimmed reads were then singled-end mapped using *STAR* on *Mus musculus* reference genome mm10, which was associated to its corresponding *Gencode* annotation during indexation, to improve accuracy of the mapping (Dobin and Gingeras, 2015; Dobin et al., 2013). For each sample, *htseq-count* (Anders et al., 2015) was used to obtain reads count of all genes contained in *Gencode* annotation file. To identify differentially expressed genes (DEG) during physiological brain development, pairwise comparisons of successive developmental stages were performed using *edgeR* (Robinson et al., 2010). We used exactly the same script than those executed for ATAC-seq analysis (*i.e.* modified script from *Sartools R* package (Varet et al., 2016, see details in **Supplementary data notebook 3**). Annotation file, based on mm10 coordinates was obtained using *BiomaRt R* package (Durinck et al., 2009; Huang et al., 2009) and was used to annotate DEG observed during brain development with *Unix* commands (see details in **Supplementary data notebook 3**). For each pairwise comparison, a file combining all information (DEG coordinates, annotation, and statistical information) were generated (**Supplementary Dataset 1**). To visualize data on *IGV* (Robinson et al., 2011), files containing mapped reads were sorted and indexed using *samtools* suite (Li et al., 2009).

### Data mining of ENCODE data: Gene ontology analyses

In order to perform Gene ontology analysis, gene lists (based on gene symbol names) were defined for each distinct datasets (*i.e.* genes associated to DMRs that are located in (or close to) captured active enhancers (H3K27ac) and genes associated to DOCR and DEG). For DOCR and DEG, only genes that are modulated between E14.5 and E15.5 and between E15.5 and E16.5 were studied. Gene ontology analyses were performed with WEB-based GEne SeT AnaLysis Toolkit (GESTALT, http://www.webgestalt.org/, using default parameters, except for the minimum number of genes for a category (set to one in our analysis). The reference gene list used for each analysis was either the genome for DOCR and DEG or all the genes of the studied capture category (*i.e.* in case of the “DMRs enhancer” list: all the genes associated to H3K27ac histone mark that are represented in the capture design (**Supplementary Dataset 3**).

### RNA extraction and purification

Each E16.5 embryonic cortex in 500µL Trizol (*Sigma*) was lysed at 4°C using *Precellys/Cryolys* (Bertin) with 1.4mm ceramic beads (Mobio) during 10 seconds (5500 rpm). mRNAs were isolated using chloroform and isopropanol, followed by ethanol precipitation. Samples were resuspended into 50µL of ultrapure RNase-free Water (*Invitrogen*). Integrity of RNA was verified on an electrophoresis gel (detection of 18S and 28S ribosomal RNA). Extracts were DNAseI treated (*Roche - ref 04 716 728 001*) in presence of RNAse inhibitors (*Promega - Ref N251A*), before RNA purification using phenol chloroforme isoamyl-alcohol and chloroforme, followed by ethanol precipitation. Samples were resuspended into 20µL of ultrapure RNase-free Water. RNA was quantified with Denovix machine (DS-11 Denovix series).

Reverse transcription was done using *Biolabs* kit (*Lunascript RT SuperMix E010G*), on less than 5µg of RNA, as recommended by the manufacter. cDNA concentration was normalized to 3ng/µL by diluting it into ultrapure water. To verify the purity of RNA extraction (absence of genomic DNA), a negative control without reverse transcriptase was also performed. Quality and Integrity of cDNA were verified by calculating 3’/5’ ratio of *Hprt1* gene, following MIQE recommendations (Bustin et al., 2009). Only cDNA preparations with a ratio lower than 5 were used for RT-qPCR (Bus**tin** et al., 2009).

### RNA-seq analysis: Differential transcriptome profile of E16.5 mouse brain cortices

The quality control of the raw data was performed with FastQC (v.0.11.8). The trimming has been done with Trim_Galore (v.0.5.0 with cutadapt 1.18 and python 3.7.1). The mapping has been done with STAR v2.6.1d. The visualisation of the mapping was performed with Samtools v1.9 and IGV. The counting of each expressed genes for each sample was performed with htseq-count v0.11.2 and the table count was obtained with R. The differentially expressed genes between EtOH samples and control (PBS samples) were found using edgeR via Sartools v1.7.3 on R v4.0.3.

the genes are normalized using TMM (Trimmed Mean of M component normalization) (**Robinson and Oshlack, 2010**). For more details see **Supplementary data notebook 5**

#### Gene ontology

Gene ontology (GO) analysis of differentially expressed genes was performed using Org.Mm.eg.db v3.18.0 data base using GO term over-representation test from ClusterProfiler v4.10.1., the annotation of genes with GO terms was done using biomaRt v2.58.2.

#### Gene set enrichment analysis

Gene set enrichment analysis (GSEA) was performed with GSEA v4.3.3 (Subramanian,A et al., 2005) in preranked mode with 1 000 permutations. Genes were pre-ranked by fold change derived from the differential expression analysis. Gene sets of size above 15 and 500 from SigDB.v2024.1.Mm.chip (Liberzon,A et al, 2015) were used.

#### Heat map representation

Heat maps were created using Morpheus (https://software.broadinstitute.org/morpheus). The gene expression was visualized using a relative color scheme (from minimum to maximum values in each row). The color scales are indicated on each heat map. Rows were organized based on their enrichment score in the biological process of interest following GSEA analysis.

### Analysis of protein-protein interaction

The DEGs, identified in the RNA-seq analysis, were imported into Cytoscape software (v3.9.1) to construct a protein-protein interaction network from STRING database (https://string-db.org/) using *stringApp* (<u>Doncheva et al., 2019</u>). We generated a physical network with a confidence cutoff of 0.7 and no maximal additional interactors. In addition, we performed MCL (Markov *Cluster* Algorithm) clustering analysis using *ClusterMarker2* (Morris et al., 2011) plugin in Cytoscape to identify interconnected regions within the protein-protein interaction network to unravel potential functional modules and interactions between genes. Next, we performed a STRING-enrichment analysis to determine the functional relevance of nodes by identifying the gene ontology (GO-terms associated with them.

### Quantitative Real-Time RT-PCR

For real-time RT-qPCR, *LunaGreen* S*YBR* mix (*Biolabs*) was used, as recommended by the manufacturer. Primers were purchased from Sigma-Aldrich. Probes sequences are listed in **Supplementary Table 6**. 384-wells PCR plates were filled using Tecan pipetting robot. RT-qPCR experiments were done using LightCycler LC480 (Roche). All reactions were made in quadruplicate for samples originated from at least three distinct litters.

Relative quantities of the target gene mRNAs were normalized to *Hprt1* and *Tpt1*. Qbase+ software was used to define the values of calibrated and normalized relative gene expression (CNRQ).

Non parametric Mann-Whitney tests were performed using R software, to determine the pvalues when comparing mRNA levels between PBS and EtOH-treated samples. To take into account multiple tests, p-values were corrected with Benjamini-Hochberg correction (padj.), using *p.adjust()* R function. For this analysis, a gene is considered as differentially expressed between PBS and EtOH-treated samples, when the adjusted p-value is below 0.05 (padj. < 0.05).

## Funding information

VM was funded by the Agence Nationale de la Recherche « HSF-EPISAME », SAMENTA ANR-13-SAMA-0008-01) and CNRS. AD was supported by a Doctoral Fellowship from the French Ministère de l’Enseignement Supérieur, de la Recherche et de l’Innovation (MESRI) and FM from a Doctoral Fellowship from the CNRS. This study contributes to the Université de Paris IdEx #ANR-18-IDEX-0001 funded by the French Government through its “Investments for the Future” program. DSD benefited from a CNRS Délégation de Recherche (2018-2020). OTT benefited from travel grants from Université Paris Diderot.

## Supporting information

Supp Table S1 Bisulfite conversion efficiency

Supp Table S2 Random datasets

Supp Table S3 Key Numbers Methylome Analysis

Supp Table S4 Datamining ATAC-seq data

Supp Table S5 Datamining ENCODE RNAseq

Supp Data Notebook1 Methylome workflow

Supp data Notebook 2 ATAC-seq data mining

Supp data Notebook 3 RNAseq data mining

Supp data Notebook 4 Integration data

Supp Dataset 1 List DMR DOCR DEG

Supp Dataset2 List integration DMR DOCR DEG

Supp Dataset 3 Capture regions

Supp Table S6 RNAseq DEG PBSvsEtOH

Suppl Table S7 data integration_DMR-RNA-Seq PAE DOCR DEG

Suppl Table S8 GO_BP_CC_MF PAE

Suppl Table S9 GSEA_Report EtOH & PBS GO global table

Suppl. Table S10 Gene list of selected GSEA GOs

Suppl. Table S11_STRING DEG PBSvsEtOH

Supp data Notebook 5 Statistical report Sartools-edgeR-RNAseq-PBSvsEtOH

## Acknowledgements

We are grateful to Slimane AIT-SI-ALI, (Université Paris Cité, CNRS Epigenetics and Cell Fate, Paris, France), Maxim GREENBERG (Université Paris Cité, CNRS Institut Jacques Monod Paris) and Deborah BOURC’HIS (Institut Curie, Paris) for helpful discussions. We are grateful to the VM team members, Aurélie de THONEL and Véronique DUBREUIL (Université Paris Cité, CNRS Epigenetics and Cell Fate, Paris, France) for helpful discussions and comments on the manuscript. We thank Renaud MASSART, former postdoc in the VM Lab, for designing methylome capture. We are grateful to Emeline MUNDWILLER and Yann MARIE (Brain and Spine Institute, Paris, France) for performing the methylome capture libraries and high-throughput sequencing of these samples. We thank Isabelle Le PARCO and the staff from the Buffon animal housing facility at the Jacques Monod Institute (Université Paris Cité, Paris, France). We thank Magali HENNION, Head of the Bioinformatics and Biostatistics Core Facility at the CNRS Epigenetics and Cell Fate Center (Université Paris Cité) for bioinformatics support.

The authors declare no conflict of interest.

## Sequences of primers used for RT-qPCR experiments.

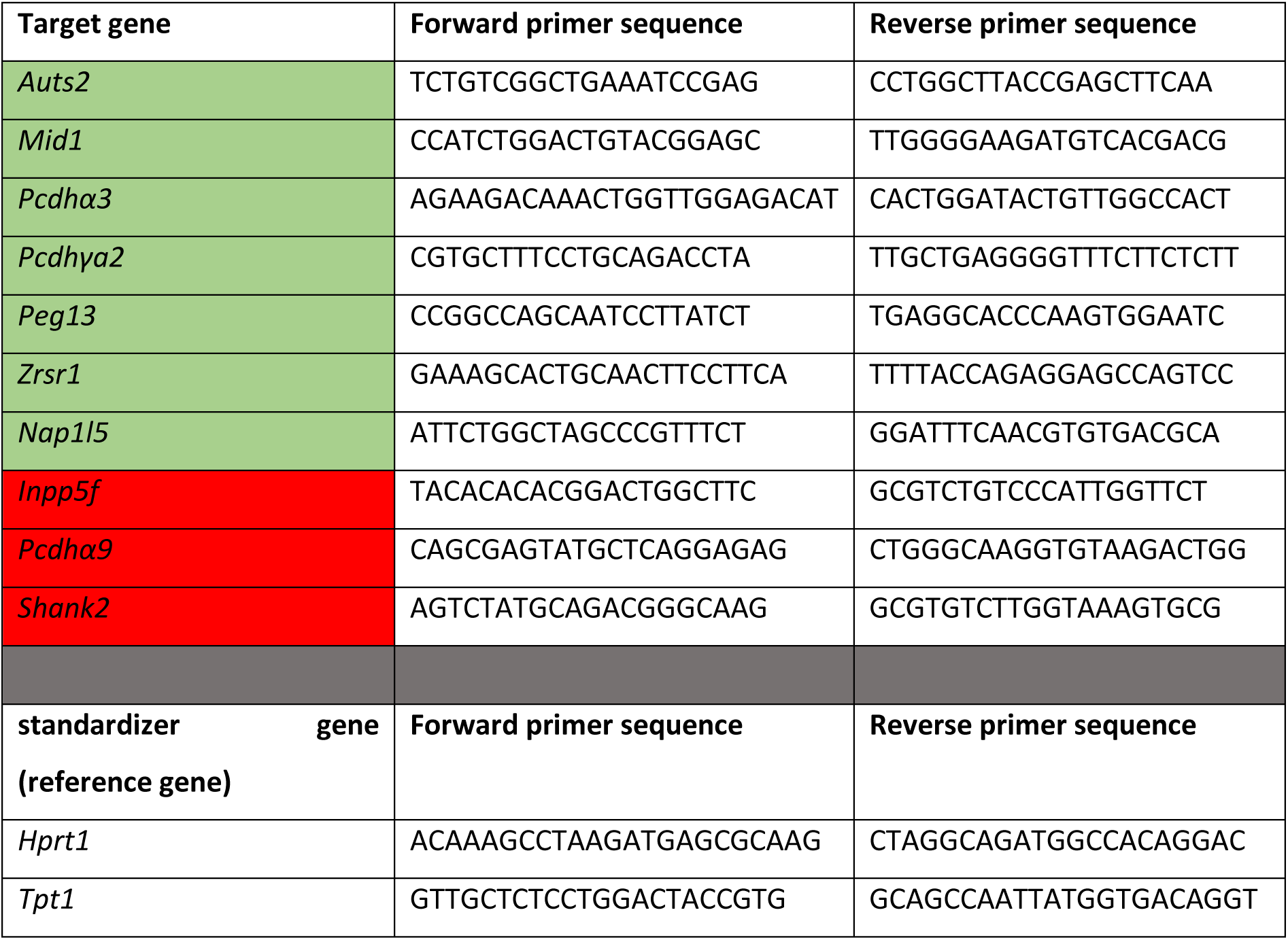

## Legends of Supplementary Figures

**Supplementary Figure 1.**
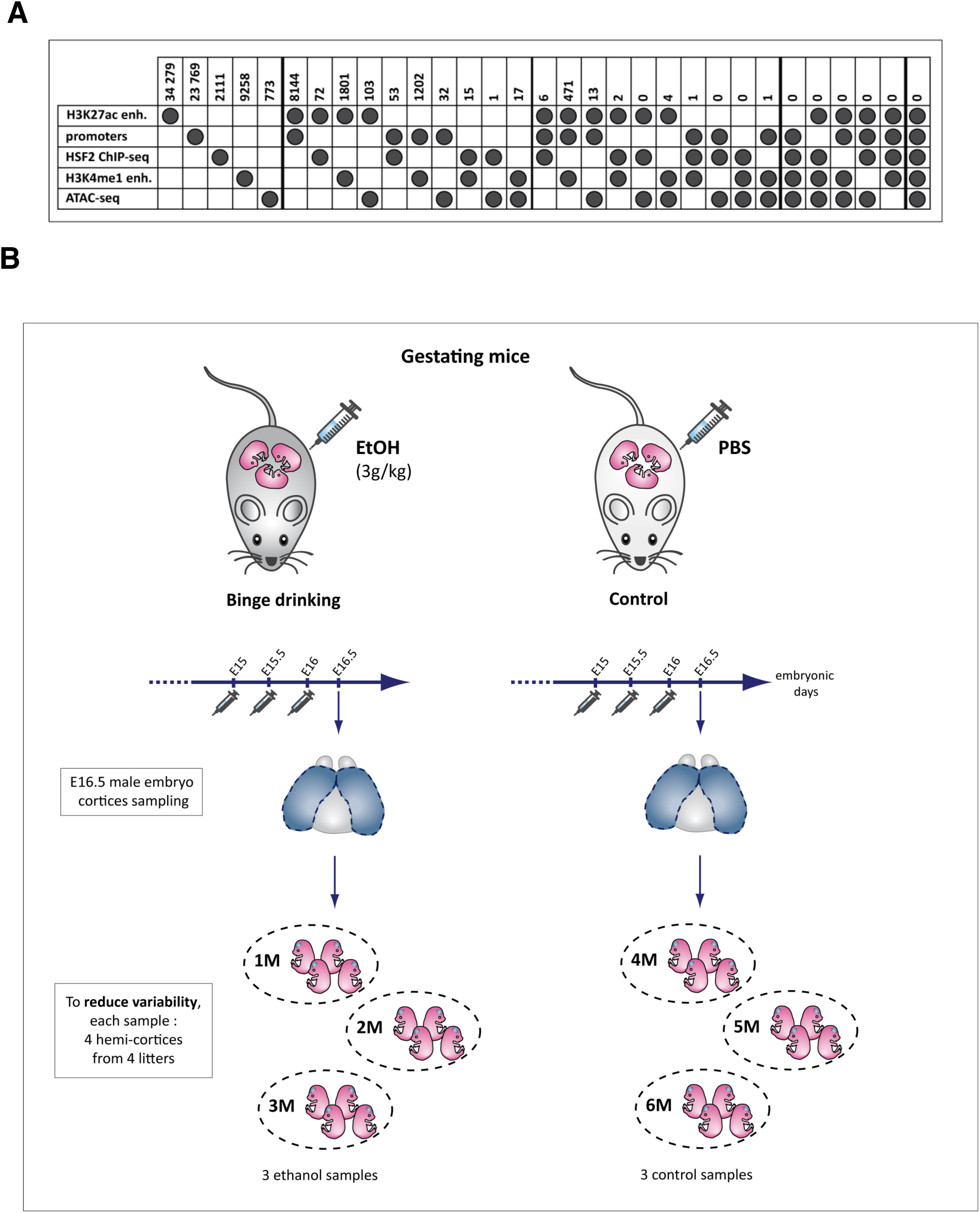
(related to **Figure 1**) **(A) Number of regions in each methylome capture category.** “Other regions” correspond to HSF2 genome-wide occupancy, a stress-responsive transcription factor involved in FASD phenotypes (El Fatimy et al., 2014; ChIP-seq data), and to differentially accessible regions and H3K4me1-marked enhancer regions associated to gene differentially expressed in response to another prenatal stress, neuroinflammation (Schang et al., 2018a; Krishnan et al., 2017, GSE197563; see **Materials and Methods).** Note that categories are not exclusive (*i.e.* some regions are counted several times, *e.g.* 8,144 regions that are both included in H3K27ac enhancers and promoters categories, are also among the 34,279 regions that are included in H3K27ac enhancers). Black dots indicated dataset(s) that are used for the counting. **(B) Sampling protocol.** To minimize intra-group and inter-individual variability, each NGS sample was composed of four hemi-cortices of four different male embryos from four different litters.

**Supplementary Figure 2.**
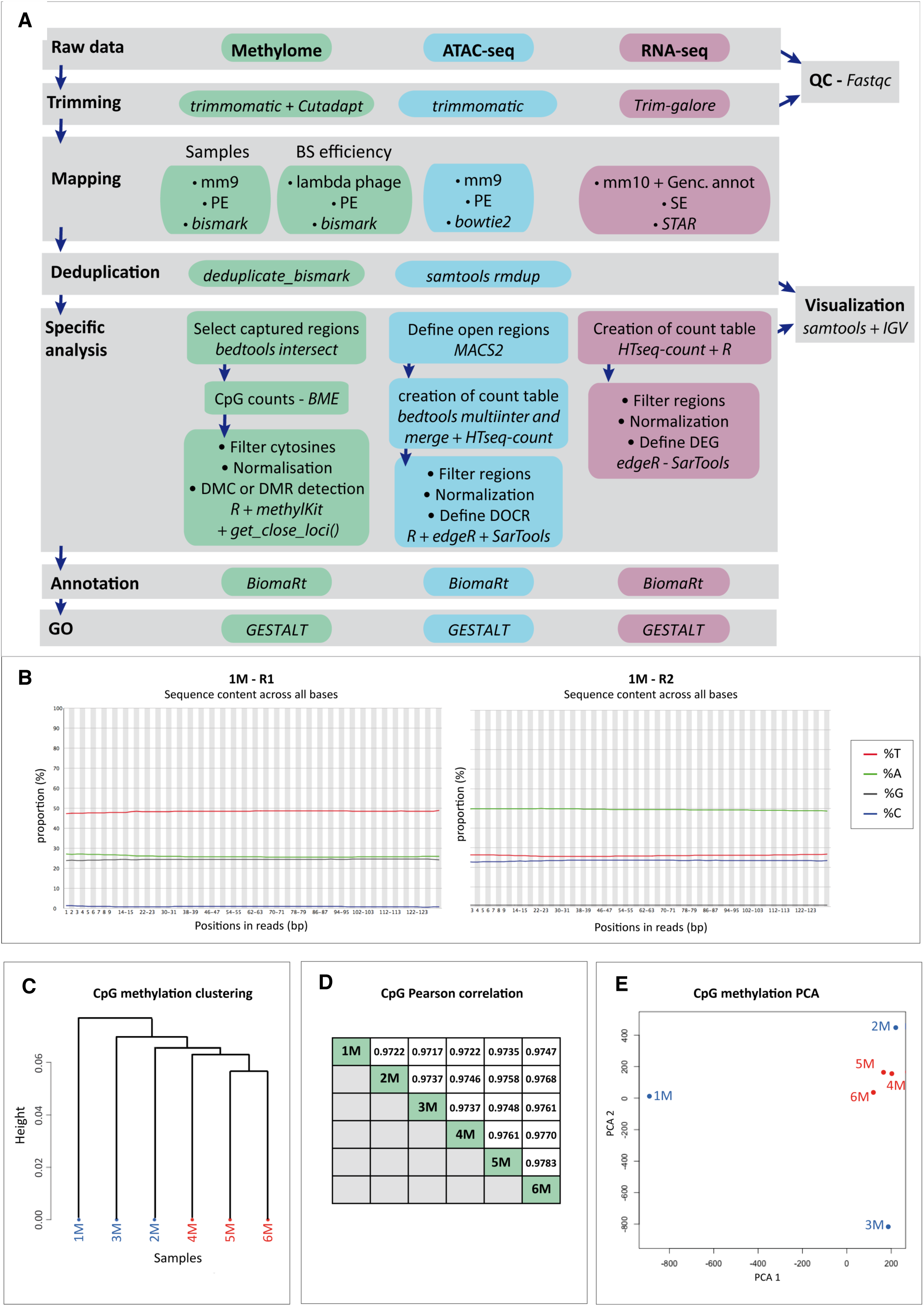
Bioinformatic workflows, quality controls and correlation between methylome samples (related to Figure 2) **(A) Bioinformatics workflows for methylome capture, ATAC-seq and RNA-seq analyses**. Methylome capture were performed in control (PBS-treated) and EtOH-treated groups (described in Figure 1A). ATAC-seq and RNA-seq data mining analyses were performed using publicly available ENCODE datasets, obtained from mice fetal forebrains at distinct developmental stages and newborn samples in physiological conditions. Key steps and tools used for each bioinformatic analysis are described here. For more details, please see **Materials and Methods** and detailed command lines in **Supplementary notebooks 1 to 4**. *QC,* quality control; *PE,* paired-end analysis; *SE,* single-end analysis; *Genc. annot.,* Gencode annotation*; BME:* bismark_methylation_extractor; *DMC*: differentially methylated isolated CpG; *DOCR*: differentially opened or closed regions upon physiological brain development stages; *DEG*, differentially expressed genes upon physiological brain development stages. **(B) *Per-base-sequence-content plots obtained with* FASTQC*. Illustrated for 1M sample.*** An imbalance is observed between bases proportions at each read position as expected after DNA conversion with sodium bisulfite (BS). The conversion of unmethylated cytosines into thymines leads to an over-representation of thymines and an underrepresentation of cytosines at a given position for R1 reads (for R2 reads, which are complementary to R1 ones, an over-representation of adenine and an underrepresentation of guanine are logically observed). These plots were obtained after the trimming of the data. Analyses represented on figures (B,C,D) were done using *MethylKit* (default parameters). CpGs were filtered (CpGs covered by less than 10 reads or having coverage > 99.9^th^ percentile were excluded) and coverage was normalized (median method). Only information from CpG positions that are in all samples are kept (1,259,111 CpGs). **(C) CpG methylation Hierarchical Cluster Analysis.** Dendrogram obtained using of ward method with correlation as distance measure. **(D) Correlation matrix.** Matrix obtained using Pearson correlation. **(E) CpG methylation Principal Components Analysis (PCA)** of Ethanol-treated (1M, 2M, 3M) and control (4M, 5M, 6M) samples.

**Supplementary Figure 3.**
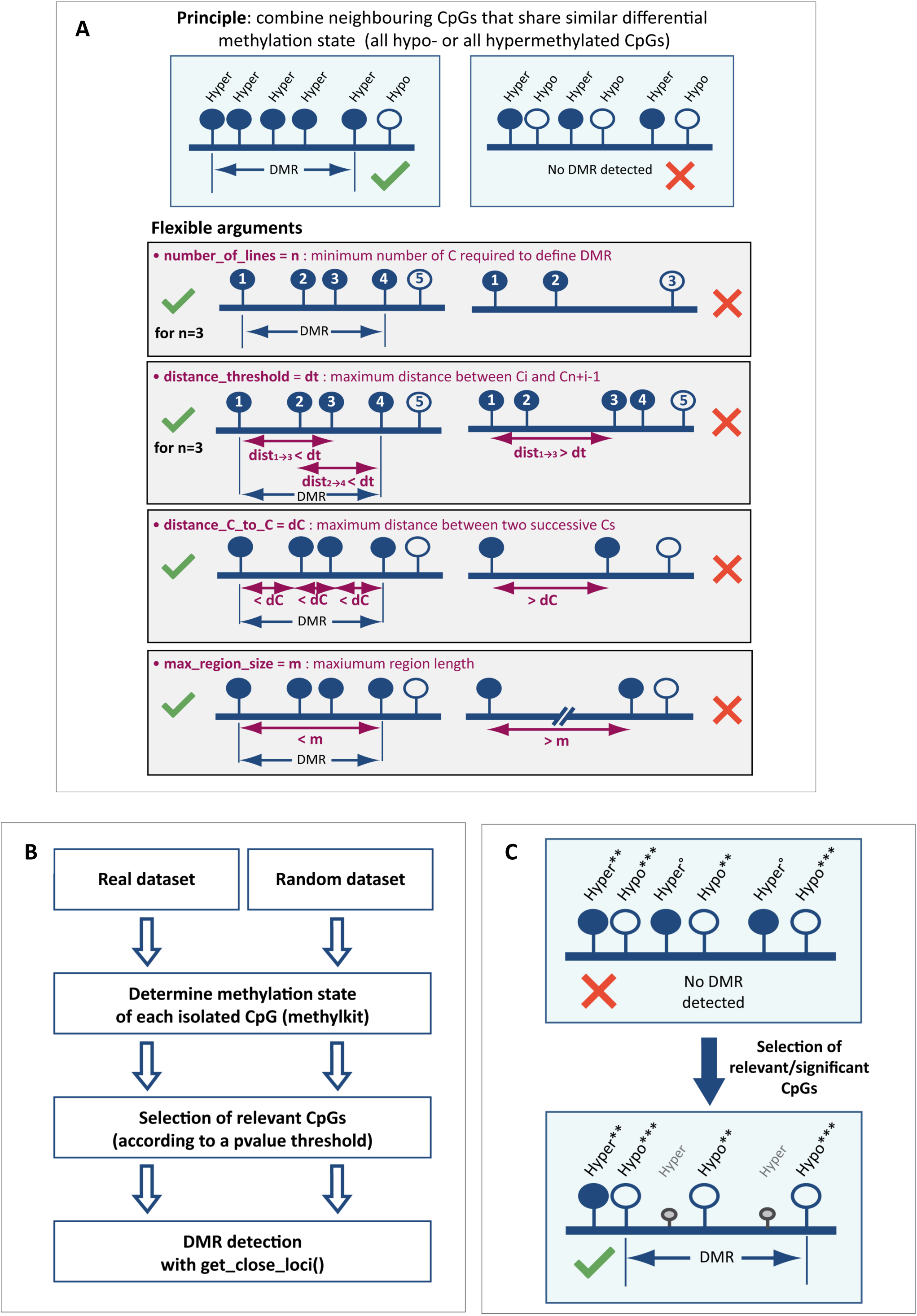
Strategy to detect DMRs (related to Figure 2) **(A) Principle and parameters of get_close_loci() function that we specifically generated for the detection of DMRs into datasets from tailor-made capture.** Filled circles represent hypermethylated cytosines observed between two datasets (*e.g.* control group *versus* test group), whereas empty circles represent hypomethylated cytosines. **(B) Main steps** of the approach performed to detect relevant DMRs in the real dataset from tailor-made methylome capture. **(C) Selection of relevant CpGs is essential for the detection of DMRs**. DMR detection could be compromised if CpGs with no reliable methylation state are kept by the method. In this example, the conservation of the two unreliably defined hypermethylated CpGs during DMRs detection process, prevents the detection of hypomethylated DMR, although this DMR exists. Therefore, the p-value threshold for the methylation status of each CpG must be defined to identify statistically relevant methylation information. Filled circles represent hypermethylated cytosines observed between two datasets (*e.g.* control group *versus* test group), whereas empty circles represent hypomethylated cytosines. *** significant p-value pval < 0.07); small empty circle, non-significant p-value (pval ≥ 0.07).

**Supplementary Figure 4.**
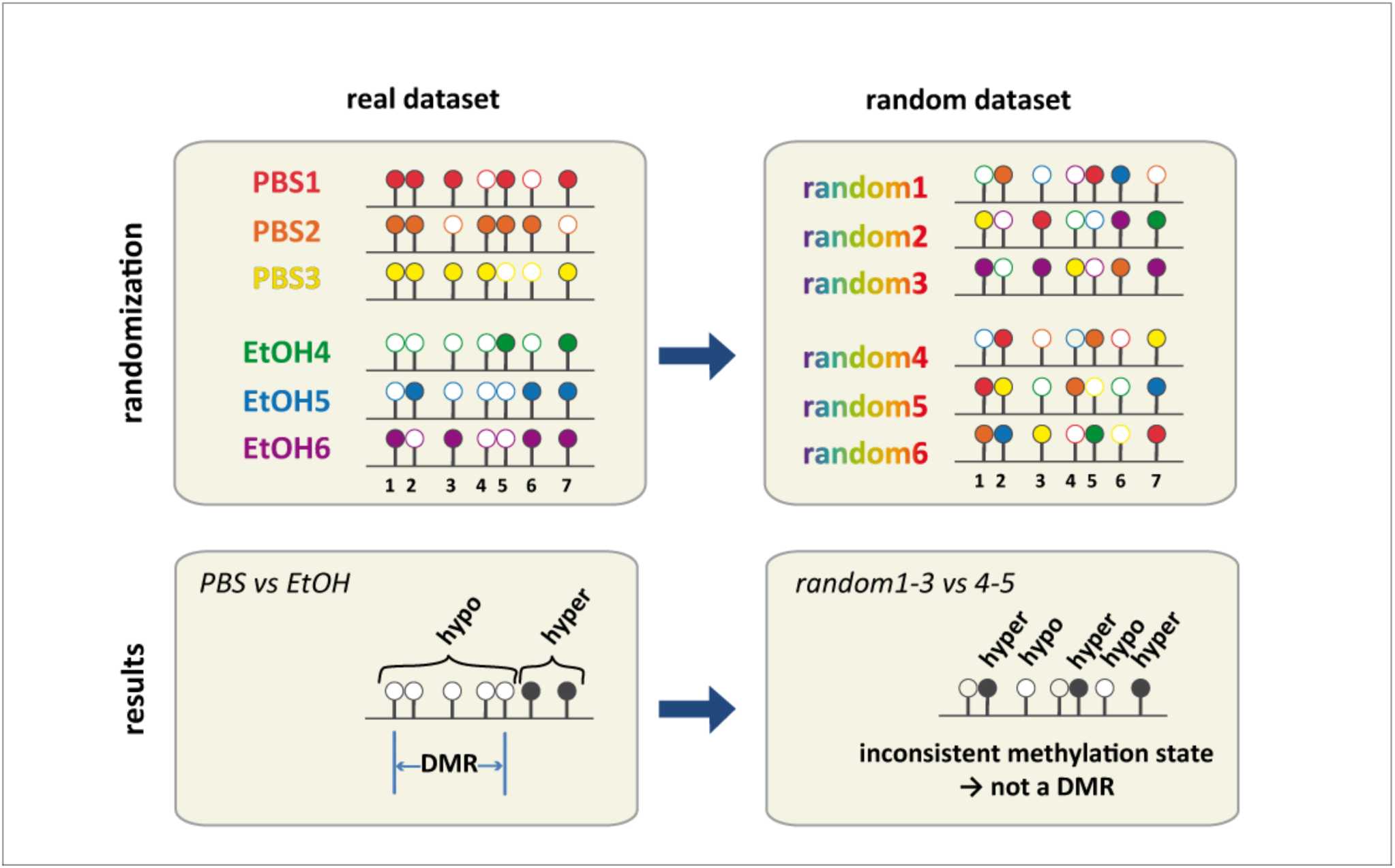
Creation of a random dataset to define relevant parameters for the detection of DMRs. (related to Figure 2) Two random data sets were generated based on our real dataset (composed of 3 PBS-treated control samples, and 3 EtOH-treated ones). In order to randomly redistribute the values without assigning unobserved values to a given CpG position, randomization was performed separately for each cytosine site (position by position, random shuffling of the percent of methylated cytosines observed in real samples). In the upper panel, filled circles represent methylated cytosines, whereas empty circles represent unmethylated cytosines.

**Supplementary Figure 5.**
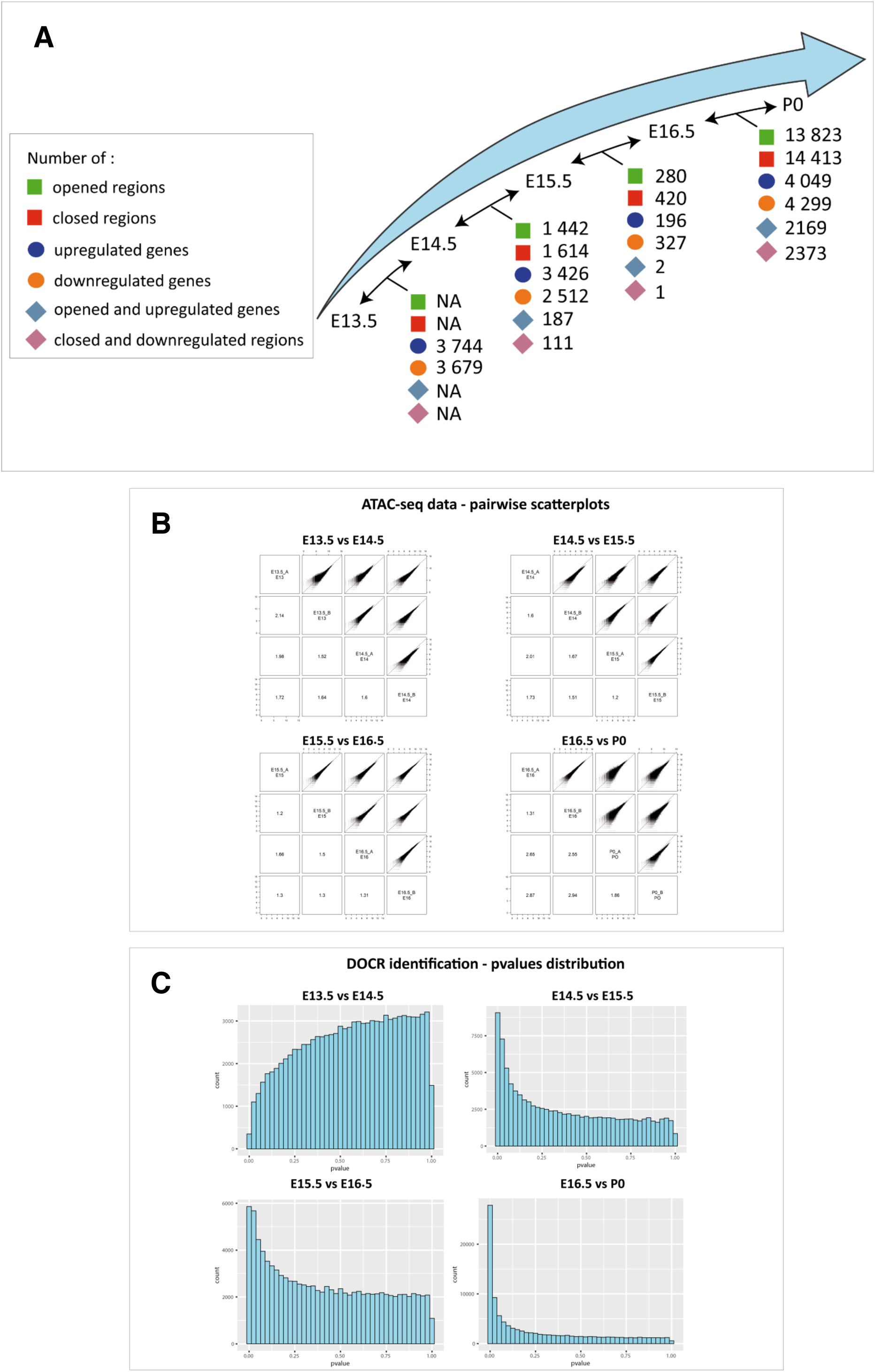
Number of differentially opened or closed regions (DOCRs) or differentially expressed genes (DEGs) identified in the developing brain under physiological conditions. (related to Figure 4) **(A)** Available ENCODE ATAC-seq and RNA-seq data are used to identify differentially opened or closed regions (DOCRs) and differentially expressed genes (DEGs) during brain development, under physiological conditions. DOCRs and DEGs are obtained in a pairwise comparison manner between two successive developmental stages (embryonic days E13.5, E14.5, E15.5, E16.5 and postnatal day P0). No DOCR is detected between E13.5 and E14.5 development stages. Indeed, it is not recommended to use *edgeR* to detect DOCRs between these samples (these data do not follow the statistical model of EdgeR tool), since p-value distribution is not suitable for this comparison (high number of regions with high p-values, see in **Supplementary Figure 5C**, E13.14 DOCR adjusted p-values distribution). Between E15.5 and E16.5 developmental stages, few DOCRs and DEGs are identified in the developing brain under physiological conditions, compared to E14.5 and E15.5 developmental stages comparisons. It can be explained by the quite equivalent inter- and intra-variabilities observed for E15.5 and E16.5 samples, whereas inter-group variability is greater between E14.5 and E15.5 samples (**Supplementary Figure 5B** and **Supplementary Figure 6A**). Many DOCRs and DEGs are identified between E16.5 and P0 stages, compared to other pairwise comparisons. This result is not surprising: since a longer span of time (a few days) separates these two stages (*versus* only one day for the other stage comparisons), it is plausible that more biological events have occurred that can imply chromatin accessibility. This is consistent with scatterplots and *Simple Error Ratio Estimate* (*SERE*) values observed between these samples (**Supplementary Figure 5B** and **Supplementary Figure 6A**). Few regions are both identified as DOCRs and DEGs between each stage, compared to the number of observed DOCRs and DEGs taken separately. DOCRs and DEGs are identified using *edgeR*, without logFC threshold but Benjamini Hochberg p-value adjustment was performed and level of controlled false positive rate was set to 0.05). **(B) Pairwise scatterplots and matrix of pairwise *Simple Error Ratio Estimate* (*SERE*) values for ATAC-seq samples.** Comparisons were performed between two successive developmental stages. *SERE* statistic was used as a similarity index (the more dissimilar the samples are, the higher the SERE value is). Replicates for a given stage, are globally similar, except at E13.5. Indeed, replicates at E13.5 are more dissimilar between each other than to E14.5 samples. This observation could explain why we do not obtain the expected p-values distribution profile for sample comparison (see C) and why identification of DOCRs between these two developmental stages is compromised (see (A)). High inter- and intragroup variabilities are observed for E15.5 and E16.5 samples. This could explain the lower number of DOCRs identified for these sample comparisons, compared to other developmental stages comparisons (Figure 4A). Scatterplots and SERE values were obtained using *SARTools R* package. **(C) Raw p-values distribution obtained for ATAC-seq samples pairwise comparisons allowing DOCR identification.** For all comparisons except E13.5 *versus* E14.5 stages comparison, p-values distribution profiles are as expected (*i.e.* a globally uniform distribution with a peak around 0), meaning that DOCRs could be detected with *edgeR* statistic tool. For E13.5 *versus* E14.5 stages comparison, distribution does not follow the expected profile (see (A)), therefore preventing the use of *edgeR* statistical model for DOCR identification. This result can be due to the high dissimilarity observed between E13.5 replicates (see (B)). These p-value histograms were obtained using *ggplot2 R* package, with data from *SARTools R* package.

**Supplementary Figure 6.**
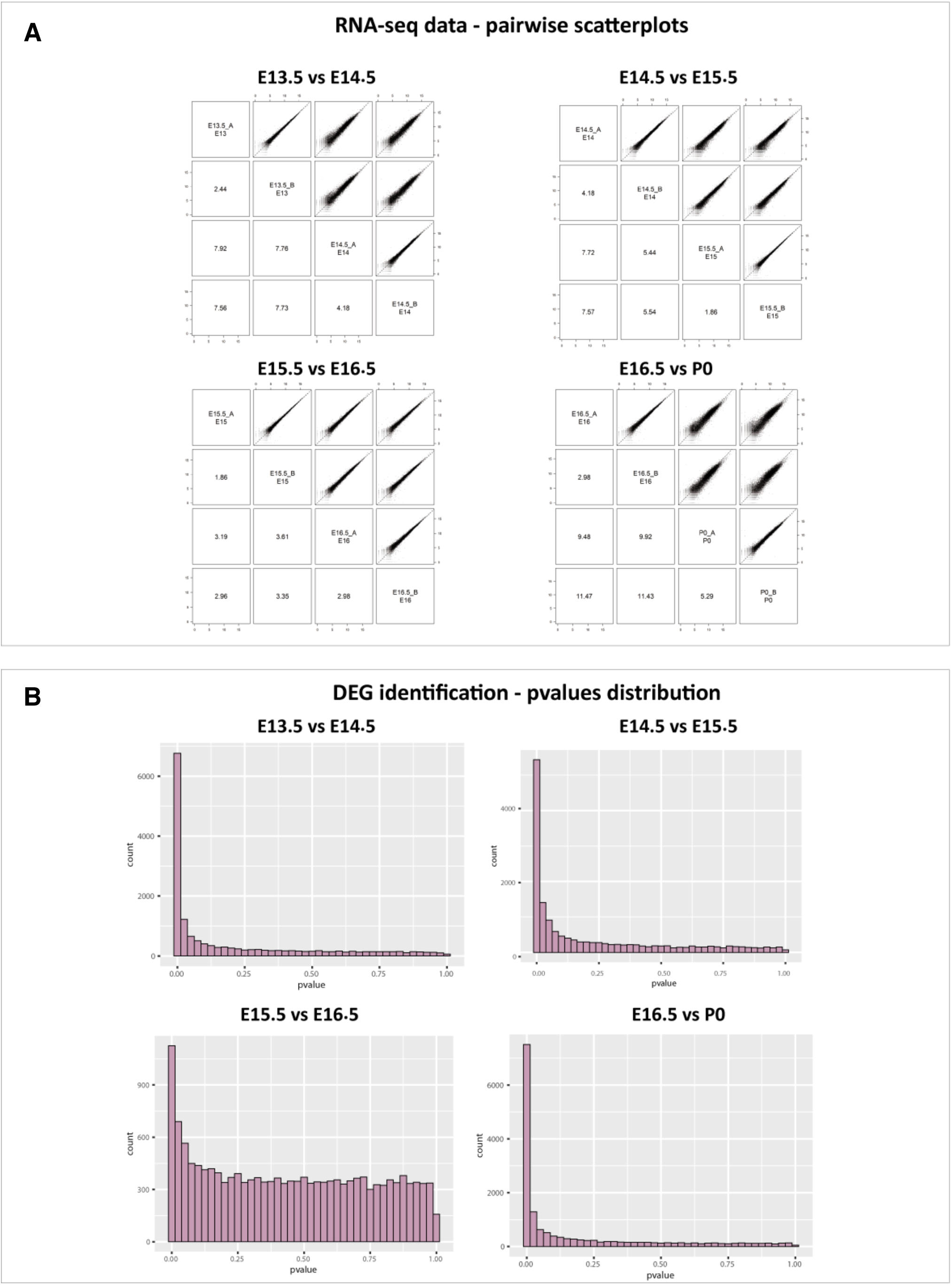
(related to **Figure 4**) **(A) Pairwise scatterplots and matrix of pairwise Simple Error Ratio Estimate (*SERE*) values for RNA-seq samples.** Comparisons were performed between two successive developmental stages. *SERE* statistic is used as a similarity index (the more dissimilar the samples are, the higher the SERE value is). For each pairwise comparison, variability between replicates is lower than those of samples from distinct stages. Nevertheless, variability between replicates is higher than those observed for ATAC-seq data comparisons (**Supplementary Figure 5**). Inter- and intragroup variabilities are nearly equivalent for E15.5 and E16.5 samples. These similarities can explain the lower number of DEGs identified for these sample comparisons, compared to other developmental stages comparisons (**Supplementary Figure 5A**). Scatterplots and SERE values were obtained using *SARTools R* package. **(B) Raw p-values distribution obtained for ATAC-seq samples pairwise comparisons allowing DEG identification.** For all comparisons, p-values distribution profiles are as expected (*i.e.* a globally uniform distribution with a peak around 0), meaning that DEGs can be detected with *edgeR* statistic tool These p-value histograms were obtained using *ggplot2 R* package, with data from *SARTools R* package.

**Supplementary Figure 7.**
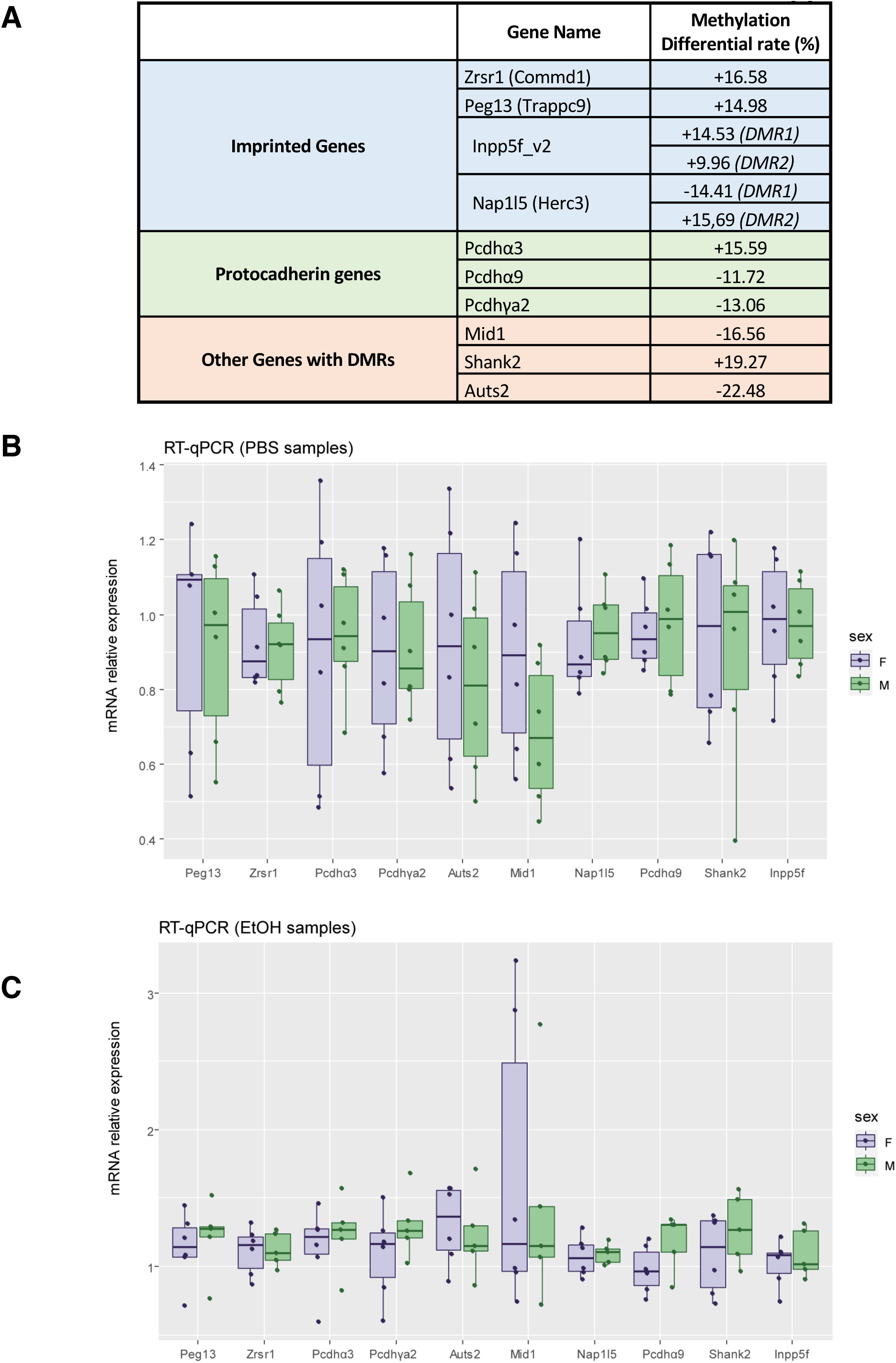
Mono-allelically expressed genes and important neurodevelopmental genes show concomitant perturbations of methylation patterns and alterations of associated gene expression upon PAE. **(A) *Methylation status of monoallelic expression genes (imprinted genes and clustered* Pcdh *genes*) *and some relevant genes for neurodevelopment*** **(B)** *Comparison of mRNA expression **does** no**t reveal** sexual dimorphism **in the expression of these** genes in **fetal** mouse cortices*. (related to Figure 6) Quantitative RT-PCR of genes of interest, in (A) PBS- or (B) EtOH-treated samples. Boxplots showing Relative mRNA levels from E16.5 fetal male or female cortices, upon PBS- (A) or EtOH-treatment (B). Each dot represents calibrated and normalized relative gene expression (CNRQ) value of each sample. mRNA levels are normalized to the expression of *Tpt1* and *Hprt1* genes. Results are generated using n = 6 females per condition (PBS or EtOH) and n = 5 or 6 males (6 for PBS, 5 for EtOH conditions) samples, from 3 distinct litters in each condition. Significant differences between PBS- and EtOH-treated samples were determined using non-parametric Mann-Whitney test. To take into account multiple tests, p-values were corrected into adjusted p-values (padj.), using Benjamini-Hochberg correction (padj. < 0.05 is considered as statistically significant result). For each gene, no statistically significant difference was observed between male and female samples (padj. always higher than 0.05).

**Supplement Figure 8.**
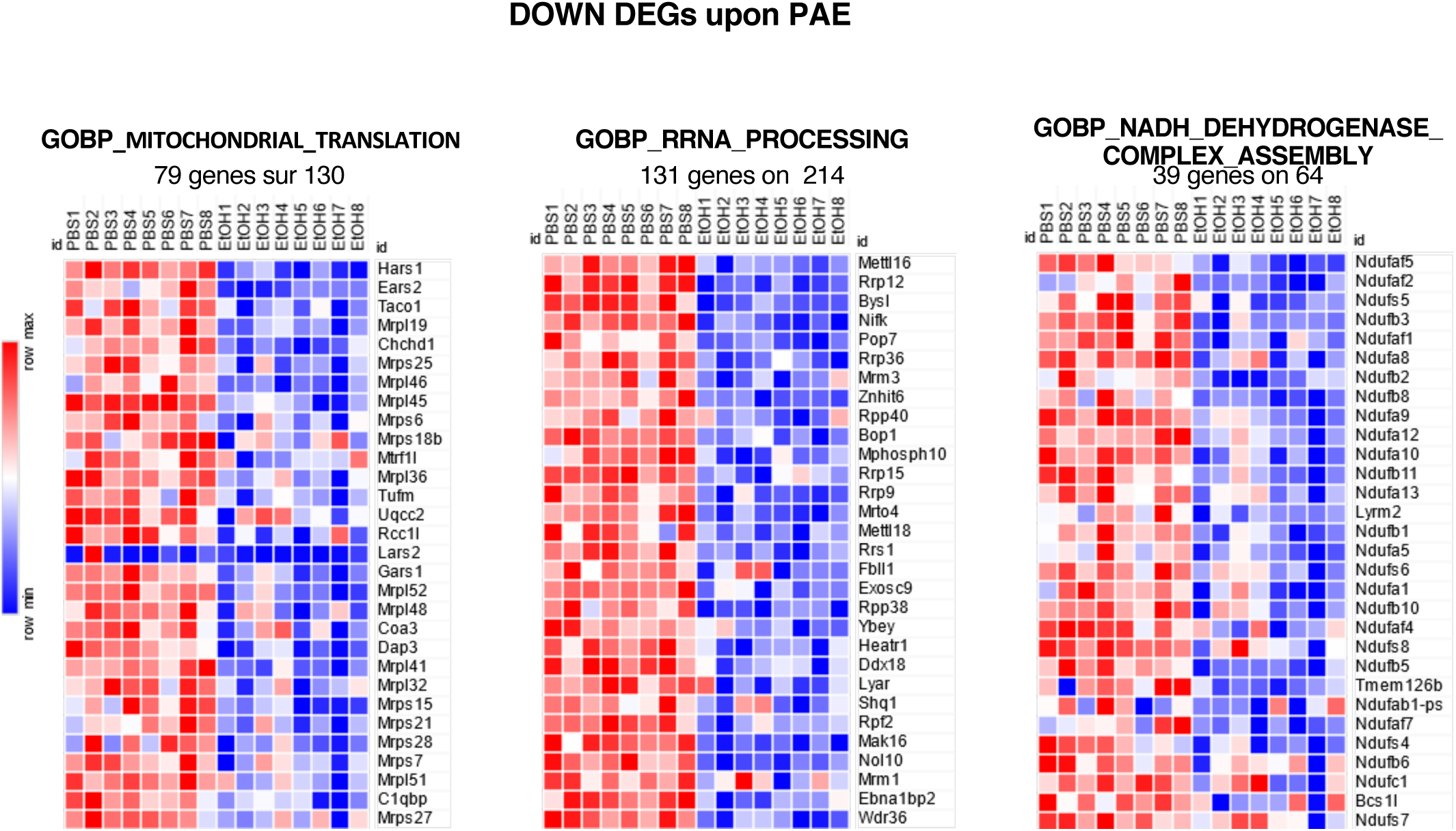
Heatmap representing the level of expression in each sample of significant down regulated genes of selected GO_Biological Process gene sets. The number of DEGs involved and the number of genes in the selected gene set are indicated.

## Notes

### Competing Interest Statement

The authors have declared no competing interest.

### Summary of Updates

This new version of the paper adds to the deciphering of the immediate effects of prenatal alcohol exposure on genome-wide gene expression in mouse prenatal brain cortices, using RNA-seq experiment analysis.

## Bibliography

Afgan, E., Baker, D., van den Beek, M., Blankenberg, D., Bouvier, D., Čech, M., Chilton, J., Clements, D., Coraor, N., Eberhard, C., et al. (2016). The Galaxy platform for accessible, reproducible and collaborative biomedical analyses: 2016 update. Nucleic Acids Res. 44, W3–W10.

Akalin, A., Kormaksson, M., Li, S., Garrett-Bakelman, F.E., Figueroa, M.E., Melnick, A., Mason, C.E., and others (2012). methylKit: a comprehensive R package for the analysis of genome-wide DNA methylation profiles. Genome Biol 13, R87.

Allum, F., Shao, X., Guénard, F., Simon, M.-M., Busche, S., Caron, M., Lambourne, J., Lessard, J., Tandre, K., Hedman, Å.K., et al. (2015). Characterization of functional methylomes by next-generation capture sequencing identifies novel disease-associated variants. Nature Communications 6, 7211.

Anders, S., Pyl, P.T., and Huber, W. (2015). HTSeq--a Python framework to work with high-throughput sequencing data. Bioinformatics 31, 166–169.

Bale, T.L., Baram, T.Z., Brown, A.S., Goldstein, J.M., Insel, T.R., McCarthy, M.M., Nemeroff, C.B., Reyes, T.M., Simerly, R.B., Susser, E.S., et al. (2010). Early Life Programming and Neurodevelopmental Disorders. Biological Psychiatry 68, 314–319.

Barnett, D.W., Garrison, E.K., Quinlan, A.R., Stromberg, M.P., and Marth, G.T. (2011). BamTools: a C++ API and toolkit for analyzing and managing BAM files. Bioinformatics 27, 1691–1692.

Bhat, P., Chow, A., Emert, B., Ettlin, O., Quinodoz, S.A., Strehle, M., Takei, Y., Burr, A., Goronzy, I.N., Chen, A.W., Huang, W., Ferrer, J.L.M., Soehalim, E., Goh, S.-T., Chari, T., Sullivan, D.K., Blanco, M.R., Guttman, M., 2024. Genome organization around nuclear speckles drives mRNA splicing efficiency. Nature 629, 1165–1173.

Bock, C. (2012). Analysing and interpreting DNA methylation data. Nature Reviews Genetics 13, 705–719.

Bolger, A.M., Lohse, M., and Usadel, B. (2014). Trimmomatic: a flexible trimmer for Illumina sequence data. Bioinformatics 30, 2114–2120.

Bourgeron, T. (2015). From the genetic architecture to synaptic plasticity in autism spectrum disorder. Nat. Rev. Neurosci. 16, 551–563.

Burd, L., Cotsonas-Hassler, T.M., Martsolf, J.T., and Kerbeshian, J. (2003). Recognition and management of fetal alcohol syndrome. Neurotoxicology and Teratology 25, 681–688.

Bustin, S.A., Benes, V., Garson, J.A., Hellemans, J., Huggett, J., Kubista, M., Mueller, R., Nolan, T., Pfaffl, M.W., Shipley, G.L., et al. (2009). The MIQE Guidelines: Minimum Information for Publication of Quantitative Real-Time PCR Experiments. Clinical Chemistry 55, 611–622.

Cahill, M.E., Xie, Z., Day, M., Photowala, H., Barbolina, M.V., Miller, C.A., Weiss, C., Radulovic, J., Sweatt, J.D., Disterhoft, J.F., et al. (2009). Kalirin regulates cortical spine morphogenesis and disease-related behavioral phenotypes. Proc. Natl. Acad. Sci. U.S.A. 106, 13058–13063.

Carloni, S., Mazzoni, E., and Balduini, W. (2004). Caspase-3 and calpain activities after acute and repeated ethanol administration during the rat brain growth spurt. J Neurochem 89, 197– 203.

Chater-Diehl, E.J., Laufer, B.I., Singh, S.M. (2017) Changes to histone modifications following prenatal alcohol exposure: An emerging picture. Alcohol 60, 41–52. PMID: 28431792

Clark, S.J., Statham, A., Stirzaker, C., Molloy, P.L., and Frommer, M. (2006). DNA methylation: bisulphite modification and analysis. Nat Protoc 1, 2353–2364.

Cobben, J.M., Krzyzewska, I.M., Venema, A., Mul, A.N., Polstra, A., Postma, A.V., Smigiel, R., Pesz, K., Niklinski, J., Chomczyk, M.A., et al. (2019). DNA methylation abundantly associates with fetal alcohol spectrum disorder and its subphenotypes. Epigenomics 11, 767– 785.

Davies, W., Isles, A.R., Humby, T., and Wilkinson, L.S. (2008). What are imprinted genes doing in the brain? Adv. Exp. Med. Biol. 626, 62–70.

Dent, C.L., and Isles, A.R. (2014). Brain-expressed imprinted genes and adult behaviour: the example of Nesp and Grb10. Mamm. Genome 25, 87–93.

Desroches Altamirano, C., Kang, M.-K., Jordan, M.A., Borianne, T., Dilmen, I., Gnädig, M., von Appen, A., Honigmann, A., Franzmann, T.M., Alberti, S., 2024. eIF4F is a thermo-sensing regulatory node in the translational heat shock response. Mol Cell 84, 1727–1741.

Devi, R., Pelletier, L., Prosser, S.L., 2021. Charting the complex composite nature of centrosomes, primary cilia and centriolar satellites. Curr Opin Struct Biol 66, 32–40.

Dobin, A., and Gingeras, T.R. (2015). Mapping RNA-seq Reads with STAR: Mapping RNA-seq Reads with STAR. In Current Protocols in Bioinformatics, A. Bateman, W.R. Pearson, L.D. Stein, G.D. Stormo, and J.R. Yates, eds. (Hoboken, NJ, USA: John Wiley & Sons, Inc.), p. 11.14.1-11.14.19.

Dobin, A., Davis, C.A., Schlesinger, F., Drenkow, J., Zaleski, C., Jha, S., Batut, P., Chaisson, M., and Gingeras, T.R. (2013). STAR: ultrafast universal RNA-seq aligner. Bioinformatics 29, 15–21.

Doncheva, N.T., Morris, J.H., Gorodkin, J., Jensen, L.J., 2019. Cytoscape StringApp: Network Analysis and Visualization of Proteomics Data. J. Proteome Res. 18, 623–632.

Downing, C., Johnson, T.E., Larson, C., Leakey, T.I., Siegfried, R.N., Rafferty, T.M., and Cooney, C.A. (2011). Subtle decreases in DNA methylation and gene expression at the mouse Igf2 locus following prenatal alcohol exposure: effects of a methyl-supplemented diet. Alcohol 45, 65–71.

Durinck, S., Spellman, P.T., Birney, E., and Huber, W. (2009). Mapping identifiers for the integration of genomic datasets with the R/Bioconductor package biomaRt. Nat Protoc 4, 1184–1191.

El Fatimy, R., Miozzo, F., Le Mouel, A., Abane, R., Schwendimann, L., Saberan-Djoneidi, D., de Thonel, A., Massaoudi, I., Paslaru, L., Hashimoto-Torii, K., et al. (2014). Heat shock factor 2 is a stress-responsive mediator of neuronal migration defects in models of fetal alcohol syndrome. EMBO Mol Med 6, 1043–1061.

El Hajj, N., Dittrich, M., Böck, J., Kraus, T.F.J., Nanda, I., Müller, T., Seidmann, L., Tralau, T., Galetzka, D., Schneider, E., et al. (2016). Epigenetic dysregulation in the developing Down syndrome cortex. Epigenetics 11, 563–578.

Gentleman, R.C., Carey, V.J., Bates, D.M., Bolstad, B., Dettling, M., Dudoit, S., Ellis, B., Gautier, L., Ge, Y., Gentry, J., et al. (2004). Bioconductor: open software development for computational biology and bioinformatics. Genome Biol 5, R80.

Gibbard, W.B., Wass, P., and Clarke, M.E. (2003). The neuropsychological implications of prenatal alcohol exposure. The Canadian Child and Adolescent Psychiatry Review.

Gräff, J., Kim, D., Dobbin, M.M., and Tsai, L.-H. (2011). Epigenetic Regulation of Gene Expression in Physiological and Pathological Brain Processes. Physiological Reviews 91, 603–649.

Guerri, C., Bazinet, A., and Riley, E.P. (2009). Foetal Alcohol Spectrum Disorders and Alterations in Brain and Behaviour. Alcohol and Alcoholism 44, 108–114.

Guo, J.U., Ma, D.K., Mo, H., Ball, M.P., Jang, M.-H., Bonaguidi, M.A., Balazer, J.A., Eaves, H.L., Xie, B., Ford, E., et al. (2011). Neuronal activity modifies the DNA methylation landscape in the adult brain. Nat Neurosci 14, 1345–1351.

Hashimoto-Torii, K., Kawasawa, Y.I., Kuhn, A., and Rakic, P. (2011). Combined transcriptome analysis of fetal human and mouse cerebral cortex exposed to alcohol. Proceedings of the National Academy of Sciences 108, 4212–4217.

Hashimoto-Torii, K., Torii, M., Fujimoto, M., Nakai, A., El Fatimy, R., Mezger, V., Ju, M.J., Ishii, S., Chao, S., Brennand, K.J., et al. (2014). Roles of Heat Shock Factor 1 in Neuronal Response to Fetal Environmental Risks and Its Relevance to Brain Disorders. Cell Neuron 82, 560–572.

Haycock, P.C. (2009). Fetal Alcohol Spectrum Disorders: The Epigenetic Perspective. Biol. Reprod. 81, 607–617.

Haycock, P.C., and Ramsay, M. (2009). Exposure of Mouse Embryos to Ethanol During Preimplantation Development: Effect on DNA Methylation in the H19 Imprinting Control Region. Biology of Reproduction 81, 618–627.

Heinz, S., Benner, C., Spann, N., Bertolino, E., Lin, Y.C., Laslo, P., Cheng, J.X., Murre, C., Singh, H., and Glass, C.K. (2010). Simple combinations of lineage-determining transcription factors prime cis-regulatory elements required for macrophage and B cell identities. Mol. Cell 38, 576–589.

Hellemans, K.G.C., Sliwowska, J.H., Verma, P., and Weinberg, J. (2010). Prenatal alcohol exposure: Fetal programming and later life vulnerability to stress, depression and anxiety disorders. Neuroscience & Biobehavioral Reviews 34, 791–807.

Honda, A., Usui, H., Sakimura, K., and Igarashi, M. (2017). Rufy3 is an adapter protein for small GTPases that activates a Rac guanine nucleotide exchange factor to control neuronal polarity. J. Biol. Chem. 292, 20936–20946.

Huang, D.W., Sherman, B.T., and Lempicki, R.A. (2009). Systematic and integrative analysis of large gene lists using DAVID bioinformatics resources. Nat Protoc 4, 44–57.

Hutson, J.R., Stade, B., Lehotay, D.C., Collier, C.P., Kapur, B.M. (2012) Folic acid transport to the human fetus is decreased in pregnancies with chronic alcohol exposure. PLoS One 7, e38057.

Ikonomidou, C., Bittigau, P., Ishimaru, M.J., Wozniak, D.F., Koch, C., Genz, K., Price, M.T., Stefovska, V., Hörster, F., Tenkova, T., et al. (2000). Ethanol-induced apoptotic neurodegeneration and fetal alcohol syndrome. Science 287, 1056–1060.

Ishii, S., Torii, M., Son, A.I., Rajendraprasad, M., Morozov, Y.M., Kawasawa, Y.I., Salzberg, A.C., Fujimoto, M., Brennand, K., Nakai, A., et al. (2017). Variations in brain defects result from cellular mosaicism in the activation of heat shock signalling. Nature Communications 8, 15157.

Jones, K.L., and Smith, D.W. (1973). Recognition of the fetal alcohol syndrome in early infancy. Lancet 302, 999–1001.

Kaminen-Ahola, N., Ahola, A., Maga, M., Mallitt, K.-A., Fahey, P., Cox, T.C., Whitelaw, E., and Chong, S. (2010). Maternal Ethanol Consumption Alters the Epigenotype and the Phenotype of Offspring in a Mouse Model. PLoS Genetics 6, e1000811.

Kashima, R., Hata, A., 2018. The role of TGF-β superfamily signaling in neurological disorders. Acta Biochim Biophys Sin (Shanghai) 50, 106–120.

Khalid, O., Kim, J.J., Kim, H.-S., Hoang, M., Tu, T.G., Elie, O., Lee, C., Vu, C., Horvath, S., Spigelman, I., et al. (2014). Gene expression signatures affected by alcohol-induced DNA methylomic deregulation in human embryonic stem cells. Stem Cell Research 12, 791–806.

Kim, K.Q., Zaher, H.S., 2022. Canary in a coal mine: collided ribosomes as sensors of cellular conditions. Trends Biochem Sci 47, 82–97.

Kleiber, M.L., Mantha, K., Stringer, R.L., and Singh, S.M. (2013). Neurodevelopmental alcohol exposure elicits long-term changes to gene expression that alter distinct molecular pathways dependent on timing of exposure. Journal of Neurodevelopmental Disorders 5, 1.

Kleiber, M.L., Diehl, E.J., Laufer, B.I., Mantha, K., Chokroborty-Hoque, A., Alberry, B., and Singh, S.M. (2014). Long-term genomic and epigenomic dysregulation as a consequence of prenatal alcohol exposure: a model for fetal alcohol spectrum disorders. Frontiers in Genetics 5.

Kodituwakku, P.W. (2007). Defining the behavioral phenotype in children with fetal alcohol spectrum disorders: a review. Neurosci Biobehav Rev 31, 192–201.

Krishnan, M.L., Van Steenwinckel, J., Schang, A.-L., Yan, J., Arnadottir, J., Le Charpentier, T., Csaba, Z., Dournaud, P., Cipriani, S., Auvynet, C., et al. (2017). Integrative genomics of microglia implicates DLG4 (PSD95) in the white matter development of preterm infants. Nat Commun 8, 428.

Krueger, F., and Andrews, S.R. (2011). Bismark: a flexible aligner and methylation caller for Bisulfite-Seq applications. Bioinformatics 27, 1571–1572.

Krueger, F., Kreck, B., Franke, A., and Andrews, S.R. (2012). DNA methylome analysis using short bisulfite sequencing data. Nature Methods 9, 145–151.

Langmead, B., and Salzberg, S.L. (2012). Fast gapped-read alignment with Bowtie 2. Nat Methods 9, 357–359.

LaSalle, J.M., Powell, W.T., and Yasui, D.H. (2013). Epigenetic layers and players underlying neurodevelopment. Trends in Neurosciences 36, 460–470.

Laufer, B.I., Mantha, K., Kleiber, M.L., Diehl, E.J., Addison, S.M.F., and Singh, S.M. (2013). Long-lasting alterations to DNA methylation and ncRNAs could underlie the effects of fetal alcohol exposure in mice. Dis Model Mech 6, 977–992.

Laufer, B.I., Kapalanga, J., Castellani, C.A., Diehl, E.J., Yan, L., and Singh, S.M. (2015). Associative DNA methylation changes in children with prenatal alcohol exposure. Epigenomics 7, 1259–1274.

Laufer BI, Chater-Diehl EJ, Kapalanga J, Singh SM. (2017) Long-term alterations to DNA methylation as a biomarker of prenatal alcohol exposure: From mouse models to human children with fetal alcohol spectrum disorders. Alcohol 60, 67–75. PMID: 28187949.

Legault, L.M., Doiron, K., Breton-Larrivée, M., Langford-Avelar, A., Lemieux, A., Caron, M., Jerome-Majewska, L.A., Sinnett, D., and McGraw, S. (2021). Pre-implantation alcohol exposure induces lasting sex-specific DNA methylation programming errors in the developing forebrain. Clin Epigenetics 13, 164.

Lemoine, P., Harousseau, H., Borteyru, J.P., and Menuet, J.C. (1968). Les enfants de parents alcooliques. Anomalies observées. A propos de 127 cas.

Li, H., Handsaker, B., Wysoker, A., Fennell, T., Ruan, J., Homer, N., Marth, G., Abecasis, G., Durbin, R., and others (2009). The sequence alignment/map format and SAMtools. Bioinformatics 25, 2078–2079.

Liao, Y., Wang, J., Jaehnig, E.J., Shi, Z., and Zhang, B. (2019). WebGestalt 2019: gene set analysis toolkit with revamped UIs and APIs. Nucleic Acids Research 47, W199–W205.

Light, S.E.W., and Jontes, J.D. (2017). δ-Protocadherins: Organizers of neural circuit assembly. Seminars in Cell & Developmental Biology 69, 83–90.

Lindquist, S., 1981. Regulation of protein synthesis during heat shock. Nature 293, 311–314.

Liu, Y., Balaraman, Y., Wang, G., Nephew, K.P., and Zhou, F.C. (2009). Alcohol exposure alters DNA methylation profiles in mouse embryos at early neurulation. Epigenetics 4, 500– 511.

Lussier, A.A., Weinberg, J., and Kobor, M.S. (2017). Epigenetics studies of fetal alcohol spectrum disorder: where are we now? Epigenomics 9, 291–311.

Lussier, A.A., Morin, A.M., MacIsaac, J.L., Salmon, J., Weinberg, J., Reynolds, J.N., Pavlidis, P., Chudley, A.E., and Kobor, M.S. (2018). DNA methylation as a predictor of fetal alcohol spectrum disorder. Clin Epigenet 10, 5.

Lussier, A.A., Bodnar, T.S., Moksa, M., Hirst, M., Kobor, M.S., and Weinberg, J. (2021). Prenatal Adversity Alters the Epigenetic Profile of the Prefrontal Cortex: Sexually Dimorphic Effects of Prenatal Alcohol Exposure and Food-Related Stress. Genes (Basel) 12, 1773.

Martin, M. (2011). Cutadapt removes adapter sequences from high-throughput sequencing reads. EMBnet j. 17, 10.

Matsunaga, Y., Noda, M., Murakawa, H., Hayashi, K., Nagasaka, A., Inoue, S., Miyata, T., Miura, T., Kubo, K.-I., and Nakajima, K. (2017). Reelin transiently promotes N-cadherin-dependent neuronal adhesion during mouse cortical development. Proc. Natl. Acad. Sci. U.S.A. 114, 2048–2053.

Mattson, S.N., Crocker, N., and Nguyen, T.T. (2011). Fetal Alcohol Spectrum Disorders: Neuropsychological and Behavioral Features. Neuropsychology Review 21, 81–101.

Miozzo, F., Arnould, H., de Thonel, A., Schang, A.L., Sabéran-Djoneidi, D., Baudry, A., Schneider, B., Mezger, V. (2018) Alcohol exposure promotes DNA methyltransferase DNMT3A upregulation through reactive oxygen species-dependent mechanisms. Cell Stress Chaperones. 23,115–126. PMID: 28712054

Molumby, M.J., Anderson, R.M., Newbold, D.J., Koblesky, N.K., Garrett, A.M., Schreiner, D., Radley, J.J., and Weiner, J.A. (2017). γ-Protocadherins Interact with Neuroligin-1 and Negatively Regulate Dendritic Spine Morphogenesis. Cell Rep 18, 2702–2714.

Morris JH, Apeltsin L, Newman AM, Baumbach J, Wittkop T, Su G, Bader GD, Ferrin TE. clusterMaker: a multi-algorithm clustering plugin for Cytoscape. BMC Bioinformatics. 2011 Nov 9;12:436.. PMID: 22070249; PMCID: PMC3262844

O’Connor, M.J., and Paley, B. (2009). Psychiatric conditions associated with prenatal alcohol exposure. Developmental Disabilities Research Reviews 15, 225–234.

Odabasi, E., Batman, U., Firat-Karalar, E.N., 2020. Unraveling the mysteries of centriolar satellites: time to rewrite the textbooks about the centrosome/cilium complex. Mol Biol Cell 31, 866–872.

Olney, J.W., Tenkova, T., Dikranian, K., Qin, Y.-Q., Labruyere, J., and Ikonomidou, C. (2002). Ethanol-induced apoptotic neurodegeneration in the developing C57BL/6 mouse brain. Brain Res. Dev. Brain Res. 133, 115–126.

Pachinger, C., Dobbelaere, J., Rumpf-Kienzl, C., Raina, S., Garcia-Baucells, J., Sarantseva, M., Brauneis, A., Dammermann, A., 2024. Centriolar satellites are sites of translation of centrosomal and ciliary proteins. 10.1101/2024.02.22.581531

Paley, B., O’Connor, M.J. (2011). Behavioral interventions for children and adolescents with fetal alcohol spectrum disorders. Alcohol Res Health. 34, 64–75. PMID: 23580043

Pang, W., Yi, X., Li, L., Liu, L., Xiang, W., and Xiao, L. (2021). Untangle the Multi-Facet Functions of Auts2 as an Entry Point to Understand Neurodevelopmental Disorders. Front Psychiatry 12, 580433.

Perez, J.D., Rubinstein, N.D., and Dulac, C. (2016). New Perspectives on Genomic Imprinting, an Essential and Multifaceted Mode of Epigenetic Control in the Developing and Adult Brain. Annu. Rev. Neurosci. 39, 347–384.

Phillips, G.R., LaMassa, N., and Nie, Y.M. (2017). Clustered protocadherin trafficking. Seminars in Cell & Developmental Biology 69, 131–139.

Pignataro, L., Varodayan, F.P., Tannenholz, L.E., and Harrison, N.L. (2009). The regulation of neuronal gene expression by alcohol. Pharmacology & Therapeutics 124, 324–335.

Portales-Casamar, E., Lussier, A.A., Jones, M.J., MacIsaac, J.L., Edgar, R.D., Mah, S.M., Barhdadi, A., Provost, S., Lemieux-Perreault, L.P., Cynader, M.S., Chudley, A., Dubé, M.P., Reynolds, J.N., Pavlidis, P., and Kobor, M.S. (2016). DNA methylation signature of human fetal alcohol spectrum disorder. Epigenetics & chromatin 9, 25.

Popova, S., Lange, S., Burd, L., and Rehm, J. (2012). Health Care Burden and Cost Associated with Fetal Alcohol Syndrome: Based on Official Canadian Data. PLoS ONE 7, e43024.

Popova, S., Lange, S., Shield, K., Mihic, A., Chudley, A.E., Mukherjee, R.A.S., Bekmuradov, D., and Rehm, J. (2016). Comorbidity of fetal alcohol spectrum disorder: a systematic review and meta-analysis. The Lancet 387, 978–987.

Quinlan, A.R., and Hall, I.M. (2010). BEDTools: a flexible suite of utilities for comparing genomic features. Bioinformatics 26, 841–842.

Remmers, C., Sweet, R.A., and Penzes, P. (2014). Abnormal kalirin signaling in neuropsychiatric disorders. Brain Research Bulletin 103, 29–38.

Riggs, C.L., Kedersha, N., Ivanov, P., Anderson, P., 2020. Mammalian stress granules and P bodies at a glance. J Cell Sci 133, jcs242487. 10.1242/jcs.242487

Rim, E.Y., Clevers, H., Nusse, R., 2022. The Wnt Pathway: From Signaling Mechanisms to Synthetic Modulators. Annu Rev Biochem 91, 571–598

Robinson, J.T., Thorvaldsdóttir, H., Winckler, W., Guttman, M., Lander, E.S., Getz, G., and Mesirov, J.P. (2011). Integrative genomics viewer. Nat Biotechnol 29, 24–26.

Robinson, M.D., McCarthy, D.J., and Smyth, G.K. (2010). edgeR: a Bioconductor package for differential expression analysis of digital gene expression data. Bioinformatics 26, 139– 140.

Saxonov, S., Berg, P., and Brutlag, D.L. (2006). A genome-wide analysis of CpG dinucleotides in the human genome distinguishes two distinct classes of promoters. Proceedings of the National Academy of Sciences 103, 1412–1417.

Schang, A.-L., Steenwinckel, J. van, Lipecki, J., Rich-Griffin, C., Woolley-Allen, K., Dyer, N., Charpentier, T.L., Schäfer, P., Fleiss, B., Ott, S., et al. (2018a). Epigenome and transcriptome landscapes highlight dual roles of proinflammatory players in a perinatal model of white matter injury (Developmental Biology).

Schang, A.-L., Sabéran-Djoneidi, D., and Mezger, V. (2018b). The impact of epigenomic next-generation sequencing approaches on our understanding of neuropsychiatric disorders. Clin Genet 93, 467–480.

Shannon, P., Markiel, A., Ozier, O., Baliga, N.S., Wang, J.T., Ramage, D., Amin, N., Schwikowski, B., Ideker, T., 2003. Cytoscape: A Software Environment for Integrated Models of Biomolecular Interaction Networks. Genome Res. 13, 2498–2504.

Sugimoto, M., and Abe, K. (2007). X Chromosome Reactivation Initiates in Nascent Primordial Germ Cells in Mice. PLoS Genet 3, e116.

The R Core Team (2018). R - A Language and Environment for statistical computing. R Foundation for Statistical Computing, Vienna, Austria.

Varet, H., Brillet-Guéguen, L., Coppée, J.-Y., and Dillies, M.-A. (2016). SARTools: A DESeq2- and EdgeR-Based R Pipeline for Comprehensive Differential Analysis of RNA-Seq Data. PLoS ONE 11, e0157022.

van der Werf, I.M., Kooy, R.F., and Vandeweyer, G. (2015). A robust protocol to increase NimbleGen SeqCap EZ multiplexing capacity to 96 samples. PloS One 10, e0123872.

Winter, J., Basilicata, M.F., Stemmler, M.P., and Krauss, S. (2016). The MID1 protein is a central player during development and in disease. Front Biosci (Landmark Ed) 21, 664–682.

Xu, J., Pope, S.D., Jazirehi, A.R., Attema, J.L., Papathanasiou, P., Watts, J.A., Zaret, K.S., Weissman, I.L., and Smale, S.T. (2007). Pioneer factor interactions and unmethylated CpG dinucleotides mark silent tissue-specific enhancers in embryonic stem cells. Proceedings of the National Academy of Sciences 104, 12377–12382.

Yamagata, M., Duan, X., and Sanes, J.R. (2018). Cadherins Interact With Synaptic Organizers to Promote Synaptic Differentiation. Front. Mol. Neurosci. 11, 142.

Zhang, Y., Liu, T., Meyer, C.A., Eeckhoute, J., Johnson, D.S., Bernstein, B.E., Nusbaum, C., Myers, R.M., Brown, M., Li, W., et al. (2008). Model-based analysis of ChIP-Seq (MACS). Genome Biol. 9, R137.

