## Supplementary material for "Immediate perturbations of DNA methylation and transcriptome upon acute prenatal alcohol exposure in the mouse developing brain cortex": Supp Table S1 Bisulfite conversion efficiency

| | | 1M | 2M | 3M | 4M | 5M | 6M |
|---|---|---|---|---|---|---|---|
| **deduplication** | Total number of mapped reads | 22 055 | 63 017 | 52 100 | 62 391 | 80 753 | 86 277 |
| | Total count of deduplicated leftover sequences (% of total) | 79,16 | 45,89 | 54,89 | 54,25 | 49,39 | 59,75 |
| ***bismark methylation extractor*** | **Total number of methylation call strings (=R1+ R2 reads)** | **34918** | **57842** | **57196** | **67688** | **79770** | **103108** |
| | **Total number of C's analysed:** | **480 679** | **1010776** | **976435** | **1176089** | **1458199** | **2041451** |
| | **Total methylated C's in CpG context:** | **1307** | **1873** | **1916** | **2102** | **2369** | **2751** |
| | Total methylated C's in CHG context: | 1671 | 2560 | 2625 | 2804 | 3147 | 3872 |
| | Total methylated C's in CHH context: | 7617 | 8635 | 9137 | 9188 | 9601 | 9803 |
| | Total C to T conversions in CpG context: | 136744 | 297737 | 285086 | 346866 | 429779 | 606148 |
| | Total C to T conversions in CHG context: | 146887 | 315798 | 302944 | 368922 | 458882 | 646756 |
| | Total C to T conversions in CHH context: | 186453 | 384173 | 374727 | 446207 | 554421 | 772121 |
| | **C methylated in CpG context:** | **0,95** | **0,63** | **0,67** | **0,60** | **0,55** | **0,45** |
| | C methylated in CHG context: | 1,12 | 0,80 | 0,86 | 0,75 | 0,68 | 0,60 |
| | C methylated in CHH context: | 3,92 | 2,20 | 2,38 | 2,02 | 1,70 | 1,25 |
| | **Total methylation (i.e. total C non converted by BS) %** | 2,20 | 1,29 | 1,40 | 1,20 | 1,04 | 0,80 |
| | **BS conversion efficiency (%)** | **97,80** | **98,71** | **98,60** | **98,80** | **98,96** | **99,20** |

DNA of Lambda phage virus were added to each sample during the sequencing. These sequences are used as DNA-spike in, which allows estimating bisulfite sodium conversion (BS), since these sequences are completely unmethylated before the conversion. Reads that mapped to lambda phage reference genome were deduplicated before the estimation of bisulfite sodium conversion (BS). The high percent observed for the bisulfite (BS) conversion efficiency for all samples allows us to analyze the data.
