## Supplementary material for "Immediate perturbations of DNA methylation and transcriptome upon acute prenatal alcohol exposure in the mouse developing brain cortex": Supp Table S2 Random datasets

**Suppl. Table 2: Comparison results between real and random datasets, and parameters finally chosen for DMRs detection using *get_close_loci()* function.**

**with first random dataset**

| pvalue thresh. | Min nb of C | Nb of selected C: real data | Nb of selected C: random data | Nb ofdetected DMR: real data | Nb ofdetected DMR: adjusted real data | Nb ofdetected DMR: random data | adjust. real ÷ random |
|---|---|---|---|---|---|---|---|
| 0.008 | 3 | 12 419 | 7 225 | 114 | 28 | 9 | 3.1 |
| | 4 | | | 16 | 2 | 0 | - |
| 0.01 | 3 | 15 089 | 8 942 | 160 | 40 | 17 | 2.4 |
| | 4 | | | 21 | 5 | 0 | - |
| 0.03 | 3 | 38 843 | 26 785 | 1080 | 453 | 263 | 1.7 |
| | 4 | | | 240 | 85 | 20 | 4.3 |
| | 5 | | | 64 | 22 | 3 | 7.3 |
| 0.05 | 3 | 60 936 | 44 899 | 2603 | 1380 | 924 | 1.5 |
| | 4 | | | 672 | 229 | 127 | 2.4 |
| | 5 | | | 225 | 94 | 18 | 5.2 |
| 0.08 | 3 | 92 657 | 73 225 | 5552 | 3506 | 2563 | 1.3 |
| | 4 | | | 1684 | 935 | 461 | 2.0 |
| | 5 | | | 581 | 287 | 81 | 3.5 |
| 0.1 | 3 | 118 453 | 95 687 | 8331 | 5659 | 4358 | 1.2 |
| | 4 | | | 2733 | 1679 | 912 | 1.8 |
| | 5 | | | 1016 | 560 | 190 | 2.9 |

**with second random dataset**

| pvalue thresh. | Min nb of C | Nb of selected C: real data | Nb of selected C: random data | Nb of detected DMR: real data | Nb of detected DMR: adjusted real data | Nb of detected DMR: random data | adjust. real ÷ random |
|---|---|---|---|---|---|---|---|
| 0.05 | 3 | 60 936 | 44 584 | 2603 | 1280 | 814 | 1.6 |
| | 4 | | | 672 | 281 | 98 | 2.9 |
| | 5 | | | 225 | 87 | 15 | 5.8 |
| 0.06 | 5 | 71 157 | 53 441 | 329 | 138 | 28 | 4.9 |
| 0.07 | 5 | 81 494 | 62 842 | 432 | 184 | 42 | 4.4 |
| 0.08 | 3 | 92 657 | 73 170 | 5552 | 3467 | 2368 | 1.5 |
| | 4 | | | 1684 | 932 | 429 | 2.2 |
| | 5 | | | 581 | 263 | 72 | 3.7 |

**Arguments values finally used for DMR detection in the real dataset**

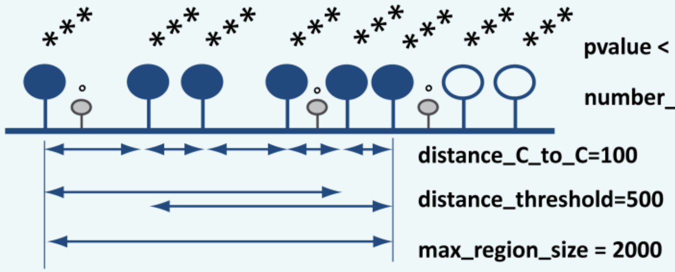


Two random datasets (for details on random dataset obtention, see **Supplementary Figure 3** and **Materials & Methods**) were compared to the actual data, by testing different combinations of DMR detection parameters (p-value threshold, minimum number of CpGs in the region), and by fixing some parameters (*distance_C_to_C* = 100, *distance_threshold*=500, *max_region_size* = 2000). Depending on the p-value threshold chosen, a greater or lesser number of CpGs is selected. In each case, more CpGs are selected in the real dataset than in the random one, for a given p-value threshold. To compare the number of DMRs obtained in the real dataset and in the random one, the same number of CpGs must be considered at the outset. Thus, by random selection of relevant CpGs into the real dataset, the number of CpGs in the real dataset was adjusted to the number of CpGs found in the random dataset. ***Abbreviations***: ***Min nb of C***: minimum number of CpGs required to obtain a DMR; ***Nb of selected C***: number of CpG taken into account, depending on the chosen p-value threshold; ***adjust. real data ÷ random***: ratio of the number of DMRs detected in the adjusted real dataset and the number of

DMRs detected in the random dataset. On the scheme, filled circles represent hypermethylated cytosines observed between two datasets (*e.g.* control group *versus* test group), whereas empty circles represent hypomethylated cytosines. *: non-significant p-value (pval ≥ 0.07); *** significant  p-value (pval < 0.07).
