## Supplementary material for "Immediate perturbations of DNA methylation and transcriptome upon acute prenatal alcohol exposure in the mouse developing brain cortex": Supp Table S3 Key Numbers Methylome Analysis

**Supplementary Table 3: Summary of key number of major steps of the bioinformatic analysis of the methylome capture.**

| | | 1M-R1 | 1M-R2 | 2M-R1 | 2M-R2 | 3M-R1 | 3M-R2 | 4M-R1 | 4M-R2 | 5M-R1 | 5M-R2 | 6M-R1 | 6M-R2 |
|---|---|---|---|---|---|---|---|---|---|---|---|---|---|
| **Raw data** | Initial number of reads | 67 884 931 | 67 884 931 | 66 493 707 | 66 493 707 | **66 654 738** | **66 654 738** | **68 515 383** | **68 515 383** | **75 016 568** | **75 016 568** | **73 103 368** | **73 103 368** |
| **Trimmomatic** | Nb of reads after trimming | 45 188 206 | 45 188 206 | 48 803 924 | 48 803 924 | 52 093 098 | 52 093 098 | 48 406 088 | 48 406 088 | 54 295 610 | 54 295 610 | 54 238 624 | 54 238 624 |
| | % of remaining reads | 66,57 | 66,57 | 73,40 | 73,40 | 78,15 | 78,15 | 70,65 | 70,65 | 72,38 | 72,38 | 74,19 | 74,19 |
| **Cutadapt** | % of removed bases | 19,9 | 19,2 | 19,9 | 19,3 | 19,9 | 19,2 | 19,9 | 19,2 | 19,9 | 19,2 | 19,9 | 19,2 |
| **Bismark** *-paired end mapping - mm9 - non directional library* | Nb of mapped reads | 27 132 662 | | 33 834 717 | | 32 974 018 | | 33 399 024 | | 37 335 974 | | 37 685 044 | |
| | Mapping efficiency (mapped reads ÷ trimmed reads - %) | **60,0** | | **69,3** | | **63,3** | | **69,0** | | **68,8** | | **69,5** | |
| | Total number of C's analysed | **1 393 221 218** | | **1 723 029 465** | | **1 688 063 509** | | **1 671 822 446** | | **1 915 763 720** | | **1 954 602 883** | |
| | Total 5mC in CpG context | 38 343 757 | | 50 120 323 | | 46 162 661 | | 46 221 886 | | 51 128 425 | | 52 101 907 | |
| | % 5mC in CpG context () | **42,1** | | **41,5** | | **42,8** | | **43,7** | | **41,1** | | **40,9** | |
| **deduplication** | Total number of alignments analysed | 27 132 662 | | 33 834 713 | | 32 974 016 | | **33 399 023** | | **37 335 971** | | **37 685 043** | |
| | Total count of deduplicated leftover sequences (% of total) | 85,87 | | 78,88 | | 80,66 | | 79,61 | | 78,46 | | 80,85 | |
| **bismark_ methylation_ extractor** | Total number of methylation call strings (=R1 + R2 reads) | 46 595 798 | | 53 377 408 | | 53 191 184 | | 53 179 200 | | 58 586 780 | | 60 938 390 | |
| | Total number of C's analysed: | 786 358 255 | | 905 431 146 | | 945 954 930 | | 886 146 203 | | 1 021 808 647 | | 1 076 902 457 | |
| | Total 5mC in CpG context: | 21 654 699 | | 26 665 187 | | 26 052 368 | | 24 693 496 | | 27 465 624 | | 28 834 224 | |
| | Total 5mC in CHG context: | 609 486 | | 754 954 | | 744 991 | | 712 498 | | 803 420 | | 829 707 | |
| | Total 5mC in CHH context: | 2 007 478 | | 2 260 093 | | 2 376 417 | | 2 251 004 | | 2 554 514 | | 2 630 633 | |
| | Tot. C→T conversion (CpG) | 29 956 981 | | 36 584 824 | | 34 103 744 | | 31 538 942 | | 38 906 686 | | 41 911 091 | |
| | Tot. C→T conversion (CHG) | 199 634 753 | | 235 186 432 | | 237 395 182 | | 223 667 975 | | 256 270 802 | | 271 045 886 | |
| | Tot. C→T conversion (CHH) | 532 494 858 | | 603 979 656 | | 645 282 228 | | 603 282 288 | | 695 807 601 | | 731 650 916 | |
| | 5mC in CpG context (%) | **42,0** | | **42,2** | | **43,3** | | **43,9** | | **41,4** | | **40,8** | |
| | 5mC in CHG context (%) | **0,3** | | **0,3** | | **0,3** | | **0,3** | | **0,3** | | **0,3** | |
| | 5mC in CHH context (%) | **0,4** | | **0,4** | | **0,4** | | **0,4** | | **0,4** | | **0,4** | |
| | Total C in CpG (5C + 5mC) | 51611680 | | 63250011 | | 60156112 | | 56232438 | | 66372310 | | 70745315 | |
| **bedtools intersect** | % deduplicated reads on capture regions | 57,36 | | 55,38 | | 54,80 | | 59,44 | | 59,34 | | 61,20 | |
| **Statistical analysis with methylkit + get-close_loci()** | Nb CpG sites covered in all samples (unfiltered) | 3 016 144 | | | | | | | | | | | |
| | Filtered positions (min. cov. = 10, high. Perc = 99.9 ; coverage normalisation | 1 676 294 | | 1 920 575 | | 1 876 952 | | 1 766 642 | | 2 080 535 | | 2 149 843 | |
| | Filtered CpG positions covered by ALL samples | 1 259 111 | | | | | | | | | | | |
| | DMC (Meth.diff 5%, qval<0,05) | 4 | | | | | | | | | | | |
| | DMR all | 432 | | | | | | | | | | | |

For more details about the bioinformatic workflow, please see Materiel & Methods and command line in Supp. data notebook n°1.
