## Supplementary material for "Immediate perturbations of DNA methylation and transcriptome upon acute prenatal alcohol exposure in the mouse developing brain cortex": Supp Table S4 Datamining ATAC-seq data

**Supplementary Table 4. Summary of key numbers at major steps of the ATAC-seq data bioinformatics analysis.**

| | | **E13_A_R1** | **E13_A_R2** | **E13_B_R1** | **E13_B_R2** | **E14_A_R1** | **E14_A_R2** | **E14_B_R1** | **E14_B_R2** | **E15_A_R1** | **E15_A_R2** | **E15_B_R1** | **E15_B_R2** |
|---|---|---|---|---|---|---|---|---|---|---|---|---|---|
| | | ENCFF401VUV | ENCFF898NRO | ENCFF721LGJ | ENCFF777UKE | ENCFF633MTW | ENCFF666DRJ | ENCFF048MTG | ENCFF890LGM | ENCFF906VXU | ENCFF500SXI | ENCFF248PXW | ENCFF825UHO |
| raw data | Initial number of reads | 49 691 764 | 49 691 764 | 51 407 232 | 51 407 232 | 48 416 233 | 48 416 233 | 35 924 683 | 35 924 683 | 43 503 968 | 43 503 968 | 53 935 170 | 53 935 170 |
| trimming | Nb of reads after trimming | 49687965 | 49687965 | 51403240 | 51403240 | 48412609 | 48412609 | 35922216 | 35922216 | 43500617 | 43500617 | 53929938 | 53929938 |
| | % of remaining reads | 99,99% | 99,99% | 99,99% | 99,99% | 99,99% | 99,99% | 99,99% | 99,99% | 99,99% | 99,99% | 99,99% | 99,99% |
| | % of removed reads | 0,01% | 0,01% | 0,01% | 0,01% | 0,01% | 0,01% | 0,01% | 0,01% | 0,01% | 0,01% | 0,01% | 0,01% |
| mapping | total number of paired reads mapped | 36 397 426 | | 38 620 494 | | 36 652 887 | | 26 378 618 | | 31 398 457 | | 38 691 368 | |
| | Mapping efficiency (%) | 73,25% | | 75,13% | | 75,71% | | 73,43% | | 72,18% | | 71,74% | |
| | Total nb of mapped reads analyzed | 95 412 250 | | 99 287 128 | | 93 808 282 | | 69 372 774 | | 83 421 228 | | 103 835 880 | |
| rdeduplication | Total nb of mapped reads analyzed after rmdup | 83 165 510 | | 88 247 440 | | 82 678 024 | | 61 178 310 | | 71 934 002 | | 85 449 160 | |
| | % of leftover mapped reads after rmdup | 87,16% | | 88,88% | | 88,14% | | 88,19% | | 86,23% | | 82,29% | |
| chrM removal | nb of reads after ChrM removal | 80 625 623 | | 86 068 230 | | 80 727 371 | | 60 068 252 | | 69 554 553 | | 82 526 570 | |
| | % of leftover mapped reads after ChRM removal | 96,95% | | 97,53% | | 97,64% | | 98,19% | | 96,69% | | 96,58% | |
| Peak calling | total tags in treatment | 77 237 227 | | | | 63 606 160 | | | | 68 870 966 | | | |
| | tags after filtering (= rm dup) in treatment | 76 334 462 | | | | 62 971 395 | | | | 68 107 975 | | | |
| | Nb of peaks detected in narrowPeaks | 70 364 | | | | 69 303 | | | | 69 128 | | | |
| bedtools multiinter + bedtools merge | Nb of regions in at least one sample | 115 752 | | | | | | | | | | | |
| count table | SAM alignment record pairs processed | 41 645 400 | | 44 222 921 | | 30 948 315 | | 41 454 717 | | 42 466 253 | | 35 842 612 | |
| statistical analysis | DOCR selection criteria | BH p-value adjustment and the level of controlled false positive rate is set to 0.05 | | | | | | | | | | | |
| | samples | **E13.14 up** | | **E14.15 up** | | **E15.16 up** | | **E16.P0 up** | | **E14.16 up** | | **E13.14 down** | |
| | nb of DOCR (mm9) | 4 | | 1442 | | 280 | | 13823 | | 6522 | | 1 | |
| | nb of DOCR (mm10) | 4 | | 1441 | | 280 | | 13817 | | 6522 | | 1 | |

**Suppl. Table 4 (continued)**

| | | E16_A_R1 | E16_A_R2 | E16_B_R1 | E16_B_R2 | P0_A_R1 | P0_A_R2 | P0_B_R1 | P0_B_R2 |
|---|---|---|---|---|---|---|---|---|---|
| | | ENCFF058IAE | ENCFF765HUX | ENCFF776GDQ | ENCFF588XZG | ENCFF197GTC | ENCFF209GGJ | ENCFF296GZG | ENCFF664RZO |
| raw data | Initial number of reads | 42 278 369 | 42 278 369 | 43 919 645 | 43 919 645 | 41 466 599 | 41 466 599 | 47548519 | 47548519 |
| trimming | Nb of reads after trimming | 42274976 | 42274976 | 43915585 | 43915585 | 41463017 | 41463017 | 47544874 | 47544874 |
| | % of remaining reads | 99,99% | 99,99% | 99,99% | 99,99% | 99,99% | 99,99% | 99,99% | 99,99% |
| | % of removed reads | 0,01% | 0,01% | 0,01% | 0,01% | 0,01% | 0,01% | 0,01% | 0,01% |
| mapping | total number of paired reads mapped | 31 885 090 | | 31 813 067 | | 30 876 756 | | 35 937 329 | |
| | Mapping efficiency (%) | 75,42% | | 72,44% | | 74,47% | | 75,59% | |
| | Total nb of mapped reads analyzed | 81 628 676 | | 84 688 118 | | 79 746 404 | | 91 825 542 | |
| rdeduplication | Total nb of mapped reads analyzed after rmdup | 67 723 330 | | 70 768 240 | | 65 301 028 | | 80 135 886 | |
| | % of leftover mapped reads after rmdup | 82,97% | | 83,56% | | 81,89% | | 87,27% | |
| chrM removal | nb of reads after ChrM removal | 65 147 743 | | 67 855 478 | | 63 265 360 | | 77 800 611 | |
| | % of leftover mapped reads after ChRM removal | 96,20% | | 95,88% | | 96,88% | | 97,09% | |
| Peak calling | total tags in treatment | 60 540 712 | | | | 64 754 019 | | | |
| | tags after filtering (= rm dup) in treatment | 59 578 110 | | | | 63 577 084 | | | |
| | Nb of peaks detected in narrowPeaks | 78 996 | | | | 81 249 | | | |
| bedtools multiinter + bedtools merge | Nb of regions in at least one sample | 115752 | | | | | | | |
| count table | SAM alignment record pairs processed | 33 458 239 | | 34 861 682 | | 33 198 116 | | 40 507 551 | |
| statistical analysis | DOCR selection criteria | BH p-value adjustment and the level of controlled false positive rate is set to 0.05 | | | | | | | |
| | samples | **E14.15 down** | | **E15.16 down** | | **E16.P0 down** | | **E14.16 down** | |
| | nb of DOCR (mm9) | 1614 | | 420 | | 14413 | | 4545 | |
| | nb of DOCR (mm10) | 1614 | | 420 | | 14413 | | 4545 | |
