## Supplementary material for "Immediate perturbations of DNA methylation and transcriptome upon acute prenatal alcohol exposure in the mouse developing brain cortex": Supp Table S5 Datamining ENCODE RNAseq

**Supplementary Table 5. Summary of key numbers at major steps of the RNA-seq data bioinformatics analysis.**

| | | P0_rep1 | P0_rep2 | E13.5_rep1 | E13.5_rep2 | E14.5_rep1 | E14.5_rep2 | E15.5_rep1 | E15.5_rep2 | E16.5_rep1 | E16.5_rep2 |
|---|---|---|---|---|---|---|---|---|---|---|---|
| **Raw data** | Number (nb) of reads | 53 468 133 | 52 580 657 | 55 578 426 | 58 648 041 | 52 984 225 | 44 427 005 | 57 991 263 | 51 931 736 | 46 904 248 | 38 483 829 |
| **1er trimming** | nb of remaining reads | 53 161 252 | 52 303 312 | 52 466 124 | 55 863 362 | 51 021 076 | 42 875 143 | 55 705 362 | 50 251 431 | 44 750 545 | 35 969 262 |
| **2nd trimming** | nb pf remaining reads | | 51 880 675 | | | 50 697 328 | | | | | |
| **Mapping (mm10)** | **UNIQUE READS:** | | | | | | | | | | |
| | Uniquely mapped reads number | 45 476 798 | 43 211 812 | 44 614 345 | 46 673 269 | 38 703 337 | 36 051 036 | 47 872 767 | 43 277 786 | 39 003 795 | 31 116 470 |
| | Uniquely mapped reads % | 85.55 | 83.29 | 85.03 | 83.55 | 81.45 | 84.08 | 85.94 | 86.12 | 87.16 | 86.51 |
| | mismatch rate per base (%) | 0.42 | 0.40 | 0.24 | 0.22 | 0.25 | 0.23 | 0.2 | 0.2 | 0.21 | 0.26 |
| | deletion rate per base (%) | 0.01 | 0.01 | 0.01 | 0.01 | 0.02 | 0.01 | 0.01 | 0.01 | 0.01 | 0.01 |
| | insertion rate per base (%) | 0.01 | 0.01 | 0.01 | 0.01 | 0.01 | 0.01 | 0.01 | 0.01 | 0.01 | 0.01 |
| | **MULTI-MAPPING READS:** | | | | | | | | | | |
| | nb of reads mapped to multiple loci | 0 | 0 | 0 | 0 | 0 | 0 | 0 | 0 | 0 | 0 |
| | % of reads mapped to too many loci | 11.15 | 13.48 | 10.9 | 11.58 | 13.92 | 11.54 | 9.93 | 9.99 | 9.14 | 10.16 |
| | **UNMAPPED READS:** | | | | | | | | | | |
| | % of reads unmapped: too many mismatches | 0.00 | 0 | 0 | 0 | 0 | 0 | 0 | 0 | 0 | 0 |
| | % of reads unmapped: too short | 2.81 | 2.83 | 2.62 | 3.15 | 3.52 | 3.29 | 2.84 | 2.58 | 2.54 | 2.1 |
| | % of reads unmapped: other | 0.49 | 0.4 | 1.44 | 1.73 | 1.1 | 1.08 | 1.29 | 1.3 | 1.16 | 1.23 |
| **DEG detection (edgeR)** | DEG selection criteria | BH p-value adjustment and the level of controlled false positive rate is set to 0.05 | | | | | | | | | |
| | samples | **E13.14 up** | **E14.15 up** | **E15.16 up** | **E16.P0 up** | **E14.16 up** | **E13.14 down** | **E14.15 down** | **E15.16 down** | **E16.P0 down** | **E14.16 down** |
| | nb de DEG | **3744** | **3426** | **196** | **4049** | **2791** | **3679** | **2512** | **327** | **4299** | **2067** |
