## Supplementary material for "Immediate perturbations of DNA methylation and transcriptome upon acute prenatal alcohol exposure in the mouse developing brain cortex": Supp Data Notebook1 Methylome workflow: Supp. data Notebook 1 - Methylome workflow - Duchateau et al.html

Annexe 3.1 Methylome workflow


### Supplementary data Notebook 1 - Methylome capture workflow¶

##### *Agathe Duchateau*¶

#### Raw data¶

Raw data were obtained from two distinct runs of sequencing, performed on the same samples (same libraries):

- **1M, 2M and 3M** are **ethanol**-treated samples
- **4M, 5M and 6M** are **PBS**-treated samples

After the first sequencing, 8 fastq files per sample are obtained (four files corresponding to R1 reads and four files for R2 reads).
After the second sequencing, 2 fastq files per sample are obtained (one file corresponding to R1 reads and one file for R2 reads).

#### 1.Tools used for the analysis¶

Following tools and R packages were used to analyse these data. Version of each tool is indicated.

In [ ]:

```
# ----------- bash tools -------------------
# FASTQC (version 0.11.3)
# Trimmomatic (version 0.33)
# cutadapt (version 1.9.1 with Python 2.7.11)
# bismark_genome_preparation (version 0.14.4)
# bismark (version 0.14.5)
# deduplicate_bismark (version 0.14.5)
# bismark_methylation_extractor (version 0.14.4)
# bedtools (merge ; intersect ; getfasta ; multiinter = multiIntersectBed) (version 2.25.0, on MAC)
# samtools (sort ; index ; view) (version 0.1.19-44428cd, on MAC)
# seqkit locate (version 0.10.2, in a conda environment v4.7.11)
# findMotifsGenome (HOMER suite v4.9.1, in a conda environment v4.7.11)

# ----------- R (version 3.2.0) packages ------------
library(methylKit) #v0.9.4
library(devEMF) #v3.6
library(data.table) #v1.10.4

# ----------- R (version 3.5.2) packages ------------
library(ggplot2) # v3.2.1
library(reshape2) # v1.4.3
libary(BiomaRt) # v2.38.0
library(dplyr) # v0.8.3
library(eulerr) # v5.1.0
library(BSgenome) #v 1.48.0
library(BSgenome.Mmusculus.UCSC.mm9) #1.4.0
library(org.Mm.eg.db) #3.7.0

# ------------ other tools ------------
# IGV (version 2.8.0)
# UCSC Table browser
# geneimprint database
# homer_annotatePeaks (Galaxeast platform - Galaxy Tool version 0.0.5 - dependency: homer v4.7
# WEB-based GEne SeT AnaLysis Toolkit
```

#### 2.Quality control of raw data and files merging¶

**FASTQC** was used to verify the quality of the sequencing raw data.

###### Conclusions (first sequencing)¶

- There are around 4 000 000 - 4 500 000 reads per file.
- "Per Base Sequence Quality" is globally better for R1 reads than for R2 ones.
- There are not remaining adapter sequences.
- There is a warning for "Sequence Length Distribution". It seems that there are many small sequences but it doesn't compromise the analysis.
- There is an imbalance in term of "CG content", as expected since bisulfite treatment was performed on the samples.
- Sometimes, an over representation of "AAAAAA", "GGGGGGG" or "TTTTTGGGG…GGGGG" patterns can be observed, notably for R2 reads.
- Sometimes (R1 and/or R2 reads), "per tile sequence quality" is not perfect, reflecting some overloading zone in the flowcell but the problem is limited to very small areas.

###### Conclusions (second sequencing)¶

- There are more reads in this sequencing compared to the first one: around 52 millions reads per file here.
- "Per Base Sequence Quality" is globally better for R1 reads than for R2 ones.
- There is a warning for "Per Base Sequence Quality" category for all samples.
- Adapter sequences (Illumina Universal Adapters) are still presented into the reads.
- There is an imbalance in term of "CG content", as expected since bisulfite treatment was performed on the samples.
- For almost all samples, an over representation of "AAAAAA", "GGGGGGG" or "TTTTTTT" patterns can be observed.
- Read extremities are not as expected.

All R1 reads, on one hand, and all R2reads, on another hand, were kept and merged together using **cat** function. Thus, for a given sample, we obtained one file with all R1 reads, and another file with R2 reads.

In [ ]:

```
%%bash

cd /Users/UMR7216/Documents/Mezger/methylome/donnees-brutes-cat

cat 1M_S1_L001_R1_001.fastq.gz 1M_S1_L002_R1_001.fastq.gz 1M_S1_L003_R1_001.fastq.gz 1M_S1_L004_R1_001.fastq.gz 1M_R1.fastq.gz > fus_1M_R1.fastq.gz
cat 1M_S1_L001_R2_001.fastq.gz 1M_S1_L002_R2_001.fastq.gz 1M_S1_L003_R2_001.fastq.gz 1M_S1_L004_R2_001.fastq.gz 1M_R2.fastq.gz > fus_1M_R2.fastq.gz

cat 2M_S2_L001_R1_001.fastq.gz 2M_S2_L002_R1_001.fastq.gz 2M_S2_L003_R1_001.fastq.gz 2M_S2_L004_R1_001.fastq.gz 2M_R1.fastq.gz > fus_2M_R1.fastq.gz
cat 2M_S2_L001_R2_001.fastq.gz 2M_S2_L002_R2_001.fastq.gz 2M_S2_L003_R2_001.fastq.gz 2M_S2_L004_R2_001.fastq.gz 2M_R2.fastq.gz > fus_2M_R2.fastq.gz

cat 3M_S3_L001_R1_001.fastq.gz 3M_S3_L002_R1_001.fastq.gz 3M_S3_L003_R1_001.fastq.gz 3M_S3_L004_R1_001.fastq.gz 3M_R1.fastq.gz > fus_3M_R1.fastq.gz
cat 3M_S3_L001_R2_001.fastq.gz 3M_S3_L002_R2_001.fastq.gz 3M_S3_L003_R2_001.fastq.gz 3M_S3_L004_R2_001.fastq.gz 3M_R2.fastq.gz > fus_3M_R2.fastq.gz

cat 4M_S4_L001_R1_001.fastq.gz 4M_S4_L002_R1_001.fastq.gz 4M_S4_L003_R1_001.fastq.gz 4M_S4_L004_R1_001.fastq.gz 4M_R1.fastq.gz > fus_4M_R1.fastq.gz
cat 4M_S4_L001_R2_001.fastq.gz 4M_S4_L002_R2_001.fastq.gz 4M_S4_L003_R2_001.fastq.gz 4M_S4_L004_R2_001.fastq.gz 4M_R2.fastq.gz > fus_4M_R2.fastq.gz

cat 5M_S5_L001_R1_001.fastq.gz 5M_S5_L002_R1_001.fastq.gz 5M_S5_L003_R1_001.fastq.gz 5M_S5_L004_R1_001.fastq.gz 5M_R1.fastq.gz > fus_5M_R1.fastq.gz
cat 5M_S5_L001_R2_001.fastq.gz 5M_S5_L002_R2_001.fastq.gz 5M_S5_L003_R2_001.fastq.gz 5M_S5_L004_R2_001.fastq.gz 5M_R2.fastq.gz > fus_5M_R2.fastq.gz

cat 6M_S6_L001_R1_001.fastq.gz 6M_S6_L002_R1_001.fastq.gz 6M_S6_L003_R1_001.fastq.gz 6M_S6_L004_R1_001.fastq.gz 6M_R1.fastq.gz > fus_6M_R1.fastq.gz
cat 6M_S6_L001_R2_001.fastq.gz 6M_S6_L002_R2_001.fastq.gz 6M_S6_L003_R2_001.fastq.gz 6M_S6_L004_R2_001.fastq.gz 6M_R2.fastq.gz > fus_6M_R2.fastq.gz
```

#### 3.Quality control of merged raw data¶

**FASTQC** was used to verify the quality of reads after the merging.

###### Conclusions¶

- "Per Base Sequence Quality" is globally better for R1 reads than for R2 ones.
- There is a warning for "Per Base Sequence Quality" category for all samples.
- Adapter sequences (Illumina Universal Adapters) are still presented into the reads.
- There is a warning for "Sequence Length Distribution". It seems that there are small sequences (35 bases) but it doesn't compromise the analysis.
- There is still an imbalance in term of "CG content", as expected since bisulfite treatment was performed on the samples.
- For almost all samples, an over representation of "AAAAAA", "GGGGGGG" or "TTTTTTT" patterns can be observed.
- Read extremities are not as expected.

#### 4.Trimming¶

To decrease methylation call errors from poor quality data, a trimming was performed on low quality base calls before read mapping. The remaining sequencing adapters and reads having bad sequencing quality were trimmed using **Trimmomatic** with default parameters.

In [ ]:

```
%%bash

mkdir -p /Users/UMR7216/Documents/Mezger/methylome/cat-trim

cd /Users/UMR7216/Documents/Mezger/methylome/donnees-brutes-cat

!/bin/bash
for r1 in *R1.fastq.gz
    do
    r2=$r1   
    r2="${r1/R1/R2}"  
    r1p="t_${r1/.fastq.gz/.paired.fastq.gz}"
    r1u="t_${r1/.fastq.gz/.unpaired.fastq.gz}"
    r2p="t_${r2/.fastq.gz/.paired.fastq.gz}"
    r2u="t_${r2/.fastq.gz/.unpaired.fastq.gz}"
    data='/Users/UMR7216/Documents/Mezger/methylome/cat-trim'
    java -jar /Users/UMR7216/Desktop/Bioinfo_TOOLS/Trimmomatic-0.33/trimmomatic-0.33.jar PE -threads 12 -phred33 $r1 $r2 $data/$r1p $data/$r1u $data/$r2p $data/$r2u ILLUMINACLIP:/Users/UMR7216/Desktop/Bioinfo_TOOLS/Trimmomatic-0.33/adapters/TruSeq3-PE-2.fa:2:30:10:1 LEADING:3 TRAILING:3 SLIDINGWINDOW:4:15 
done
```

#### 5.Quality control of filtered data (after Trimmomatic) and second trimming¶

**FASTQC** was used to verify the quality of filtered reads.

###### Conclusions¶

- "Per Base Sequence Quality" of R1 and R2 reads is better than those of unfiltered reads.
- Sequencing adapters (Illumina Universal Adapters) are well removed.
- There is a warning for "Sequence Length Distribution", as before the trimming.
- There is still an imbalance in term of "CG content", as expected since bisulfite treatment was performed on the samples.
- For almost all samples, an over representation of "GGGGGGG" pattern is observed among R1 reads and "A" or "AA" or "AAA" or "C" or "T" or "AAA…AAA", or "GGG…GGG" patterns are observed among R2 reads.
- Read extremities are not as expected.

According to these observations, trimming has been done properly but it was not sufficient for read extremities. Thus, files were unzipped and **cutadapt** was used to remove 10 bases at each read extremity (both at 5' and 3' extremities).

In [ ]:

```
%%bash 

# To unzip files
cd /Users/UMR7216/Documents/Mezger/methylome/cat-trim

#!/bin/bash
for file in t_fus_*M_R*.paired.fastq.gz
do
    gzcat $file > /Volumes/Disque_2_To/methylome/fusion/cat/${file%.fastq.gz}.fastq
done

# To remove 10 bases at each read extremity
cd /Volumes/Disque_2_To/methylome/fusion/cat/

#!/bin/bash
for ftrim in t_fus_*M_R*.paired.fastq
do 
    cutadapt -u -10 -u +10 -o /Users/UMR7216/Documents/Mezger/methylome/cat-trim/c10${ftrim%paired.fastq}fastq $ftrim 
done
```

#### 6.Quality control of filtered data (after Cutadapt)¶

**FASTQC** was used to verify the quality of filtered reads.

###### Conclusions¶

- "Per Base Sequence Quality" of R1 and R2 reads is still satisfying.
- Sequencing adapters (Illumina Universal Adapters) are well removed.
- There is still an imbalance in term of "CG content", as expected since bisulfite treatment was performed on the samples.
- For almost all samples, an over representation of "GGGGGGG" pattern is observed among R1 reads and "A" or "AA" or "AAA" or "GGG…GGG" patterns are observed among R2 reads. There are also "empty" reads.
- Read extremities are as expected.

According to these observations, cutadapt worked properly.

#### 7.Verification of bisulfite conversion¶

Depending on the experimental conditions, sodium bisulfite conversion may not be complete (Clark *et al.*, 2006), leaving certain unmethylated cytosines unchanged whether to modify them into thymines. To ensure that sodium bisulfite treatment correctly converted unmethylated cytosines, we evaluated the conversion efficiency by looking at conversion rate of a DNA spike-in (i.e. unmethylated known sequences of lambda phage), which was added in each sample preparation. This efficiency is obtained by:

- mapping filtered reads on lambda phage reference genome (Genbank accession number **NC\_001416**, FASTA file).
- calculating the ratio of converted reads (containing thymines) to the total number of reads covering these cytosines. Informations are obtained in mapping report of each sample.

##### 7.1.Lambda phage genome preparation¶

To take into account methylation profile (sodium bisulfite conversion) of filtered reads, **bismark** mapper was used since it is adapted for alignment of bisulfite converted reads. To map filtered reads on lambda phage reference genome, this genome must be prepared, using **bismark\_genome\_preparation**.

In [ ]:

```
%%bash 

cd /Users/UMR7216/Desktop/Bioinfo_TOOLS/bismark_v0.14.5

./bismark_genome_preparation --path_to_bowtie 
/Users/UMR7216/Desktop/Bioinfo_TOOLS/bowtie2-2.1.0 
--bowtie2 --verbose 
/Users/UMR7216/Documents/Mezger/capture-methylome/Diderot-CaptureEpi-2015-479-27264242/genome-lambda_phage
```

##### 7.2. Mapping on lambda phage genome¶

Then, filtered reads are mapped using **bismark**, with paired-end parameters. Libraries are considered as non-directional for the mapping.

In [ ]:

```
%%bash

mkdir -p  /Volumes/DD-Agathe/methylome/MALE-FUSION/align-lambda/1M
mkdir -p  /Volumes/DD-Agathe/methylome/MALE-FUSION/align-lambda/2M
mkdir -p  /Volumes/DD-Agathe/methylome/MALE-FUSION/align-lambda/3M
mkdir -p  /Volumes/DD-Agathe/methylome/MALE-FUSION/align-lambda/4M
mkdir -p  /Volumes/DD-Agathe/methylome/MALE-FUSION/align-lambda/5M
mkdir -p  /Volumes/DD-Agathe/methylome/MALE-FUSION/align-lambda/6M

cd /Users/UMR7216/Documents/Mezger/methylome/cat-trim

#!/bin/bash
for file1 in c10t_fus*M_R1.fastq
    do
    file2=$file1   
    file2="${file1/R1/R2}"
    director="/Volumes/DD-Agathe/methylome/MALE-FUSION/align-lambda/${file1:9:2}"
    bismark -N 1 /Users/UMR7216/Documents/Mezger/methylome/genome-lambda_phage/ 
    --path_to_bowtie /Users/UMR7216/Desktop/Bioinfo_TOOLS/bowtie2-2.1.0 
    --non_directional --un 
    -o $director -1 /Users/UMR7216/Documents/Mezger/methylome/cat-trim/$file1 
    -2 /Users/UMR7216/Documents/Mezger/methylome/cat-trim/$file2
done
```

##### 7.3. Deduplication and extraction of methylation state to estimate bisulfite treatment efficiency¶

Duplicate reads are generated from amplification of a unique original DNA fragment and map exactly to the same genomic location (same chromosome, starting and ending positions). Since some biases can be introduced by possible preferential PCR amplification, the estimation of methylation level is biased toward the over-amplified DNA molecules, having different methylation state from other DNA molecules if we keep duplicate reads.

To obtain a more accurate estimation of bisulfite treatment efficiency, reads that mapped to lambda phage genome were deduplicated, using **deduplicate\_bismark**, a tool from **bismark** suite, that is adapted for the deduplication of bisulfite converted reads which require special attention due to the loss of complementarity between strands after PCR amplification (this tool is able to identify duplicates, even if original strands are no more complementary upon sodium bisulfite conversion and PCR amplification).

After deduplication, **bismark\_methylation\_extractor**, another tool from **bismark** suite, was used to extract methylation percent (it gives the number of methylated and unmethylated cytosines that covered a given cytosine site) and thereby estimate sodium bisulfite conversion efficiency.

In [ ]:

```
%%bash

# Deduplication
#!/bin/bash
for fbam in /Volumes/DDMezger/methylome/fusion/align-lambda/*M/c10t_fus_*M_R1.fastq_bismark_bt2_pe.bam
do
    deduplicate_bismark -p --bam $fbam
done

# Creation of the folder for the output
mkdir -p /Volumes/DDMezger/methylome/fusion/align-lambda/bismark_methylation_extractor/

# Copy and paste deduplicated .bam files in folder 
#/Volumes/DDMezger/methylome/fusion/align-lambda/bismark_methylation_extractor/

#Extraction of methylation state using bismark_methylation_extractor 
cd /Volumes/DDMezger/methylome/fusion/align-lambda/bismark_methylation_extractor/

#!/bin/bash
for file in d_fus_PE_c10t_*M.bam
do
dir="/Volumes/DDMezger/methylome/fusion/align-lambda/bismark_methylation_extractor/"
bismark_methylation_extractor -p -o $dir --report --no_overlap --bedGraph --counts --comprehensive --buffer_size 50G $file
done
```

With a bisulfite conversion **ratio equal or higher than 97,8%** for every samples, bisulfite treatment worked correctly and data can be analyzed (Supp. Table S1 in Chapter 4 of the thesis, paper Duchateau et al., in preparation).

#### 8. Mapping on mouse genome (mm9)¶

Filtered reads were paired-end mapped on **Mus musculus reference genome mm9** using **bismark** mapper. Whole genome was used as a reference. As for mapping on lambda phage genome, mm9 genome must be prepared before mapping, to take into account read methylation profile. For the mapping, libraries were considered to be non directional.

In [ ]:

```
%%bash

# 8.1. Reference genome (mm9) preparation
bismark_genome_preparation 
--path_to_bowtie /Users/UMR7216/Desktop/Bioinfo_TOOLS/bowtie2-2.1.0 
--bowtie2 --verbose /Users/UMR7216/Documents/Mezger/methylome/genome-mm9

# 8.2. Mapping (mm9)
# Creation of the work directory
mkdir -p /Volumes/DD-Agathe/methylome/MALE-FUSION/align-mm9/1M
mkdir -p /Volumes/DD-Agathe/methylome/MALE-FUSION/align-mm9/2M
mkdir -p /Volumes/DD-Agathe/methylome/MALE-FUSION/align-mm9/3M
mkdir -p /Volumes/DD-Agathe/methylome/MALE-FUSION/align-mm9/4M
mkdir -p /Volumes/DD-Agathe/methylome/MALE-FUSION/align-mm9/5M
mkdir -p /Volumes/DD-Agathe/methylome/MALE-FUSION/align-mm9/6M

# Mapping
cd /Users/UMR7216/Documents/Mezger/methylome/cat-trim

bismark -N 1 /Users/UMR7216/Documents/Mezger/methylome/genome-mm9/ 
--path_to_bowtie /Users/UMR7216/Desktop/Bioinfo_TOOLS/bowtie2-2.1.0 
--non_directional --un -o /Volumes/DD-Agathe/methylome/MALE-FUSION/align-mm9/1M 
-1 /Users/UMR7216/Documents/Mezger/methylome/cat-trim/c10t_fus_1M_R1.fastq 
-2 /Users/UMR7216/Documents/Mezger/methylome/cat-trim/c10t_fus_1M_R2.fastq

bismark -N 1 /Users/UMR7216/Documents/Mezger/methylome/genome-mm9/ 
--path_to_bowtie /Users/UMR7216/Desktop/Bioinfo_TOOLS/bowtie2-2.1.0 
--non_directional --un -o /Volumes/DD-Agathe/methylome/MALE-FUSION/align-mm9/2M 
-1 /Users/UMR7216/Documents/Mezger/methylome/cat-trim/c10t_fus_2M_R1.fastq 
-2 /Users/UMR7216/Documents/Mezger/methylome/cat-trim/c10t_fus_2M_R2.fastq

bismark -N 1 /Users/UMR7216/Documents/Mezger/methylome/genome-mm9/ 
--path_to_bowtie /Users/UMR7216/Desktop/Bioinfo_TOOLS/bowtie2-2.1.0 
--non_directional --un -o /Volumes/DD-Agathe/methylome/MALE-FUSION/align-mm9/3M 
-1 /Users/UMR7216/Documents/Mezger/methylome/cat-trim/c10t_fus_3M_R1.fastq 
-2 /Users/UMR7216/Documents/Mezger/methylome/cat-trim/c10t_fus_3M_R2.fastq

bismark -N 1 /Users/UMR7216/Documents/Mezger/methylome/genome-mm9/ 
--path_to_bowtie /Users/UMR7216/Desktop/Bioinfo_TOOLS/bowtie2-2.1.0 
--non_directional --un -o /Volumes/DD-Agathe/methylome/MALE-FUSION/align-mm9/4M 
-1 /Users/UMR7216/Documents/Mezger/methylome/cat-trim/c10t_fus_4M_R1.fastq 
-2 /Users/UMR7216/Documents/Mezger/methylome/cat-trim/c10t_fus_4M_R2.fastq

bismark -N 1 /Users/UMR7216/Documents/Mezger/methylome/genome-mm9/ 
--path_to_bowtie /Users/UMR7216/Desktop/Bioinfo_TOOLS/bowtie2-2.1.0 
--non_directional --un -o /Volumes/DD-Agathe/methylome/MALE-FUSION/align-mm9/5M 
-1 /Users/UMR7216/Documents/Mezger/methylome/cat-trim/c10t_fus_5M_R1.fastq 
-2 /Users/UMR7216/Documents/Mezger/methylome/cat-trim/c10t_fus_5M_R2.fastq

bismark -N 1 /Users/UMR7216/Documents/Mezger/methylome/genome-mm9/ 
--path_to_bowtie /Users/UMR7216/Desktop/Bioinfo_TOOLS/bowtie2-2.1.0 
--non_directional --un -o /Volumes/DD-Agathe/methylome/MALE-FUSION/align-mm9/6M 
-1 /Users/UMR7216/Documents/Mezger/methylome/cat-trim/c10t_fus_6M_R1.fastq 
-2 /Users/UMR7216/Documents/Mezger/methylome/cat-trim/c10t_fus_6M_R2.fastq
```

#### 8.3. Deduplication¶

Filtered reads files were then compressed.

To obtain a more accurate estimation of bisulfite treatment efficiency, reads that mapped to mm9 genome were deduplicated, using **deduplicate\_bismark**.

In [ ]:

```
%%bash

# To compress files containing trimmed reads 
cd /Volumes/DDMezger/methylome/fusion/cat-trim/

#!/bin/bash
for fcut in c10t_fus*M_R*.fastq
do
zip ${fcut%.fastq}.fastq.zip $fcut
done

# To remove duplicates
#!/bin/bash
for fbam in /Volumes/DDMezger/methylome/fusion/align-mm9/*M/c10t_fus_*M_R1.fastq_bismark_bt2_pe.bam
do
deduplicate_bismark -p --bam $fbam
done
```

#### 9.Filename simplification¶

Name of files containing deduplicated mapped reads was simplified for subsequent analyses:

- c10t\_fus\_1M\_R1.fastq\_bismark\_bt2\_pe.bam was renamed **fus\_PE\_c10t\_1M.bam**
- c10t\_fus\_1M\_R1.fastq\_bismark\_bt2\_pe.deduplicated.bam -> **d\_fus\_PE\_c10t\_1M.bam**
- c10t\_fus\_2M\_R1.fastq\_bismark\_bt2\_pe.bam -> **fus\_PE\_c10t\_2M.bam**
- c10t\_fus\_2M\_R1.fastq\_bismark\_bt2\_pe.deduplicated.bam -> **d\_fus\_PE\_c10t\_2M.bam**
- c10t\_fus\_3M\_R1.fastq\_bismark\_bt2\_pe.bam -> **fus\_PE\_c10t\_3M.bam**
- c10t\_fus\_3M\_R1.fastq\_bismark\_bt2\_pe.deduplicated.bam -> **d\_fus\_PE\_c10t\_3M.bam**
- c10t\_fus\_4M\_R1.fastq\_bismark\_bt2\_pe.bam -> **fus\_PE\_c10t\_4M.bam**
- c10t\_fus\_4M\_R1.fastq\_bismark\_bt2\_pe.deduplicated.bam -> **d\_fus\_PE\_c10t\_4M.bam**
- c10t\_fus\_5M\_R1.fastq\_bismark\_bt2\_pe.bam -> **fus\_PE\_c10t\_5M.bam**
- c10t\_fus\_5M\_R1.fastq\_bismark\_bt2\_pe.deduplicated.bam -> **d\_fus\_PE\_c10t\_5M.bam**
- c10t\_fus\_6M\_R1.fastq\_bismark\_bt2\_pe.bam -> **fus\_PE\_c10t\_6M.bam**
- c10t\_fus\_6M\_R1.fastq\_bismark\_bt2\_pe.deduplicated.bam -> **d\_fus\_PE\_c10t\_6M.bam**

#### 10.Extraction of data corresponding to the methylome capture¶

To identify reads that map to captured regions, intersection between captured regions coordinates and mapped reads was investigated, using **bedtools intersect**. To avoid removal of potential interesting sequences, at the border of capture regions, each region was extended up and downstream with 150 bases, before the intersection (extended capture regions was named ***enlarged capture***). This allows to keep reads that mapped totally or partially in captured regions. **Excel** was used to systematically remove 150bp from start position, and add 150bp from end position of each captured region. **bedtools merge** was used to merge enlarged captured regions that overlap with each other.

To estimate proportion of reads that mapped captured regions, number of lines of file containing whole mapped reads were compared to number of lines of file containing reads included in the capture. For that, bam files have to be converted into sam files, using **samtools view**. sam files get a header of 37 lines. These lines were not take into account for estimating proportion of reads in the methylome capture (in other words, one read is represented by one line, except the 37 first lines corresponding to the header).

In [ ]:

```
%%bash

# -----------------
# Excel was used to add +/- 150bp at each captured region coordinate. 
# File containing enlarged captured regions is called Capture_76800_et150seq.bed
# Then, bedtools merge was used to merge enlarged captured regions that overlap with each other.

cd /Users/UMR7216/Documents/Mezger/capture-methylome/Diderot-CaptureEpi-2015-479-27264242/capture-regions

# Data must first be sorted by chromosome (-k1,1 option) and by start position (-k2,2 option)
sort -k1,1 -k2,2n Capture_76800_et150seq.bed>Capture_76800_et150seq.sorted.bed
bedtools merge -i Capture_76800_et150seq.sorted.bed> Capture_76800_et150seq.merge.sorted.bed


# -----------------
# Intersection between enlarged captured regions and mapped (+ deduplicated) reads

mkdir -p /Volumes/DDMezger/methylome/fusion/align-mm9/intersect-capture/1M
mkdir -p /Volumes/DDMezger/methylome/fusion/align-mm9/intersect-capture/2M
mkdir -p /Volumes/DDMezger/methylome/fusion/align-mm9/intersect-capture/3M
mkdir -p /Volumes/DDMezger/methylome/fusion/align-mm9/intersect-capture/4M
mkdir -p /Volumes/DDMezger/methylome/fusion/align-mm9/intersect-capture/5M
mkdir -p /Volumes/DDMezger/methylome/fusion/align-mm9/intersect-capture/6M

#!/bin/bash
for file in /Volumes/DDMezger/methylome/fusion/align-mm9/*M/d_fus_PE_c10t_*M.bam
do
director="/Volumes/DDMezger/methylome/fusion/align-mm9/intersect-capture/${file:45:2}/inter_d_fus_PE_c10t_${file:45:2}.bam"
bedtools intersect -abam $file 
-b /Users/UMR7216/Documents/Mezger/methylome/capture-regions/Capture_76800_et150seq.merge.sorted.bed > $director
done


# -----------------
# Proportion of mapped reads into captured regions: 
# convert .bam into .sam files and count the number of lines
# input = deduplicated reads (all, not only those included in a capture region)
#!/bin/bash
for filebam in /Volumes/DDMezger/methylome/fusion/align-mm9/*M/d_fus_PE_c10t_*M.bam
do
samtools view -h -o ${filebam%.bam}.sam $filebam
done

#!/bin/bash 
for filesam in /Volumes/DDMezger/methylome/fusion/align-mm9/*M/d_fus_PE_c10t_*M.sam
do
wc -l $filesam
done

#Number of lines without header (which is composed of 37 lines)  
# 46 595 798 lines for 1M
# 53 377 408 lines for 2M
# 53 191 184 lines for 3M
# 53 179 200 lines for 4M
# 58 586 780 lines for 5M
# 60 938 390 lines for 6M
# As expected, it corresponds to the number ofdeduplicated reads pairs (R1+R2), if we divided these numbers by two. 

# input = deduplicated reads that are in captured regions
#!/bin/bash
for filebam in /Volumes/DDMezger/methylome/fusion/align-mm9/intersect-capture/*M/inter_d_fus_PE_c10t_*M.bam
do
samtools view -h -o ${filebam%.bam}.sam $filebam
done

#!/bin/bash 
for filesam in /Volumes/DDMezger/methylome/fusion/align-mm9/intersect-capture/*M/inter_d_fus_PE_c10t_*M.sam
do
wc -l $filesam
done

#Number of lines without header (which is composed of 37 lines) 
# 26 727 731 lines for 1M, meaning that 57,36 % of mapped reads are into the capture.
# 29 559 673 lines for 2M, meaning that 55,38 % of mapped reads are into the capture.
# 29 150 857 lines for 3M, meaning that 54,80 % of mapped reads are into the capture.
# 31 607 993 lines for 4M, meaning that 59,44% of mapped reads are into the capture.
# 34 767 160 lines for 5M, meaning that 59,34% of mapped reads are into the capture.
# 37 293 752 lines for 6M, meaning that 61,20% of mapped reads are into the capture.
```

#### 11.Visualization of the mapping¶

**IGV** was used to visualize mapped reads. In order to use it, files containing mapped reads were sorted and indexed by **samtools sort** and **samtools index** respectively.

Using IGV, methylation state can be visualized:

- *blue* color reprensented *unmethylated* cytosines, in CpG context.
- *red* color represented *methylated* cytosines, in CpG context.

In [ ]:

```
%%bash

# ----------------------
# To sort files containing whole mapped reads 
# (not only those that are in captured regions)
#!/bin/bash
for fbam in /Volumes/DDMezger/methylome/fusion/align-mm9/*M/*PE_c10t_*M.bam
do
samtools sort $fbam ${fbam%.bam}_sorted
done

# To index mapped reads
#!/bin/bash
for fbam in /Volumes/DDMezger/methylome/fusion/align-mm9/*M/*PE_c10t_*M_sorted.bam
do
samtools index $fbam ${fbam%.bam}.bai 
done 


# -----------------
# Verification of captured selection on IGV
# Fot that, files must be sorted and indexed

# To sort files containing reads that are located in captured regions
#!/bin/bash
for fbam in /Volumes/DDMezger/methylome/fusion/align-mm9/intersect-capture/*M/inter_d_fus_PE_c10t_*M.bam
do
samtools sort $fbam ${fbam%.bam}_sorted
done

# To index mapped reads
samtools index input_sorted.bam ouput_sorted.bai
#!/bin/bash
for fbam in /Volumes/DDMezger/methylome/fusion/align-mm9/intersect-capture/*M/inter_d_fus_PE_c10t_*M_sorted.bam
do
samtools index $fbam ${fbam%.bam}.bai 
done
```

#### 12.Methylation state extraction¶

**bismark\_methylation\_extractor (BME)** was used to obtain methylation profile of mapped cytosines in CpG context only. bismark\_methylation\_extractor was runned on files containing **whole mapped reads**, not only those observed in captured regions because BME doesn't work properly with files that are organized differently from bismark output. It could be due to the fact that BME can't identify paired-end reads anymore if reads are reorganized / splitted by another tool. Thus, selection of CpG that are in captured regions is done after.

**Detailed options for BME:**

- -p: to specify that input file is Bismark result file generated from **paired-end** read data.
- -o: to define ouput directory
- --report: to obtain a report
- --no\_overlap: to take into account paired-end reads and prevent double counting of paired-end relsults, when R1 and R2 reads overlapped.
- --bedGraph: to obtain a bedGraph
- --counts: to count number of unmethylated and methylated cytosines, for a given CpG site.
- --comprehensive: to obtain CpG state of all strands (OT, OB, CTOT and CTOB strand, meaning respectively original top strand, original bottom strand, complementary to OT stand or complementary to OB strand), not only OT.
- --buffer\_size 50G: This allows to specify the main memory sort buffer when sorting the methylation information. Default to 2G.

The bedGraph output (tab-delimited; 0-based start coords, 1-based end coords) contains following information:

- chromosome
- start position
- end position
- methylation percentage

The coverage output (tab-delimited, 1-based genomic coords) contains following information:

- chromosome
- start position
- end position
- methylation percentage
- count of methylated CpG
- count of non-methylated CpG

In [ ]:

```
%%bash

# To create folders that will contain bismark_methylation_extractor results
mkdir -p /Volumes/DDMezger/methylome/fusion/etat_methylation/bismark_methylation_extractor/1M
mkdir -p /Volumes/DDMezger/methylome/fusion/etat_methylation/bismark_methylation_extractor/2M
mkdir -p /Volumes/DDMezger/methylome/fusion/etat_methylation/bismark_methylation_extractor/3M
mkdir -p /Volumes/DDMezger/methylome/fusion/etat_methylation/bismark_methylation_extractor/4M
mkdir -p /Volumes/DDMezger/methylome/fusion/etat_methylation/bismark_methylation_extractor/5M
mkdir -p /Volumes/DDMezger/methylome/fusion/etat_methylation/bismark_methylation_extractor/6M

#Files containing deduplicated rads were copied into the following folder : 
#/Volumes/DDMezger/methylome/fusion/align-mm9/bam/

cd /Volumes/DDMezger/methylome/fusion/align-mm9/bam/

#!/bin/bash
for file in d_fus_PE_c10t_*M.bam
do
dir="/Volumes/DDMezger/methylome/fusion/etat_methylation/bismark_methylation_extractor/${file:14:2}/"
bismark_methylation_extractor -p -o $dir --report --no_overlap 
--bedGraph --counts --comprehensive --buffer_size 50G $file
done
```

#### 13.Selection to only analyse CpG that are in captured regions¶

To only work with captured regions, .cov files from bismark\_methylation\_extractor (i.e. files containing methylated / unmethylated cytosines count per sample and per cytosine site in CpG context), were intersect with file containing enlarged capture regions, using **bedtools intersect**.

In [ ]:

```
%%bash

# To create folders that will contain outputs
mkdir -p /Volumes/DDMezger/methylome/fusion/etat_methylation/bismark_methylation_extractor/1M/capt
mkdir -p /Volumes/DDMezger/methylome/fusion/etat_methylation/bismark_methylation_extractor/2M/capt
mkdir -p /Volumes/DDMezger/methylome/fusion/etat_methylation/bismark_methylation_extractor/3M/capt
mkdir -p /Volumes/DDMezger/methylome/fusion/etat_methylation/bismark_methylation_extractor/4M/capt
mkdir -p /Volumes/DDMezger/methylome/fusion/etat_methylation/bismark_methylation_extractor/5M/capt
mkdir -p /Volumes/DDMezger/methylome/fusion/etat_methylation/bismark_methylation_extractor/6M/capt

# .cov.zip repositories from bismark_methylation_extractor were copied into the following folder :
# /Volumes/DDMezger/methylome/fusion/etat_methylation/coverage2cytosine/covdezip
# nb: it could be confusing because coverage2cytosine is also a tool from bismark suite, but .cov files actually come from BME.

# .cov files were then unziped to run bedtools intersect 

cd /Volumes/DDMezger/methylome/fusion/etat_methylation/coverage2cytosine/covdezip

#!/bin/bash
for file in d_fus_PE_c10t_*M.bismark.cov
do
director="inter_d_fus_PE_c10t_${file:14:2}.cov"
bedtools intersect -abam $file 
-b /Users/UMR7216/Documents/Mezger/methylome/capture-regions/Capture_76800_et150seq.merge.sorted.bed > $director
done
# nb: it would have been better to use -a parameter whether than -abam one, 
# since .cov files are not .bam files. 
# However, same results are obtained with -a or -abam parameters.

cd /Volumes/DDMezger/methylome/fusion/etat_methylation/coverage2cytosine/covdezip

wc -l inter_d_fus_PE_c10t_1M.cov 
# 3 327 263 inter_d_fus_PE_c10t_1M.cov, 
# meaning that 3 327 263 CpG sites that are included in a given captured region, are covered.

wc -l inter_d_fus_PE_c10t_2M.cov 
# 3 371 259 inter_d_fus_PE_c10t_2M.cov

wc -l inter_d_fus_PE_c10t_3M.cov 
# 3 384 718 inter_d_fus_PE_c10t_3M.cov

wc -l inter_d_fus_PE_c10t_4M.cov 
# 3 329 265 inter_d_fus_PE_c10t_4M.cov

wc -l inter_d_fus_PE_c10t_5M.cov 
# 3 396 821 inter_d_fus_PE_c10t_5M.cov

wc -l inter_d_fus_PE_c10t_6M.cov 
# 3 403 563 inter_d_fus_PE_c10t_6M.cov
```

#### 14.Statistical analysis¶

**methylkit** (R package) was used to perform a statistical analysis of methylome data.

##### 14.1. Data formatting¶

.cov files from **bismark\_methylation\_extractor** were copied into a specific folder for the statistical analysis. Then, data was formatted in order to use **methylkit** properly.

At the end of **bismark\_methylation\_extractor**, data are organized in a dataframe as follow:

- chr: chromosome
- start: position of the cytosine
- end: position of the cytosine
- %meth: methylation percent
- count C: number of unconverted cytosines (*i.e.* methylated cytosines for a given position)
- count T: number of converted cytosines (*i.e.* unmethylated cytosines for a given position)

To use **methylKit**, data must be organized in a dataframe as follow:

- chrBase: ID of the position
- chr: chromosome
- base: position of the cytosine
- strand
- coverage: total number of C (*i.e.* unconverted cytosines) and T (*i.e.* converted cytosines) counted at a given position
- freqC: frequence of observed C, meaning ratio of the number of unconverted cytosines (methylated cytosines) and the total cytosine count (C+T) at the position.
- freqT: frequence of observed T, meaning ratio of the number of converted cytosines (unmethylated cytosines) and the total cytosine count (C+T) at the position.

Since we don't have strand informations in .cov files from **bismark\_methylation\_extractor**, we defined a fictive strand for each CpG position.

In [ ]:

```
%%bash 

# To create work directory
mkdir -p /Volumes/DDMezger/methylome/fusion/methylkit/capture

# Copy and paste .cov files from BME, named inter_d_fus_PE_c10t_*M.cov 
# into /Volumes/DDMezger/methylome/fusion/methylkit/capture
```

In [ ]:

```
%%R

# ---------------------------
# Data formatting
# For 1M sample: 
# ---------------------------

setwd ("/Volumes/DDMezger/methylome/fusion/methylkit/capture/")

# To load input file (.cov file)
Data01MAI <- read.table("/Volumes/DDMezger/methylome/fusion/methylkit/capture/inter_d_fus_PE_c10t_1M.cov", header=FALSE, sep="", col.names=c("chr", "base","end","methpercent","countmeth", "countunmeth"), na.strings="NA", dec=".", strip.white=TRUE)

# To check first rows to verify the loading
head(Data01MAI)

# To copy the R object 
Data1MAI = Data01MAI

# To calculate the coverage 
Data1MAI$coverage <- with(Data1MAI, countmeth+countunmeth)

# Verification  
head(Data1MAI)

# To calculate T or C frequencies 
Data1MAI$freqCL <- with (Data1MAI, (countmeth/coverage)*100) 
Data1MAI$freqTL <- with (Data1MAI, (countunmeth/coverage)*100)

# To modify freqC and freqT format (decimal values rather than integers),
# to avoid troubles when loading data on methylKit
Data1MAI$freqC = with (Data1MAI,format(Data1MAI$freqCL,digits=2))
Data1MAI$freqT = with (Data1MAI,format(Data1MAI$freqTL,digits=2))

# Verification 
head(Data1MAI)

# To create chr.Base column 
Data1MAI$chrBase <- with(Data1MAI, paste(Data1MAI$chr,Data1MAI$base,sep="."))

# Verification 
head(Data1MAI)

# To create a column with strand 
# /!\ this strand is not necessary the real one
Data1MAI$strand <- with(Data1MAI, paste("R"))

# Verification 
head(Data1MAI)

# To generate a dataframe containing only columns of interest
Data1MBI= Data1MAI[,c("chrBase","chr","base","strand","coverage","freqC","freqT")]

# Verification 
head(Data1MBI)

# To save the dataframe
write.table(Data1MBI, file ="cov1_capt_CpG_dc10t_fus_1M.txt", sep=" ", 
            quote = FALSE, na="NA", dec = ".", row.names=FALSE, col.names=TRUE)
# quote=FALSE option to avoid enclosing non-numeric data in quotation marks in the output files.
# col.names=TRUE option is necessary to add an header into the output file.


#---------------------------------------------------------
#Same process for other samples (2M, 3M, 4M, 5M and 6M)
```

##### 14.2. Data loading on methylkit and quality controls¶

Then, files containing methylation state of cytosines in CpG context were loaded on **methylKit**.
  
  
Two quality controls of the data were done:

- **getMethylationStats()**: this command plots an histogram of methylation percent distribution. Using genome-wide dataset, two peaks on both ends of the histogram are expected, since in a given cell, cytosines are either methylated or not. Therefore, if many cells are analysed, similar pattern will be observed, with lot of cytosine position with high methylation and lot of methylation with low methylation.
- **getCoverageStats()**: this command plots an histogram of read coverage per cytosine position. If a peak is observed on the right end of the histogram, it indicates that somes PCR duplicates may exist and can biased the statistical analysis.

In [ ]:

```
%%R

# ---------------------------
# Data loading on methylKit
# ---------------------------

library(methylKit)

setwd("/Volumes/DDMerger/methylome/fusion/methylkit/capture/")

# Data loading
male.list.capt=list("cov1_capt_CpG_dc10t_fus_1M.txt", "cov1_capt_CpG_dc10t_fus_2M.txt", 
                    "cov1_capt_CpG_dc10t_fus_3M.txt", "cov1_capt_CpG_dc10t_fus_4M.txt", 
                    "cov1_capt_CpG_dc10t_fus_5M.txt", "cov1_capt_CpG_dc10t_fus_6M.txt")

Captnf = read(male.list.capt, sample.id=list("1M","2M","3M","4M","5M","6M"), 
              assembly="mm9", header = TRUE, context="CpG", 
              resolution = "base", treatment=c(1,1,1,0,0,0))

Captnf

setwd("/Volumes/DDMezger/methylome/fusion/methylkit/capture")

library(methylKit)

M.list.captBME=list("cov1_capt_CpG_dc10t_fus_1M.txt", 
                    "cov1_capt_CpG_dc10t_fus_2M.txt",
                    "cov1_capt_CpG_dc10t_fus_3M.txt", 
                    "cov1_capt_CpG_dc10t_fus_4M.txt", 
                    "cov1_capt_CpG_dc10t_fus_5M.txt", 
                    "cov1_capt_CpG_dc10t_fus_6M.txt")

# ------------------------------------------
# Quality control n°1 - getMethylationStats
# ------------------------------------------

#Default parameters - .png output
png(filename="methylation_stat_capt1Mfus_nf.png")
getMethylationStats(Captnf[[1]],plot=T,both.strands=F) 
dev.off()

png(filename="methylation_stat_capt2Mfus_nf.png")
getMethylationStats(Captnf[[2]],plot=T,both.strands=F) 
dev.off()
 
png(filename="methylation_stat_capt3Mfus_nf.png")
getMethylationStats(Captnf[[3]],plot=T,both.strands=F) 
dev.off()
 
png(filename="methylation_stat_capt4Mfus_nf.png")
getMethylationStats(Captnf[[4]],plot=T,both.strands=F) 
dev.off()
 
png(filename="methylation_stat_capt5Mfus_nf.png")
getMethylationStats(Captnf[[5]],plot=T,both.strands=F) 
dev.off()
 
png(filename="methylation_stat_capt6Mfus_nf.png")
getMethylationStats(Captnf[[6]],plot=T,both.strands=F) 
dev.off()

#for nclass=10 - .png output
png(filename="methylation_stat_capt1Mfus_nf_class10.png")
getMethylationStats(Captnf[[1]],plot=T,both.strands=F,nclass=10) 
dev.off()
 
png(filename="methylation_stat_capt2Mfus_nf_class10.png")
getMethylationStats(Captnf[[2]],plot=T,both.strands=F,nclass=10) 
dev.off()
 
png(filename="methylation_stat_capt3Mfus_nf_class10.png")
getMethylationStats(Captnf[[3]],plot=T,both.strands=F,nclass=10) 
dev.off()

png(filename="methylation_stat_capt4Mfus_nf_class10.png")
getMethylationStats(Captnf[[4]],plot=T,both.strands=F,nclass=10) 
dev.off()
 
png(filename="methylation_stat_capt5Mfus_nf_class10.png")
getMethylationStats(Captnf[[5]],plot=T,both.strands=F,nclass=10) 
dev.off()
 
png(filename="methylation_stat_capt6Mfus_nf_class10.png")
getMethylationStats(Captnf[[6]],plot=T,both.strands=F,nclass=10) 
dev.off()

#Default parameters - .emf output
library(devEMF)

emf(file="methylation stat capt 1M fus nf.emf")
getMethylationStats(Captnf[[1]],plot=T,both.strands=F) 
dev.off()

emf(file="methylation stat capt 2M fus nf.emf")
getMethylationStats(Captnf[[2]],plot=T,both.strands=F)
dev.off()

emf(file="methylation stat capt 3M fus nf.emf")
getMethylationStats(Captnf[[3]],plot=T,both.strands=F) 
dev.off()

emf(file="methylation stat capt 4M fus nf.emf")
getMethylationStats(Captnf[[4]],plot=T,both.strands=F) 
dev.off()

emf(file="methylation stat capt 5M fus nf.emf")
getMethylationStats(Captnf[[5]],plot=T,both.strands=F) 
dev.off()

emf(file="methylation stat capt 6M fus nf.emf")
getMethylationStats(Captnf[[6]],plot=T,both.strands=F) 
dev.off()

#for nclass=10 - .emf output
emf(file="methylation stat capt 1M fus nf class10.emf")
getMethylationStats(Captnf[[1]],plot=T,both.strands=F,nclass=10)
dev.off()

emf(file="methylation stat capt 2M fus nf class10.emf")
getMethylationStats(Captnf[[2]],plot=T,both.strands=F,nclass=10)
dev.off()

emf(file="methylation stat capt 3M fus nf class10.emf")
getMethylationStats(Captnf[[3]],plot=T,both.strands=F,nclass=10)
dev.off()

emf(file="methylation stat capt 4M fus nf class10.emf")
getMethylationStats(Captnf[[4]],plot=T,both.strands=F,nclass=10)
dev.off()

emf(file="methylation stat capt 5M fus nf class10.emf")
getMethylationStats(Captnf[[5]],plot=T,both.strands=F,nclass=10)
dev.off()

emf(file="methylation stat capt 6M fus nf class10.emf")
getMethylationStats(Captnf[[6]],plot=T,both.strands=F,nclass=10)
dev.off()

#Tables
getMethylationStats(Captnf[[1]],plot=F,both.strands=F)
getMethylationStats(Captnf[[2]],plot=F,both.strands=F)
getMethylationStats(Captnf[[3]],plot=F,both.strands=F)
getMethylationStats(Captnf[[4]],plot=F,both.strands=F)
getMethylationStats(Captnf[[5]],plot=F,both.strands=F)
getMethylationStats(Captnf[[6]],plot=F,both.strands=F) 


# ------------------------------------------
# Quality control n°2 - getCoverageStats
# ------------------------------------------

#Default parameters - .png output
png(filename="coverage Stat capt1M fus nf.png")
getCoverageStats(Captnf[[1]],plot=T,both.strands=F) 
dev.off()

png(filename="coverage Stat capt2M fus nf.png")
getCoverageStats(Captnf[[2]],plot=T,both.strands=F)
dev.off()
 
png(filename="coverage Stat capt3M fus nf.png")
getCoverageStats(Captnf[[3]],plot=T,both.strands=F)
dev.off()
 
png(filename="coverage Stat capt4M fus nf.png")
getCoverageStats(Captnf[[4]],plot=T,both.strands=F)
dev.off()
 
png(filename="coverage Stat capt5M fus nf.png")
getCoverageStats(Captnf[[5]],plot=T,both.strands=F)
dev.off()
 
png(filename="coverage Stat capt6M fus nf.png")
getCoverageStats(Captnf[[6]],plot=T,both.strands=F)
dev.off()

#Default parameters - .emf output
emf(file="coverage Stat capt1M fus nf.emf")
getCoverageStats(Captnf[[1]],plot=T,both.strands=F) 
dev.off()

emf(file="coverage Stat capt2M fus nf.emf")
getCoverageStats(Captnf[[2]],plot=T,both.strands=F)
dev.off()

emf(file="coverage Stat capt3M fus nf.emf")
getCoverageStats(Captnf[[3]],plot=T,both.strands=F)
dev.off()

emf(file="coverage Stat capt4M fus nf.emf")
getCoverageStats(Captnf[[4]],plot=T,both.strands=F)
dev.off()

emf(file="coverage Stat capt5M fus nf.emf")
getCoverageStats(Captnf[[5]],plot=T,both.strands=F)
dev.off()

emf(file="coverage Stat capt6M fus nf.emf")
getCoverageStats(Captnf[[6]],plot=T,both.strands=F)
dev.off()
```

###### Conclusions:¶

- For **getMethylationStats**, histogram profiles of the captured data are similar than those expected for genome-wide dataset (*i.e.* high number of CpG that are never methylated, and high number of CpG that are always methylated). In our captured data, there are more CpG that are never methylated than CpG that are always methylated. This imbalance can be due to the choice of specific genomic regions in the capture. Indeed, capture is composed of promoters that are majorly unmethylated in mouse genome.
- For **getCoverageStats**, no peak is observed on the right end of each histogram, meaning that deduplication step works properly (there are no PCR duplicates, thus no biases for the statistical analysis at this level).

##### 14.3. Data filtering and quality controls of filtered data¶

To increase the power of statistical tests and avoid biases, data were filtered:

- cytosine positions that are abnormal too much covered (**> 99.9th percentile**) were excluded from the analysis. Indeed, regions displaying abnormally high read coverage can comprise a sizeable proportion of the experiment and thereby introduce considerable bias (Kruger et al., 2012).
- cytosine position displaying low read coverage (**number of reads < 10**) were also excluded from the analysis.

Then, read coverage was **normalized** before statistical analysis, using median to calculate scaling factor. Same quality controls than those done on unfiltered data were performed on filtered data.

In [ ]:

```
%%R

# ------------------------------------------
# To filter the data
# ------------------------------------------

# To remove cytosines that are not enough (lo.count) or too much covered (hi.perc)
Captfmincov10= filterByCoverage(Captnf, lo.count=10, hi.perc=99.9)

# To normalize the coverage of the data
normCaptfmincov10= normalizeCoverage(Captfmincov10, method="median")

normCaptfmincov10


# ------------------------------------------
# Quality control n°1 : getMethylationStats
# ------------------------------------------

#Default parameters - .png output
png(filename="methylation stat capt1M fus f et normalise mincov10.png")
getMethylationStats(normCaptfmincov10[[1]],plot=T,both.strands=F) 
dev.off()

png(filename="methylation stat capt2M fus f et normalise mincov10.png")
getMethylationStats(normCaptfmincov10[[2]],plot=T,both.strands=F) 
dev.off()

png(filename="methylation stat capt3M fus f et normalise mincov10.png")
getMethylationStats(normCaptfmincov10[[3]],plot=T,both.strands=F) 
dev.off()

png(filename="methylation stat capt4M fus f et normalise mincov10.png")
getMethylationStats(normCaptfmincov10[[4]],plot=T,both.strands=F) 
dev.off()

png(filename="methylation stat capt5M fus f et normalise mincov10.png")
getMethylationStats(normCaptfmincov10[[5]],plot=T,both.strands=F) 
dev.off()

png(filename="methylation stat capt6M fus f et normalise mincov10.png")
getMethylationStats(normCaptfmincov10[[6]],plot=T,both.strands=F) 
dev.off()


#for nclass=10 - .png output
png(filename="methylation stat capt1M fus f et normalise mincov10nclass10.png")
getMethylationStats(normCaptfmincov10[[1]],plot=T,both.strands=F,nclass=10)
dev.off()
 
png(filename="methylation stat capt2M fus f et normalise mincov10nclass10.png")
getMethylationStats(normCaptfmincov10[[2]],plot=T,both.strands=F,nclass=10)
dev.off()

png(filename="methylation stat capt3M fus f et normalise mincov10nclass10.png")
getMethylationStats(normCaptfmincov10[[3]],plot=T,both.strands=F,nclass=10)
dev.off()
 
png(filename="methylation stat capt4M fus f et normalise mincov10nclass10.png")
getMethylationStats(normCaptfmincov10[[4]],plot=T,both.strands=F,nclass=10)
dev.off()
 
png(filename="methylation stat capt5M fus f et normalise mincov10nclass10.png")
getMethylationStats(normCaptfmincov10[[5]],plot=T,both.strands=F,nclass=10)
dev.off()
 
png(filename="methylation stat capt6M fus f et normalise mincov10nclass10.png")
getMethylationStats(normCaptfmincov10[[6]],plot=T,both.strands=F,nclass=10)
dev.off()

#Default parameters - .emf output
library(devEMF)

emf(file="methylation stat capt1M fus f et normalise mincov10.emf")
getMethylationStats(normCaptfmincov10[[1]],plot=T,both.strands=F) 
dev.off()

emf(file="methylation stat capt2M fus f et normalise mincov10.emf")
getMethylationStats(normCaptfmincov10[[2]],plot=T,both.strands=F) 
dev.off()

emf(file="methylation stat capt3M fus f et normalise mincov10.emf")
getMethylationStats(normCaptfmincov10[[3]],plot=T,both.strands=F) 
dev.off()

emf(file="methylation stat capt4M fus f et normalise mincov10.emf")
getMethylationStats(normCaptfmincov10[[4]],plot=T,both.strands=F) 
dev.off()

emf(file="methylation stat capt5M fus f et normalise mincov10.emf")
getMethylationStats(normCaptfmincov10[[5]],plot=T,both.strands=F) 
dev.off()

emf(file="methylation stat capt6M fus f et normalise mincov10.emf")
getMethylationStats(normCaptfmincov10[[6]],plot=T,both.strands=F) 
dev.off()


#for nclass=10 - .emf output 
emf(file="methylation stat capt1M fus f et normalise mincov10 nclass10.emf")
getMethylationStats(normCaptfmincov10[[1]],plot=T,both.strands=F,nclass=10)
dev.off()

emf(file="methylation stat capt2M fus f et normalise mincov10 nclass10.emf")
getMethylationStats(normCaptfmincov10[[2]],plot=T,both.strands=F,nclass=10)
dev.off()

emf(file="methylation stat capt3M fus f et normalise mincov10 nclass10.emf")
getMethylationStats(normCaptfmincov10[[3]],plot=T,both.strands=F,nclass=10)
dev.off()

emf(file="methylation stat capt4M fus f et normalise mincov10 nclass10.emf")
getMethylationStats(normCaptfmincov10[[4]],plot=T,both.strands=F,nclass=10)
dev.off()

emf(file="methylation stat capt5M fus f et normalise mincov10 nclass10.emf")
getMethylationStats(normCaptfmincov10[[5]],plot=T,both.strands=F,nclass=10)
dev.off()

emf(file="methylation stat capt6M fus f et normalise mincov10 nclass10.emf")
getMethylationStats(normCaptfmincov10[[6]],plot=T,both.strands=F,nclass=10)
dev.off()

#Tables
getMethylationStats(normCaptfmincov10[[1]],plot=F,both.strands=F)
getMethylationStats(normCaptfmincov10[[2]],plot=F,both.strands=F) 
getMethylationStats(normCaptfmincov10[[3]],plot=F,both.strands=F) 
getMethylationStats(normCaptfmincov10[[4]],plot=F,both.strands=F) 
getMethylationStats(normCaptfmincov10[[5]],plot=F,both.strands=F) 
getMethylationStats(normCaptfmincov10[[6]],plot=F,both.strands=F) 


# ------------------------------------------
# Quality control n°2 : getCoverageStats
# ------------------------------------------

#Default parameters - .png output
png(filename="coverage Stat capt1M fus f et normalise mincov10.png")
getCoverageStats(normCaptfmincov10[[1]],plot=T,both.strands=F) 
dev.off()
 
png(filename="coverage Stat capt2M fus f et normalise mincov10.png")
getCoverageStats(normCaptfmincov10[[2]],plot=T,both.strands=F)
dev.off()

png(filename="coverage Stat capt3M fus f et normalise mincov10.png")
getCoverageStats(normCaptfmincov10[[3]],plot=T,both.strands=F)
dev.off()
 
png(filename="coverage Stat capt4M fus f et normalise mincov10.png")
getCoverageStats(normCaptfmincov10[[4]],plot=T,both.strands=F)
dev.off()

png(filename="coverage Stat capt5M fus f et normalise mincov10.png")
getCoverageStats(normCaptfmincov10[[5]],plot=T,both.strands=F)
dev.off()
 
png(filename="coverage Stat capt6M fus f et normalise mincov10.png")
getCoverageStats(normCaptfmincov10[[6]],plot=T,both.strands=F)
dev.off()
 

#Default parameters - .emf output
emf (file="coverage Stat capt1M fus f et normalise mincov10.emf")
getCoverageStats(normCaptfmincov10[[1]],plot=T,both.strands=F) 
dev.off()

emf(file="coverage Stat capt2M fus f et normalise mincov10.emf")
getCoverageStats(normCaptfmincov10[[2]],plot=T,both.strands=F)
dev.off()

emf(file="coverage Stat capt3M fus f et normalise mincov10.emf")
getCoverageStats(normCaptfmincov10[[3]],plot=T,both.strands=F)
dev.off()

emf(file="coverage Stat capt4M fus f et normalise mincov10.emf")
getCoverageStats(normCaptfmincov10[[4]],plot=T,both.strands=F)
dev.off()

emf(file="coverage Stat capt5M fus f et normalise mincov10.emf")
getCoverageStats(normCaptfmincov10[[5]],plot=T,both.strands=F)
dev.off()

emf(file="coverage Stat capt6M fus f et normalise mincov10.emf")
getCoverageStats(normCaptfmincov10[[6]],plot=T,both.strands=F)
dev.off()
```

###### Conclusions (globally similar than unfiltered and unormalized data):¶

- For **getMethylationStats**, histogram profiles of the captured data are similar than those expected for genome-wide dataset (*i.e.* high number of CpG that are never methylated, and high number of CpG that are always methylated). In our captured data, there are more CpG that are never methylated than CpG that are always methylated. This imbalance can be due to the choice of specific genomic regions in the capture. Indeed, capture is composed of promoters that are majorly unmethylated in mouse genome.
- For **getCoverageStats**, no peak is observed on the right end of each histogram, meaning that deduplication step works properly (there are no PCR duplicates, thus no biases for the statistical analysis at this levele).

##### 14.4. Creation of a MethylBase object¶

To compare samples and identify differentially methylated cytosines (**DMCs**) upon alcohol exposure, a methylBase object must be created. It is a table containing methylation state informations of cytosines that are in all samples.

In [ ]:

```
%%R

# methylBase object containing only cytosines that are covered in aLL samples:
methcaptnormcov10strict=methylKit::unite(normCaptfmincov10,destrand=FALSE)

methcaptnormcov10strict
# 1 259 111  rows = number of CpG that are covered (at least 10 times) in aLL samples

write.table(methcaptnormcov10strict, file ="pool-strict-CpG-commun-capt-norm-mincov10-99.9-male_fus.txt", sep=" ", quote = FALSE, na="NA", dec = ".", row.names=FALSE, col.names=TRUE)
```

##### 14.5. Sample comparison¶

Correlation between samples were estimated using **getCorrelation** function of **methylkit**. Dendrogram (representing
sample clusters) and PCA analysis were done using, respectively, **clusterSample** and **PCASamples** functions of **methylkit** package.

In [ ]:

```
%%R

# ------------------------------------------
# Correlation between samples
# ------------------------------------------

setwd("/Volumes/DDMezger/methylome/fusion/methylkit/capture/Captf-mincov10-norm")

png(filename="getCorrelation male fus strict capt norm mincov10 hi.perc 99.9.png")
getCorrelation(methcaptnormcov10strict,plot=TRUE)
dev.off()

# ------------------------------------------
# Dendrogram - default parameters
# ------------------------------------------
png(filename="clusterSamples methstrict fus ward dist correl capt norm mincov10 hi.perc99.9.png")
clusterSamples(methcaptnormcov10strict, dist="correlation", method="ward", plot=TRUE)
dev.off()

# ------------------------------------------
# PCA analysis - default parameters
# ------------------------------------------
png(filename="PCASamples (defaut) methstrict fus capt norm mincov10 hi.perc 99.9.png")
PCASamples(methcaptnormcov10strict, screeplot = FALSE, adj.lim=c(0.0004,0.1), 
           scale=TRUE, center=TRUE, comp=c(1,2),transpose=TRUE, sd.threshold=0, obj.return=FALSE)
dev.off()
```

##### 14.6. Research of differentially methylated CpG (DMCs) upon alcohol exposure¶

For detection of significant differentially methylated isolated CpG (**DMCs**) between control and alcohol-treated samples, filtered data were analysed with **methylKit**. Only methylation state of CpG sites that were covered in all samples was investigated for the analysis (corresponding to 1,259,111 positions).
  
  
DMCs were considered as significant when difference of methylation state between the two groups was **higher than 5%**, for a **qvalue threshold of 0.05**.

In [ ]:

```
%%R

#----------------------------------------------------
# Identification of DMCs among methylBase object containing 
# only cytosines that are covered in aLL samples
#----------------------------------------------------
Captnormcov10DiffCstrict=calculateDiffMeth(methcaptnormcov10strict)
Captnormcov10DiffCstrict

Captnormcov10Chypostrict=get.methylDiff(Captnormcov10DiffCstrict, difference=5, qvalue=0.05,type="hypo")
Captnormcov10Chypostrict

Captnormcov10ChyperCstrict=get.methylDiff(Captnormcov10DiffCstrict, difference=5, qvalue=0.05,type="hyper")
Captnormcov10ChyperCstrict

write.table(Captnormcov10Chypostrict, file ="capt-norm-mincov10-hypoCstrict-malefus.txt", 
            sep=" ", quote = FALSE, na="NA", dec = ".", row.names=FALSE, col.names=TRUE)

write.table(Captnormcov10hyperCstrict, file ="capt-norm-mincov10-hyperCstrict-malefus.txt", 
            sep=" ", quote = FALSE, na="NA", dec = ".", row.names=FALSE, col.names=TRUE)
```

##### 14.7. Research of differentially methylated regions (DMRs)¶

Due to specificity of methylome tailor-made capture we studied, we generated our own R function, called **get\_close\_closi()**
to identify differentially methylated regions (DMRs, Supp. Fig. S1B-D of the paper in preparation, Chapter 4, section 1.2, collab. Sascha Ott, Warwick University). This function combines neighbouring CpGs that share similar differential methylation state, (all hypo- or all hypermethylated CpGs, Supp. Fig. S1B and S1D of the paper in preparation, Chapter 4, section 1.2). Because MethylKit has been designed for methylome data and provides relevant statistical information about individual CpG, we used **MethylKit** statistical output as an input of get\_close\_loci() function (*i.e.* position of the CpG site, percent of methylation difference between groups for a given position, pvalue and qvalue).
  
  
With the same parameters than those used for DMCs detection, formatted, filtered and normalized data were used for DMRs detection. Only methylation state of CpG sites that were covered in all samples was investigated for the analysis (corresponding to 1,259,111 positions). To define DMRs with get\_close\_loci(), only CpG sites having a reliable methylation state must be considered, since one non relevant CpG is sufficient to affect DMR detection (Supp. Fig. S1D of the paper in preparation, Chapter 4, section 1.2). Thus a **pre-selection** of relevant CpGs was done, according to their **pvalue**. This pvalue threshold was determined using **comparisons of real and random datasets** (see below, Supp. Fig. S1E, Supp. Table 2 of the paper in preparation, Chapter 4, section 1.2).
  
  
The approach to identify DMRs can be summarized as follow:

- **a.** Creation of two random datasets that will be compared to real data.
- **b.** Definition of isolated CpG methylation state for both real and random datasets, using **methylKit** since this tool is suitable for this step.
- **c.** Selection of relevant CpGs, based on a pvalue threshold.
- **d.** Detection of DMRs with **get\_close\_loci()**, which is adapted to identify differentially methylated regions on the methylome capture we studied. Tests with various parameters were done, to identify conditions for which random and real dataset give distinct results (*i.e.* conditions for which real data results make sense and are not obtained by chance).

###### 14.7.a Creation of two random datasets¶

Two random datasets were generated and compared to real one, to define parameters of get\_close\_loci() function that were appropriated for DMRs detection.
  
  
To obtain these random datasets, methylation states from real samples, were randomly switched, **CpG position by CpG position**, between samples and replicates using our own R script (see below, Supp. Fig. S1E of the paper in preparation, Chapter 4, section 1.2). Each methylation state is thus reassigned to a given sample, but methylation state between distinct CpG positions are not shuffled. This randomization assigns biologically plausible methylation values, actually observed at a given cytosine position (this is not the case if randomization was global, i.e redistribution of CpG methylation state, regardless of the CpG position, because there would be a risk of assigning, for example, high DNA methylation rates to a cytosine position that is always detected as non-methylated, regardless the conditions, PBS or EtOH-treatment).
  
  
Randomization was done on filtered - *i.e.* CpG positions without low read coverage (number of reads < 10) or abnormal high read coverage (> 99.9th percentile) - but unormalized real dataset. Quality controls (**getMethylationStats** and **getCoverageStats** of these filtered but unormalized data were performed. Read coverage was normalized after the randomization, using median to calculate scaling factor.

###### 14.7.b. Definition of isolated CpG methylation state¶

**methylKit** was used to determine methylation state of each isolated CpGs found in all the samples.

In [ ]:

```
%%R

setwd("/Volumes/DDMerger/methylome/fusion/methylkit/capture")

# ------------------------------------------
# 1. Real data filtering
# ------------------------------------------

# To remove cytosines that are not enough or too much covered
Captfmincov10= filterByCoverage(Captnf, lo.count=10, hi.perc=99.9)

# /!\ No normalization of read coverage, before randomization /!\

# ------------------------------------------
# 2a.Quality control n°1 : getMethylationStats
# ------------------------------------------

#Default parameters - .png output
png(filename="methylation stat capt1M fus f mincov10.png")
getMethylationStats(Captfmincov10[[1]],plot=T,both.strands=F) 
dev.off()
  
png(filename="methylation stat capt2M fus f mincov10.png")
getMethylationStats(Captfmincov10[[2]],plot=T,both.strands=F) 
dev.off()
 
png(filename="methylation stat capt3M fus f mincov10.png")
getMethylationStats(Captfmincov10[[3]],plot=T,both.strands=F)
dev.off()
 
png(filename="methylation stat capt4M fus f mincov10.png")
getMethylationStats(Captfmincov10[[4]],plot=T,both.strands=F) 
dev.off()
 
png(filename="methylation stat capt5M fus f mincov10.png")
getMethylationStats(Captfmincov10[[5]],plot=T,both.strands=F) 
dev.off()
 
png(filename="methylation stat capt6M fus f mincov10.png")
getMethylationStats(Captfmincov10[[6]],plot=T,both.strands=F) 
dev.off()
 
#for nclass=10 - .png output
png(filename="methylation stat capt1M fus f mincov10nclass10.png")
getMethylationStats(Captfmincov10[[1]],plot=T,both.strands=F,nclass=10)
dev.off()
 
png(filename="methylation stat capt2M fus f mincov10nclass10.png")
getMethylationStats(Captfmincov10[[2]],plot=T,both.strands=F,nclass=10)
dev.off()
 
png(filename="methylation stat capt3M fus f mincov10nclass10.png")
getMethylationStats(Captfmincov10[[3]],plot=T,both.strands=F,nclass=10)
dev.off()
 
png(filename="methylation stat capt4M fus f mincov10nclass10.png")
getMethylationStats(Captfmincov10[[4]],plot=T,both.strands=F,nclass=10)
dev.off()
 
png(filename="methylation stat capt5M fus f mincov10nclass10.png")
getMethylationStats(Captfmincov10[[5]],plot=T,both.strands=F,nclass=10)
dev.off()
 
png(filename="methylation stat capt6M fus f mincov10nclass10.png")
getMethylationStats(Captfmincov10[[6]],plot=T,both.strands=F,nclass=10)
dev.off()
 

#Default parameters - .emf ouput
library(devEMF)

emf(file="methylation stat capt1M fus f mincov10.emf")
getMethylationStats(Captfmincov10[[1]],plot=T,both.strands=F) 
dev.off()

emf(file="methylation stat capt2M fus f mincov10.emf")
getMethylationStats(Captfmincov10[[2]],plot=T,both.strands=F) 
dev.off()

emf(file="methylation stat capt3M fus f mincov10.emf")
getMethylationStats(Captfmincov10[[3]],plot=T,both.strands=F) 
dev.off()

emf(file="methylation stat capt4M fus f mincov10.emf")
getMethylationStats(Captfmincov10[[4]],plot=T,both.strands=F) 
dev.off()

emf(file="methylation stat capt5M fus f mincov10.emf")
getMethylationStats(Captfmincov10[[5]],plot=T,both.strands=F) 
dev.off()

emf(file="methylation stat capt6M fus f mincov10.emf")
getMethylationStats(Captfmincov10[[6]],plot=T,both.strands=F) 
dev.off()

#for nclass=10 - .emf output 
emf(file="methylation stat capt1M fus f mincov10 nclass10.emf")
getMethylationStats(Captfmincov10[[1]],plot=T,both.strands=F,nclass=10)
dev.off()

emf(file="methylation stat capt2M fus f mincov10 nclass10.emf")
getMethylationStats(Captfmincov10[[2]],plot=T,both.strands=F,nclass=10)
dev.off()

emf(file="methylation stat capt3M fus f mincov10 nclass10.emf")
getMethylationStats(Captfmincov10[[3]],plot=T,both.strands=F,nclass=10)
dev.off()

emf(file="methylation stat capt4M fus f mincov10 nclass10.emf")
getMethylationStats(Captfmincov10[[4]],plot=T,both.strands=F,nclass=10)
dev.off()

emf(file="methylation stat capt5M fus f mincov10 nclass10.emf")
getMethylationStats(Captfmincov10[[5]],plot=T,both.strands=F,nclass=10)
dev.off()

emf(file="methylation stat capt6M fus f mincov10 nclass10.emf")
getMethylationStats(Captfmincov10[[6]],plot=T,both.strands=F,nclass=10)
dev.off()


#Tables
getMethylationStats(Captfmincov10[[1]],plot=F,both.strands=F) 
getMethylationStats(Captfmincov10[[2]],plot=F,both.strands=F) 
getMethylationStats(Captfmincov10[[3]],plot=F,both.strands=F) 
getMethylationStats(Captfmincov10[[4]],plot=F,both.strands=F) 
getMethylationStats(Captfmincov10[[5]],plot=F,both.strands=F) 
getMethylationStats(Captfmincov10[[6]],plot=F,both.strands=F)


# ------------------------------------------
# 2b.Quality control n°2 : getCoverageStats
# ------------------------------------------

#Default parameters - .png output
png(filename="coverage Stat capt1M fus f mincov10.png")
getCoverageStats(Captfmincov10[[1]],plot=T,both.strands=F) 
dev.off()
 
png(filename="coverage Stat capt2M fus f mincov10.png")
getCoverageStats(Captfmincov10[[2]],plot=T,both.strands=F)
dev.off()
 
png(filename="coverage Stat capt3M fus f mincov10.png")
getCoverageStats(Captfmincov10[[3]],plot=T,both.strands=F)
dev.off()
 
png(filename="coverage Stat capt4M fus f mincov10.png")
getCoverageStats(Captfmincov10[[4]],plot=T,both.strands=F)
dev.off()
 
png(filename="coverage Stat capt5M fus f mincov10.png")
getCoverageStats(Captfmincov10[[5]],plot=T,both.strands=F)
dev.off()
 
png(filename="coverage Stat capt6M fus f mincov10.png")
getCoverageStats(Captfmincov10[[6]],plot=T,both.strands=F)
dev.off()
 
#Default parameters - .emf output: 
emf (file="coverage Stat capt1M fus f mincov10.emf")
getCoverageStats(Captfmincov10[[1]],plot=T,both.strands=F) 
dev.off()

emf(file="coverage Stat capt2M fus f mincov10.emf")
getCoverageStats(Captfmincov10[[2]],plot=T,both.strands=F)
dev.off()

emf(file="coverage Stat capt3M fus f mincov10.emf")
getCoverageStats(Captfmincov10[[3]],plot=T,both.strands=F)
dev.off()

emf(file="coverage Stat capt4M fus f mincov10.emf")
getCoverageStats(Captfmincov10[[4]],plot=T,both.strands=F)
dev.off()

emf(file="coverage Stat capt5M fus f mincov10.emf")
getCoverageStats(Captfmincov10[[5]],plot=T,both.strands=F)
dev.off()

emf(file="coverage Stat capt6M fus f mincov10.emf")
getCoverageStats(Captfmincov10[[6]],plot=T,both.strands=F)
dev.off()
```

###### Conclusions (globally similar than filtered and normalized data):¶

- For **getMethylationStats**, histogram profiles of the captured data are similar than those expected for genome-wide dataset (*i.e.* high number of CpG that are never methylated, and high number of CpG that are always methylated). In our captured data, there are more CpG that are never methylated than CpG that are always methylated. This imbalance can be due to the choice of specific genomic regions in the capture. Indeed, capture is composed of promoters that are majorly unmethylated in mouse genome.
- For **getCoverageStats**, no peak is observed on the right end of each histogram, meaining that deduplication step works properly (there are no PCR duplicates, thus no biases for the statistical analysis at this level).

In [ ]:

```
%%R

# ------------------------------------------
# 3.Obtention of a methylBase object = CpG methylation state count table
# ------------------------------------------

# methylBase object containing only cytosines that are covered in aLL samples:
meth_capt_strict_BME=methylKit::unite(Captfmincov10, destrand=FALSE) 
meth_capt_strict_BME
# 1 259 111  rows = number of CpG that are covered (at least 10 times) in aLL samples 

setwd("/Volumes/DDMezger/methylome/fusion/methylkit/capture/test_randomisation/randomisation_en_ligne")

write.table(meth_capt_strict_BME, file = "pool_strict_captBME10_99.9_non_normalise.txt", 
            quote = FALSE, sep = "\t")


#-------------------------------------
# 4a. Creation of the first random dataset - row-by-row randomization
#-------------------------------------

#=========
# Part 1: creation of random dataset (unormalized data)
#=========

# We need a methylBase object as input (i.e. tables with coverage, nb of methylated/unmethylated reads for each CpG)
pool.init = read.table("pool_strict_captBME10_99.9_non_normalise.txt", header = TRUE, sep = "\t")

head(pool.init)

#To select only numerical values (no coordinates)
pool.init.num = pool.init[,5:22]
head(pool.init.num)

#Function to row-by-row randomize the data
# --> 6 samples (3 versus 3 samples)
# --> 3 concatenated values per sample that must be kept together during the shuffling.
# These 3 values correspond to coverage and nb of methylated/unmethylated reads for each CpG
randomisation.en.ligne = function(matligne){
  
  nb.ech = 6
  nb.col.grp = 3
  
  #To create variables called val.ech1, val.ech2...
  for (i in 1:nb.ech){
    assign(x = paste("val.ech", i, sep = ""), value = NULL)
  }
  
  #To combine columns that must be kept together during the shuffling
  # (i.e. 3 values = coverage, nb of meth/unmethylated CpG)
  for (i in 1:nrow(matligne)){
    val.ech1 = c(val.ech1, paste(matligne[i,1], matligne[i,2], matligne[i,nb.col.grp], sep =";"))
    val.ech2 = c(val.ech2, paste(matligne[i,nb.col.grp+1], matligne[i,nb.col.grp+2], matligne[i,nb.col.grp*2], sep =";"))
    val.ech3 = c(val.ech3, paste(matligne[i,nb.col.grp*2+1], matligne[i,nb.col.grp*2+2], matligne[i,nb.col.grp*3], sep =";"))
    val.ech4 = c(val.ech4, paste(matligne[i,nb.col.grp*3+1], matligne[i,nb.col.grp*3+2], matligne[i,nb.col.grp*4], sep =";"))
    val.ech5 = c(val.ech5, paste(matligne[i,nb.col.grp*4+1], matligne[i,nb.col.grp*4+2], matligne[i,nb.col.grp*5], sep =";"))
    val.ech6 = c(val.ech6, paste(matligne[i,nb.col.grp*5+1], matligne[i,nb.col.grp*5+2], matligne[i,nb.col.grp*6], sep =";"))
  }
  
  # To check the results
  print("val.ech1")
  print(head(val.ech1))
  print("val.ech2")
  print(head(val.ech2))
  print("val.ech3")
  print(head(val.ech3))
  print("val.ech4")
  print(head(val.ech4))
  print("val.ech5")
  print(head(val.ech5))
  print("val.ech6")
  print(head(val.ech6))
  
  # To combine the concatenated values into a global table
  concat = cbind(val.ech1,val.ech2, val.ech3, val.ech4, val.ech5, val.ech6)
  
  # Row-by-row randomization
  valeur.random = NULL
  
  for (i in 1:nrow(matligne)){
    valeur.random = c(valeur.random, sample(concat[i,], replace = FALSE))
  }
  
  head(valeur.random)
  length(valeur.random)
  
  tab.random.LbyL.concat = as.data.frame(matrix(valeur.random, 
                          ncol=ncol(concat), nrow=nrow(pool.init.num), 
                          byrow = TRUE))
  
  library(data.table)
  
  # To deconcatenate the 3 values that were previously concatenated,
      # to separate coverage, nb of methylated and unmethylated reads for a given CpG position.
  # nb : need to associate table containing concatenated values and table containing decontatenated values, otherwise, it doesn't work.   
  tab.random.LbyL.deconcat = cbind(tab.random.LbyL.concat, tstrsplit(tab.random.LbyL.concat$V1, ";"), tstrsplit(tab.random.LbyL.concat$V2, ";"), tstrsplit(tab.random.LbyL.concat$V3, ";"), 
                                   tstrsplit(tab.random.LbyL.concat$V4, ";"), tstrsplit(tab.random.LbyL.concat$V5, ";"), tstrsplit(tab.random.LbyL.concat$V6, ";"), deparse.level = 0)
  # deparse.level=0 argument constructs no labels, to prevents from having columns with extended name.
  
  #To name the columns (V1 to V24 columns).
  for (i in 1:24){
    colnames(tab.random.LbyL.deconcat)[i] = paste("V",i, sep="")
  }
  
  melange.decon.light = tab.random.LbyL.deconcat[,7:24]
  
  return(melange.decon.light)
  
}


# To test the function: 
randomisation.en.ligne(pool.init.num[1:10,])
pool.init.num[1:10,]

# To use the function
# Since pool.init.num is a huge file, the process was runned from split file, otherwise it requires too much processing power and time.
sample1_100000 = randomisation.en.ligne(pool.init.num[1:100000,])
write.table(sample1_100000, file = "sample1_100000.txt", quote = FALSE, sep = "\t")

sample100001_300000 = randomisation.en.ligne(pool.init.num[100001:300000,])
write.table(sample100001_300000, file = "sample100001_300000.txt", quote = FALSE, sep = "\t")

sample300001_500000 = randomisation.en.ligne(pool.init.num[300001:500000,])
write.table(sample300001_500000, file = "sample300001_500000.txt", quote = FALSE, sep = "\t")


# To automatate the process:
listing = list(500001:600000, 600001:700000, 700001:800000, 800001:900000, 900001:1000000,1000001:1100000, 1100001:1259111)
for (j in 1:3){ # mistake: I wrote 1:3 but I should have written 1:7, thus only three first elements of the list were taken into account.
ech = randomisation.en.ligne(pool.init.num[listing[[j]],])
write.table(ech, file = paste("sample",j,".txt", sep = ""), sep = "\t", quote = FALSE)
}


# Due to the mistake I mentioned before, I runned again the command for missing rows
# (I renamed files from the first listing to avoid overwritting their contents)
listing = list(800001:900000, 900001:1000000,1000001:1100000, 1100001:1259111)
for (j in 1:4){
ech = randomisation.en.ligne(pool.init.num[listing[[j]],])
write.table(ech, file = paste("sample",j,".txt", sep = ""), sep = "\t", quote = FALSE)
}


# Then, I renamed samples: sample1 (from row n° 500,001 to 600,000) to sample7 (from row n°1,100,001 to 1,259,111).
# To check the results : 
sample1[1,]
pool.init.num[500001,]

sample2[1,]
pool.init.num[600001,]

sample3[1,]
pool.init.num[700001,]

sample4[1,]
pool.init.num[800001,]

sample5[1,]
pool.init.num[900001,]

sample6[1,]
pool.init.num[1000001,]

sample7[1,]
pool.init.num[1100001,]
# Randomization seems to have been done correctly.


# To combine rows into a global table
sample01 = as.data.frame(read.table(file = "sample1_100000.txt", sep = "\t"))
sample02 = as.data.frame(read.table(file = "sample100001_300000.txt", sep = "\t"))
sample03 =  as.data.frame(read.table(file = "sample300001_500000.txt", sep = "\t"))
data.random.LbL.num = rbind(sample01,sample02, sample03, sample1, sample2, sample3, sample4, sample5, sample6, sample7)

# To add coordinates
data.random.LbL = cbind(pool.init[,1:3], data.random.LbL.num)

# To check
dim(data.random.LbL.num)
dim(data.random.LbL)
head(data.random.LbL)
head(pool.init)

#To save the random dataset
write.table(data.random.LbL, "data.random_pool_strict_captBME10_99.9_non.norm.txt",
            sep="\t", quote = FALSE, na="NA", dec = ".", row.names=FALSE, col.names=TRUE)


#=========
# Part 2: load the data on methylKit to normalize it
#=========

randomisation.non.norm = read.table(file = "data.random_pool_strict_captBME10_99.9_non.norm.txt", 
                                    header = TRUE, sep = "\t")

head(randomisation.non.norm)
dim(randomisation.non.norm)


# Data formatting to create methylkit inputs
# To separate data into different samples (1M_random, 2M_random, ... ,6M_random)
random_1M.non.norm = randomisation.non.norm[,1:6]
random_2M.non.norm = randomisation.non.norm[,c(1:3,7:9)]
random_3M.non.norm = randomisation.non.norm[,c(1:3,10:12)]
random_4M.non.norm = randomisation.non.norm[,c(1:3,13:15)]
random_5M.non.norm = randomisation.non.norm[,c(1:3,16:18)]
random_6M.non.norm = randomisation.non.norm[,c(1:3,19:21)]

head(random_1M.non.norm)
head(random_2M.non.norm)
head(random_3M.non.norm)
head(random_4M.non.norm)
head(random_5M.non.norm)
head(random_6M.non.norm)

# Creation of columns that are necessary for methylKit input
for(i in 1:6){
  name <- paste("random_", i,"M.non.norm", sep="")
  tmp <- get(name)
  colnames(tmp) <- c("chr", "start","end", "cov", "nbC", "nbT")
  tmp$freqC <- with(tmp, round((nbC/cov)*100, digits = 3))
  tmp$freqT <- with(tmp, round((nbT/cov)*100, digits = 3))
  tmp$strand <- with(tmp, paste("R"))
  tmp$chrBase <- with(tmp, paste(tmp$chr,tmp$start, sep ="."))
  print(name)
  print(head(tmp))
  assign(name, tmp)
}

# Extraction of columns of interest and saving in .txt files
for(i in 1:6){
  name <- paste("random_", i,"M.non.norm", sep="")
  name.output <- paste0("methylkit_random_", i, "M.non.norm")
  tmp <- get(name)
  assign(name.output, tmp[, c("chrBase","chr","start","strand","cov","freqC", "freqT")])
  print(name.output)
  tmp.output <- get(name.output)
  print(head(tmp.output))
  write.table(tmp.output, paste0("random_", i,"M_strict_captBME10_99.9_non.norm.txt"), 
              quote = FALSE, col.names= TRUE, sep = "\t")
}


# To import data on methylKit 
library(methylKit)

# To load sample files
M.list.captBME.Rand.non.norm=list("random_1M_strict_captBME10_99.9_non.norm.txt", 
                                  "random_2M_strict_captBME10_99.9_non.norm.txt",
                                  "random_3M_strict_captBME10_99.9_non.norm.txt", 
                                  "random_4M_strict_captBME10_99.9_non.norm.txt", 
                                  "random_5M_strict_captBME10_99.9_non.norm.txt", 
                                  "random_6M_strict_captBME10_99.9_non.norm.txt")

M_cov10_captBME.Rand.non.norm = read(M.list.captBME.Rand.non.norm, 
                                     sample.id=list("1M","2M","3M","4M","5M","6M"), 
                                     assembly="mm9", header = TRUE, context="CpG", 
                                     resolution = "base", treatment=c(1,1,1,0,0,0))

M_cov10_captBME.Rand.non.norm

# Normalization (data are already filtered)
M_cov10_captBME.Rand.norm = normalizeCoverage(M_cov10_captBME.Rand.non.norm, method="median")
M_cov10_captBME.Rand.norm

# To combine normalized data into a unique table 
# strict analysis: creation of a methylBase object with cytosines that are covered in aLL samples
meth_capt_strict_BME.Rand.norm = methylKit::unite(M_cov10_captBME.Rand.norm, destrand=FALSE)
meth_capt_strict_BME.Rand.norm

# To check: at this step, we expect to obtain same results than in "randomisation.non.norm" (= randomized file)
head(randomisation.non.norm)


#=========
# Part 3: Acquisition of methylation informations
#=========

DMC_capt_strict_BME.Rand.norm =calculateDiffMeth(meth_capt_strict_BME.Rand.norm)
DMC_capt_strict_BME.Rand.norm

write.table(DMC_capt_strict_BME.Rand.norm, 
            file ="pval_random.norm_apres_pool_strict_captBME10_99.9_non.norm.txt", 
            sep="\t", quote = FALSE, na="NA", dec = ".", row.names=FALSE, col.names=TRUE)
```

In [ ]:

```
%%R

#-------------------------------------
# 4b. Creation of a second random dataset - row-by-row randomization
#-------------------------------------

#=========
# Part 1: creation of random dataset (unormalized data)
#=========

# A methylBase object is needed as input (i.e. tables with coverage, nb of methylated/unmethylated reads for each CpG)
# File called pool_strict_captBME10_99.9_non_normalise.txt which has been already generated, was used as a input

library(methylKit)
library(data.table)

setwd("/Volumes/DDMezger/methylome/fusion/methylkit/capture/test_randomisation/randomisation_en_ligne")

pool.init = read.table("pool_strict_captBME10_99.9_non_normalise.txt", header = TRUE, sep = "\t")
head(pool.init)

#To select only numerical values (no coordinates)
pool.init.num = pool.init[,5:22]
head(pool.init.num)

#Function to row-by-row randomize the data
# --> 6 samples (3 versus 3 samples)
# --> 3 concatenated values per sample that must be kept together during the shuffling.
# These 3 values correspond to coverage and nb of methylated/unmethylated reads for each CpG
randomisation.en.ligne = function(matligne){
  
  nb.ech = 6
  nb.col.grp = 3
  
  # To create variables called val.ech1, val.ech2...
  for (i in 1:nb.ech){
    assign(x = paste("val.ech", i, sep = ""), value = NULL)
  }
  
  #To combine columns that must be kept together during the shuffling
  # (i.e. 3 values = coverage, nb of meth/unmethylated CpG)
  for (i in 1:nrow(matligne)){
    val.ech1 = c(val.ech1, paste(matligne[i,1], matligne[i,2], matligne[i,nb.col.grp], sep =";"))
    val.ech2 = c(val.ech2, paste(matligne[i,nb.col.grp+1], matligne[i,nb.col.grp+2], matligne[i,nb.col.grp*2], sep =";"))
    val.ech3 = c(val.ech3, paste(matligne[i,nb.col.grp*2+1], matligne[i,nb.col.grp*2+2], matligne[i,nb.col.grp*3], sep =";"))
    val.ech4 = c(val.ech4, paste(matligne[i,nb.col.grp*3+1], matligne[i,nb.col.grp*3+2], matligne[i,nb.col.grp*4], sep =";"))
    val.ech5 = c(val.ech5, paste(matligne[i,nb.col.grp*4+1], matligne[i,nb.col.grp*4+2], matligne[i,nb.col.grp*5], sep =";"))
    val.ech6 = c(val.ech6, paste(matligne[i,nb.col.grp*5+1], matligne[i,nb.col.grp*5+2], matligne[i,nb.col.grp*6], sep =";"))
  }
  
  # To check the results
  print("val.ech1")
  print(head(val.ech1))
  print("val.ech2")
  print(head(val.ech2))
  print("val.ech3")
  print(head(val.ech3))
  print("val.ech4")
  print(head(val.ech4))
  print("val.ech5")
  print(head(val.ech5))
  print("val.ech6")
  print(head(val.ech6))
  
  # To combine the concatenated values into a global table
  concat = cbind(val.ech1,val.ech2, val.ech3, val.ech4, val.ech5, val.ech6)
  
  # Row-by-row randomization
  valeur.random = NULL
  
  for (i in 1:nrow(matligne)){
    valeur.random = c(valeur.random, sample(concat[i,], replace = FALSE))
  }
  
  print(head(valeur.random))
  print(length(valeur.random))
  
  tab.random.LbyL.concat = as.data.frame(matrix(valeur.random, 
                                        ncol=ncol(concat), nrow=nrow(matligne), byrow = TRUE))
  
  # To deconcatenate the 3 values that were previously concatenated,
      # to separate coverage, nb of methylated and unmethylated reads for a given CpG position.
  # nb : need to associate table containing concatenated values and table containing decontatenated values, otherwise, it doesn't work.
  tab.random.LbyL.deconcat = cbind(tab.random.LbyL.concat, 
                                   tstrsplit(tab.random.LbyL.concat$V1, ";"), 
                                   tstrsplit(tab.random.LbyL.concat$V2, ";"), 
                                   tstrsplit(tab.random.LbyL.concat$V3, ";"), 
                                   tstrsplit(tab.random.LbyL.concat$V4, ";"), 
                                   tstrsplit(tab.random.LbyL.concat$V5, ";"), 
                                   tstrsplit(tab.random.LbyL.concat$V6, ";"), 
                                   deparse.level = 0)
  # deparse.level=0 argument constructs no labels, to prevents from having columns with extended name.
    
  #To name the columns (V1 to V24 columns).
  for (i in 1:24){
    colnames(tab.random.LbyL.deconcat)[i] = paste("V",i, sep="")
  }
  
  melange.decon.light = tab.random.LbyL.deconcat[,7:24]
  
  return(melange.decon.light)
  
}


# To test the function:
randomisation.en.ligne(pool.init.num[1:10,])

pool.init.num[1:10,]

# To use the function
# Since pool.init.num is a huge file, the process was runned from split file, otherwise it requires too much processing power and time.
# To automatate the process:
listing = list(1:100000,100001:200000,200001:300000,300001:400000,
               400001:500000,500001:600000, 600001:700000, 700001:800000,
               800001:900000, 900001:1000000,1000001:1100000, 1100001:1259111)

for (j in 1:12){
  ech = randomisation.en.ligne(pool.init.num[listing[[j]],])
  write.table(ech, file = paste("sample",j,".txt", sep = ""), sep = "\t", quote = FALSE)
  assign(x = paste("sample", j, sep = ""), value = ech)
}

# To combine rows into a global table
data.random.LbL.num = rbind(sample1,sample2, sample3, sample4, 
                            sample5, sample6, sample7, sample8, 
                            sample9, sample10, sample11, sample12)

# To add coordinates
data.random.LbL.comp = cbind(pool.init[,1:3], data.random.LbL.num[,])

#To save the random dataset
write.table(data.random.LbL.comp, "data.random_pool_strict_captBME10_99.9_non.norm.txt", sep="\t", quote = FALSE, na="NA", dec = ".", row.names=FALSE, col.names=TRUE)


#=========
# Part 2: load the data on methylKit to normalize it
#=========

randomisation.non.norm = read.table(file = "data.random_pool_strict_captBME10_99.9_non.norm.txt", 
                                    header = TRUE, sep = "\t")

# To check
head(randomisation.non.norm)
dim(randomisation.non.norm)

# Data formatting to create methylkit inputs
# To separate data into different samples (1M_random, 2M_random, ... ,6M_random)
random_1M.non.norm = randomisation.non.norm[,1:6]
random_2M.non.norm = randomisation.non.norm[,c(1:3,7:9)]
random_3M.non.norm = randomisation.non.norm[,c(1:3,10:12)]
random_4M.non.norm = randomisation.non.norm[,c(1:3,13:15)]
random_5M.non.norm = randomisation.non.norm[,c(1:3,16:18)]
random_6M.non.norm = randomisation.non.norm[,c(1:3,19:21)]

head(random_1M.non.norm)
head(random_2M.non.norm)
head(random_3M.non.norm)
head(random_4M.non.norm)
head(random_5M.non.norm)
head(random_6M.non.norm)

# Creation of columns that are necessary for methylKit input
for(i in 1:6){
  name <- paste("random_", i,"M.non.norm", sep="")
  tmp <- get(name)
  colnames(tmp) <- c("chr", "start","end", "cov", "nbC", "nbT")
  tmp$freqC <- with(tmp, round((nbC/cov)*100, digits = 3))
  tmp$freqT <- with(tmp, round((nbT/cov)*100, digits = 3))
  tmp$strand <- with(tmp, paste("R"))
  tmp$chrBase <- with(tmp, paste(tmp$chr,tmp$start, sep ="."))
  print(name)
  print(head(tmp))
  assign(name, tmp)
}


# Extraction of columns of interest and saving in .txt files
for(i in 1:6){
  name <- paste("random_", i,"M.non.norm", sep="")
  name.output <- paste0("methylkit_random_", i, "M.non.norm")
  tmp <- get(name)
  assign(name.output, tmp[, c("chrBase","chr","start","strand","cov","freqC", "freqT")])
  print(name.output)
  tmp.output <- get(name.output)
  print(head(tmp.output))
  write.table(tmp.output, paste0("random_", i,"M_strict_captBME10_99.9_non.norm.txt"), quote = FALSE, col.names= TRUE, sep = "\t")
}

# To import data on methylKit 
library(methylKit)

# To load sample files 
M.list.captBME.Rand.non.norm=list("random_1M_strict_captBME10_99.9_non.norm.txt", 
                                  "random_2M_strict_captBME10_99.9_non.norm.txt",
                                  "random_3M_strict_captBME10_99.9_non.norm.txt", 
                                  "random_4M_strict_captBME10_99.9_non.norm.txt", 
                                  "random_5M_strict_captBME10_99.9_non.norm.txt", 
                                  "random_6M_strict_captBME10_99.9_non.norm.txt")

M_cov10_captBME.Rand.non.norm = read(M.list.captBME.Rand.non.norm, 
                                     sample.id=list("1M","2M","3M","4M","5M","6M"), 
                                     assembly="mm9", header = TRUE, context="CpG", 
                                     resolution = "base", treatment=c(1,1,1,0,0,0))

M_cov10_captBME.Rand.non.norm

# Normalization (data are already filtered)
M_cov10_captBME.Rand.norm = normalizeCoverage(M_cov10_captBME.Rand.non.norm, method="median")
M_cov10_captBME.Rand.norm

# To combine normalized data into a unique table 
# strict analysis: creation of a methylBase object with cytosines that are covered in aLL samples
meth_capt_strict_BME.Rand.norm = methylKit::unite(M_cov10_captBME.Rand.norm, destrand=FALSE)

meth_capt_strict_BME.Rand.norm

# To check: at this step, we expect to obtain same results than in "randomisation.non.norm" (= randomized file)
head(randomisation.non.norm)


#=========
# Part 3: Acquisition of methylation informations
#=========

DMC_capt_strict_BME.Rand.norm =calculateDiffMeth(meth_capt_strict_BME.Rand.norm)
DMC_capt_strict_BME.Rand.norm

write.table(DMC_capt_strict_BME.Rand.norm, 
            file ="pval_random.norm_apres_pool_strict_captBME10_99.9_non.norm.txt", 
            sep="\t", quote = FALSE, na="NA", dec = ".", row.names=FALSE, col.names=TRUE)
```

###### 14.7.c. Selection of relevant CpGs: definition of a pvalue threshold¶

To define DMRs with get\_close\_loci(), selection of relevant CpGs is essential. Indeed, DMR detection could be compromised if CpGs with no reliable methylation state are kept (see below, Supp. Fig. S1D of the paper in preparation, Chapter 4, section 1.2). Thus a **pre-selection** of relevant CpGs was done, according to their pvalues.

###### 14.7.d. Modulation of parameters for DMRs detection¶

To define appropriate parameters for DMR detection, different parameters were modulated and number of DMRs found in the two datasets (real *versus* random) was compared (Fig. S1B, Supp. Table 2 of the paper in preparation, Chapter 4, section 1.2). To be more precise, two random datasets were compared to the real data, by testing different combinations of DMR detection parameters of get\_close\_loci() function (pvalue threshold, minimum number of CpGs in the region), and by fixing some parameters (distance\_C\_to\_C = 100, distance\_threshold=500, max\_region\_size = 2000). Depending on the pvalue threshold chosen, more or less CpGs are selected. In each case, more CpGs are selected in the real dataset than in the random one for a given p-value threshold. To be able to compare the number of DMRs obtained in the real dataset and in the random one, the same number of CpGs must be considered at the outset. Thus, by random selection of relevant CpGs among the real dataset, the number of CpGs in the real dataset was adjusted to the number of CpGs found in the random dataset. Similar results were obtained by using the two random datasets (Supp. Table 2 of the paper in preparation, Chapter 4, section 1.2).

In [ ]:

```
%%R

# -------------------------------
# 1.Comparison between real and random dataset (random dataset n°1)
# -------------------------------

setwd("/Volumes/DDMezger/methylome/fusion/methylkit/capture/test_randomisation/randomisation_en_ligne/premier_random")

# To load random dataset
data.random.norm = read.table("pval_random.norm_apres_pool_strict_captBME10_99.9_non.norm.txt", 
                              sep = "\t", header = TRUE)

# To load real dataset
data.real = read.table("pval_real_pool_strict_captBME10_99.9.txt", 
                       sep = "\t", header = TRUE)

head(data.random.norm)
head(data.real)

# To combine real and random dataset into a unique dataframe
pool.norm=cbind(data.real, data.random.norm[,5:7])

colnames(pool.norm) = c("chr","start","end","strand","pvalue.real",
                        "qvalue.real","meth.diff.real", "pvalue.random", 
                        "qvalue.random", "meth.diff.random")
head(pool.norm)


# Function get_close_loci to define DMRs
get_close_loci_AD3 <- function(DMCs,number_of_lines=2,
                               distance_threshold,distance_C_to_C, 
                               max_region_size=100,
                               consistent_direction=TRUE,
                               deactivate_consistency_check=FALSE) {
  result <- matrix(0,0,ncol(DMCs))
  result2 <- matrix(0,0,ncol(DMCs))
  second_last <- nrow(DMCs)-(number_of_lines-1)
  i <- 1
  while (i<=second_last) {
    first_chr <- DMCs[i,1]
    second_chr <- DMCs[i+(number_of_lines-1),1]
    if (first_chr==second_chr) {
      difference <- DMCs[i+(number_of_lines-1),2]-DMCs[i,2]
      if (difference<=distance_threshold) {
        subsetted <- DMCs[i:(i+number_of_lines-1),7]
        if (sum(subsetted>=0)==number_of_lines || sum(subsetted<0)==number_of_lines || deactivate_consistency_check) {
          
          # expand region if possible to remove redundancy
          conditions_met <- TRUE
          last_line <- i+number_of_lines-1
          while ((last_line<nrow(DMCs))&&(conditions_met)) {
            new_chr <- DMCs[last_line+1,1]
            if (first_chr != new_chr) {
              conditions_met <- FALSE
            }
            new_size <- 1+DMCs[last_line+1,2]-DMCs[i,2]
            if (new_size>max_region_size) {
              conditions_met <- FALSE
            }
            subsetted <- DMCs[i:(last_line+1),7]
            if (sum(subsetted>0)!=(last_line-i+2) && sum(subsetted<0)!=(last_line-i+2)) {
              conditions_met <- FALSE
            }
            difference_bis <-  DMCs[last_line+1,2] - DMCs[last_line,2]
            if (difference_bis > distance_C_to_C) {
              conditions_met <- FALSE
            }
            if (conditions_met) {
              last_line <- last_line+1
            }
            
          }
          
          final_set <- DMCs[i:last_line,]
          DMCs[i,3] <- DMCs[last_line,2]
          DMCs[i,4] <- last_line-i+1
          DMCs[i,6] <- median(final_set[,6])
          DMCs[i,7] <- median(final_set[,7])
          result <- rbind(DMCs[i,],result)
          i <- last_line
        } else {
          if (!consistent_direction) {
            result2 <- rbind(DMCs[i:(i+(number_of_lines-1)),],result2)  
          }
        }
      }
    }
    i <- i+1
  }
  if (nrow(result2)>0) {
    return(result2)
  } else {
    return(result)
  }
}


#-------------------------------------------------------------

# The following code was runned, by changing pvalue threshold (0.008 ; 0.01 ; 0.03 ; 0.05; 0.08; 0.1...): 
DMC_real_0.05 = data.real[data.real[,"pvalue"]<0.05,]
DMC_random_0.05 = data.random.norm[data.random.norm[,"pvalue"]<0.05,]

dim(DMC_real_0.05)
dim(DMC_random_0.05)

# To be able to compare DMRs numbers obtained in real and random datasets, it is necessary to start the analysis  
# with the same numbers of isolated CpG in the two compared datasets. Since we always observed a greater number of CpGs
# in real dataset than in random one, for a given pvalue threshold, we decided to randomly select isolated CpG of the real dataset, among CpGs that meet the pvalue threshold, to analyse same number of CpG in real and random dataset.
# To randomly select CpG in real dataset:
DMC_real_0.05_cut = DMC_real_0.05[sort(sample(1:nrow(DMC_real_0.05),nrow(DMC_random_0.05))),]
dim(DMC_real_0.05_cut)

# To replace "+" in strand column by "0" in order to avoid error message for future command lines execution
DMC_real_0.05[,4]= 0
DMC_real_0.05_cut[,4]= 0
DMC_random_0.05[,4]= 0

# To use get_close_loci function with real versus random dataset 
# whole real dataset
DMR3_real_0.05 = get_close_loci_AD3(DMC_real_0.05,
                                    number_of_lines=3,
                                    distance_threshold=500,
                                    distance_C_to_C=100, 
                                    max_region_size=2000,
                                    consistent_direction=TRUE,
                                    deactivate_consistency_check=FALSE)  
dim(DMR3_real_0.05)

# "filtered" real dataset to have as many CpG as in the random dataset
DMR3_real_0.05_cut = get_close_loci_AD3(DMC_real_0.05_cut,
                                        number_of_lines=3,
                                        distance_threshold=500,
                                        distance_C_to_C=100, 
                                        max_region_size=2000,
                                        consistent_direction=TRUE,
                                        deactivate_consistency_check=FALSE)  
dim(DMR3_real_0.05_cut)

# random dataset
DMR3_random_0.05 = get_close_loci_AD3(DMC_random_0.05,
                                      number_of_lines=3,
                                      distance_threshold=500,
                                      distance_C_to_C=100, 
                                      max_region_size=2000,
                                      consistent_direction=TRUE,
                                      deactivate_consistency_check=FALSE)  
dim(DMR3_random_0.05)

# same process, with distinct parameters
DMR4_real_0.05 = get_close_loci_AD3(DMC_real_0.05,
                                    number_of_lines=4,
                                    distance_threshold=500,
                                    distance_C_to_C=100, 
                                    max_region_size=2000,
                                    consistent_direction=TRUE,
                                    deactivate_consistency_check=FALSE)  
dim(DMR4_real_0.05)

DMR4_real_0.05_cut = get_close_loci_AD3(DMC_real_0.05_cut,
                                        number_of_lines=4,
                                        distance_threshold=500,
                                        distance_C_to_C=100, 
                                        max_region_size=2000,
                                        consistent_direction=TRUE,
                                        deactivate_consistency_check=FALSE)  
dim(DMR4_real_0.05_cut)

DMR4_random_0.05 = get_close_loci_AD3(DMC_random_0.05,
                                      number_of_lines=4,
                                      distance_threshold=500,
                                      distance_C_to_C=100, 
                                      max_region_size=2000,
                                      consistent_direction=TRUE,
                                      deactivate_consistency_check=FALSE)  
dim(DMR4_random_0.05)

DMR5_real_0.05 = get_close_loci_AD3(DMC_real_0.05,
                                    number_of_lines=5,
                                    distance_threshold=500,
                                    distance_C_to_C=100, 
                                    max_region_size=2000,
                                    consistent_direction=TRUE,
                                    deactivate_consistency_check=FALSE)  
dim(DMR5_real_0.05)

DMR5_real_0.05_cut = get_close_loci_AD3(DMC_real_0.05_cut,
                                        number_of_lines=5,
                                        distance_threshold=500,
                                        distance_C_to_C=100, 
                                        max_region_size=2000,
                                        consistent_direction=TRUE,
                                        deactivate_consistency_check=FALSE)  
dim(DMR5_real_0.05_cut)

DMR5_random_0.05 = get_close_loci_AD3(DMC_random_0.05,
                                      number_of_lines=5,
                                      distance_threshold=500,
                                      distance_C_to_C=100, 
                                      max_region_size=2000,
                                      consistent_direction=TRUE,
                                      deactivate_consistency_check=FALSE)  
dim(DMR5_random_0.05)

                                                                 
                                                                 
# ----------------------------------------------------------                                                                
                                                                
# Then, observed DMRs can be filtered, based on percent of methylation differences between control and treated groups.
# For example, we can keep only DMR having a high differential of methylation between control and treated groups (>10% ou < à -10%)

DMR5_real_0.05_filtre = DMR5_real_0.05[DMR5_real_0.05[,"meth.diff"]< -10 | DMR5_real_0.05[,"meth.diff"]> 10, ]
dim(DMR5_real_0.05_filtre)

DMR5_real_0.05_filtre

# To save informations into files                                                             
write.table(DMR5_real_0.05, file = "DMR225_real_p0.05.txt", 
            sep = "\t", quote = FALSE, dec = ".", row.names=FALSE, col.names=TRUE)

write.table(DMR5_real_0.05_filtre, file = "DMR173_real_p0.05.txt", 
            sep = "\t", quote = FALSE, dec = ".", row.names=FALSE, col.names=TRUE)

                     

#====================================================
                                                                 
 

# -------------------------------
# 2.Comparison between real and random dataset (random dataset n°2)
# -------------------------------

setwd("E:/methylome/fusion/methylkit/capture/test_randomisation/randomisation_en_ligne/second_dataset")

# To load random dataset
data.random.norm = read.table("pval_random.norm_apres_pool_strict_captBME10_99.9_non.norm.txt", 
                              sep = "\t", header = TRUE)

setwd("E:/methylome/fusion/methylkit/capture/test_randomisation/randomisation_en_ligne/premier_random")

# To load real dataset
data.real = read.table("pval_real_pool_strict_captBME10_99.9.txt", 
                       sep = "\t", header = TRUE)

head(data.random.norm)
head(data.real)

dim(data.random.norm)
dim(data.real)

# To combine real and random dataset into a unique dataframe
pool.norm=cbind(data.real, data.random.norm[,5:7])

colnames(pool.norm) = c("chr","start","end","strand",
                        "pvalue.real","qvalue.real",
                        "meth.diff.real", "pvalue.random", 
                        "qvalue.random", "meth.diff.random")
head(pool.norm)


# Function get_close_loci to define DMRs
get_close_loci_AD3 <- function(DMCs,number_of_lines=2,
                               distance_threshold,
                               distance_C_to_C, 
                               max_region_size=100,
                               consistent_direction=TRUE,
                               deactivate_consistency_check=FALSE) {
  result <- matrix(0,0,ncol(DMCs))
  result2 <- matrix(0,0,ncol(DMCs))
  second_last <- nrow(DMCs)-(number_of_lines-1)
  i <- 1
  while (i<=second_last) {
    first_chr <- DMCs[i,1]
    second_chr <- DMCs[i+(number_of_lines-1),1]
    if (first_chr==second_chr) {
      difference <- DMCs[i+(number_of_lines-1),2]-DMCs[i,2]
      if (difference<=distance_threshold) {
        subsetted <- DMCs[i:(i+number_of_lines-1),7]
        if (sum(subsetted>=0)==number_of_lines || sum(subsetted<0)==number_of_lines || deactivate_consistency_check) {
          
          # expand region if possible to remove redundancy
          conditions_met <- TRUE
          last_line <- i+number_of_lines-1
          while ((last_line<nrow(DMCs))&&(conditions_met)) {
            new_chr <- DMCs[last_line+1,1]
            if (first_chr != new_chr) {
              conditions_met <- FALSE
            }
            new_size <- 1+DMCs[last_line+1,2]-DMCs[i,2]
            if (new_size>max_region_size) {
              conditions_met <- FALSE
            }
            subsetted <- DMCs[i:(last_line+1),7]
            if (sum(subsetted>0)!=(last_line-i+2) && sum(subsetted<0)!=(last_line-i+2)) {
              conditions_met <- FALSE
            }
            difference_bis <-  DMCs[last_line+1,2] - DMCs[last_line,2]
            if (difference_bis > distance_C_to_C) {
              conditions_met <- FALSE
            }
            if (conditions_met) {
              last_line <- last_line+1
            }
            
          }
          
          final_set <- DMCs[i:last_line,]
          DMCs[i,3] <- DMCs[last_line,2]
          DMCs[i,4] <- last_line-i+1
          DMCs[i,6] <- median(final_set[,6])
          DMCs[i,7] <- median(final_set[,7])
          result <- rbind(DMCs[i,],result)
          i <- last_line
        } else {
          if (!consistent_direction) {
            result2 <- rbind(DMCs[i:(i+(number_of_lines-1)),],result2)  
          }
        }
      }
    }
    i <- i+1
  }
  if (nrow(result2)>0) {
    return(result2)
  } else {
    return(result)
  }
}


#-------------------------------------------------------------

# The following code was runned, by changing pvalue threshold (0.05; 0.06 ; 0.07 ; 0.08): 
# For pvalue threshold = 0.05
DMC_real_0.05 = data.real[data.real[,"pvalue"]<0.05,]
DMC_random_0.05 = data.random.norm[data.random.norm[,"pvalue"]<0.05,]

dim(DMC_real_0.05)
dim(DMC_random_0.05)

# To be able to compare DMRs numbers obtained in real and random datasets, it is necessary to start the analysis  
# with the same numbers of isolated CpG in the two compared datasets. Since we always observed a greater number of CpGs
# in real dataset than in random one, for a given pvalue threshold, we decided to randomly select isolated CpG, 
# of the real dataset, among CpGs that meet the pvalue threshold, to analyse same number of CpG in real and random dataset.
# To randomly select CpG in real dataset:
DMC_real_0.05_cut = DMC_real_0.05[sort(sample(1:nrow(DMC_real_0.05),nrow(DMC_random_0.05))),]
dim(DMC_real_0.05_cut)


# To replace "+" in strand column by "0" in order to avoid error message for future command lines execution
DMC_real_0.05[,4]= 0
DMC_real_0.05_cut[,4]= 0
DMC_random_0.05[,4]= 0

# To use get_close_loci function with real versus random dataset 
# whole real dataset
DMR3_real_0.05 = get_close_loci_AD3(DMC_real_0.05,
                                    number_of_lines=3,
                                    distance_threshold=500,
                                    distance_C_to_C=100, 
                                    max_region_size=2000,
                                    consistent_direction=TRUE,
                                    deactivate_consistency_check=FALSE)  
dim(DMR3_real_0.05)

# "filtered" real dataset to have as many CpG as in the random dataset
DMR3_real_0.05_cut = get_close_loci_AD3(DMC_real_0.05_cut,
                                        number_of_lines=3,
                                        distance_threshold=500,
                                        distance_C_to_C=100, 
                                        max_region_size=2000,
                                        consistent_direction=TRUE,
                                        deactivate_consistency_check=FALSE)  
dim(DMR3_real_0.05_cut)

# random dataset
DMR3_random_0.05 = get_close_loci_AD3(DMC_random_0.05,
                                      number_of_lines=3,
                                      distance_threshold=500,
                                      distance_C_to_C=100, 
                                      max_region_size=2000,
                                      consistent_direction=TRUE,
                                      deactivate_consistency_check=FALSE)  
dim(DMR3_random_0.05)

# same process, with distinct parameters
DMR4_real_0.05 = get_close_loci_AD3(DMC_real_0.05,
                                    number_of_lines=4,
                                    distance_threshold=500,
                                    distance_C_to_C=100, 
                                    max_region_size=2000,
                                    consistent_direction=TRUE,
                                    deactivate_consistency_check=FALSE)  
dim(DMR4_real_0.05)

DMR4_real_0.05_cut = get_close_loci_AD3(DMC_real_0.05_cut,
                                        number_of_lines=4,
                                        distance_threshold=500,
                                        distance_C_to_C=100, 
                                        max_region_size=2000,
                                        consistent_direction=TRUE,
                                        deactivate_consistency_check=FALSE)  
dim(DMR4_real_0.05_cut)

DMR4_random_0.05 = get_close_loci_AD3(DMC_random_0.05,
                                      number_of_lines=4,
                                      distance_threshold=500,
                                      distance_C_to_C=100, 
                                      max_region_size=2000,
                                      consistent_direction=TRUE,
                                      deactivate_consistency_check=FALSE)  
dim(DMR4_random_0.05)

DMR5_real_0.05 = get_close_loci_AD3(DMC_real_0.05,
                                    number_of_lines=5,
                                    distance_threshold=500,
                                    distance_C_to_C=100, 
                                    max_region_size=2000,
                                    consistent_direction=TRUE,
                                    deactivate_consistency_check=FALSE)  
dim(DMR5_real_0.05)

DMR5_real_0.05_cut = get_close_loci_AD3(DMC_real_0.05_cut,
                                        number_of_lines=5,
                                        distance_threshold=500,
                                        distance_C_to_C=100, 
                                        max_region_size=2000,
                                        consistent_direction=TRUE,
                                        deactivate_consistency_check=FALSE)  
dim(DMR5_real_0.05_cut)

DMR5_random_0.05 = get_close_loci_AD3(DMC_random_0.05,
                                      number_of_lines=5,
                                      distance_threshold=500,
                                      distance_C_to_C=100, 
                                      max_region_size=2000,
                                      consistent_direction=TRUE,
                                      deactivate_consistency_check=FALSE)  
dim(DMR5_random_0.05)


# ----------------------------------------------------
# For pvalue threshold = 0.06
DMC_real_0.06 = data.real[data.real[,"pvalue"]<0.06,]
DMC_random_0.06 = data.random.norm[data.random.norm[,"pvalue"]<0.06,]

dim(DMC_real_0.06)
dim(DMC_random_0.06)

# To filter real dataset to have as many CpG as in the random dataset
DMC_real_0.06_cut = DMC_real_0.06[sort(sample(1:nrow(DMC_real_0.06),nrow(DMC_random_0.06))),]
dim(DMC_real_0.06_cut)

# To replace "+" in strand column by "0" in order to avoid error message for future command lines execution
DMC_real_0.06[,4]= 0
DMC_real_0.06_cut[,4]= 0
DMC_random_0.06[,4]= 0

# To use get_close_loci function with real versus random dataset 
# whole real dataset
DMR5_real_0.06 = get_close_loci_AD3(DMC_real_0.06,
                                    number_of_lines=5,
                                    distance_threshold=500,
                                    distance_C_to_C=100, 
                                    max_region_size=2000,
                                    consistent_direction=TRUE,
                                    deactivate_consistency_check=FALSE)  
dim(DMR5_real_0.06)

# "filtered" real dataset to have as many CpG as in the random dataset
DMR5_real_0.06_cut = get_close_loci_AD3(DMC_real_0.06_cut,
                                        number_of_lines=5,
                                        distance_threshold=500,
                                        distance_C_to_C=100, 
                                        max_region_size=2000,
                                        consistent_direction=TRUE,
                                        deactivate_consistency_check=FALSE)  
dim(DMR5_real_0.06_cut)

# random dataset
DMR5_random_0.06 = get_close_loci_AD3(DMC_random_0.06,
                                      number_of_lines=5,
                                      distance_threshold=500,
                                      distance_C_to_C=100, 
                                      max_region_size=2000,
                                      consistent_direction=TRUE,
                                      deactivate_consistency_check=FALSE)  
dim(DMR5_random_0.06)

write.table(DMR5_real_0.06, file = "DMR329_real_p0.06.txt", 
            sep = "\t", quote = FALSE, dec = ".", row.names=FALSE, col.names=TRUE)


# ----------------------------------------------------
# For pvalue threshold = 0.07
DMC_real_0.07 = data.real[data.real[,"pvalue"]<0.07,]
DMC_random_0.07 = data.random.norm[data.random.norm[,"pvalue"]<0.07,]

dim(DMC_real_0.07)
dim(DMC_random_0.07)

# To filter real dataset to have as many CpG as in the random dataset
DMC_real_0.07_cut = DMC_real_0.07[sort(sample(1:nrow(DMC_real_0.07),nrow(DMC_random_0.07))),]
dim(DMC_real_0.07_cut)

# To replace "+" in strand column by "0" in order to avoid error message for future command lines execution
DMC_real_0.07[,4]= 0
DMC_real_0.07_cut[,4]= 0
DMC_random_0.07[,4]= 0

# To use get_close_loci function with real versus random dataset 
# whole real dataset
DMR5_real_0.07 = get_close_loci_AD3(DMC_real_0.07,
                                    number_of_lines=5,
                                    distance_threshold=500,
                                    distance_C_to_C=100, 
                                    max_region_size=2000,
                                    consistent_direction=TRUE,
                                    deactivate_consistency_check=FALSE)  
dim(DMR5_real_0.07)

# "filtered" real dataset to have as many CpG as in the random dataset
DMR5_real_0.07_cut = get_close_loci_AD3(DMC_real_0.07_cut,
                                        number_of_lines=5,
                                        distance_threshold=500,
                                        distance_C_to_C=100, 
                                        max_region_size=2000,
                                        consistent_direction=TRUE,
                                        deactivate_consistency_check=FALSE)  
dim(DMR5_real_0.07_cut)

# random dataset
DMR5_random_0.07 = get_close_loci_AD3(DMC_random_0.07,
                                      number_of_lines=5,
                                      distance_threshold=500,
                                      distance_C_to_C=100, 
                                      max_region_size=2000,
                                      consistent_direction=TRUE,
                                      deactivate_consistency_check=FALSE)  
dim(DMR5_random_0.07)

write.table(DMR5_real_0.07, file = "DMR432_real_p0.07.txt", 
            sep = "\t", quote = FALSE, dec = ".", row.names=FALSE, col.names=TRUE)


# ----------------------------------------------------
# For pvalue threshold = 0.08
DMC_real_0.08 = data.real[data.real[,"pvalue"]<0.08,]
DMC_random_0.08 = data.random.norm[data.random.norm[,"pvalue"]<0.08,]

dim(DMC_real_0.08)
dim(DMC_random_0.08)

# To filter real dataset to have as many CpG as in the random dataset
DMC_real_0.08_cut = DMC_real_0.08[sort(sample(1:nrow(DMC_real_0.08),nrow(DMC_random_0.08))),]
dim(DMC_real_0.08_cut)

# To replace "+" in strand column by "0" in order to avoid error message for future command lines execution
DMC_real_0.08[,4]= 0
DMC_real_0.08_cut[,4]= 0
DMC_random_0.08[,4]= 0

# To use get_close_loci function with real versus random dataset 
# whole real dataset
DMR3_real_0.08 = get_close_loci_AD3(DMC_real_0.08,
                                    number_of_lines=3,
                                    distance_threshold=500,
                                    distance_C_to_C=100, 
                                    max_region_size=2000,
                                    consistent_direction=TRUE,
                                    deactivate_consistency_check=FALSE)  
dim(DMR3_real_0.08)

# "filtered" real dataset to have as many CpG as in the random dataset
DMR3_real_0.08_cut = get_close_loci_AD3(DMC_real_0.08_cut,
                                        number_of_lines=3,
                                        distance_threshold=500,
                                        distance_C_to_C=100, 
                                        max_region_size=2000,
                                        consistent_direction=TRUE,
                                        deactivate_consistency_check=FALSE)  
dim(DMR3_real_0.08_cut)

# random dataset
DMR3_random_0.08 = get_close_loci_AD3(DMC_random_0.08,
                                      number_of_lines=3,
                                      distance_threshold=500,
                                      distance_C_to_C=100, 
                                      max_region_size=2000,
                                      consistent_direction=TRUE,
                                      deactivate_consistency_check=FALSE)  
dim(DMR3_random_0.08)

# same process, with distinct parameters
DMR4_real_0.08 = get_close_loci_AD3(DMC_real_0.08,
                                    number_of_lines=4,
                                    distance_threshold=500,
                                    distance_C_to_C=100, 
                                    max_region_size=2000,
                                    consistent_direction=TRUE,
                                    deactivate_consistency_check=FALSE)  
dim(DMR4_real_0.08)

DMR4_real_0.08_cut = get_close_loci_AD3(DMC_real_0.08_cut,
                                        number_of_lines=4,
                                        distance_threshold=500,
                                        distance_C_to_C=100, 
                                        max_region_size=2000,
                                        consistent_direction=TRUE,
                                        deactivate_consistency_check=FALSE)  
dim(DMR4_real_0.08_cut)

DMR4_random_0.08 = get_close_loci_AD3(DMC_random_0.08,
                                      number_of_lines=4,
                                      distance_threshold=500,
                                      distance_C_to_C=100, 
                                      max_region_size=2000,
                                      consistent_direction=TRUE,
                                      deactivate_consistency_check=FALSE)  
dim(DMR4_random_0.08)

DMR5_real_0.08 = get_close_loci_AD3(DMC_real_0.08,
                                    number_of_lines=5,
                                    distance_threshold=500,
                                    distance_C_to_C=100, 
                                    max_region_size=2000,
                                    consistent_direction=TRUE,
                                    deactivate_consistency_check=FALSE)  
dim(DMR5_real_0.08)

DMR5_real_0.08_cut = get_close_loci_AD3(DMC_real_0.08_cut,
                                        number_of_lines=5,
                                        distance_threshold=500,
                                        distance_C_to_C=100, 
                                        max_region_size=2000,
                                        consistent_direction=TRUE,
                                        deactivate_consistency_check=FALSE)  
dim(DMR5_real_0.08_cut)

DMR5_random_0.08 = get_close_loci_AD3(DMC_random_0.08,
                                      number_of_lines=5,
                                      distance_threshold=500,
                                      distance_C_to_C=100, 
                                      max_region_size=2000,
                                      consistent_direction=TRUE,
                                      deactivate_consistency_check=FALSE)  
dim(DMR5_random_0.08)
```

Finally, parameters chosen to define DMRs, are as follow: **association of at least 5 CpG having a same methylation state, in a region of maximum 2000 bases, with a maximum distance of 100 bases between two successive selected CpG, according to pvalue threshold (pval < 0.07**, Supp. Table 2 of the paper in preparation, Chapter 4, section 1.2). These parameters allow us to make a compromise: identify a large normal of relevant DMRs (true positive), without detecting too many false positives.
  
  
Using these parameters, **432 DMRs are identified in alcohol-treated group, compared to control one.** These DMRs are called "DMR432" afterwards, later in the document.

#### 15. Annotation of DMR432 (mm9)¶

DMR432 were then annotated using **homer\_annotatePeaks**, a tool available on **Galaxeast** platform.

In [ ]:

```
%%bash

cd /mnt/e/methylome/fusion/methylkit/capture/test_randomisation/randomisation_en_ligne/premier_random

# Data formatting to use homer_annotatePeaks on Galaxeast
# To remove the header
awk 'NR!=1 {print $0}' DMR432_real_p0.07.txt > sh_DMR432_real_p0.07.txt

# To select only .bed informations (chr / start / end coordinates of DMRs)
cut -f1,2,3 sh_DMR432_real_p0.07.txt > bed_sh_DMR432_real_p0.07.txt

head bed_sh_DMR432_real_p0.07.txt

# Annotation of bed file with homer_annotatePeaks on Galaxeast:
# 283: homer_annotatePeaks on bed_sh_DMR432_real_p0.07_genome_mm9
# 284: homer_annotatePeaks on bed_sh_DMR432_real_p0.07_genome_mm9.log
```

#### 16.Conversion of region coordinates: mm9 to mm10¶

In order to combine methylome data with results of others analyses, mm9 coordinates of DMR432 were converted into mm10 coordinates, using **LiftOver** tool proposed by **UCSC**. For that, only chromosomal coordinates (chromosome;start;end) are kept to generate a .bed file. We used the file called **bed\_sh\_DMR432\_real\_p0.07.txt** in G:\methylome\fusion\post\_methylkit\DMR432\_pval0.07\bed.

Default settings of **LiftOver** tool have been retained:

- Minimum ratio of bases that must remap: 0.95
- BED 4 to BED 6 Options
  - Don’t allow multiple output regions
  - Minimum hit size in query: 0
  - Minimum chain size in target:0
- BED 12 Options
  - Min ratio of alignment blocks or exons that must map:1
  - If thickStart/thickEnd is not mapped, use the closest mapped base: No

All DMR432 were successfully converted. File containing mm10 coordinates, called hglft\_genome\_56b84\_3fda70.bed was renamed (**mm10\_DMR432.bed**) and saved in the following folder: G:\methylome\fusion\post\_methylkit\DMR432\_pval0.07\bed\mm10

Both mm9 and mm10 coordinates of the DMR432 were associated into a unique file to keep both annotations.

In [ ]:

```
%%bash 

----------------------------------------------------------------------
# To generate file containing both mm9 and mm10 coordinates of DMR432
----------------------------------------------------------------------

cd /mnt/g/methylome/fusion/post_methylkit/DMR432_pval0.07/bed/mm10

paste /mnt/g/methylome/fusion/post_methylkit/DMR432_pval0.07/bed/bed_sh_DMR432_real_p0.07.txt mm10_DMR432.bed > DMR432_mm9_mm10.txt

head DMR432_mm9_mm10.txt

# mm9 coordinates are in the first columns, followed by mm10 coordinates
```

#### 17.Annotation of DMR432 (mm10)¶

Syntaxic annotation of regions of interest was performed using **bedtools intersect** with an annotation file (**biomart\_mm10.txt**) obtained with **BiomaRt** R package. This file contains:

- ensembl\_gene\_id: gene name
- chromosome\_name: chromosome/scaffold number
- strand
- start\_position: gene start (bp)
- end\_position: gene end (bp)
- entrezgene\_id:NCBI gene ID
- gene\_biotype: gene type
- mgi\_symbol: MGI symbol
- entrezgene\_accession: NCBI gene accession
- entrezgene\_description: NCBI gene description
- uniprot\_gn\_symbol: UniProtKB gene name symbol

In [ ]:

```
%%R

#--------------------------------------
# 1. Obtention of biomart_mm10.txt file
#--------------------------------------

# To load BiomaRt package
library("biomaRt")

# 1. To select database and reference genome
mm10 = useMart("ensembl", dataset="mmusculus_gene_ensembl")
# mm10 Mus musculus version used: Ensembl 97 Jul 2019 http://jul2019.archive.ensembl.org 

# 2. To create the dataset with information of interest
# It's necessary to do it in two steps, because there is a limitation 
# in the number of attributes that can be collected at the same time.
annot_mm10_part1<-getBM(attributes=c("ensembl_gene_id","chromosome_name",
                                     "strand", "start_position",
                                     "end_position","entrezgene_id",
                                     "gene_biotype","mgi_symbol",
                                     "entrezgene_accession"), mart=mm10)

annot_mm10_part2<-getBM(attributes=c("ensembl_gene_id","entrezgene_description",
                                     "uniprot_gn_symbol"), mart=mm10)

# Verifications
head(annot_mm10_part1)
head(annot_mm10_part2)
dim(annot_mm10_part1)
dim(annot_mm10_part2)

# To merge all the informations in one R object
annot_mm10 = merge(annot_mm10_part1, annot_mm10_part2, by="ensembl_gene_id", all=TRUE)

# Verifications
dim(annot_mm10)
head(annot_mm10)

# To save the file
write.table(annot_mm10,"E:/projet_DU_AD/results/annotation_mm10/Biomart/biomart_mm10.txt", quote= FALSE, sep="\t",row.names=FALSE)
```

In [ ]:

```
%%bash

#--------------------------------------
# 2. Combination of differentially methylated regions (DMR432-mm10) with biomart annotation file 
#--------------------------------------

# 2.1 Dataset formatting
mkdir -p /mnt/g/methylome/fusion/post_methylkit/DMR432_pval0.07/annotations/Biomart/mm10

cd /mnt/g/methylome/fusion/annotation/AD/mm10/Biomart/

head biomart_mm10.txt
```

In [ ]:

```
%%R

biomart_mm10 =read.table("G:/methylome/fusion/annotation/AD/mm10/Biomart/biomart_mm10.txt",
                         sep="\t", na.strings = "NA", fill=TRUE, quote="", header=TRUE)

head(biomart_mm10)

# To change "-1" and "1" strand encoding by "-" and "+"
for (i in 1:nrow(biomart_mm10)){
  if(biomart_mm10[i,"strand"]==1){
    biomart_mm10[i,"strand"]="+"
  }else if (biomart_mm10[i,"strand"]==-1){
    biomart_mm10[i,"strand"]="-"
  }else{
  }
}

# To change chromosome format 
biomart_mm10$chr = with(biomart_mm10,paste("chr",biomart_mm10[,"chromosome_name"],sep=""))

head(biomart_mm10)

# To combine informations, we will use bedtools intersect. It seems to work with a .bed containing exactly 9 columns. 
# uniprot_genename and entrezgene_description informations are removed, 
# because bedtools intersect doesn't work if these columns are kept.
# (bedtools intersect is not able to detect the file format if these columns are kept).
biomart_mm10_bis=biomart_mm10[,c("chr","start_position","end_position",
                                 "strand","entrezgene_accession", "gene_biotype",
                                 "mgi_symbol","ensembl_gene_id", "entrezgene_id")] 

write.table(biomart_mm10_bis,"G:/methylome/fusion/annotation/AD/mm10/Biomart/biomart_mm10_all_info.bed", sep = "\t", col.names=TRUE, row.names=FALSE, quote=FALSE)
```

In [ ]:

```
%%bash 

# 2.2 To sort files
# To sort Biomart annotation file
cd /mnt/g/methylome/fusion/annotation/AD/mm10/Biomart/

sed '1d' biomart_mm10_all_info.bed | sort -k1,1 -k2,2n > biomart_mm10_all_info_sorted.bed

head biomart_mm10_all_info_sorted.bed

# To sort mm10 .bed files containing differentially methylated regions (DMR432)
cd /mnt/g/methylome/fusion/post_methylkit/DMR432_pval0.07/bed/mm10

sort -k1,1 -k2,2n mm10_DMR432.bed > mm10_DMR432_sorted.bed

head mm10_DMR432_sorted.bed


#----------------------------------
# 3. To combine information, using bedtools intersect
#----------------------------------

bedtools intersect -wao -a /mnt/g/methylome/fusion/post_methylkit/DMR432_pval0.07/bed/mm10/mm10_DMR432_sorted.bed 
-b /mnt/g/methylome/fusion/annotation/AD/mm10/Biomart/biomart_mm10_all_info_sorted.bed > /mnt/g/methylome/fusion/post_methylkit/DMR432_pval0.07/annotations/Biomart/mm10/DMR432_annot_syntaxique_mm10_biomart.bed

head /mnt/g/methylome/fusion/post_methylkit/DMR432_pval0.07/annotations/Biomart/mm10/DMR432_annot_syntaxique_mm10_biomart.bed

wc -l /mnt/g/methylome/fusion/post_methylkit/DMR432_pval0.07/annotations/Biomart/mm10/DMR432_annot_syntaxique_mm10_biomart.bed
# 619 /mnt/g/methylome/fusion/post_methylkit/DMR432_pval0.07/annotations/Biomart/mm10/DMR432_annot_syntaxique_mm10_biomart.bed
# There are more rows than there are DMR432, meaning that some DMRs have several annotations and are thus represented by distinct rows
```

In [ ]:

```
%%R

#------------------------------------------------------------------------
# 4. To oncatenate annotation informations: one row = one DMR
#------------------------------------------------------------------------

# 4.1. To load DMR432 file containing biomart annotation.
DMR432_annot_mm10 = read.table(file = "G:/methylome/fusion/post_methylkit/DMR432_pval0.07/annotations/Biomart/mm10/DMR432_annot_syntaxique_mm10_biomart.bed", 
                               header = FALSE, sep = "\t", fill=TRUE)

head(DMR432_annot_mm10)
dim(DMR432_annot_mm10)

# To create the ID = chr-start-end
DMR432_annot_mm10_ID = cbind("ID" = paste(DMR432_annot_mm10$V1, 
                                          DMR432_annot_mm10$V2, DMR432_annot_mm10$V3, sep = ";"),
                             DMR432_annot_mm10)

head(DMR432_annot_mm10_ID)

# 4.2. To concatenate informations 
library(dplyr)

# Liste of unique ID  
liste_chrom_unique <- unique(DMR432_annot_mm10_ID$ID)


# empty matrice which will be filled during the loop
# Nb of rows = nb of unique ID | Nb of columns = nb of variables
treatment_file <- data.frame(matrix(NA,nrow=length(liste_chrom_unique),ncol=ncol(DMR432_annot_mm10_ID)))

colnames(treatment_file) <- colnames(DMR432_annot_mm10_ID)
treatment_file$ID <- liste_chrom_unique
for(i in 1:length(liste_chrom_unique)){

  # Filter according to ID k 
   DMR432_filt <- DMR432_annot_mm10_ID %>%
     filter(ID == liste_chrom_unique[i])


  # Collapsing of distincts elements of a variables for ID k 
  # If variable is composed of only one element ==> we keep this element only once 
  # If variable is composed of m distinct elements ==> even if elements can be repeated, we keep all the elements (with repetitions)
   for(j in 2:length(colnames(treatment_file))){
     if(length(unique(DMR432_filt[,j])) == 1){
       treatment_file[i,j] <- paste(unique(DMR432_filt[,j]), collapse=';')
     } else {
       treatment_file[i,j] <- paste(DMR432_filt[,j], collapse=';')  
     }
   }
 }

dim(treatment_file)

# To rename properly the column
colnames(treatment_file) = c("ID","chr_DMR","start_DMR","end_DMR",
                             "chr_annot","start_annot","end_annot", 
                             "strand", "entrezgene_accession", "gene_biotype",
                             "mgi_symbol", "ENSEMBL_ID","Entrez_ID","overlap_length")

head(treatment_file)

# To save the data into a file
write.table(treatment_file, "G:/methylome/fusion/post_methylkit/DMR432_pval0.07/annotations/Biomart/mm10/DMR432_unique_annot_syntaxique_mm10_biomart.txt", 
            col.names = TRUE, sep = "\t", quote=FALSE, row.names=FALSE)
```

#### 18. Combination of annotation and statistical informations¶

All data (*i.e.* statistical informations, mm9 and mm10 coordinates regions, annotation informations) are combined in a single file.

In [ ]:

```
%%bash

#------------------------------------------------------------------------------
# To combine DMR432 annotation and statistical informations
#------------------------------------------------------------------------------

#------------------------------------------------------------------------------
# 1. To have a common ID between the two files, add mm10 coordinates to file containing methylation information of DMR432: 
mkdir -p /mnt/g/methylome/fusion/post_methylkit/DMR432_pval0.07/rawdata/mm10

cd /mnt/g/methylome/fusion/post_methylkit/DMR432_pval0.07/rawdata/mm10

#It works only when files are in this order
paste /mnt/g/methylome/fusion/post_methylkit/DMR432_pval0.07/bed/mm10/DMR432_mm9_mm10.txt /mnt/g/methylome/fusion/post_methylkit/DMR432_pval0.07/rawdata/nh_DMR432_real_p0.07.txt > nh_DMR432_real_p0.07_mm9_mm10.txt

head nh_DMR432_real_p0.07_mm9_mm10.txt

wc -l nh_DMR432_real_p0.07_mm9_mm10.txt
# 432 nh_DMR432_real_p0.07_mm9_mm10.txt
```

In [ ]:

```
%%R
#------------------------------------------------------------------------------
# 2. To load file containing methylation information of DMR432 .
DMR432_meth = read.table(file = "G:/methylome/fusion/post_methylkit/DMR432_pval0.07/rawdata/mm10/nh_DMR432_real_p0.07_mm9_mm10.txt", 
                         sep = "\t", header=FALSE)

head(DMR432_meth)
dim(DMR432_meth)

colnames(DMR432_meth) = c("chr_mm9","start_mm9","end_mm9", 
                          "chr_mm10","start_mm10","end_mm10", 
                          "chr_mm9bis","start_mm9bis","end_mm9bis", 
                          "CpG_nb","pvalue","qvalue","meth.diff")

DMR432_meth$ID =  paste(DMR432_meth$chr_mm10, DMR432_meth$start_mm10, DMR432_meth$end_mm10, sep = ";")

# To add the ID and delete columns that are duplicated
DMR432_meth_ID = DMR432_meth[, c("ID","chr_mm9","start_mm9",
                                 "end_mm9","chr_mm10","start_mm10", 
                                 "end_mm10","CpG_nb","pvalue","qvalue",
                                 "meth.diff")]

#------------------------------------------------------------------------------
# 3. To combine DMR432 annotation and statistical informations
DMR432_meth_annot= merge(x=treatment_file, DMR432_meth_ID, by = "ID")

write.table(DMR432_meth_annot,
            "G:/methylome/fusion/post_methylkit/DMR432_pval0.07/annotations/Biomart/mm10/DMR432_unique_annot_mm10_BMT_meth.txt",
            col.names = TRUE, sep = "\t", quote=FALSE, row.names=FALSE)
```

#### 19.Identification of CpG islands into DMR432¶

In order to see if DMR432 observed upon prenatal alcohol exposure are located into CpG islands (CGi), DMR432 coordinates (mm10) were compared to mouse CGi ones (mm10 coordinates). For that, **list of CGi** was obtained on **UCSC Table browser** website, using following parameters:

- clade:Mammal
- genome: Mouse
- assembly: Dec. 2011 (GRCm38/mm10)
- group: Expression and Regulation
- track: CpG islands
- table: cpgIslandExt
- output format: Selected fields from primary related tables:
  - chrom: Reference sequence chromosome or scaffold
  - chromStart: Start position in chromosome
  - chromEnd: End position in chromosome
  - name: CpG Island
  - length: Island Length
  - cpgNum: Number of CpGs in island

List of CGi was saved into a file called **cpgIslandExt\_mm10\_ucsc.txt** (in folder /mnt/g/methylome/fusion/annotation/AD/mm10/CG\_islands).
  
  
**bedtools intersect** was used to identify overlaps between CGi and DMR432.

In [ ]:

```
%%bash

# --------------------------------------------------------------------
# 1. Obtention of CGi coordinates (mm10) and data formatting
# based on a script kindly given by Olivier Kirsh, UMR7216
# --------------------------------------------------------------------

cd /mnt/g/methylome/fusion/annotation/AD/mm10/CG_islands

head cpgIslandExt_mm10_ucsc.txt

wc -l cpgIslandExt_mm9_ucsc.txt
#16 023 cpgIslandExt_mm10_ucsc.txt

# replaced by
awk '{print $1"\t"$2"\t"$3"\t"$4"_"$5"\t"$6"\t"$7}' cpgIslandExt_mm10_ucsc.txt | sed '1d' | sort -k1,1 -k2,2n | head

awk '{print $1"\t"$2"\t"$3"\t"$4"_"$5"\t"$6"\t"$7}' cpgIslandExt_mm10_ucsc.txt | sed '1d' | sort -k1,1 -k2,2n > mm10_CpG_Island_sorted.bed

wc -l mm10_CpG_Island_sorted.bed
#16 023 mm10_CpG_Island_sorted.bed
```

In [ ]:

```
%%R 

#------------------------------------------------------------------------------
# 2. DMR432 file formatting
#------------------------------------------------------------------------------

# Three first columns of DMR432 file must contained .bed informations (chr / start / end) 

DMR432_meth_annot = DMR432_meth_annot[,c(2:4, 1, 5:24)]

head(DMR432_meth_annot)

write.table(DMR432_meth_annot, 
            "G:/methylome/fusion/post_methylkit/DMR432_pval0.07/annotations/Biomart/mm10/DMR432_unique_annot_mm10_BMT_meth.bed",
            col.names = TRUE, sep = "\t", quote=FALSE, row.names=FALSE)
```

In [ ]:

```
%%bash

cd /mnt/g/methylome/fusion/post_methylkit/DMR432_pval0.07/annotations/Biomart/mm10/

sed '1d' DMR432_unique_annot_mm10_BMT_meth.bed | sort -k1,1 -k2,2n  > DMR432_unique_annot_mm10_BMT_meth_sorted.bed
    
#------------------------------------------------------------------------------
# 3. Intersection of CGi and DMR432 coordinates (mm10)
#------------------------------------------------------------------------------

bedtools intersect -wao -a DMR432_unique_annot_mm10_BMT_meth_sorted.bed 
    -b /mnt/g/methylome/fusion/annotation/AD/mm10/CG_islands/mm10_CpG_Island_sorted.bed > DMR432_unique_annot_mm10_BMT_meth_cgi.bed

head DMR432_unique_annot_mm10_BMT_meth_cgi.bed
```

#### 20. Search for imprinted genes¶

In order to see if DMR432 observed upon prenatal alcohol exposure are located into imprinted genes, DMR432 gene names were compared to mouse imprinted genes from **geneimprint** website.

In [ ]:

```
%%bash

# List of imprinted genes from geneimprint website 
sample="AF313042
AF357341
AF357355
AF357359
AF357425
AF357426
AF357428
Air
AK155734
Ampd3
Ano1
Asb4
Ascl2
Atp10a
B830012L14Rik
Begain
Blcap
Calcr
Ccdc40
Cd81
Cdkn1c
Commd1
Copg2
Cpa4
Ctnna3
Dcn
Ddc
Dhcr7
Dio3
Dlk1
Dlx5
Ftx
Gab1
Gabra5
Gabrb3
Gabrg3
Gatm
Glis3
Gnai3
Gnas
Gpr1
Grb10
Gtl2
H13
H19
Htr2a
Hymai
Igf2
Igf2as
Igf2r
Il6
Impact
Inpp5f V2
Ins1
Ins2
Jpx
Kcnk9
Kcnq1
Kcnq1ot1
Klf14
L3mbtl
Lin28a
Lin28b
LOC101055709
Magel2
Magi2
Mcts2
Mest
Mir127
Mir134
Mir136
Mir154
Mir184
Mir296
Mir298
Mir335
Mir337
Mir370
Mir376b
Mir380
Mir410
Mir411
Mir431
Mirg
Mkrn1-ps1
Mkrn3
Musd2
Nap1l4
Nap1l5
Nctc1
Ndn
Nespas
Ngfb
Nnat
Ntm
Osbpl5
Peg10
Peg12
Peg13
Peg3
Peg3os
Phlda2
Plagl1
Pon1
Pon2
Pon3
Ppp1r9a
Pwcr1
Rasgrf1
Rb1
Rhox5
Rian
Rtl1
Sdhd
Sfmbt2
Sgce
Slc22a18
Slc22a2
Slc22a3
Slc38a4
Snrpn
Snurf
Snx14
Tfpi2
Tgfb1
Th
Tnfrsf22
Tnfrsf23
Tnfrsf26
Trp73
Trpm5
Tsix
Tspan32
Tssc4
U2af1-rs1
Ube3a
Usp29
Wt1
Xist
Zc3h12c
Zcchc13
Zdbf2
Zfat
Zfp127as
Zfp264
Zim1
Zim2
Zim3
Znf264
Zrsr1"

# Search for these imprinted genes names, among DMR432.
for i in ${sample}
do
echo "---------------------------------------------------"
echo "${i}"
grep "${i}" DMR432_unique_annot_mm10_BMT_meth_cgi.bed
done
```

nb: after further research, Dlx5 and Glis3, which are found among DMR432, are not imprinted genes in mouse.

Imprinted genes that are found among DMR432 are thus:

- Commd1 / Zrsr1
- Gab1
- Gnas
- Grb10
- H13
- Impact
- Inpp5f
- Nap1l5
- Peg10
- Peg13

#### 21. Number of DMRs observed upon PAE, in each methylome capture category¶

##### 21.1 Capture Information¶

DMR432 are observed among capture regions we have chosen. Capture is composed of regions, based on ENCODE available data (for active enhancers and promoters regions) or based on previous lab results (“other regions” group). Captured regions correspond to:

- (A) **active enhancers** in adult (8 weeks-old) mouse cortex (characterized by the **H3K27ac** histone mark, regions of +/- 500 bases from the middle of the peaks)
- (B) **mouse promoter regions** (+/- 500 bases from the transcription start site - TSS). We estimated that more than 75% of known promoters regions are included in the capture. Due to repetitive sequences that prevent the design of specific probes of some regions, and also because of annotation database information differences, some promoters regions couldn’t be studied.
- (C) **“Other regions”** group corresponds to:
  - (i) **HSF2 binding sites** found in unstressed mice cortices of embryos at E16.5 development stage (results of a ChIP-seq experiment, R. El Fatimy, A.L. Mouël, I. Massaoudi)
  - (ii) **differentially opened or closed regions** identified in isolated oligodendrocyte precursors (O4+ cells) from cortices of 5 days-old mice, after an **inflammatory stress** from P1 to P5, a period equivalent to the third trimester of pregnancy in human (results of an ATAC-seq experiment, A.L. Schang and D. Sabéran-Djoneidi, (Schang et al., 2018))
  - (iii) **enhancers** (characterized by the **H3K4me1** histone mark, +/- 100 bases from the middle of the peaks) of adult (8 weeks-old) mouse cortex among genes (and at +/-20 kb from genes ends) that were differentially expressed in microglia (CD11B+ cells), at different stages (i.e. postnatal day 1 (P1) + P5 + P10 or at P5 + P10 or at P45) upon an **inflammatory stress** from P1 to P5 (results of a microarray experiment, A.L. Schang, Krishnan et al., 2017).

To see the repartition of DMRs among methylome capture categories (promoters, active enhancers, others), DMR432 were intersected with methylome capture regions. Before that, capture was analysed to better characterize it notably we defined:

- 1. number of regions per capture category, since all regions of interest can't be captured due to technical limits. For details about the number of regions in each of this sub-category of “other regions” group, please see Supp. Fig. S1A of the paper in preparation, Chapter 4, section 1.2).
- 1. number of common region between categories (which regions are included into two or more capture categories?)
- 1. number of CpG in the capture, compared to the number of CpG in mouse genome (mm9)
- 1. number of nucleotides included in the (enlarged) capture
- 1. size of captured regions

In [ ]:

```
%%bash

# --------------------------------------
# 1. Number of regions per capture category
# --------------------------------------

# ========
# 1.a. Number of regions in unenlarged capture (original file)
# ========

cd /mnt/h/methylome/fusion/capture/capture_regions

conda activate chip

# Unenlarged capture file (original file)
head 151019_MM9_EDC_RM_EPI_capture_targets.bed

# To sort the file containing captured regions
sort -k1,1 -k2,2n 151019_MM9_EDC_RM_EPI_capture_targets.bed > Capture76800_NON_agrandi_sorted.bed

wc -l 151019_MM9_EDC_RM_EPI_capture_targets.bed
# 76 848 151019_MM9_EDC_RM_EPI_capture_targets.bed

wc -l Capture76800_NON_agrandi_sorted.bed
# 76 848 Capture76800_NON_agrandi_sorted.bed
# --> In the capture (not enlarged capture), 76 848 distinct regions are considered.

md5sum 151019_MM9_EDC_RM_EPI_capture_targets.bed
# 4c46a7f4730efffe28e912865b903aa9  151019_MM9_EDC_RM_EPI_capture_targets.bed

md5sum Capture76800_NON_agrandi_sorted.bed
# aeca37fbb3a9b5035dc04da071222ece  Capture76800_NON_agrandi_sorted.bed

# Check: to merge overlapped regions, to see if 76 848 regions in the capture are unique
bedtools merge -i Capture76800_NON_agrandi_sorted.bed > Capture76800_NON_agrandi_merge_sorted.bed

wc -l Capture76800_NON_agrandi_merge_sorted.bed
# 76 848 Capture76800_NON_agrandi_merge_sorted.bed
# --> Regions of the capture are unique and non-overlapping. 


# ========
# 1.b. Number of regions in enlarged capture 
# ========

# As a reminder, each region was extended up and downstream with 150 bases for the analysis
# to avoid removal of potential interesting sequences, at the border of capture regions.
# After this extension, bedtools merge was used to merge enlarged captured regions that overlaps with each other.

cd /mnt/h/methylome/fusion/capture/capture_regions

wc -l Capture_76800_et150seq.merge.sorted.bed
# 58 611 Capture_76800_et150seq.merge.sorted.bed
# --> In the enlarged capture, 58 611 distinct regions are considered.


# ========
# 1.c. Number of regions per capture category
# ========

cd /mnt/h/methylome/capture

# Number of regions of interest, in each category.
# nb: it corresponds to the regions we wanted to capture, 
# but not all these regions are in the capture, for diverse technical reasons.

wc -l *
#   (Cii) 823 ATAC-capture.txt
#   (B) 33 577 clean_500upstream500downstream_mouse_bis.txt
#   (A) 36 824 H3K27Ac_Cx_+_-100.txt 
#   (Ci) 4 226 HSF2MACSwo.bed
#   (Ciii) 2 317 P1_5_10_bed_+-20k_H3K4me1WB+-100.txt
#   (Ciii) 1 126 P45_bed_+-20k_H3K4me1WB+-100.txt
#   (Ciii) 6 304 P5_10_txt_+-20k_H3K4me1WB+-100.bed

# To sort these files, and keep unique regions
sort -k1,1 -k2,2n clean_500upstream500downstream_mouse_bis.txt | uniq > clean_promoters_uniq_sorted.txt
sort -k1,1 -k2,2n ATAC-capture.txt | uniq > ATAC_uniq_sorted.txt
sort -k1,1 -k2,2n H3K27Ac_Cx_+_-100.txt | uniq > H3K27Ac_uniq_sorted.txt
sort -k1,1 -k2,2n HSF2MACSwo.bed | uniq > ChIPHSF2_uniq_sorted.txt
sort -k1,1 -k2,2n P1_5_10_bed_+-20k_H3K4me1WB+-100.txt | uniq > P1_5_10_H3K4me1_uniq_sorted.txt
sort -k1,1 -k2,2n P5_10_txt_+-20k_H3K4me1WB+-100.bed | uniq > P5_10_H3K4me1_uniq_sorted.txt
sort -k1,1 -k2,2n P45_bed_+-20k_H3K4me1WB+-100.txt | uniq > P45_H3K4me1_uniq_sorted.txt

# To create a file corresponding to "other regions" group
cat ATAC_uniq_sorted.txt 
ChIPHSF2_uniq_sorted.txt 
P1_5_10_H3K4me1_uniq_sorted.txt 
P5_10_H3K4me1_uniq_sorted.txt P45_H3K4me1_uniq_sorted.txt | sort -k1,1 -k2,2n > all_except_prom_and_H3K27ac_sorted.txt

wc -l all_except_prom_and_H3K27ac_sorted.txt
# 14 800 all_except_prom_and_H3K17ac_sorted.txt

# To keep unique regions
uniq all_except_prom_and_H3K27ac_sorted.txt > all_except_prom_and_H3K17ac_uniq_sorted.txt

wc -l all_except_prom_and_H3K27ac_uniq_sorted.txt
#14 376 all_except_prom_and_H3K17ac_uniq_sorted.txt

# To merge overlapping regions, in order to avoid double counting of a given region
sample="clean_promoters ATAC H3K27Ac ChIPHSF2 P1_5_10_H3K4me1 P5_10_H3K4me1 P45_H3K4me1 all_except_prom_and_H3K27ac"

for file in ${sample}
do
echo $file
wc -l ${file}_uniq_sorted.txt
bedtools merge -i ${file}_uniq_sorted.txt > ${file}uniq_merge_sorted.txt
wc -l ${file}uniq_merge_sorted.txt
done

# clean_promoters
# 26 998 clean_promoters_uniq_sorted.txt
# 23 963 clean_promotersuniq_merge_sorted.txt

# ATAC-seq
# 823 ATAC_uniq_sorted.txt
# 823 ATACuniq_merge_sorted.txt

# H3K27Ac enhancers
# 36 825 H3K27Ac_uniq_sorted.txt
# 35 178 H3K27Acuniq_merge_sorted.txt

# ChIPHSF2
# 4 227 ChIPHSF2_uniq_sorted.txt
# 4 227 ChIPHSF2uniq_merge_sorted.txt

# P1_5_10_H3K4me1
# 2 318 P1_5_10_H3K4me1_uniq_sorted.txt
# 2 318 P1_5_10_H3K4me1uniq_merge_sorted.txt

# P5_10_H3K4me1
# 6 305 P5_10_H3K4me1_uniq_sorted.txt
# 6 305 P5_10_H3K4me1uniq_merge_sorted.txt

# P45_H3K4me1
# 1 127 P45_H3K4me1_uniq_sorted.txt
# 1 127 P45_H3K4me1uniq_merge_sorted.txt

# all_except_prom_and_H3K27ac
# 14 376 all_except_prom_and_H3K27ac_uniq_sorted.txt
# 14 356 all_except_prom_and_H3K27acuniq_merge_sorted.txt

# --> promoters and H3K27ac enhancers lists contain overlapping regions, that are now merged


# Since we considered enlarged capture for the analysis, 
# each region of these files were enlarged in the same way.
for file in ${sample}
do
echo $file
awk '{ print $1"\t"$2-150"\t"$3+150 }' ${file}uniq_merge_sorted.txt > agrandi_${file}_uniq_sorted.txt
bedtools merge -i agrandi_${file}_uniq_sorted.txt > agrandi_${file}_uniq_merge_sorted.txt
echo ${file}uniq_merge_sorted.txt
head ${file}uniq_merge_sorted.txt
echo agrandi_${file}_uniq_sorted.txt
head agrandi_${file}_uniq_sorted.txt
done

# Number of enlarged regions in each category
for file in ${sample}
do
wc -l agrandi_${file}_uniq_sorted.txt
wc -l agrandi_${file}_uniq_merge_sorted.txt
done 

# 23 963 agrandi_clean_promoters_uniq_sorted.txt
# 23 669 agrandi_clean_promoters_uniq_merge_sorted.txt
# 823 agrandi_ATAC_uniq_sorted.txt
# 823 agrandi_ATAC_uniq_merge_sorted.txt
# 35 178 agrandi_H3K27Ac_uniq_sorted.txt
# 34 201 agrandi_H3K27Ac_uniq_merge_sorted.txt
# 4 227 agrandi_ChIPHSF2_uniq_sorted.txt
# 3 875 agrandi_ChIPHSF2_uniq_merge_sorted.txt
# 2 318 agrandi_P1_5_10_H3K4me1_uniq_sorted.txt
# 2 318 agrandi_P1_5_10_H3K4me1_uniq_merge_sorted.txt
# 6 305 agrandi_P5_10_H3K4me1_uniq_sorted.txt
# 6 304 agrandi_P5_10_H3K4me1_uniq_merge_sorted.txt
# 1 127 agrandi_P45_H3K4me1_uniq_sorted.txt
# 1 127 agrandi_P45_H3K4me1_uniq_merge_sorted.txt
# 14 356 agrandi_all_except_prom_and_H3K27ac_uniq_sorted.txt
# 13 977 agrandi_all_except_prom_and_H3K27ac_uniq_merge_sorted.txt
# --> When enlarged, some regions overlapped (for promoters, H3K27ac enhancers, HSF2 binding site, H3K4me1 enhancers categories).

# Intersection between these files containing enlarged regions
# that are separated by category, and the enlarged capture 
# to identify regions that are really considered in the capture
# nb: -wao option writes the original A and B entries plus the number 
# of base pairs of overlap between the two features. 
# However, A features w/o overlap are also reported with a NULL B feature and overlap.
for file in ${sample}
do
echo $file
bedtools intersect -wao -a agrandi_${file}_uniq_merge_sorted.txt -b /mnt/h/methylome/fusion/capture/capture_regions/Capture_76800_et150seq.merge.sorted.bed > wao_capt_agrandi_${file}.txt
wc -l wao_capt_agrandi_${file}.txt
done 

# clean_promoters
# 24 012 wao_capt_agrandi_clean_promoters.txt
# ATAC
# 828 wao_capt_agrandi_ATAC.txt
# H3K27Ac
# 34 279 wao_capt_agrandi_H3K27Ac.txt
# ChIPHSF2
# 3 896 wao_capt_agrandi_ChIPHSF2.txt
# P1_5_10_H3K4me1
# 2 318 wao_capt_agrandi_P1_5_10_H3K4me1.txt
# P5_10_H3K4me1
# 6 304 wao_capt_agrandi_P5_10_H3K4me1.txt
# P45_H3K4me1
# 1 128 wao_capt_agrandi_P45_H3K4me1.txt
# all_except_prom_and_H3K27ac
# 14 004 wao_capt_agrandi_all_except_prom_and_H3K17ac.txt


# Intersection between these files containing enlarged regions
# that are separated by category, and the enlarged capture 
# to identify regions that are really considered in the capture
# Here without -wao option
sample="clean_promoters ATAC H3K27Ac ChIPHSF2 P1_5_10_H3K4me1 P5_10_H3K4me1 P45_H3K4me1 all_except_prom_and_H3K27ac"
for file in ${sample}
do
echo $file
bedtools intersect -a /mnt/h/methylome/fusion/capture/capture_regions/Capture_76800_et150seq.merge.sorted.bed -b agrandi_${file}_uniq_merge_sorted.txt > in_big_capt_${file}.txt
wc -l in_big_capt_${file}.txt
done

# clean_promoters
# 23 769 in_big_capt_clean_promoters.txt
# ATAC
# 773 in_big_capt_ATAC.txt
# H3K27Ac
# 34 279 in_big_capt_H3K27Ac.txt
# ChIPHSF2
# 2 111 in_big_capt_ChIPHSF2.txt
# P1_5_10_H3K4me1
# 2 316 in_big_capt_P1_5_10_H3K4me1.txt
# P5_10_H3K4me1
# 6 248 in_big_capt_P5_10_H3K4me1.txt
# P45_H3K4me1
# 1 128 in_big_capt_P45_H3K4me1.txt
# all_except_prom_and_H3K27ac
# 12 108 in_big_capt_all_except_prom_and_H3K27ac.txt
# --> It corresponds to captured regions that are really considered, in each category. 


 
# =============================================================================


# --------------------------------------
# 2.1. Number of common regions between categories (which regions are included into two or more capture categories?)
# --------------------------------------

# To sort files of interest
sort -k1,1 -k2,2n in_big_capt_all_except_prom_and_H3K27ac.txt > in_big_capt_all_except_prom_and_H3K27ac_sorted.txt
sort -k1,1 -k2,2n in_big_capt_clean_promoters.txt > in_big_capt_clean_promoters_sorted.txt
sort -k1,1 -k2,2n in_big_capt_H3K27Ac.txt > in_big_capt_H3K27Ac_sorted.txt

# Common regions between promoters and H3K27Ac enhancers
bedtools intersect -a in_big_capt_clean_promoters_sorted.txt -b in_big_capt_H3K27Ac_sorted.txt > common_in_capt_prom_H3K27.txt

# Common regions between promoters and other regions (= all except H3K27ac enhancers)
bedtools intersect -a in_big_capt_clean_promoters_sorted.txt -b in_big_capt_all_except_prom_and_H3K27ac_sorted.txt > common_prom_all_sans_H3K27ac.txt

# Common regions between H3K27ac enhancers and other regions (=all except promoters)
bedtools intersect -a in_big_capt_H3K27Ac_sorted.txt -b in_big_capt_all_except_prom_and_H3K27ac_sorted.txt > common_H3K27_all_sans_prom.txt

wc -l common*
#  1 969 common_H3K27_all_sans_prom.txt
#  8 144 common_in_capt_prom_H3K27.txt
#  1 285 common_prom_all_sans_H3K27ac.txt

# Common regions of all categories (promoters, H3K27ac promoters, other group)
bedtools multiinter -i in_big_capt_clean_promoters_sorted.txt in_big_capt_H3K27Ac_sorted.txt in_big_capt_all_except_prom_and_H3K17ac_sorted.txt > in_big_capt_all.txt
 
awk '$5~ /1,2,3/ {print $5}' in_big_capt_all.txt | wc -l
# 490 
 
# Venn diagram representation, using eulerr package
library(eulerr)

VennDiag <- euler(c( "enhancers (H3K27ac)" = 34279, "other regions" = 12108, "promoters" = 23769,"promoters&enhancers (H3K27ac)" = 8144, "enhancers (H3K27ac)&other regions" = 1969, "promoters&other regions" = 1285, "promoters&enhancers (H3K27ac)&other regions" = 490))
setwd("H:/methylome/capture/redo/agrandi/ok")
pdf("VennDiagramm - Capture repartition.pdf")
plot(VennDiag, counts = TRUE, font=1, cex=1, alpha=0.5,
     fill=c("darkgreen", "darkred", "grey"))
dev.off()


# --------------------------------------
# 2.2. Number of common regions between categories (which regions are included into two or more capture categories?)
# MORE detailed analysis. Here categories are 
# H3K27ac enhancers, promoters, HSF2 binding site, 
# differentially opened or closed regions upon inflammatory stress and H3K4me1 enhancers
# --------------------------------------

cd /mnt/h/methylome/capture/redo/agrandi/ok/

# To group H3K4me1 enhancers  
cat in_big_capt_P1_5_10_H3K4me1.txt in_big_capt_P5_10_H3K4me1.txt in_big_capt_P45_H3K4me1.txt | sort -k1,1 -k2,2n > in_big_capt_all_H3K4me1_unmerged_sorted.txt 

# To merge overlapping regions
bedtools merge -i in_big_capt_all_H3K4me1_unmerged_sorted.txt > in_big_capt_all_H3K4me1_merged.txt

# To obtain unique ID
sort -k1,1 -k2,2n in_big_capt_all_H3K4me1_merged.txt | uniq >  in_big_capt_all_H3K4me1_merged_uniq_sorted.txt
wc -l in_big_capt_all_H3K4me1_merged_uniq_sorted.txt
wc -l in_big_capt_P1_5_10_H3K4me1.txt in_big_capt_P5_10_H3K4me1.txt
wc -l in_big_capt_P5_10_H3K4me1.txt 
wc -l in_big_capt_P45_H3K4me1.txt
wc -l in_big_capt_all_H3K4me1_unmerged_sorted.txt

data1="H3K27Ac" 
data2="ChIPHSF2"
data3="clean_promoters"
data4="global_H3K4me1"
data5="ATAC"

cp in_big_capt_all_H3K4me1_merged_uniq_sorted.txt in_big_capt_global_H3K4me1.txt

# Two-by-two file crossing
bedtools intersect -a in_big_capt_$data1.txt -b in_big_capt_$data2.txt > /mnt/h/methylome/capture/redo/agrandi/ok/croisement/in_big_capt_common_${data1}_${data2}.txt

bedtools intersect -a in_big_capt_$data1.txt -b in_big_capt_$data3.txt > /mnt/h/methylome/capture/redo/agrandi/ok/croisement/in_big_capt_common_${data1}_${data3}.txt

bedtools intersect -a in_big_capt_$data1.txt -b in_big_capt_$data4.txt > /mnt/h/methylome/capture/redo/agrandi/ok/croisement/in_big_capt_common_${data1}_${data4}.txt

bedtools intersect -a in_big_capt_$data1.txt -b in_big_capt_$data5.txt > /mnt/h/methylome/capture/redo/agrandi/ok/croisement/in_big_capt_common_${data1}_${data5}.txt

bedtools intersect -a in_big_capt_$data2.txt -b in_big_capt_$data3.txt > /mnt/h/methylome/capture/redo/agrandi/ok/croisement/in_big_capt_common_${data2}_${data3}.txt

bedtools intersect -a in_big_capt_$data2.txt -b in_big_capt_$data4.txt > /mnt/h/methylome/capture/redo/agrandi/ok/croisement/in_big_capt_common_${data2}_${data4}.txt

bedtools intersect -a in_big_capt_$data2.txt -b in_big_capt_$data5.txt > /mnt/h/methylome/capture/redo/agrandi/ok/croisement/in_big_capt_common_${data2}_${data5}.txt

bedtools intersect -a in_big_capt_$data3.txt -b in_big_capt_$data4.txt > /mnt/h/methylome/capture/redo/agrandi/ok/croisement/in_big_capt_common_${data3}_${data4}.txt

bedtools intersect -a in_big_capt_$data3.txt -b in_big_capt_$data5.txt > /mnt/h/methylome/capture/redo/agrandi/ok/croisement/in_big_capt_common_${data3}_${data5}.txt

bedtools intersect -a in_big_capt_$data4.txt -b in_big_capt_$data5.txt > /mnt/h/methylome/capture/redo/agrandi/ok/croisement/in_big_capt_common_${data4}_${data5}.txt


# Three-by-three file crossing 
bedtools intersect -a in_big_capt_$data3.txt -b /mnt/h/methylome/capture/redo/agrandi/ok/croisement/in_big_capt_common_${data1}_${data2}.txt > /mnt/h/methylome/capture/redo/agrandi/ok/croisement/in_big_capt_common_${data1}_${data2}_${data3}.txt

bedtools intersect -a in_big_capt_$data4.txt -b /mnt/h/methylome/capture/redo/agrandi/ok/croisement/in_big_capt_common_${data1}_${data2}.txt > /mnt/h/methylome/capture/redo/agrandi/ok/croisement/in_big_capt_common_${data1}_${data2}_${data4}.txt

bedtools intersect -a in_big_capt_$data5.txt -b /mnt/h/methylome/capture/redo/agrandi/ok/croisement/in_big_capt_common_${data1}_${data2}.txt > /mnt/h/methylome/capture/redo/agrandi/ok/croisement/in_big_capt_common_${data1}_${data2}_${data5}.txt

bedtools intersect -a in_big_capt_$data4.txt -b /mnt/h/methylome/capture/redo/agrandi/ok/croisement/in_big_capt_common_${data1}_${data3}.txt > /mnt/h/methylome/capture/redo/agrandi/ok/croisement/in_big_capt_common_${data1}_${data3}_${data4}.txt

bedtools intersect -a in_big_capt_$data5.txt -b /mnt/h/methylome/capture/redo/agrandi/ok/croisement/in_big_capt_common_${data1}_${data3}.txt > /mnt/h/methylome/capture/redo/agrandi/ok/croisement/in_big_capt_common_${data1}_${data3}_${data5}.txt

bedtools intersect -a in_big_capt_$data5.txt -b /mnt/h/methylome/capture/redo/agrandi/ok/croisement/in_big_capt_common_${data1}_${data4}.txt > /mnt/h/methylome/capture/redo/agrandi/ok/croisement/in_big_capt_common_${data1}_${data4}_${data5}.txt

bedtools intersect -a in_big_capt_$data4.txt -b /mnt/h/methylome/capture/redo/agrandi/ok/croisement/in_big_capt_common_${data2}_${data3}.txt > /mnt/h/methylome/capture/redo/agrandi/ok/croisement/in_big_capt_common_${data2}_${data3}_${data4}.txt

bedtools intersect -a in_big_capt_$data5.txt -b /mnt/h/methylome/capture/redo/agrandi/ok/croisement/in_big_capt_common_${data2}_${data3}.txt > /mnt/h/methylome/capture/redo/agrandi/ok/croisement/in_big_capt_common_${data2}_${data3}_${data5}.txt

bedtools intersect -a in_big_capt_$data5.txt -b /mnt/h/methylome/capture/redo/agrandi/ok/croisement/in_big_capt_common_${data3}_${data4}.txt > /mnt/h/methylome/capture/redo/agrandi/ok/croisement/in_big_capt_common_${data3}_${data4}_${data5}.txt

bedtools intersect -a in_big_capt_$data5.txt -b /mnt/h/methylome/capture/redo/agrandi/ok/croisement/in_big_capt_common_${data2}_${data4}.txt > /mnt/h/methylome/capture/redo/agrandi/ok/croisement/in_big_capt_common_${data2}_${data4}_${data5}.txt

cd /mnt/h/methylome/capture/redo/agrandi/ok/croisement/

# Four-by-four file crossing
bedtools intersect -a in_big_capt_common_${data2}_${data3}.txt -b in_big_capt_common_${data4}_${data5}.txt > /mnt/h/methylome/capture/redo/agrandi/ok/croisement/in_big_capt_common_${data2}_${data3}_${data4}_${data5}.txt

bedtools intersect -a in_big_capt_common_${data1}_${data3}.txt -b in_big_capt_common_${data4}_${data5}.txt > /mnt/h/methylome/capture/redo/agrandi/ok/croisement/in_big_capt_common_${data1}_${data3}_${data4}_${data5}.txt

bedtools intersect -a in_big_capt_common_${data1}_${data2}.txt -b in_big_capt_common_${data4}_${data5}.txt > /mnt/h/methylome/capture/redo/agrandi/ok/croisement/in_big_capt_common_${data1}_${data2}_${data4}_${data5}.txt

bedtools intersect -a in_big_capt_common_${data1}_${data2}.txt -b in_big_capt_common_${data3}_${data5}.txt > /mnt/h/methylome/capture/redo/agrandi/ok/croisement/in_big_capt_common_${data1}_${data2}_${data3}_${data5}.txt

bedtools intersect -a in_big_capt_common_${data1}_${data2}.txt -b in_big_capt_common_${data3}_${data4}.txt > /mnt/h/methylome/capture/redo/agrandi/ok/croisement/in_big_capt_common_${data1}_${data2}_${data3}_${data4}.txt

#  All files crossing
bedtools intersect -a /mnt/h/methylome/capture/redo/agrandi/ok/croisement/in_big_capt_common_${data1}_${data2}_${data3}_${data4}.txt -b in_big_capt_$data5.txt > /mnt/h/methylome/capture/redo/agrandi/ok/croisement/in_big_capt_common_${data1}_${data2}_${data3}_${data4}_${data5}.txt


# Results: 
cd /mnt/h/methylome/capture/redo/agrandi/ok/croisement/2par2

wc -l *
#      1 in_big_capt_common_ChIPHSF2_ATAC.txt
#     53 in_big_capt_common_ChIPHSF2_clean_promoters.txt
#     15 in_big_capt_common_ChIPHSF2_global_H3K4me1.txt
#     32 in_big_capt_common_clean_promoters_ATAC.txt
#   1202 in_big_capt_common_clean_promoters_global_H3K4me1.txt
#     17 in_big_capt_common_global_H3K4me1_ATAC.txt
#    103 in_big_capt_common_H3K27Ac_ATAC.txt
#     72 in_big_capt_common_H3K27Ac_ChIPHSF2.txt
#   8144 in_big_capt_common_H3K27Ac_clean_promoters.txt
#   1801 in_big_capt_common_H3K27Ac_global_H3K4me1.txt
#  11440 total

cd /mnt/h/methylome/capture/redo/agrandi/ok/croisement/3par3/
wc -l *
#     0 in_big_capt_common_ChIPHSF2_clean_promoters_ATAC.txt
#     1 in_big_capt_common_ChIPHSF2_clean_promoters_global_H3K4me1.txt
#     0 in_big_capt_common_ChIPHSF2_global_H3K4me1_ATAC.txt
#     1 in_big_capt_common_clean_promoters_global_H3K4me1_ATAC.txt
#     0 in_big_capt_common_H3K27Ac_ChIPHSF2_ATAC.txt
#     6 in_big_capt_common_H3K27Ac_ChIPHSF2_clean_promoters.txt
#     2 in_big_capt_common_H3K27Ac_ChIPHSF2_global_H3K4me1.txt
#    13 in_big_capt_common_H3K27Ac_clean_promoters_ATAC.txt
#   471 in_big_capt_common_H3K27Ac_clean_promoters_global_H3K4me1.txt
#     4 in_big_capt_common_H3K27Ac_global_H3K4me1_ATAC.txt
#   498 total

cd /mnt/h/methylome/capture/redo/agrandi/ok/croisement/4par4 
wc -l *
# 0 in_big_capt_common_ChIPHSF2_clean_promoters_global_H3K4me1_ATAC.txt
# 0 in_big_capt_common_H3K27Ac_ChIPHSF2_clean_promoters_ATAC.txt
# 0 in_big_capt_common_H3K27Ac_ChIPHSF2_clean_promoters_global_H3K4me1.txt
# 0 in_big_capt_common_H3K27Ac_ChIPHSF2_global_H3K4me1_ATAC.txt
# 0 in_big_capt_common_H3K27Ac_clean_promoters_global_H3K4me1_ATAC.txt
# 0 total

# input files (+ others files): 
cd /mnt/h/methylome/capture/redo/agrandi/ok
wc -l in_big_capt_*
#   12108 in_big_capt_all_except_prom_and_H3K27ac_sorted.txt
#   12108 in_big_capt_all_except_prom_and_H3K27ac.txt
#    9258 in_big_capt_all_H3K4me1_merged.txt
#    9258 in_big_capt_all_H3K4me1_merged_uniq_sorted.txt
#    9692 in_big_capt_all_H3K4me1_unmerged_sorted.txt
#   80961 in_big_capt_all.txt
#     773 in_big_capt_ATAC.txt
#    2111 in_big_capt_ChIPHSF2.bed
#    2111 in_big_capt_ChIPHSF2.txt
#   23769 in_big_capt_clean_promoters_sorted.txt
#    4487 in_big_capt_clean_promoters_sorted.xlsx
#   23769 in_big_capt_clean_promoters.txt
#    9258 in_big_capt_global_H3K4me1.txt
#   34279 in_big_capt_H3K27Ac_sorted.txt
#   34279 in_big_capt_H3K27Ac.txt
#    2316 in_big_capt_P1_5_10_H3K4me1.txt
#    1128 in_big_capt_P45_H3K4me1.txt
#    6248 in_big_capt_P5_10_H3K4me1.txt
#  277913 total


# =============================================================================


# --------------------------------------
# 3. Number of CpG in the capture, compared to the number of CpG in mouse genome (mm9)
# --------------------------------------

# a. Capture file must be converted into a .fasta file
bedtools getfasta -fo Capture76800_NON_agrandi_merge_sorted.fa 
    -fi /Users/UMR7216/Documents/Mezger/methylome/genome-mm9/mm9.fa -bed Capture76800_NON_agrandi_merge_sorted.bed 

bedtools getfasta -fo Capture_76800_et150seq.merge.sorted.fa 
    -fi /Users/UMR7216/Documents/Mezger/methylome/genome-mm9/mm9.fa -bed Capture_76800_et150seq.merge.sorted.bed 
# Feature (chr13:120283547-120284433) beyond the length of chr13 size (120284312 bp).  Skipping.


# b. Number of CpGs, using seqkit tool
conda activate chip
conda install seqkit

# I copied and pasted files into H:\methylome\fusion\capture\capture_region
cd /mnt/h/methylome/fusion/capture/capture_regions/

#  Number of CpGs ("forward" strand only)
seqkit locate --ignore-case --pattern "(CG)" 
    --only-positive-strand Capture_76800_et150seq.merge.sorted.fa > /mnt/h/methylome/fusion/capture/nb_CpG_1strand_Capture_76800_et150seq.merge.sorted.txt

seqkit locate --ignore-case --pattern "(CG)" 
    --only-positive-strand Capture76800_NON_agrandi_merge_sorted.fa > /mnt/h/methylome/fusion/capture/nb_CpG_1strand_Capture76800_NON_agrandi_merge_sorted.txt

# Number of CpGs ("forward and reverse" strands)
seqkit locate --ignore-case --pattern "(CG)" 
    Capture_76800_et150seq.merge.sorted.fa > /mnt/h/methylome/fusion/capture/nb_CpG_2strands_Capture_76800_et150seq.merge.sorted.txt

seqkit locate --ignore-case --pattern "(CG)" 
    Capture76800_NON_agrandi_merge_sorted.fa > /mnt/h/methylome/fusion/capture/nb_CpG_2strands_Capture76800_NON_agrandi_merge_sorted.txt

#  Number of CpGs ("forward" strand only)
wc -l /mnt/h/methylome/fusion/capture/nb_CpG_1strand_Capture_76800_et150seq.merge.sorted.txt
# 1 822 332 /mnt/h/methylome/fusion/capture/nb_CpG_1strand_Capture_76800_et150seq.merge.sorted.txt

wc -l /mnt/h/methylome/fusion/capture/nb_CpG_1strand_Capture76800_NON_agrandi_merge_sorted.txt
# 1 537 452 /mnt/h/methylome/fusion/capture/nb_CpG_1strand_Capture76800_NON_agrandi_merge_sorted.txt

# Number of CpGs ("forward and reverse" strands)
wc -l /mnt/h/methylome/fusion/capture/nb_CpG_2strands_Capture_76800_et150seq.merge.sorted.txt
# 3 644 663 /mnt/h/methylome/fusion/capture/nb_CpG_2strands_Capture_76800_et150seq.merge.sorted.txt

wc -l /mnt/h/methylome/fusion/capture/nb_CpG_2strands_Capture76800_NON_agrandi_merge_sorted.txt
# 3 074 903 /mnt/h/methylome/fusion/capture/nb_CpG_2strands_Capture76800_NON_agrandi_merge_sorted.txt
```

In [ ]:

```
%%R 

# c. Number of CpG in mouse genome
# analysis based on a script kindly given by Olivier Kirsh, UMR7216

source("https://bioconductor.org/biocLite.R")
biocLite("Biostrings")
biocLite("BSgenome")

# To choose species of interest
biocLite("BSgenome.Mmusculus.UCSC.mm9")
library("BSgenome.Mmusculus.UCSC.mm9") 

# Once the "BSgenome.Mmusculus.UCSC.mm9" is loaded, mouse genome is in R memory as the Mmusculus object
#Mmusculus
#Mouse genome:
  # organism: Mus musculus (Mouse)
  # provider: UCSC
  # provider version: mm9
  # release date: Jul. 2007
  # release name: NCBI Build 37
  # 35 sequences:
  #   chr1         chr2         chr3         chr4         chr5         chr6         chr7         chr8         chr9         chr10        chr11        chr12        chr13       
  #   chr14        chr15        chr16        chr17        chr18        chr19        chrX         chrY         chrM         chr1_random  chr3_random  chr4_random  chr5_random 
  #   chr7_random  chr8_random  chr9_random  chr13_random chr16_random chr17_random chrX_random  chrY_random  chrUn_random                                                    
  # (use 'seqnames()' to see all the sequence names, use the '$' or '[[' operator to access a given sequence)
  
# How many chromosomes?
length(names(Mmusculus))
#[1] 35

# Which chromosomes?
names(Mmusculus)
#[1] "chr1"         "chr2"         "chr3"         "chr4"         "chr5"         "chr6"         "chr7"         "chr8"         "chr9"         "chr10"        "chr11"       
#[12] "chr12"        "chr13"        "chr14"        "chr15"        "chr16"        "chr17"        "chr18"        "chr19"        "chrX"         "chrY"         "chrM"        
#[23] "chr1_random"  "chr3_random"  "chr4_random"  "chr5_random"  "chr7_random"  "chr8_random"  "chr9_random"  "chr13_random" "chr16_random" "chr17_random" "chrX_random" 
#[34] "chrY_random"  "chrUn_random"

# To select main chrom 
mychrom_main = names(Mmusculus)[1:22]

# how many CpG in main chromosomes?
c1= countPattern("CG",Mmusculus$chr1)
c2= countPattern("CG",Mmusculus$chr2)         
c3= countPattern("CG",Mmusculus$chr3)
c4= countPattern("CG",Mmusculus$chr4)    
c5= countPattern("CG",Mmusculus$chr5)
c6= countPattern("CG",Mmusculus$chr6)    
c7= countPattern("CG",Mmusculus$chr7)
c8= countPattern("CG",Mmusculus$chr8)    
c9= countPattern("CG",Mmusculus$chr9)
c10= countPattern("CG",Mmusculus$chr10)    
c11= countPattern("CG",Mmusculus$chr11)
c12= countPattern("CG",Mmusculus$chr12)    
c13= countPattern("CG",Mmusculus$chr13)
c14= countPattern("CG",Mmusculus$chr14)    
c15= countPattern("CG",Mmusculus$chr15)
c16= countPattern("CG",Mmusculus$chr16)    
c17= countPattern("CG",Mmusculus$chr17)
c18= countPattern("CG",Mmusculus$chr18)    
c19= countPattern("CG",Mmusculus$chr19)
c20= countPattern("CG",Mmusculus$chrX)    
c21= countPattern("CG",Mmusculus$chrY)
c22= countPattern("CG",Mmusculus$chrM)    

sumCumul = 0
for (i in 1:22){
  sum_chr=get(paste0("c",i))
  sumCumul=sum(sumCumul, sum_chr)
  print(paste0("c",i))
  print(sumCumul)
}
#[1] "c1"
# [1] 1471973
# etc...
# [1] "c22"
# [1] 21 342 779
# --> There are 21 342 779 CpG in mm9


# d. Proportion of CpG of the mouse genome, included in the enlarged and merged capture
# (nb of CpG in the capture / nb of CpG in mm9)*100
(1 822 332/21 342 779)*100
# --> 8.538401 % of mm9 CpG are in the capture.
```

In [ ]:

```
%%bash 

# =============================================================================


# --------------------------------------
# 4. Number of nucleotides included in the (enlarged and merged) capture
# --------------------------------------

# "forward" strand only
cd /mnt/h/methylome/fusion/capture/capture_regions

grep -v ">" Capture_76800_et150seq.merge.sorted.fa | wc
#   58610   58610 81374638

grep -v ">" Capture76800_NON_agrandi_merge_sorted.fa | wc
#  76848   76848 62493124
```

In [ ]:

```
%%R


# =============================================================================


# --------------------------------------
# 5.size of captured regions (enlarged and merged capture regions)
# --------------------------------------

# boxplots and descriptive statistics
# global capture (enlarged and mergerd regions)
setwd("H:/methylome/fusion/capture/capture_regions")

capt_all = read.delim("Capture_76800_et150seq.merge.sorted.bed", header = FALSE)

capt_all$dim = with(capt_all,V3-V2)

head(capt_all)

library(ggplot2)

p <- ggplot(capt_all, aes(x="",y=dim)) + geom_boxplot(outlier.colour="black", outlier.shape=1,
                                                       outlier.size=1, notch=FALSE)
p

ggsave("H:/methylome/fusion/capture/capture_regions/size_Capture_76800_et150seq.merge.sorted.png", p)

summary(capt_all)
#       V1              V2                  V3                 dim      
# chr2   : 4812   Min.   :     4745   Min.   :     5935   Min.   : 360  
# chr7   : 4146   1st Qu.: 37192195   1st Qu.: 37193568   1st Qu.:1327  
# chr11  : 4091   Median : 74374495   Median : 74376068   Median :1376  
# chr1   : 3646   Mean   : 75964980   Mean   : 75966367   Mean   :1387  
# chr4   : 3552   3rd Qu.:109578877   3rd Qu.:109580862   3rd Qu.:1415  
# chr5   : 3478   Max.   :197067546   Max.   :197068282   Max.   :7487  
# (Other):34886


# promoters regions
setwd("H:/methylome/capture/redo/agrandi/ok")

capt_prom = read.delim("in_big_capt_clean_promoters_sorted.txt", header = FALSE)

capt_prom$region_size = with(capt_prom,V3-V2)

dim(capt_prom)

p_prom <- ggplot(capt_prom, aes(x="",y=region_size)) + geom_boxplot(outlier.colour="black", 
                                                                    outlier.shape=1, outlier.size=1, notch=FALSE)

p_prom

ggsave("H:/methylome/capture/redo/agrandi/ok/size_in_big_capt_clean_promoters_sorted.png", p_prom)

summary(capt_prom)
#       V1              V2                  V3             region_size  
# chr7   : 2161   Min.   :     4802   Min.   :     5935   Min.   :   7  
# chr2   : 2017   1st Qu.: 36334770   1st Qu.: 36335925   1st Qu.:1300  
# chr11  : 1756   Median : 73929260   Median : 73930560   Median :1300  
# chr4   : 1467   Mean   : 75371059   Mean   : 75372386   Mean   :1327  
# chr5   : 1357   3rd Qu.:109605564   3rd Qu.:109606823   3rd Qu.:1300  
# chr1   : 1324   Max.   :197003259   Max.   :197004559   Max.   :5721  
# (Other):13687  


# H3K27ac enhancers regions
setwd("H:/methylome/capture/redo/agrandi/ok")

capt_enh = read.delim("in_big_capt_H3K27Ac_sorted.txt", header = FALSE)

capt_enh$region_size = with(capt_enh,V3-V2)

dim(capt_enh)

p_enh <- ggplot(capt_enh, aes(x="",y=region_size)) + geom_boxplot(outlier.colour="black", 
                                                                  outlier.shape=1, outlier.size=1, notch=FALSE)

p_enh

ggsave("H:/methylome/capture/redo/agrandi/ok/size_in_big_capt_H3K27Ac_sorted.png", p_enh)

summary(capt_enh)
#       V1              V2                  V3             region_size  
# chr2   : 2737   Min.   :   233769   Min.   :   235069   Min.   :   2  
# chr11  : 2392   1st Qu.: 37884352   1st Qu.: 37885652   1st Qu.:1300  
# chr1   : 2188   Median : 74097424   Median : 74098724   Median :1300  
# chr7   : 2148   Mean   : 76148775   Mean   : 76150132   Mean   :1357  
# chr4   : 2100   3rd Qu.:108868036   3rd Qu.:108869336   3rd Qu.:1300  
# chr5   : 2081   Max.   :196956843   Max.   :196958143   Max.   :5962  
# (Other):20633 


# Other regions
setwd("H:/methylome/capture/redo/agrandi/ok")

capt_other = read.delim("in_big_capt_all_except_prom_and_H3K17ac_sorted.txt", header = FALSE)

capt_other$region_size = with(capt_other,V3-V2)

dim(capt_other)

p_other <- ggplot(capt_enh, aes(x="",y=region_size)) + geom_boxplot(outlier.colour="black", 
                                                                    outlier.shape=1, outlier.size=1, notch=FALSE)

p_other

ggsave("H:/methylome/capture/redo/agrandi/ok/size_in_big_capt_all_except_prom_and_H3K17ac_sorted.png", p_other)

summary(capt_other)
#       V1             V2                  V3             region_size    
# chr2   : 977   Min.   :   112367   Min.   :   112909   Min.   :   2.0  
# chr11  : 921   1st Qu.: 35437864   1st Qu.: 35438256   1st Qu.: 500.0  
# chr1   : 798   Median : 75762600   Median : 75763100   Median : 500.0  
# chr10  : 763   Mean   : 76632301   Mean   : 76632801   Mean   : 500.1  
# chr5   : 723   3rd Qu.:110600375   3rd Qu.:110600875   3rd Qu.: 500.0  
# chr4   : 722   Max.   :197067607   Max.   :197068217   Max.   :6142.0  
# (Other):7204
```

##### 21.2 Number of DMRs in each methylome capture category¶

To see the repartition of DMRs among methylome capture category (promoters, H3K27ac enhancers, others), DMR432 were intersected with methylome capture regions, using our own R function, called **find\_overlaps\_AD\_table**.
  
  
For this analysis, a distance tolerance of 500bases between DMR432 and captured regions was allowed.

In [ ]:

```
%%bash

# ------------------------------------------------------------------
# Intersection between DMR432 and enlarged + merged captured regions
# ------------------------------------------------------------------

# a.DMR432 file
# File must be sorted by coordinates
cd /mnt/e/methylome/fusion/post_methylkit/DMR432_pval0.07/bed

sort -k1,1 -k2,2n bed_sh_DMR432_real_p0.07.txt > sorted_bed_sh_DMR432_real_p0.07.txt
```

In [ ]:

```
%%R

# b.Function to intersect captured and DMR432 regions
find_overlaps_AD_table <- function (regions_1,regions_2,tolerance=0)
{
  common_region_table = NULL
  number_of_regions_1 <- nrow(regions_1)
  number_of_regions_2 <- nrow(regions_2)
  
  overlapped_is <- c()
  overlapped_js <- c()
  
  for (i in 1:number_of_regions_1) {
    
    for (j in 1:number_of_regions_2) {
      
      if (as.character(regions_1[i,1])==as.character(regions_2[j,1])) {
        start_1 <- as.numeric(regions_1[i,2])
        end_1 <- as.numeric(regions_1[i,3])
        expanded_start_1 <- start_1 - tolerance
        expanded_end_1 <- end_1 + tolerance
        start_2 <- as.numeric(regions_2[j,2])
        end_2 <- as.numeric(regions_2[j,3])
        
        if (expanded_end_1>=start_2) {
          
          if (end_2>=expanded_start_1) {
            
            overlapped_is <- c(overlapped_is,i)
            overlapped_js <- c(overlapped_js,j)
            
            first_region <- paste(regions_1[i,1],start_1,end_1,sep="_")
            second_region <- paste(regions_2[j,1],start_2,end_2,sep="_")
            overlap_size <- min(end_1,end_2)-max(start_1,start_2)+1
            common_region_table = rbind(common_region_table,c(as.character(regions_1[i,1]),start_1,end_1, 
                                                              as.character(regions_2[j,1]),start_2,end_2, overlap_size))
            
          }
        } 
      }
    }
  }
  print(paste("Number matched in first data set:",length(unique(overlapped_is))))
  print(paste("Number matched in second data set:",length(unique(overlapped_js))))
  if (length(common_region_table) > 0){
    colnames(common_region_table) = c("chr_reg1", "start_reg1", "end_reg1", "chr_reg2", "start_reg2", "end_reg2","overlap_size")
  }else{
  }
  return(as.data.frame(common_region_table))
}


# To load DMR432 file
DMR432 = read.table(file="H:/methylome/fusion/post_methylkit/DMR432_pval0.07/bed/sorted_bed_sh_DMR432_real_p0.07.txt", header = FALSE, sep = "\t")

head(DMR432)
dim(DMR432)

# c. To load files containing captured regions, separated by category 
ChIP_capt =  read.table(file="H:/methylome/capture/redo/agrandi/ok/in_big_capt_ChIPHSF2.txt",  header = FALSE, sep = "\t")
head(ChIP_capt)
dim(ChIP_capt)
# [1] 2111    3


prom_capt = read.table(file="H:/methylome/capture/redo/agrandi/ok/in_big_capt_clean_promoters.txt",   header = FALSE, sep = "\t")
head(prom_capt)
dim(prom_capt)
# [1] 23769     3


ATAC_capt = read.table(file="H:/methylome/capture/redo/agrandi/ok/in_big_capt_ATAC.txt",  header = FALSE, sep = "\t")
head(ATAC_capt)
dim(ATAC_capt)
# [1] 773   3


H3K27_capt = read.table(file="H:/methylome/capture/redo/agrandi/ok/in_big_capt_H3K27Ac.txt",   header = FALSE, sep = "\t")
head(H3K27_capt)
dim(H3K27_capt)
# [1] 34279     3


P1_5_10_H3K4me1_capt = read.table(file="H:/methylome/capture/redo/agrandi/ok/in_big_capt_P1_5_10_H3K4me1.txt",   header = FALSE, sep = "\t")
head(P1_5_10_H3K4me1_capt)
dim(P1_5_10_H3K4me1_capt)
# [1] 2316    3


P5_10_H3K4me1_capt = read.table(file="H:/methylome/capture/redo/agrandi/ok/in_big_capt_P5_10_H3K4me1.txt",  header = FALSE, sep = "\t")
head(P5_10_H3K4me1_capt)
dim(P5_10_H3K4me1_capt)
# [1] 6248    3


P45_H3K4me1_capt = read.table(file="H:/methylome/capture/redo/agrandi/ok/in_big_capt_P45_H3K4me1.txt",  header = FALSE, sep = "\t")
head(P45_H3K4me1_capt)
dim(P45_H3K4me1_capt)
# [1] 1128    3


others_capt = read.table(file="H:/methylome/capture/redo/agrandi/ok/in_big_capt_all_except_prom_and_H3K27ac_sorted.txt",  header = FALSE, sep = "\t")
head(others_capt)
dim(others_capt)
# [1] 12108     3


# d1. Intersection between DMR432 and captured regions (ALL DMR432)
dir.create("H:/methylome/fusion/post_methylkit/DMR432_pval0.07/capture_separee/with_merge_file/DMR432_All/dist500", recursive=TRUE)


# large intersection (max. distance between DMR432 and captured region = 500bases)
setwd("H:/methylome/fusion/post_methylkit/DMR432_pval0.07/capture_separee/with_merge_file/DMR432_All/dist500")

for (file2 in list("ChIP_capt", "prom_capt", "H3K27_capt", "ATAC_capt", 
                   "P1_5_10_H3K4me1_capt", "P5_10_H3K4me1_capt", 
                   "P45_H3K4me1_capt", "others_capt")){
   name_dataset1 <- "DMR432"
   dataset1 <- get("DMR432")
   
   name_dataset2 <- file2
   dataset2 <- get(file2)
   output_name <- paste(name_dataset1, substr(name_dataset2,1,7),"dist500", sep = "_")
   print(output_name)
   assign(output_name, find_overlaps_AD_table(dataset1, dataset2, tolerance = 500))
   data=get(output_name)
   write.table(data, file = paste(name_dataset1, "_", name_dataset2, ".txt", sep = ""), sep ="\t", col.names=TRUE, row.names = FALSE, quote = FALSE)
 }
# [1] "DMR432_ChIP_ca_dist500"
# [1] "Number matched in first data set: 3"
# [1] "Number matched in second data set: 3"
# [1] "DMR432_prom_ca_dist500"
# [1] "Number matched in first data set: 134"
# [1] "Number matched in second data set: 131"
# [1] "DMR432_H3K27_c_dist500"
# [1] "Number matched in first data set: 345"
# [1] "Number matched in second data set: 345"
# [1] "DMR432_ATAC_ca_dist500"
# [1] "Number matched in first data set: 6"
# [1] "Number matched in second data set: 6"
# [1] "DMR432_P1_5_10_dist500"
# [1] "Number matched in first data set: 6"
# [1] "Number matched in second data set: 6"
# [1] "DMR432_P5_10_H_dist500"
# [1] "Number matched in first data set: 16"
# [1] "Number matched in second data set: 17"
# [1] "DMR432_P45_H3K_dist500"
# [1] "Number matched in first data set: 7"
# [1] "Number matched in second data set: 6"
# [1] "DMR432_others__dist500"
# [1] "Number matched in first data set: 35"
# [1] "Number matched in second data set: 35"


for (file2 in c("ChIP_capt", "prom_capt", "H3K27_capt", "ATAC_capt", 
                "P1_5_10_H3K4me1_capt", "P5_10_H3K4me1_capt", 
                "P45_H3K4me1_capt", "others_capt")){
   name_dataset1 <- "DMR432"
   name_dataset2 <- file2
   output_name <- paste(name_dataset1, substr(name_dataset2,1,7),"dist500", sep = "_")
   print(output_name)
   data=get(output_name)
   print(dim(data))
   print(head(data))
 }


# ==============================================


# d2. Intersection between DMR432 and captured regions (Hyper DMR432 / Hypo DMR432)

# To split Hypo and Hyper DMR432
DMR432_meth = read.table(file="H:/methylome/fusion/post_methylkit/DMR432_pval0.07/rawdata/DMR432_real_p0.07.txt", 
                         header = TRUE, sep = "\t")

head(DMR432_meth)
dim(DMR432_meth)

DMR432_Hyper = DMR432_meth[DMR432_meth[,"meth.diff"]>0,]
DMR432_Hypo = DMR432_meth[DMR432_meth[,"meth.diff"]<0,]

head(DMR432_Hyper)
dim(DMR432_Hyper)
# [1] 257   7

head(DMR432_Hypo)
dim(DMR432_Hypo)
# [1] 175   7


setwd("H:/methylome/fusion/post_methylkit/DMR432_pval0.07/capture_separee/with_merge_file/")
 
# large intersection (max. distance between DMR432 and captured region = 500bases)
dir.create("H:/methylome/fusion/post_methylkit/DMR432_pval0.07/capture_separee/with_merge_file/DMR432_Hyper/dist500", 
            recursive=TRUE)

dir.create("H:/methylome/fusion/post_methylkit/DMR432_pval0.07/capture_separee/with_merge_file/DMR432_Hypo/dist500", 
            recursive=TRUE)

for (file1 in c("Hyper","Hypo")){
   name_dataset1 <- paste("DMR432",file1, sep = '_')
   dataset1 <- get(name_dataset1)
   
   for (file2 in list("ChIP_capt", "prom_capt", "H3K27_capt", 
                      "ATAC_capt", "P1_5_10_H3K4me1_capt", 
                      "P5_10_H3K4me1_capt", "P45_H3K4me1_capt", "others_capt")){
     name_dataset2 <- file2
     dataset2 <- get(file2)
     output_name <- paste(name_dataset1, substr(name_dataset2,1,7),"dist500", sep = "_")
     print(output_name)
     assign(output_name, find_overlaps_AD_table(dataset1, dataset2, tolerance=500))
     data=get(output_name)
     print(dim(data))
     print(head(data))
     write.table(data, file = paste(name_dataset1, name_dataset2, "dist500.txt", sep = "_"), 
                 sep ="\t", col.names=TRUE, row.names = FALSE, quote = FALSE)
   }
 }
# [1] "DMR432_Hyper_ChIP_ca_dist500"
# [1] "Number matched in first data set: 0"
# [1] "Number matched in second data set: 0"
# [1] 0 0
# [1] "DMR432_Hyper_prom_ca_dist500"
# [1] "Number matched in first data set: 54"
# [1] "Number matched in second data set: 52"
# [1] 55  7
# [1] "DMR432_Hyper_H3K27_c_dist500"
# [1] "Number matched in first data set: 222"
# [1] "Number matched in second data set: 223"
# [1] 225   7
# [1] "DMR432_Hyper_ATAC_ca_dist500"
# [1] "Number matched in first data set: 3"
# [1] "Number matched in second data set: 3"
# [1] 3 7
# [1] "DMR432_Hyper_P1_5_10_dist500"
# [1] "Number matched in first data set: 2"
# [1] "Number matched in second data set: 2"
# [1] 2 7
# [1] "DMR432_Hyper_P5_10_H_dist500"
# [1] "Number matched in first data set: 7"
# [1] "Number matched in second data set: 7"
# [1] 7 7
# [1] "DMR432_Hyper_P45_H3K_dist500"
# [1] "Number matched in first data set: 3"
# [1] "Number matched in second data set: 2"
# [1] 3 7
# [1] "DMR432_Hyper_others__dist500"
# [1] "Number matched in first data set: 15"
# [1] "Number matched in second data set: 14"
# [1] 15  7
# [1] "DMR432_Hypo_ChIP_ca_dist500"
# [1] "Number matched in first data set: 3"
# [1] "Number matched in second data set: 3"
# [1] 3 7
# [1] "DMR432_Hypo_prom_ca_dist500"
# [1] "Number matched in first data set: 80"
# [1] "Number matched in second data set: 80"
# [1] 80  7
# [1] "DMR432_Hypo_H3K27_c_dist500"
# [1] "Number matched in first data set: 123"
# [1] "Number matched in second data set: 123"
# [1] 124   7
# [1] "DMR432_Hypo_ATAC_ca_dist500"
# [1] "Number matched in first data set: 3"
# [1] "Number matched in second data set: 3"
# [1] 3 7
# [1] "DMR432_Hypo_P1_5_10_dist500"
# [1] "Number matched in first data set: 4"
# [1] "Number matched in second data set: 4"
# [1] 4 7
# [1] "DMR432_Hypo_P5_10_H_dist500"
# [1] "Number matched in first data set: 9"
# [1] "Number matched in second data set: 10"
# [1] 10  7
# [1] "DMR432_Hypo_P45_H3K_dist500"
# [1] "Number matched in first data set: 4"
# [1] "Number matched in second data set: 4"
# [1] 4 7
# [1] "DMR432_Hypo_others__dist500"
# [1] "Number matched in first data set: 20"
# [1] "Number matched in second data set: 21"
# [1] 21  7
```

Then, files were manually renamed and folders were reorganized.

- In /mnt/h/methylome/fusion/post\_methylkit/DMR432\_pval0.07/capture\_separee/with\_merge\_file/DMR432\_All/dist500, are the files:

  - DMR432\_all\_ATAC\_capt\_dist500.txt
  - DMR432\_all\_ChIP\_capt\_dist500.txt
  - DMR432\_all\_H3K27\_capt\_dist500.txt
  - DMR432\_all\_others\_capt\_dist500.txt
  - DMR432\_all\_P1\_5\_10\_H3K4me1\_capt\_dist500.txt
  - DMR432\_all\_P45\_H3K4me1\_capt\_dist500.txt
  - DMR432\_all\_P5\_10\_H3K4me1\_capt\_dist500.txt
  - DMR432\_all\_prom\_capt\_dist500.txt
- In /mnt/h/methylome/fusion/post\_methylkit/DMR432\_pval0.07/capture\_separee/with\_merge\_file/DMR432\_Hyper/dist500:

  - DMR432\_Hyper\_ATAC\_capt\_dist500.txt
  - DMR432\_Hyper\_ChIP\_capt\_dist500.txt
  - DMR432\_Hyper\_H3K27\_capt\_dist500.txt
  - DMR432\_Hyper\_others\_capt\_dist500.txt
  - DMR432\_Hyper\_P1\_5\_10\_H3K4me1\_capt\_dist500.txt
  - DMR432\_Hyper\_P45\_H3K4me1\_capt\_dist500.txt
  - DMR432\_Hyper\_P5\_10\_H3K4me1\_capt\_dist500.txt
  - DMR432\_Hyper\_prom\_capt\_dist500.txt
- In /mnt/h/methylome/fusion/post\_methylkit/DMR432\_pval0.07/capture\_separee/with\_merge\_file/DMR432\_Hypo/dist500:

  - DMR432\_Hypo\_ATAC\_capt\_dist500.txt
  - DMR432\_Hypo\_ChIP\_capt\_dist500.txt
  - DMR432\_Hypo\_H3K27\_capt\_dist500.txt
  - DMR432\_Hypo\_others\_capt\_dist500.txt
  - DMR432\_Hypo\_P1\_5\_10\_H3K4me1\_capt\_dist500.txt
  - DMR432\_Hypo\_P45\_H3K4me1\_capt\_dist500.txt
  - DMR432\_Hypo\_P5\_10\_H3K4me1\_capt\_dist500.txt
  - DMR432\_Hypo\_prom\_capt\_dist500.txt

##### 21.3 Enrichment of each capture category¶

To determine whether methylation defects were randomly distributed among captured regions or not, we performed **hypergeometric tests** using **phyper** function of R, either on all DMRs, or by separating hypo-DMRs and hyper-DMRs.

In [ ]:

```
%%R

# -------------------------------------------------------------
# Hypergeometric tests
# to identify potential enrichment for a given capture element
# -------------------------------------------------------------

setwd("H:/methylome/fusion/post_methylkit/DMR432_pval0.07/capture_separee/with_merge_file")

DMR432_capture = read.table("DMR432_capt_merge_sep.txt", sep="\t", row.names=1, header=TRUE)

colnames(DMR432_capture)=c("nb_tot_capt", "all_large", "hyper_large", "hypo_large")

DMR432_capture
#                                   nb_tot_capt all_large hyper_large hypo_large
# promoteur                               23769      135       55          80
# H3K27ac                                 34279      349       225         124
# ChIP HSF2                                2111      3         0           3
# ATACseq                                   773      6         3           3
# P1_5_10_H3K4me1                          2316      6         2           4
# P5_10_H3K4me1                            6248      17        7           10
# P45_H3K4me1                              1128      7         3           4
# others_all_except_prom_et_H3K27ac       12108      36        15          21


# Preparation of the results table
treatment_file_low = matrix(NA, ncol=4, nrow=nrow(DMR432_capture))
treatment_file_low[,1] = DMR432_capture[,1]
rownames(treatment_file_low) = rownames(DMR432_capture)
colnames(treatment_file_low) = colnames(DMR432_capture)
treatment_file_low

treatment_file_up = matrix(NA, ncol=4, nrow=nrow(DMR432_capture))
treatment_file_up[,1] = DMR432_capture[,1]
rownames(treatment_file_up) = rownames(DMR432_capture)
colnames(treatment_file_up) = colnames(DMR432_capture)
treatment_file_up

n_capt_tot = 58611 #number of regions in the capture (enlarged and merged capture regions)

for (colonne in 2:4) {
   for (ligne in 1:nrow(DMR432_capture)) {
     k = DMR432_capture[ligne,colonne] #nb of DMR that are into (or close to) a region of captured element we studied 
     l = DMR432_capture[ligne,1] #nb of captured regions in the captured element considered
     
     if (colonne == 2) {
       n_DMR = 432 # number of DMR in DMR432 (all DMRs)
     } else if (colonne == 3){
       n_DMR = 257 # number of Hyper DMRs among DMR432
     } else if (colonne == 4){
       n_DMR = 175 # number of Hypo DMRs among DMR432
     }
     
     value_low = phyper(k-1, n_DMR, n_capt_tot - n_DMR, l, lower.tail = FALSE)
     value_up = phyper(k, n_DMR, n_capt_tot - n_DMR, l, lower.tail = TRUE)
     
     print(paste("Pour :", colnames(DMR432_capture)[colonne], "-", rownames(DMR432_capture)[ligne], "k vaut : ", k, "et l vaut : ", l))
     print(paste("valeur low : ", value_low, "- valeur up : ", value_up))
     
     treatment_file_low[ligne,colonne]= value_low
     treatment_file_up[ligne,colonne]= value_up
   }
 }


head(treatment_file_low)
#                 nb_tot_capt   all_large  hyper_large    hypo_large
# promoteur             23769 9.999757e-01 1.000000e+00  0.0946983013
# H3K27ac               34279 3.384778e-23 1.591491e-24  0.0004681597
# ChIP HSF2              2111 9.999816e-01 1.000000e+00  0.9533009556
# ATACseq                 773 5.058000e-01 6.606032e-01  0.4067426552
# P1_5_10_H3K4me1        2316 9.994818e-01 9.996408e-01  0.9182647152
# P5_10_H3K4me1          6248 9.999999e-01 9.999997e-01  0.9922111214

head(treatment_file_up) 
#                 nb_tot_capt      all_large   hyper_large   hypo_large
# promoteur             23769 3.742334e-05 5.335081e-11 0.92852096
# H3K27ac               34279 1.000000e+00 1.000000e+00 0.99973984
# ChIP HSF2              2111 1.063351e-04 7.882790e-05 0.12118705
# ATACseq                 773 6.552772e-01 5.599107e-01 0.79887453
# P1_5_10_H3K4me1        2316 1.641787e-03 2.094954e-03 0.17494295
# P5_10_H3K4me1          6248 2.579597e-07 1.516861e-06 0.01679286

write.table(treatment_file_low, "hypergeom_test_low.txt", sep="\t", row.names = TRUE, col.names = TRUE, quote=FALSE)

write.table(treatment_file_up, "hypergeom_test_up.txt", sep="\t", row.names = TRUE, col.names = TRUE, quote=FALSE)
```

**Conclusion**
Genomic regions were not randomly altered by prenatal alcohol exposure in their DNA methylation status:

- brain active enhancers, characterized by the H3K27ac histone mark, are significantly over-represented among the DMRs identified in the capture, (Fig. 2B, Fig. 3 and Table 2 of the paper in preparation, Chapter 4, section 1.2).
- In contrast, promoters and other regions of interest are not significantly affected by the binge drinking stress (Fig. 2A and 2B of the paper in preparation, Chapter 4, section 1.2).

### 22.Motifs enrichment¶

Transcription factor binding site enrichment within the DMRs located in active enhancer regions (H3K27ac mark), was explored using **findMotifsGenome** (**HOMER** suite). This tool allows the identification of enrichment in known (associated with a transcription factor) or unknown patterns among the DMRs.
  
  
In this analysis, DMRs are located into H3K27ac enhancer regions, which are genomic regions that does not necessarily represent the whole-genome. Thus, background considered in this analysis is restricted to all H3K27ac enhancers that are in the capture.
  
  
In order to estimate the threshold value at which the pattern search are unreliable, this pattern search was also performed on a "random" dataset, gathering randomly generated regions, but sharing the same characteristics as the real dataset (i.e. as many regions and identical region size).

In [ ]:

```
%%bash 

#-----------------------------------------------
# A. Motifs enrichment (among DMR that are in - or close to - H3K27ac enhancers)
#-----------------------------------------------

# a. Motifs enrichment in real dataset
# a1. Extraction of DMRs (bed coordinates) that are in - or close to - H3K27ac enhancers  
cd /mnt/h/methylome/fusion/post_methylkit/DMR432_pval0.07/capture_separee/with_merge_file/DMR432_All/dist500
sed '1d' DMR432_all_H3K27_capt_dist500.txt | awk '{ print $1"\t"$2"\t"$3 }' | sort -k1,1 -k2,2n | uniq > DMR432_all_H3K27_dist500.bed

cd /mnt/h/methylome/fusion/post_methylkit/DMR432_pval0.07/capture_separee/with_merge_file/DMR432_Hyper/dist500/
sed '1d' DMR432_Hyper_H3K27_capt_dist500.txt | awk '{ print $1"\t"$2"\t"$3 }' | sort -k1,1 -k2,2n | uniq > DMR432_hyper_H3K27_dist500.bed

cd /mnt/h/methylome/fusion/post_methylkit/DMR432_pval0.07/capture_separee/with_merge_file/DMR432_Hypo/dist500/
sed '1d' DMR432_Hypo_H3K27_capt_dist500.txt | awk '{ print $1"\t"$2"\t"$3 }' | sort -k1,1 -k2,2n | uniq > DMR432_hypo_H3K27_dist500.bed

mkdir -p /mnt/h/methylome/fusion/post_methylkit/DMR432_pval0.07/homer/H3K27ac/all/real/with_bg
mkdir -p /mnt/h/methylome/fusion/post_methylkit/DMR432_pval0.07/homer/H3K27ac/hyper/real/with_bg
mkdir -p /mnt/h/methylome/fusion/post_methylkit/DMR432_pval0.07/homer/H3K27ac/hypo/real/with_bg

conda activate chip 

# enhancers real All
findMotifsGenome.pl /mnt/h/methylome/fusion/post_methylkit/DMR432_pval0.07/capture_separee/with_merge_file/DMR432_All/dist500/DMR432_all_H3K27_dist500.bed mm9 /mnt/h/methylome/fusion/post_methylkit/DMR432_pval0.07/homer/H3K27ac/all/real -size given -mset vertebrates -bg /mnt/h/methylome/capture/redo/agrandi/ok/in_big_capt_H3K27Ac.txt

# enhancers real Hyper
findMotifsGenome.pl /mnt/h/methylome/fusion/post_methylkit/DMR432_pval0.07/capture_separee/with_merge_file/DMR432_Hyper/dist500/DMR432_hyper_H3K27_dist500.bed mm9 /mnt/h/methylome/fusion/post_methylkit/DMR432_pval0.07/homer/H3K27ac/hyper/real/with_bg -size given -mset vertebrates -bg /mnt/h/methylome/capture/redo/agrandi/ok/in_big_capt_H3K27Ac.txt

# enhancers real Hypo
findMotifsGenome.pl /mnt/h/methylome/fusion/post_methylkit/DMR432_pval0.07/capture_separee/with_merge_file/DMR432_Hypo/dist500/DMR432_hypo_H3K27_dist500.bed mm9 /mnt/h/methylome/fusion/post_methylkit/DMR432_pval0.07/homer/H3K27ac/hypo/real/with_bg -size given -mset vertebrates -bg /mnt/h/methylome/capture/redo/agrandi/ok/in_big_capt_H3K27Ac.txt
```

In [ ]:

```
%%R

# b. Motifs enrichment in random dataset 
# b.1 To create a random dataset
enh_all = read.table("H:/methylome/fusion/post_methylkit/DMR432_pval0.07/capture_separee/with_merge_file/DMR432_All/dist500/DMR432_all_H3K27_dist500.bed", sep = "\t", header=FALSE)

enh_hyper = read.table("H:/methylome/fusion/post_methylkit/DMR432_pval0.07/capture_separee/with_merge_file/DMR432_Hyper/dist500/DMR432_hyper_H3K27_dist500.bed", sep = "\t", header=FALSE)

enh_hypo = read.table("H:/methylome/fusion/post_methylkit/DMR432_pval0.07/capture_separee/with_merge_file/DMR432_Hypo/dist500/DMR432_hypo_H3K27_dist500.bed", sep = "\t", header=FALSE)

head(enh_all)
head(enh_hyper)
head(enh_hypo)

dim(enh_all)
dim(enh_hyper)
dim(enh_hypo)

# To load all captured H3K27ac enhancer file 
enh_capt = read.table("H:/methylome/capture/redo/agrandi/ok/in_big_capt_H3K27Ac.txt", sep="\t", header=FALSE)

head(enh_capt)
dim(enh_capt)


# Function to create random dataset, having same number of regions than real dataset, with regions of similar length
randomisation= function(treatment_file, file_name, background, folder){
  colnames(treatment_file) = c("chr","start","end")
  treatment_file$start = as.numeric(as.character(treatment_file$start))
  treatment_file$end = as.numeric(as.character(treatment_file$end))
    # To determine DMR length 
  treatment_file$length = with(treatment_file, end-start)
  print(file_name)
  print(head(treatment_file))
  
  # To select same number of regions than in the real dataset
  random_dataset = background[sample(1:nrow(background), size=nrow(treatment_file)),]
  
  # to determine median coordinate of randomly selected regions
  random_dataset$middle = with(random_dataset,V2+((V3-V2)/2))
  
  pool.real.random=cbind(treatment_file, random_dataset)
  
  # To create regions of same length than those in real dataset
  pool.real.random$end_random = with(pool.real.random,floor(middle+(length/2)))
  pool.real.random$start_random = with(pool.real.random,ceiling(middle-(length/2)))
  
  # To create.bed file for randomly selected regions
  bed_random_dataset = pool.real.random[,c("V1","start_random","end_random")]
  
  write.table(bed_random_dataset, file=paste(folder,"random_",file_name,".bed",sep=""), sep = "\t", quote = FALSE, dec = ".", row.names=FALSE, col.names=TRUE)
}


randomisation(treatment_file = enh_all , file_name = "DMR432_enh_all_dist500", 
              background = enh_capt, 
              folder= "H:/methylome/fusion/post_methylkit/DMR432_pval0.07/random_data/homer_analysis/")

randomisation(treatment_file = enh_hyper , file_name = "DMR432_enh_hyper_dist500", 
              background = enh_capt,  
              folder= "H:/methylome/fusion/post_methylkit/DMR432_pval0.07/random_data/homer_analysis/")

randomisation(treatment_file = enh_hypo , file_name = "DMR432_enh_hypo_dist500", 
              background = enh_capt,  
              folder= "H:/methylome/fusion/post_methylkit/DMR432_pval0.07/random_data/homer_analysis/")
```

In [ ]:

```
%%bash

# b.2 To sort random files
cd /mnt/h/methylome/fusion/post_methylkit/DMR432_pval0.07/random_data/homer_analysis/

sample="enh_all enh_hyper enh_hypo"
for file in $sample
do 
	echo $file
	sort -k1,1 -k2,2n random_DMR432_${file}_dist500.bed > random_DMR432_${file}_dist500_sorted.bed
done


# b.3 To search motif enrichment (HOMER) - random dataset
mkdir -p /mnt/h/methylome/fusion/post_methylkit/DMR432_pval0.07/homer/H3K27ac/all/random/with_bg
mkdir -p /mnt/h/methylome/fusion/post_methylkit/DMR432_pval0.07/homer/H3K27ac/hyper/random/with_bg
mkdir -p /mnt/h/methylome/fusion/post_methylkit/DMR432_pval0.07/homer/H3K27ac/hypo/random/with_bg

conda activate chip 

#enhancers random All :
findMotifsGenome.pl /mnt/h/methylome/fusion/post_methylkit/DMR432_pval0.07/random_data/homer_analysis/random_DMR432_enh_all_dist500_sorted.bed mm9 /mnt/h/methylome/fusion/post_methylkit/DMR432_pval0.07/homer/H3K27ac/all/random/with_bg -size given -mset vertebrates -bg /mnt/h/methylome/capture/redo/agrandi/ok/in_big_capt_H3K27Ac.txt

#enhancers random Hyper :
findMotifsGenome.pl /mnt/h/methylome/fusion/post_methylkit/DMR432_pval0.07/random_data/homer_analysis/random_DMR432_enh_hyper_dist500_sorted.bed mm9 /mnt/h/methylome/fusion/post_methylkit/DMR432_pval0.07/homer/H3K27ac/hyper/random/with_bg -size given -mset vertebrates -bg /mnt/h/methylome/capture/redo/agrandi/ok/in_big_capt_H3K27Ac.txt

#enhancers random Hypo :
findMotifsGenome.pl /mnt/h/methylome/fusion/post_methylkit/DMR432_pval0.07/random_data/homer_analysis/random_DMR432_enh_hypo_dist500_sorted.bed mm9 /mnt/h/methylome/fusion/post_methylkit/DMR432_pval0.07/homer/H3K27ac/hypo/random/with_bg -size given -mset vertebrates -bg /mnt/h/methylome/capture/redo/agrandi/ok/in_big_capt_H3K27Ac.txt
```

#### 23. Combination of DMR432 informations: annotation, statistical, CGi overlap and capture repartition¶

All data (*i.e.* statistical informations, mm9 and mm10 coordinates regions, annotation informations, CGi overlap, capture repartition) were combined into a single file.

In [ ]:

```
%%R 

setwd("H:/methylome/fusion/post_methylkit/DMR432_pval0.07/annotations/Biomart/mm10")

# To load DMR432 global informations file
DMR432 = read.table("DMR432_unique_annot_mm10_BMT_meth_cgi.bed", 
                    header = FALSE, fill=TRUE, quote="", sep="\t", na.strings = "")
head(DMR432)
dim(DMR432)

# To load files containing DMR in each captured element type.
ATAC = read.table("H:/methylome/fusion/post_methylkit/DMR432_pval0.07/capture_separee/with_merge_file/DMR432_All/dist500/DMR432_all_ATAC_capt_dist500.txt", 
                  sep = "\t", header=TRUE)
head(ATAC)
dim(ATAC)

prom = read.table("H:/methylome/fusion/post_methylkit/DMR432_pval0.07/capture_separee/with_merge_file/DMR432_All/dist500/DMR432_all_prom_capt_dist500.txt", 
                  sep = "\t", header=TRUE)
head(prom)
dim(prom)

ChIP = read.table("H:/methylome/fusion/post_methylkit/DMR432_pval0.07/capture_separee/with_merge_file/DMR432_All/dist500/DMR432_all_ChIP_capt_dist500.txt", 
                  sep = "\t", header=TRUE)
head(ChIP)
dim(ChIP)

H3K27 = read.table("H:/methylome/fusion/post_methylkit/DMR432_pval0.07/capture_separee/with_merge_file/DMR432_All/dist500/DMR432_all_H3K27_capt_dist500.txt", 
                   sep = "\t", header=TRUE)
head(H3K27)
dim(H3K27)

others = read.table("H:/methylome/fusion/post_methylkit/DMR432_pval0.07/capture_separee/with_merge_file/DMR432_All/dist500/DMR432_all_others_capt_dist500.txt",
                    sep = "\t", header=TRUE)
head(others)
dim(others)

P1_P5_P10 = read.table("H:/methylome/fusion/post_methylkit/DMR432_pval0.07/capture_separee/with_merge_file/DMR432_All/dist500/DMR432_all_P1_5_10_H3K4me1_capt_dist500.txt", 
                       sep = "\t", header=TRUE)
head(P1_P5_P10)
dim(P1_P5_P10)

P5_P10 = read.table("H:/methylome/fusion/post_methylkit/DMR432_pval0.07/capture_separee/with_merge_file/DMR432_All/dist500/DMR432_all_P5_10_H3K4me1_capt_dist500.txt", 
                    sep = "\t", header=TRUE)
head(P5_P10)
dim(P5_P10)

P45 = read.table("H:/methylome/fusion/post_methylkit/DMR432_pval0.07/capture_separee/with_merge_file/DMR432_All/dist500/DMR432_all_P45_H3K4me1_capt_dist500.txt", 
                 sep = "\t", header=TRUE)
head(P45)
dim(P45)

# To create columns that will be fill with informations 
DMR432$promoters=NA #yes/no
DMR432$promoters_ID= NA #chr;start;end
DMR432$promoters_overlap=NA #overlap-size

DMR432$H3K27ac=NA #yes/no
DMR432$H3K27ac_ID= NA #chr;start;end
DMR432$H3K27ac_overlap=NA #overlap-size

DMR432$others=NA #yes/no
DMR432$others_ID= NA #chr;start;end
DMR432$others_overlap=NA #overlap-size

DMR432$ATAC=NA #yes/no
DMR432$ATAC_ID= NA #chr;start;end
DMR432$ATAC_overlap=NA #overlap-size

DMR432$ChIP=NA #yes/no
DMR432$ChIP_ID= NA #chr;start;end
DMR432$ChIP_overlap=NA #overlap-size

DMR432$P1_5_10=NA #yes/no
DMR432$P1_5_10_ID= NA #chr;start;end
DMR432$P1_5_10_overlap=NA #overlap-size

DMR432$P5_10=NA #yes/no
DMR432$P5_10_ID= NA #chr;start;end
DMR432$P5_10_overlap=NA #overlap-size

DMR432$P45=NA #yes/no
DMR432$P45_ID= NA #chr;start;end
DMR432$P45_overlap=NA #overlap-size


#---------------------------
# For promoters informations
#---------------------------

# To create the ID (chr;start;end) of DMR432 global information file
DMR432$mm10_ID = with(DMR432, paste(V15,V16,V17,sep=";"))

# To complete promoters informations 
treatment_file = prom #data
column = "promoters"

# To create the ID (chr;start;end)
treatment_file$ID = with(treatment_file, paste(chr_reg1,start_reg1,end_reg1,sep=";"))

# Is the DMR in the captured element type?
for (i in 1:nrow(DMR432)){
  DMR432_ID <- DMR432$mm10_ID[i]
  for(j in 1:nrow(treatment_file)){
    if(DMR432_ID == treatment_file$ID[j]){
      DMR432[i,column]= "yes"
      
      # If case contains "NA" value:
      if (is.na(DMR432[i,paste(column,"ID",sep="_")])==TRUE){
        DMR432[i,paste(column,"ID",sep="_")]=paste(treatment_file$chr_reg2[j],
                                                   treatment_file$start_reg2[j],
                                                   treatment_file$end_reg2[j],sep=";") 
        DMR432[i,paste(column,"overlap",sep="_")]=treatment_file$overlap_size[j]
      }else{
        #If another element type was already found : keep old and new element type 
        DMR432[i,paste(column,"ID",sep="_")]=paste(DMR432[i,paste(column,"ID",sep="_")],
                                                   paste(treatment_file$chr_reg2[j],
                                                         treatment_file$start_reg2[j],
                                                         treatment_file$end_reg2[j],
                                                         sep=";"), sep="_") 
        DMR432[i,paste(column,"overlap",sep="_")]=paste(DMR432[i,paste(column,"overlap",sep="_")],
                                                        treatment_file$overlap_size[j], sep=";")
      }
    }
  }
}

DMR432[is.na(DMR432[,column]),column] = "no"
DMR432[is.na(DMR432[,paste(column,"ID",sep="_")]),paste(column,"ID",sep="_")] = "."
DMR432[is.na(DMR432[,paste(column,"overlap",sep="_")]),paste(column,"overlap",sep="_")] = "."

View(DMR432)

#-----------------------------------
# For H3K27ac enhancers informations
#-----------------------------------
treatment_file = H3K27 #data
column = "H3K27ac"

# To create the ID (chr;start;end)
treatment_file$ID = with(treatment_file, paste(chr_reg1,start_reg1,end_reg1,sep=";"))

# Is the DMR in the captured element type?
for (i in 1:nrow(DMR432)){
  DMR432_ID <- DMR432$mm10_ID[i]
  for(j in 1:nrow(treatment_file)){
    if(DMR432_ID == treatment_file$ID[j]){
      DMR432[i,column]= "yes"
      
      # If case contains "NA" value:
      if (is.na(DMR432[i,paste(column,"ID",sep="_")])==TRUE){
        DMR432[i,paste(column,"ID",sep="_")]=paste(treatment_file$chr_reg2[j],
                                                   treatment_file$start_reg2[j],
                                                   treatment_file$end_reg2[j],
                                                   sep=";") 
        DMR432[i,paste(column,"overlap",sep="_")]=treatment_file$overlap_size[j]
      }else{
        #If another element type was already found : keep old and new element type 
        DMR432[i,paste(column,"ID",sep="_")]=paste(DMR432[i,paste(column,"ID",sep="_")],
                                                   paste(treatment_file$chr_reg2[j],
                                                         treatment_file$start_reg2[j],
                                                         treatment_file$end_reg2[j],sep=";"), sep="_") 
        DMR432[i,paste(column,"overlap",sep="_")]=paste(DMR432[i,paste(column,"overlap",sep="_")],
                                                        treatment_file$overlap_size[j], sep=";")
      }
    }
  }
}

DMR432[is.na(DMR432[,column]),column] = "no"
DMR432[is.na(DMR432[,paste(column,"ID",sep="_")]),paste(column,"ID",sep="_")] = "."
DMR432[is.na(DMR432[,paste(column,"overlap",sep="_")]),paste(column,"overlap",sep="_")] = "."


# For other group informations
treatment_file = others #data
column = "others"

# To create the ID (chr;start;end)
treatment_file$ID = with(treatment_file, paste(chr_reg1,start_reg1,end_reg1,sep=";"))

# Is the DMR in the captured element type?
for (i in 1:nrow(DMR432)){
  DMR432_ID <- DMR432$mm10_ID[i]
  for(j in 1:nrow(treatment_file)){
    if(DMR432_ID == treatment_file$ID[j]){
      DMR432[i,column]= "yes"
      
      # If case contains "NA" value:
      if (is.na(DMR432[i,paste(column,"ID",sep="_")])==TRUE){
        DMR432[i,paste(column,"ID",sep="_")]=paste(treatment_file$chr_reg2[j],
                                                   treatment_file$start_reg2[j],
                                                   treatment_file$end_reg2[j],sep=";") 
        DMR432[i,paste(column,"overlap",sep="_")]=treatment_file$overlap_size[j]
      }else{
        #If another element type was already found : keep old and new element type 
        DMR432[i,paste(column,"ID",sep="_")]=paste(DMR432[i,paste(column,"ID",sep="_")],
                                                   paste(treatment_file$chr_reg2[j],
                                                         treatment_file$start_reg2[j],
                                                         treatment_file$end_reg2[j],sep=";"), 
                                                   sep="_") 
        DMR432[i,paste(column,"overlap",sep="_")]=paste(DMR432[i,paste(column,"overlap",sep="_")],
                                                        treatment_file$overlap_size[j], sep=";")
      }
    }
  }
}

DMR432[is.na(DMR432[,column]),column] = "no"
DMR432[is.na(DMR432[,paste(column,"ID",sep="_")]),paste(column,"ID",sep="_")] = "."
DMR432[is.na(DMR432[,paste(column,"overlap",sep="_")]),paste(column,"overlap",sep="_")] = "."


#----------------------
# For ChIP informations
#----------------------
treatment_file = ChIP #data
column = "ChIP"

# To create the ID (chr;start;end)
treatment_file$ID = with(treatment_file, paste(chr_reg1,start_reg1,end_reg1,sep=";"))

# Is the DMR in the captured element type?
for (i in 1:nrow(DMR432)){
  DMR432_ID <- DMR432$mm10_ID[i]
  for(j in 1:nrow(treatment_file)){
    if(DMR432_ID == treatment_file$ID[j]){
      DMR432[i,column]= "yes"
      
      # If case contains "NA" value:
      if (is.na(DMR432[i,paste(column,"ID",sep="_")])==TRUE){
        DMR432[i,paste(column,"ID",sep="_")]=paste(treatment_file$chr_reg2[j],
                                                   treatment_file$start_reg2[j],
                                                   treatment_file$end_reg2[j],sep=";") 
        DMR432[i,paste(column,"overlap",sep="_")]=treatment_file$overlap_size[j]
      }else{
        #If another element type was already found : keep old and new element type 
        DMR432[i,paste(column,"ID",sep="_")]=paste(DMR432[i,paste(column,"ID",sep="_")],
                                                   paste(treatment_file$chr_reg2[j],
                                                         treatment_file$start_reg2[j],
                                                         treatment_file$end_reg2[j],sep=";"), 
                                                   sep="_") 
        DMR432[i,paste(column,"overlap",sep="_")]=paste(DMR432[i,paste(column,"overlap",sep="_")],
                                                        treatment_file$overlap_size[j], sep=";")
      }
    }
  }
}

DMR432[is.na(DMR432[,column]),column] = "no"
DMR432[is.na(DMR432[,paste(column,"ID",sep="_")]),paste(column,"ID",sep="_")] = "."
DMR432[is.na(DMR432[,paste(column,"overlap",sep="_")]),paste(column,"overlap",sep="_")] = "."

#----------------------
# For Atac-seq informations
#----------------------
treatment_file = ATAC #data
column = "ATAC"

# To create the ID (chr;start;end)
treatment_file$ID = with(treatment_file, paste(chr_reg1,start_reg1,end_reg1,sep=";"))
# Is the DMR in the captured element type?
for (i in 1:nrow(DMR432)){
  DMR432_ID <- DMR432$mm10_ID[i]
  for(j in 1:nrow(treatment_file)){
    if(DMR432_ID == treatment_file$ID[j]){
      DMR432[i,column]= "yes"
      
      # If case contains "NA" value:
      if (is.na(DMR432[i,paste(column,"ID",sep="_")])==TRUE){
        DMR432[i,paste(column,"ID",sep="_")]=paste(treatment_file$chr_reg2[j],
                                                   treatment_file$start_reg2[j],
                                                   treatment_file$end_reg2[j],sep=";") 
        DMR432[i,paste(column,"overlap",sep="_")]=treatment_file$overlap_size[j]
      }else{
        #If another element type was already found : keep old and new element type 
        DMR432[i,paste(column,"ID",sep="_")]=paste(DMR432[i,paste(column,"ID",sep="_")],
                                                   paste(treatment_file$chr_reg2[j],
                                                         treatment_file$start_reg2[j],
                                                         treatment_file$end_reg2[j],sep=";"), 
                                                   sep="_") 
        DMR432[i,paste(column,"overlap",sep="_")]=paste(DMR432[i,paste(column,"overlap",sep="_")],
                                                        treatment_file$overlap_size[j], sep=";")
      }
    }
  }
}

DMR432[is.na(DMR432[,column]),column] = "no"
DMR432[is.na(DMR432[,paste(column,"ID",sep="_")]),paste(column,"ID",sep="_")] = "."
DMR432[is.na(DMR432[,paste(column,"overlap",sep="_")]),paste(column,"overlap",sep="_")] = "."


#----------------------
# For P1_P5_P10 informations
#----------------------
treatment_file = P1_P5_P10 #data
column = "P1_5_10"

# To create the ID (chr;start;end)
treatment_file$ID = with(treatment_file, paste(chr_reg1,start_reg1,end_reg1,sep=";"))

# Is the DMR in the captured element type?
for (i in 1:nrow(DMR432)){
  DMR432_ID <- DMR432$mm10_ID[i]
  for(j in 1:nrow(treatment_file)){
    if(DMR432_ID == treatment_file$ID[j]){
      DMR432[i,column]= "yes"
      
      # If case contains "NA" value:
      if (is.na(DMR432[i,paste(column,"ID",sep="_")])==TRUE){
        DMR432[i,paste(column,"ID",sep="_")]=paste(treatment_file$chr_reg2[j],
                                                   treatment_file$start_reg2[j],
                                                   treatment_file$end_reg2[j],sep=";") 
        DMR432[i,paste(column,"overlap",sep="_")]=treatment_file$overlap_size[j]
      }else{
        #If another element type was already found : keep old and new element type 
        DMR432[i,paste(column,"ID",sep="_")]=paste(DMR432[i,paste(column,"ID",sep="_")],
                                                   paste(treatment_file$chr_reg2[j],
                                                         treatment_file$start_reg2[j],
                                                         treatment_file$end_reg2[j],sep=";"), 
                                                   sep="_") 
        DMR432[i,paste(column,"overlap",sep="_")]=paste(DMR432[i,paste(column,"overlap",sep="_")],
                                                        treatment_file$overlap_size[j], sep=";")
      }
    }
  }
}

DMR432[is.na(DMR432[,column]),column] = "no"
DMR432[is.na(DMR432[,paste(column,"ID",sep="_")]),paste(column,"ID",sep="_")] = "."
DMR432[is.na(DMR432[,paste(column,"overlap",sep="_")]),paste(column,"overlap",sep="_")] = "."

#----------------------
# For P5_P10 informations
#----------------------
treatment_file = P5_P10 #data
column = "P5_10"

# To create the ID (chr;start;end)
treatment_file$ID = with(treatment_file, paste(chr_reg1,start_reg1,end_reg1,sep=";"))

# Is the DMR in the captured element type?
for (i in 1:nrow(DMR432)){
  DMR432_ID <- DMR432$mm10_ID[i]
  for(j in 1:nrow(treatment_file)){
    if(DMR432_ID == treatment_file$ID[j]){
      DMR432[i,column]= "yes"
      
      # If case contains "NA" value:
      if (is.na(DMR432[i,paste(column,"ID",sep="_")])==TRUE){
        DMR432[i,paste(column,"ID",sep="_")]=paste(treatment_file$chr_reg2[j],
                                                   treatment_file$start_reg2[j],
                                                   treatment_file$end_reg2[j],sep=";") 
        DMR432[i,paste(column,"overlap",sep="_")]=treatment_file$overlap_size[j]
      }else{
        #If another element type was already found : keep old and new element type 
        DMR432[i,paste(column,"ID",sep="_")]=paste(DMR432[i,paste(column,"ID",sep="_")],
                                                   paste(treatment_file$chr_reg2[j],
                                                         treatment_file$start_reg2[j],
                                                         treatment_file$end_reg2[j],sep=";"), 
                                                   sep="_") 
        DMR432[i,paste(column,"overlap",sep="_")]=paste(DMR432[i,paste(column,"overlap",sep="_")],
                                                        treatment_file$overlap_size[j], sep=";")
      }
    }
  }
}

DMR432[is.na(DMR432[,column]),column] = "no"
DMR432[is.na(DMR432[,paste(column,"ID",sep="_")]),paste(column,"ID",sep="_")] = "."
DMR432[is.na(DMR432[,paste(column,"overlap",sep="_")]),paste(column,"overlap",sep="_")] = "."

#----------------------
# For P45 informations
#----------------------
treatment_file = P45 #data
column = "P45"

# To create the ID (chr;start;end)
treatment_file$ID = with(treatment_file, paste(chr_reg1,start_reg1,end_reg1,sep=";"))

# Is the DMR in the captured element type?
for (i in 1:nrow(DMR432)){
  DMR432_ID <- DMR432$mm10_ID[i]
  for(j in 1:nrow(treatment_file)){
    if(DMR432_ID == treatment_file$ID[j]){
      DMR432[i,column]= "yes"
      
      # If case contains "NA" value:
      if (is.na(DMR432[i,paste(column,"ID",sep="_")])==TRUE){
        DMR432[i,paste(column,"ID",sep="_")]=paste(treatment_file$chr_reg2[j],
                                                   treatment_file$start_reg2[j],
                                                   treatment_file$end_reg2[j],sep=";") 
        DMR432[i,paste(column,"overlap",sep="_")]=treatment_file$overlap_size[j]
      }else{
        #If another element type was already found : keep old and new element type 
        DMR432[i,paste(column,"ID",sep="_")]=paste(DMR432[i,paste(column,"ID",sep="_")],
                                                   paste(treatment_file$chr_reg2[j],
                                                         treatment_file$start_reg2[j],
                                                         treatment_file$end_reg2[j],sep=";"), 
                                                   sep="_") 
        DMR432[i,paste(column,"overlap",sep="_")]=paste(DMR432[i,paste(column,"overlap",sep="_")],
                                                        treatment_file$overlap_size[j], sep=";")
      }
    }
  }
}

DMR432[is.na(DMR432[,column]),column] = "no"
DMR432[is.na(DMR432[,paste(column,"ID",sep="_")]),paste(column,"ID",sep="_")] = "."
DMR432[is.na(DMR432[,paste(column,"overlap",sep="_")]),paste(column,"overlap",sep="_")] = "."

View(DMR432)

colnames(DMR432)=c("DMR_mm10_chr","DMR_mm10_start","DMR_mm10_end",
                   "DMR_mm10_ID", "DMR_annot_chr","DMR_annot_start",
                   "DMR_annot_end","strand","entrezgene_accession",
                   "gene_biotype","mgi_symbol", "ENSEMBL_ID", 
                   "Entrez_ID", "annot_overlap_length", "DMR_mm9_chr", 
                   "DMR_mm9_start", "DMR_mm9_end","DMR_mm10_chr",
                   "DMR_mm10_start","DMR_mm10_end", "CpG_nb", "pvalue", 
                   "qvalue",  "meth.diff", "CGi_chr","CGi_start",
                   "CGi_end","CGi_name","CGi_length","CGi_CG_nb",  
                   "CGI_overlap_length","promoters","promoters_ID",
                   "promoters_overlap","H3K27ac","H3K27ac_ID",
                   "H3K27ac_overlap","others","others_ID","others_overlap",
                   "ATAC","ATAC_ID","ATAC_overlap","HSF2_naive_ChIP",
                   "HSF2_naive_ChIP_ID","HSF2_ChIP_overlap",
                   "P1_5_10","P1_5_10_ID","P1_5_10_overlap",
                   "P5_10","P5_10_ID","P5_10_overlap","P45","P45_ID",
                   "P45_overlap","DMR_mm10_ID")

DMR432_clean = DMR432[,c("DMR_mm10_chr","DMR_mm10_start","DMR_mm10_end",
                         "DMR_mm10_ID", "DMR_annot_chr","DMR_annot_start",
                         "DMR_annot_end","strand", "entrezgene_accession",
                         "gene_biotype","mgi_symbol", "ENSEMBL_ID", "Entrez_ID", 
                         "annot_overlap_length", "DMR_mm9_chr", 
                         "DMR_mm9_start", "DMR_mm9_end","CpG_nb", "pvalue", 
                         "qvalue","meth.diff","CGi_chr","CGi_start","CGi_end",
                         "CGi_name","CGi_length","CGi_CG_nb", "CGI_overlap_length",
                         "promoters","promoters_ID","promoters_overlap","H3K27ac",
                         "H3K27ac_ID","H3K27ac_overlap","others","others_ID",
                         "others_overlap","ATAC","ATAC_ID","ATAC_overlap",
                         "HSF2_naive_ChIP","HSF2_naive_ChIP_ID","HSF2_ChIP_overlap",
                         "P1_5_10","P1_5_10_ID","P1_5_10_overlap","P5_10","P5_10_ID",
                         "P5_10_overlap","P45","P45_ID","P45_overlap")]

DMR432_capture = DMR432[,c("DMR_mm10_chr","DMR_mm10_start","DMR_mm10_end",
                           "DMR_annot_chr","DMR_annot_start","DMR_annot_end",
                           "strand","entrezgene_accession","gene_biotype",
                           "mgi_symbol", "pvalue", "qvalue",  "meth.diff",
                           "promoters","promoters_ID","promoters_overlap",
                           "H3K27ac","H3K27ac_ID","H3K27ac_overlap","others",
                           "others_ID","others_overlap","ATAC","ATAC_ID",
                           "ATAC_overlap","HSF2_naive_ChIP","HSF2_naive_ChIP_ID",
                           "HSF2_ChIP_overlap","P1_5_10","P1_5_10_ID","P1_5_10_overlap",
                           "P5_10","P5_10_ID","P5_10_overlap","P45","P45_ID","P45_overlap")]

write.table(DMR432_clean,"DMR432_unique_annot_mm10_BMT_meth_cgi_capt_dist500.txt", 
            sep="\t", quote=FALSE, row.names = FALSE, col.names = TRUE)

write.table(DMR432_capture,"DMR432_unique_annot_light_mm10_BMT_capt_dist500.txt", 
            sep="\t", quote=FALSE, row.names = FALSE, col.names = TRUE)
```

#### 24.Manual re-annotation of DMRs included in protocadherins regions¶

##### 24.1. Pcdhs information retrieval¶

There is a limitation in the use of Biomart to annotate DMR432, at the level of protocadherins (Pcdhs). Indeed, since Pcdhs are organized into clusters, and share common genomic regions, if a DMR is included in a Pcdh region, it is systematically associated to almost all Pcdhs of a cluster.
  
  
In order to annotate more precisely the DMRs, the few DMRs located in Pcdhs have been manually reannotated. For that, DMR432 file was loaded on **UCSC Genome browser** in order to identify the Pcdh (exon when possible) in which each DMR concerned is.
  
  
Here are the mm10 coordinates of concerned DMR432, followed by their mm9 coordinates and the Pcdhs that were manually associated to these DMRs.

| mm10\_chr | mm10\_start | mm10\_end | mm9\_chr | mm9\_start | mm9\_end | Pcdh |
| --- | --- | --- | --- | --- | --- | --- |
| chr18 | 36946321 | 36946496 | chr18 | 37105975 | 37106150 | Pcdha3 (exon) |
| chr18 | 36998102 | 36998372 | chr18 | 37157756 | 37158026 | Pcdha9 (exon) |
| chr18 | 37005677 | 37005756 | chr18 | 37165331 | 37165410 | Pcdha11 (exon, according to Gencode), Pcdha10 (according to NCBI RefSeq genes) |
| chr18 | 37669155 | 37669362 | chr18 | 37828809 | 37829016 | Pcdhga2 (exon) |
| chr18 | 37720141 | 37720375 | chr18 | 37879795 | 37880029 | Pcdhgb4 (part of promoter) + intronic region |
| chr18 | 37765890 | 37766023 | chr18 | 37925544 | 37925677 | Pcdhga12 (promoter) |
| chr18 | 37694545 | 37694935 | chr18 | 37854199 | 37854589 | Pcdhga5 (exon) |
| chr18 | 37005158 | 37005610 | chr18 | 37164812 | 37165264 | Pcdha11 (promoter+exon, according to Gencode), Pcdha10 (according to NCBI RefSeq genes) |
| chr18 | 37751794 | 37751861 | chr18 | 37911448 | 37911515 | Pcdhgb7 (exon) |
| chr18 | 37755743 | 37756176 | chr18 | 37915397 | 37915830 | Pcdhga11 (promoter+exon) |

These informations are saved into a file called **Annotation\_Pcdh\_clean.txt** in the folder H:/methylome/fusion/post\_methylkit/DMR432\_pval0.07/annotations/Biomart/mm10/

##### 24.2. Correction of the information of these DMRs in the overall table¶

In [ ]:

```
%%R

# To load file
setwd("H:/methylome/fusion/post_methylkit/DMR432_pval0.07/annotations/Biomart/mm10/")

DMR432 = read.table("DMR432_unique_annot_mm10_BMT_meth_cgi_capt_dist500.txt", 
                    header = TRUE, fill=TRUE, quote="", sep="\t", na.strings = "", 
                    stringsAsFactors = FALSE)

Pcdh = read.table("Annotation_Pcdh_clean.txt", header = TRUE, fill=TRUE, sep="\t", 
                  na.strings = "", stringsAsFactors = FALSE)

head(DMR432)
head(Pcdh)

dim(DMR432)
dim(Pcdh)

# Pcdh reannotation : remove Pcdh "imprecise listing" and replace it with unique Pcdh information identified with UCSC genome browser
Pcdh$DMR_mm10_ID <- as.character(Pcdh$DMR_mm10_ID)
for (i in 1:nrow(Pcdh)){
  ID_Pcdh <- Pcdh$DMR_mm10_ID[i]
  row_to_modify <- which(DMR432$DMR_mm10_ID == ID_Pcdh)
  DMR432[row_to_modify,6:14] <- Pcdh[i,6:14]
}

write.table(DMR432, "DMR432_uniq_annot_mm10_BMT_meth_cgi_capt_dist500_clean_Pcdh.txt", 
            row.names = FALSE, col.names = TRUE, quote=FALSE, sep ="\t")
```

#### 25. Define DMR432 that belongs to more than one capture element¶

In [ ]:

```
%%R

setwd("H:/methylome/fusion/post_methylkit/DMR432_pval0.07/annotations/Biomart/mm10")

# To load file
DMR432 = as.data.frame(read.table("H:/methylome/fusion/post_methylkit/DMR432_pval0.07/annotations/Biomart/mm10/DMR432_uniq_annot_mm10_BMT_meth_cgi_capt_dist500_clean_Pcdh.txt", 
                    header = TRUE, sep = "\t", quote="", fill=TRUE))

head(DMR432)
dim(DMR432)

# For maximum distance = 1000bases
prom1000 = DMR432[DMR432[,"promoters"]=="yes",]
dim(prom1000)

enh1000 = DMR432[DMR432[,"H3K27ac"]=="yes",]
dim(enh1000)

others1000 = DMR432[DMR432[,"others"]=="yes",]
dim(others1000)

prom_enh1000 = DMR432[DMR432[,"promoters"]=="yes" & DMR432[,"H3K27ac"]=="yes",]
dim(prom_enh1000)
View(prom_enh1000)

prom_others1000 = DMR432[DMR432[,"promoters"]=="yes" & DMR432[,"others"]=="yes",]
dim(prom_others1000)
View(prom_others1000)

enh_others1000 = DMR432[DMR432[,"H3K27ac"]=="yes" & DMR432[,"others"]=="yes",]
dim(enh_others1000)
View(enh_others1000)

prom_enh_other1000 = DMR432[DMR432[,"H3K27ac"]=="yes" & DMR432[,"others"]=="yes" & DMR432[,"promoters"]=="yes",]
dim(prom_enh_other1000)
View(prom_enh_other1000)


#Venn Diagram representation
VennDiag1000 <- euler(c( "enhancers (H3K27ac)" = 277, "other regions" = 9, "promoters" = 73,
                     "promoters&enhancers (H3K27ac)" = 47, "enhancers (H3K27ac)&other regions" = 12, 
                     "promoters&other regions" = 5, "promoters&enhancers (H3K27ac)&other regions" = 9))
setwd("H:/methylome/fusion/post_methylkit/DMR432_pval0.07/annotations/Biomart/mm10")
pdf("VennDiagramm - DMR432 - dist1000 repartition.pdf")
plot(VennDiag1000, counts = TRUE, font=1, cex=1, alpha=0.5,
     fill=c("darkgreen", "darkred", "grey"))
dev.off()


# Same process for Hyper-DMR
# For maximum distance = 1000bases
prom1000_hyper = DMR432[DMR432[,"promoters"]=="yes" & as.vector(DMR432[,"meth.diff"])>=0,]
dim(prom1000_hyper)

enh1000_hyper = DMR432[DMR432[,"H3K27ac"]=="yes" & as.vector(DMR432[,"meth.diff"])>=0,]
dim(enh1000_hyper)

others1000_hyper = DMR432[DMR432[,"others"]=="yes" & as.vector(DMR432[,"meth.diff"])>=0,]
dim(others1000_hyper)

prom_enh1000_hyper = DMR432[DMR432[,"promoters"]=="yes" & DMR432[,"H3K27ac"]=="yes" & as.vector(DMR432[,"meth.diff"])>=0,]
dim(prom_enh1000_hyper)
View(prom_enh1000_hyper)

prom_others1000_hyper = DMR432[DMR432[,"promoters"]=="yes" & DMR432[,"others"]=="yes" & as.vector(DMR432[,"meth.diff"])>=0,]
dim(prom_others1000_hyper)
View(prom_others1000_hyper)

enh_others1000_hyper = DMR432[DMR432[,"H3K27ac"]=="yes" & DMR432[,"others"]=="yes" & as.vector(DMR432[,"meth.diff"])>=0,]
dim(enh_others1000_hyper)
View(enh_others1000_hyper)

prom_enh_other1000_hyper = DMR432[DMR432[,"H3K27ac"]=="yes" & DMR432[,"others"]=="yes" & DMR432[,"promoters"]=="yes" & as.vector(DMR432[,"meth.diff"])>=0,]
dim(prom_enh_other1000_hyper)
View(prom_enh_other1000_hyper)


# Venn Diagram representation
library(eulerr)

VennDiag1000_hyper <- euler(c( "enhancers (H3K27ac)" = 191, "other regions" = 3, "promoters" = 30,
                         "promoters&enhancers (H3K27ac)" = 21, "enhancers (H3K27ac)&other regions" = 9, 
                         "promoters&other regions" = 2, "promoters&enhancers (H3K27ac)&other regions" = 1))
setwd("H:/methylome/fusion/post_methylkit/DMR432_pval0.07/annotations/Biomart/mm10")
pdf("VennDiagramm - DMR432_hyper - dist1000 repartition.pdf")
plot(VennDiag1000_hyper, counts = TRUE, font=1, cex=1, alpha=0.5,
     fill=c("darkgreen", "darkred", "grey"))
dev.off()


# Same process for Hypo-DMR
# For maximum distance = 1000bases
prom1000_hypo = DMR432[DMR432[,"promoters"]=="yes" & as.vector(DMR432[,"meth.diff"])<0,]
dim(prom1000_hypo)

enh1000_hypo = DMR432[DMR432[,"H3K27ac"]=="yes" & as.vector(DMR432[,"meth.diff"])<0,]
dim(enh1000_hypo)

others1000_hypo = DMR432[DMR432[,"others"]=="yes" & as.vector(DMR432[,"meth.diff"])<0,]
dim(others1000_hypo)

prom_enh1000_hypo = DMR432[DMR432[,"promoters"]=="yes" & DMR432[,"H3K27ac"]=="yes" & as.vector(DMR432[,"meth.diff"])<0,]
dim(prom_enh1000_hypo)
View(prom_enh1000_hypo)

prom_others1000_hypo = DMR432[DMR432[,"promoters"]=="yes" & DMR432[,"others"]=="yes" & as.vector(DMR432[,"meth.diff"])<0,]
dim(prom_others1000_hypo)
View(prom_others1000_hypo)

enh_others1000_hypo = DMR432[DMR432[,"H3K27ac"]=="yes" & DMR432[,"others"]=="yes" & as.vector(DMR432[,"meth.diff"])<0,]
dim(enh_others1000_hypo)
View(enh_others1000_hypo)

prom_enh_other1000_hypo = DMR432[DMR432[,"H3K27ac"]=="yes" & DMR432[,"others"]=="yes" & DMR432[,"promoters"]=="yes" & as.vector(DMR432[,"meth.diff"])<0,]
dim(prom_enh_other1000_hypo)
View(prom_enh_other1000_hypo)


# Venn diagram representation
library(eulerr)

VennDiag1000_hypo <- euler(c( "enhancers (H3K27ac)" = 86, "other regions" = 6, "promoters" = 43,
                               "promoters&enhancers (H3K27ac)" = 26, "enhancers (H3K27ac)&other regions" = 3, 
                               "promoters&other regions" = 3, "promoters&enhancers (H3K27ac)&other regions" = 8))
setwd("H:/methylome/fusion/post_methylkit/DMR432_pval0.07/annotations/Biomart/mm10")
pdf("VennDiagramm - DMR432_hypo - dist1000 repartition.pdf")
plot(VennDiag1000_hypo, counts = TRUE, font=1, cex=1, alpha=0.5,
     fill=c("darkgreen", "darkred", "grey"))
dev.off()
```

#### 26. Gene ontology of DMR432 into H3K27ac enhancers¶

Gene ontology analysis was performed on DMR432 associated to H3K27ac enhancers.
  
  
Since DMRs were observed in methylome capture that doesn't include all mouse genes, the universe (or background) must be adapted to avoid biases. Thus, universe of the analysis can't be the all mouse genes. We considered, as universe, all genes that overlapped with the K3K27ac enhancers that are in the capture.

Analysis procedure was as follows:

- 1. Extraction of the ID (gene names) of DMRs that are in H3K27ac enhancers.
- 1. Conversion of the coordinates (mm9 to mm10) of all the H3K27ac enhancers included in the capture.
- 1. Annotaion (Biomart annotation) of all the H3K27ac enhancers included in the capture to identify genes that are associated to these promoters.
     *nb : Pcdhs re-annotation is also needed.*
- 1. Extraction of the ID (gene names) of all the H3K27ac enhancers included in the capture.
- 1. Gene ontology analysis using all the H3K27ac enhancers included in the capture as universe.

##### 26.1. Extraction of the ID (gene names) of DMRs that are in H3K27ac enhancers¶

In [ ]:

```
%%R

#--------------------------------------------------------
# 1. To extract the ID (gene names) of DMRs that are in H3K27ac enhancers
#--------------------------------------------------------

# To load file
setwd("H:/methylome/fusion/post_methylkit/DMR432_pval0.07/annotations/Biomart/mm10/")

DMR432 = read.table("DMR432_unique_annot_mm10_BMT_meth_cgi_capt_dist500.txt", header = TRUE, fill=TRUE, quote="", sep="\t", na.strings = "", stringsAsFactors = FALSE)
Pcdh = read.table("Annotation_Pcdh_clean.txt", header = TRUE, fill=TRUE, sep="\t", na.strings = "", stringsAsFactors = FALSE)

head(DMR432)
head(Pcdh)

dim(DMR432)
dim(Pcdh)

# Pcdh reannotation: remove Pcdh "imprecise listing" and replace it with unique Pcdh information identified with UCSC genome browser
Pcdh$DMR_mm10_ID <- as.character(Pcdh$DMR_mm10_ID)
for (i in 1:nrow(Pcdh)){
  ID_Pcdh <- Pcdh$DMR_mm10_ID[i]
  row_to_modify <- which(DMR432$DMR_mm10_ID == ID_Pcdh)
  DMR432[row_to_modify,6:14] <- Pcdh[i,6:14]
}

#already done:
#write.table(DMR432, "DMR432_uniq_annot_mm10_BMT_meth_cgi_capt_dist500_clean_Pcdh.txt", 
# row.names = FALSE, col.names = TRUE, quote=FALSE, sep ="\t")


# Extraction of DMRs corresponding to H3K27ac enhancers (all, hyper, hypo)
DMR_all_enh = DMR432[DMR432[,"H3K27ac"]=="yes",]
DMR_hyper_enh = DMR_all_enh[DMR_all_enh[,"meth.diff"]>0,]         
DMR_hypo_enh = DMR_all_enh[DMR_all_enh[,"meth.diff"]<0,]          

head(DMR_all_enh)
head(DMR_hyper_enh)
head(DMR_hypo_enh)

dim(DMR_all_enh)
dim(DMR_hyper_enh)
dim(DMR_hypo_enh)

# Function to extract ID
unlist_gene = function(treatment_file, gene_ID, file_name, folder){
  # ID Eextraction - Biomart mm10 annotation: 
  # Obtention of a matrix with only one column, containing ENSEMBL ID
  gene_name_ID = as.character(treatment_file[,gene_ID])
  
  # Split element of interest to extract all the informations (all the ID homonyms)
  list_gene_splitted <- strsplit(gene_name_ID,split=";")
    vec_gene_splitted <- unique(unlist(list_gene_splitted))

  # remove "."  
  vec_gene_splitted_filter = vec_gene_splitted[vec_gene_splitted!="."]
  
  # Present data as a dataframe
  table_VGS_filter = as.data.frame(vec_gene_splitted_filter)
  
  write.table(table_VGS_filter, paste(folder,gene_ID,"_",file_name,".txt",sep=""), 
              quote=FALSE, col.names=FALSE, row.names=FALSE)
}


# Function application: extract Entrez accession ID
unlist_gene(treatment_file=DMR_all_enh, gene_ID="entrezgene_accession", file_name="DMR432_all_enhancers",
            folder="H:/methylome/fusion/post_methylkit/DMR432_pval0.07/GO/")

unlist_gene(treatment_file=DMR_hyper_enh, gene_ID="entrezgene_accession", file_name="DMR432_hyper_enhancers",
            folder="H:/methylome/fusion/post_methylkit/DMR432_pval0.07/GO/")

unlist_gene(treatment_file=DMR_hypo_enh, gene_ID="entrezgene_accession", file_name="DMR432_hypo_enhancers",
            folder="H:/methylome/fusion/post_methylkit/DMR432_pval0.07/GO/")
```

##### 26.2. Conversion of the coordinates (mm9 to mm10) of all H3K27ac enhancers included in the capture¶

As for DMR432 analysis, **LiftOver** tool proposed by **UCSC** was used to convert mm9 coordinates of all H3K27ac enhancers that are in the capture into mm10 coordinates.
  
  
For the analysis, we used the .bed file called **in\_big\_capt\_H3K27Ac\_sorted.txt** in H:\methylome\capture\redo\agrandi\ok.
  
  
Default settings of **LiftOver** tool have been retained :

- Minimum ratio of bases that must remap: 0.95
- BED 4 to BED 6 Options
  - Don’t allow multiple output regions
  - Minimum hit size in query: 0
  - Minimum chain size in target:0
- BED 12 Options
  - Min ratio of alignment blocks or exons that must map:1
  - If thickStart/thickEnd is not mapped, use the closest mapped base : No

Some regions (10/34279) can't be converted for distinct reasons:

In [ ]:

```
#--------------------------------------------------------
# 2. Convert coordinates (mm9 to mm10) of all H3K27ac enhancers included in the capture
#--------------------------------------------------------

# Successfully converted 34269 records. 

# Conversion failed on 10 records:
#Partially deleted in new (Sequence insufficiently intersects one chain)
chr11	88407882	88408998
#Partially deleted in new (Sequence insufficiently intersects one chain)
chr11	88459561	88460861
#Partially deleted in new (Sequence insufficiently intersects one chain)
chr13	83724370	83725670
#Partially deleted in new (Sequence insufficiently intersects one chain)
chr5	30644712	30645730
#Partially deleted in new (Sequence insufficiently intersects one chain)
chr5	33015542	33016518
#Split in new (Sequence insufficiently intersects multiple chains)
chr5	128845131	128846431
#Partially deleted in new (Sequence insufficiently intersects one chain)
chr7	3303200	3304500
#Partially deleted in new (Sequence insufficiently intersects one chain)
chr7	3389443	3390743
#Partially deleted in new (Sequence insufficiently intersects one chain)
chr7	3456791	3458845
#Partially deleted in new (Sequence insufficiently intersects one chain)
chr7	89393667	89394967
```

##### 26.3. Annotation of all H3K27ac enhancers included in the capture¶

To annotate all the H3K27ac enhancers included in the capture using **BiomaRt**, annotation file named **biomart\_mm10\_all\_info\_sorted.bed** (in folder G:/projet\_DU\_AD/results/annotation\_mm10/Biomart/) was used. This file is already formatted (chromosome is named properly, *i.e* chr1 and not 1 - ; strand is not "+/-" but "1/-1" ; remove of the column called "uniprot\_genename" and "entrezgene\_description" because bedtools intersect doesn't work if these columns are kept).

In [ ]:

```
%%bash

#--------------------------------------------------------
# 3. Annotatation of (Biomart annotation) all the H3K27ac enhancers 
# included in the capture to identify genes that are associated to these regions.
#--------------------------------------------------------

# 1.To sort files
# Biomart annotation file is already sorted
# To sort file containing all H3K27ac enhancers of the capture (mm10 coordinates)

cd /mnt/h/methylome/capture/redo/agrandi/ok/mm10/

sort -k1,1 -k2,2n mm10_in_big_capt_H3K27Ac.bed > mm10_in_big_capt_H3K27Ac_sorted.bed

head mm10_in_big_capt_H3K27Ac_sorted.bed

# 2.Annotation, using bedtools intersect 
bedtools intersect -wao 
        -a mm10_in_big_capt_H3K27Ac_sorted.bed 
        -b /mnt/h/methylome/capture/redo/agrandi/ok/mm10/Biomart/biomart_mm10_all_info_sorted.bed > /mnt/h/methylome/capture/redo/agrandi/ok/mm10/Biomart/annot_BMT_mm10_in_big_capt_H3K27Ac.bed

head /mnt/h/methylome/capture/redo/agrandi/ok/mm10/Biomart/annot_BMT_mm10_in_big_capt_H3K27Ac.bed

wc -l /mnt/h/methylome/capture/redo/agrandi/ok/mm10/Biomart/annot_BMT_mm10_in_big_capt_H3K27Ac.bed
# 40 707 /mnt/h/methylome/capture/redo/agrandi/ok/mm10/Biomart/annot_BMT_mm10_in_big_capt_H3K27Ac.bed
```

In [ ]:

```
%%R

# 3.To concatenate annotation informations : one row = one region of interest / DMR
# 3.1. To load DMR432 file containing biomart annotation.

enh_annot = read.table(file = "H:/methylome/capture/redo/agrandi/ok/mm10/Biomart/annot_BMT_mm10_in_big_capt_H3K27Ac.bed", 
                       header = FALSE, sep = "\t", fill=TRUE)

head(enh_annot)
dim(enh_annot)

# To create the ID = chr-start-end
enh_annot_ID = cbind("ID" = paste(enh_annot$V1, enh_annot$V2, enh_annot$V3, sep = ";"),enh_annot)

head(enh_annot_ID)

# 3.2. To concatenate informations
library(dplyr)

# Liste of unique ID 
liste_chrom_unique <- unique(enh_annot_ID$ID)

# empty matrice which will be filled during the loop
# Nb of rows = nb of unique ID | Nb of columns = nb of variables
treatment_file <- data.frame(matrix(NA,nrow=length(liste_chrom_unique),ncol=ncol(enh_annot_ID)))
colnames(treatment_file) <- colnames(enh_annot_ID)

treatment_file$ID <- liste_chrom_unique
for(i in 1:length(liste_chrom_unique)){
  # Filter according to ID k 
  enh_filt <- enh_annot_ID %>%
    filter(ID == liste_chrom_unique[i])
  
    # Collapsing of distincts elements of a variables for ID k 
    # If variable is composed of only one element ==> we keep this element only once 
    # If variable is composed of m distinct elements ==> even if elements can be repeated, we keep all the elements (with repetitions)
   for(j in 2:length(colnames(treatment_file))){
     if(length(unique(enh_filt[,j])) == 1){
       treatment_file[i,j] <- paste(unique(enh_filt[,j]), collapse=';')
     } else {
       treatment_file[i,j] <- paste(enh_filt[,j], collapse=';')  
     }
   }
 }

head(treatment_file)
dim(treatment_file)

# To rename properly the column
colnames(treatment_file) = c("mm10_ID_H3K27ac","mm10_chr_H3K27ac",
                             "mm10_start_H3K27ac","mm10_end_H3K27ac",
                             "mm10_chr_annot","mm10_start_annot",
                             "mm10_end_annot", "strand","entrezgene_accession", 
                             "gene_biotype", "mgi_symbol", "ENSEMBL_ID",
                             "Entrez_ID","overlap_length")

head(treatment_file)

# To save the data into a file
write.table(treatment_file, "H:/methylome/capture/redo/agrandi/ok/mm10/Biomart/annot_uniq_BMT_mm10_in_big_capt_H3K27Ac.bed",
            col.names = TRUE, sep = "\t", quote=FALSE, row.names=FALSE)
```

*nb: for a more precise annotation, H3K27ac enhancers that are located into **protocadherins** were manually reannotated, as we did for DMR432 regions*

##### 26.4. Extraction of the ID (gene names) of all H3K27ac enhancers included in the capture¶

In [ ]:

```
%%R 

#--------------------------------------------------------
# 4. Extract the ID (gene names) of all H3K27ac enhancers included in the capture
#--------------------------------------------------------

# ========== For H3K27ac enhancers ===============

# Function to extract ID
unlist_gene = function(treatment_file, gene_ID, file_name, folder){
  # ID ENSEMBL extraction - Biomart mm10 annotation: 
  # Obtention of a matrix with only one column, containing ENSEMBL ID
  gene_name_ID = as.character(treatment_file[,gene_ID])
  
  # To split element of interest to extract all the informations (all the ID homonyms)
   list_gene_splitted <- strsplit(gene_name_ID,split=";")
   vec_gene_splitted <- unique(unlist(list_gene_splitted))
  
  # To remove "."
  vec_gene_splitted_filter = vec_gene_splitted[vec_gene_splitted!="."]
  
  # To present data as a dataframe
   table_VGS_filter = as.data.frame(vec_gene_splitted_filter)
   
   write.table(table_VGS_filter, paste(folder,gene_ID,"_",file_name,".txt",sep=""), 
               quote=FALSE, col.names=FALSE, row.names=FALSE)
 }

# Function application: to extract Entrez accession ID
unlist_gene(treatment_file=treatment_file, gene_ID="entrezgene_accession", 
            file_name="in_big_capt_H3K27Ac",  folder="H:/methylome/fusion/post_methylkit/DMR432_pval0.07/GO/rawdata/")
```

##### 26.5. Gene ontology analysis¶

For the Gene ontology analysis, all the promoters / H3K27ac enhancers included in the capture were used as a background.
**WEB-based GEne SeT AnaLysis Toolkit (WebGestalt)** was used to performed the Gene Ontology analysis.
