## Supplementary material for "Immediate perturbations of DNA methylation and transcriptome upon acute prenatal alcohol exposure in the mouse developing brain cortex": Supp data Notebook 2 ATAC-seq data mining: Supp. data Notebook 2 - ATAC-seq workflow - Duchateau et al..html

Annexe 3.2 ATAC-seq workflow


### Supplementary data Notebook 2 - ATAC-seq workflow: analysis of the physiological evolution of chromatin accessibility throughout brain development using ENCODE available data¶

##### *Agathe Duchateau, Délara Sabéran Djoneidi, Yoann Devriendt*¶

First part of the analysis (quality control of the raw data, trimming, mapping and removal of duplicates and mitochondrial chromosome) was done using tools of Galaxy platform (**usegalaxy.eu**, analysis done by Délara Sabéran Djoneidi and Yoann Devriendt).
Second part of the analysis (from mapping to the end of the analysis) was done on Unix and R environments (analysis done by Agathe Duchateau).

#### Download ENCODE data¶

To define chromatin accessibility profile in the developing brain, ATAC-seq data time-course analysis was performed, using available public ENCODE data. These ATAC-seq experiments were done using embryonic and newborn forebrains of mixed sex C57BL/6N mice (experiments from Bing Ren laboratory, UCSD).
Developmental stages studied in duplicates were :

E11.5 (ENCODE accession : ENCFF419LDW + ENCFF963YIU, ENCFF083TDB + ENCFF680UAR)
  
E12.5 (ENCODE accession : ENCFF413XTH + ENCFF119TXW, ENCFF199UBT + ENCFF171APM)
  
E13.5 (ENCODE accession : ENCFF401VUV + ENCFF898NRO, ENCFF721LGJ + ENCFF777UKE)
  
E14.5 (ENCODE accession : ENCFF048MTG + ENCFF890LGM, ENCFF633MTW + ENCFF666DRJ)
  
E15.5 (ENCODE accession : ENCFF248PXW + ENCFF825UHO, ENCFF906VXU + ENCFF500SXI)
  
E16.5 (ENCODE accession :ENCFF058IAE + ENCFF765HUX, ENCFF776GDQ + ENCFF588XZG)
  
and postnatal day 0 (P0, Encode accession : ENCFF197GTC + ENCFF209GGJ, ENCFF296GZG + ENCFF664RZO).

These data were obtained in Bing Ren lab (UCSD) using an ATAC-seq protocol on frozen tissue. In brief, frozen tissues were pulverized in liquid nitrozen using pestle and mortar, isolated nuclei were then permeabilized and tagmentation performed during 30mn at 37°C. Library were generated by PCR amplification of DNA extracted using **minElute PCR purification Kit**. Size selection were performed using **Ampure XP beads**. For more explanations on experimental procedures, please see Experiment Summary on ENCODE database, pages available with Encode accession of samples cited above.

#### Quality control and statistics¶

After simplifying fastq.gz file names, **FASTQC Read Quality reports** tool (Galaxy Version 0.72) was used to verify the quality of the sequenced raw data, this tool is also used after triming step (see below).
In order to follow the steps of the workflow and identify the rate of remaining reads, **Samtools stats tool** (Galaxy Version 2.0.2+galaxy1) was run after each treatment step. These statistics are summarised in a table.

#### Trimming¶

Reads having bad sequencing quality were trimmed using **Trimmomatic** tool (Galaxy Version 0.32.3) with the following (default) parameters :
Adapter sequences to use : **Nextera (paired ended)**
Maximum mismatch count which will still allow a full match to be performed : **2**

How accurate the match between the two 'adapter ligated' reads must be for PE palindrome read alignment : **30**

How accurate the match between any adapter etc. sequence must be against a read : **10**

Select Trimmomatic operation to perform : **cut bases off the start of a read, if below a threshold quality (LEADING)**
**cut bases off the end of a read, if below a threshold quality (TRAILING)**

Minimum quality required to keep a base : **3**

#### Mapping¶

Trimmed reads were then paired-end mapped on Mus musculus reference genome mm9, using **Bowtie2** (Galaxy Version 2.3.4.3) with the default parameters but with the **Sensitive end-to-end** (**--sensitive**) option.

#### Removal of duplicates and reads on mitochondrial chromosome¶

Then, reads were deduplicated using **Samtools rmdup** (Galaxy Version 2.0.1) indicating that the input files are paired-end bam files.
Reads that mapped to mitochondrial chromosome were removed using tools on Galaxy platform, by splitting the mapped reads per chromosome using **Bam-splitter** tool (Galaxy Version 2.4.0). Then all the files were merged except the reads mapped on mitochondrial chromosome, using **Merge-Bam Files** tool (Galaxy Version 1.2.0) that uses the **Picard** merge command.

#### Md5sum identification¶

**Secure Hash** (Galaxy Version 0.0.1) were used to define Md5sum for each Merged Bam file of deduplicated mapped reads without mitochondrial chromosome.

E11.5  
ENCFF419LDW + ENCFF963YIU 5d6ac81ede8c5ea207a07c75924a8aab

#### Setting up of bioinformatic tools in a conda environment¶

Following tools were installed in a conda environment to continue ATAC-seq analysis. Version of each tool is indicated. R packages which are cited below were also used to analyse the data.

In [ ]:

```
conda create -n atac #conda version: 4.6.14

conda activate atac

conda install MACS2 # version 2.1.2
conda install bedtools # version 2.28.0
conda install htseq # version 0.11.2

# R version 3.5.2
# R package used for analysis : 
library(edgeR) # version 3.24.3
library(ggplot2) # version 3.2.1
library(reshape2) # version1.4.3
library(GGally) # version 1.4.0
library(SARTools) # version 1.6.9
library(BiomaRt) #version 2.38.0
```

#### Input files download¶

Merged Bam file of deduplicated mapped reads without mitochondrial chromosome generated on Galaxy platform were downloaded locally and rename following this nomenclature : stage\_rep\_ss\_mito.bam (e.g. E15\_B\_ss\_mito.bam).
MD5sum of files of Galaxy server and downloaded files were compared and are similar.

#### Peak calling to find potential accessible chromatin regions¶

**MACS2** tool was used to identify enriched regions, corresponding to potential open chromatin regions.

**Selected parameters :**

- macs2 callpeak : Peakcalling module
- -t treated\_file.bam : input file. In case of more than one file, MACS2 merges files before performing the peakcalling step.
- --name E11 : prefix name that will be given to output files
- --outdir folder : output folder
- --format BAMPE : it indicates that paired-end bam files are used.
- --gsize mm : size of the reference genome (mm for Mus musculus, mm9 = 1.87e+09)
- --tsize 50 : size of mapped reads
- --qvalue 0.05 : qvalue threshold (= FDR value threshold). Default :0.05. qvalues are based on pvalues and obtained using the Benjamini-Hochberg method
- --keep-dup 1 : if duplicates exists, it keeps only 1 read.
- --bdg : to obtain bedGraph file
- --mfold MFOLD MFOLD : select the regions within MFOLD range of high-confidence enrichment ratio against background to build model. Fold-enrichment in regions must be lower than upper limit, and higher than the lower limit. Default :5 50
- --bw BW : band width for picking regions to compute fragment size. Default :300

In [ ]:

```
%%bash 

mkdir -p /mnt/g/projet_DU_DSD/results/06_Peaks_calling/dup_pool/

# Samples name
sample="E11 E12 E13 E14 E15 E16 P0"

cd /mnt/g/projet_DU_DSD/results/05_BAM_wt_mt/

for ech in ${sample}
do
    echo  "---------------------------"
    echo  "Peaks_calling - MACS2 "
    echo  "samples : ${ech}_ss_mito.bam"
    echo  "---------------------------"
    macs2 callpeak -t ${ech}_A_ss_mito.bam ${ech}_B_ss_mito.bam --name ${ech} --outdir /mnt/g/projet_DU_DSD/results/06_Peaks_calling/dup_pool/${ech} --format BAMPE --gsize mm --tsize 50 --qvalue 0.05 --keep-dup 1 --bdg --mfold 5 50 --bw 300
done
```

#### Obtaining a table with detected peaks from all samples¶

**bedtools multiinter** was used to create a table containing all detected peaks, whether they are common between the samples or not. We obtain a table with one region per row and one sample per column. The value "1" is assigned to a sample if the region is observed in this sample, otherwise "0" is assigned.

**Selected parameters :**

- -header : print a header line
- -i : input file
- -names : prefix name to describe each input file

In [ ]:

```
%%bash

#---------------------------
# 1.Files have to be sorted
#---------------------------

# Samples name
sample="E13 E14 E15 E16 P0"

for ech in ${sample}
do
    echo "sorted des pics de l'ech : ${ech}"
    sort -k1,1 -k2,2n /mnt/g/projet_DU_DSD/results/06_Peaks_calling/dup_pool/${ech}/${ech}_peaks.narrowPeak > /mnt/g/projet_DU_DSD/results/06_Peaks_calling/dup_pool/${ech}/${ech}_sorted.narrowPeak 
    head /mnt/g/projet_DU_DSD/results/06_Peaks_calling/dup_pool/${ech}/${ech}_sorted.narrowPeak 
done

#---------------------------
# 2.Obtention of the table with all detected peaks
#---------------------------

mkdir -p /mnt/g/projet_DU_DSD/results/07_all_peaks_MI/dup_pool

cd /mnt/g/projet_DU_DSD/results/06_Peaks_calling/dup_pool/	

bedtools multiinter -header -i E13/E13_sorted.narrowPeak E13/E13_sorted.narrowPeak E14/E14_sorted.narrowPeak E14/E14_sorted.narrowPeak E15/E15_sorted.narrowPeak E15/E15_sorted.narrowPeak E16/E16_sorted.narrowPeak E16/E16_sorted.narrowPeak P0/P0_sorted.narrowPeak P0/P0_sorted.narrowPeak -names E13_A E13_B E14_A E14_B E15_A E15_B E16_A E16_B P0_A P0_B > /mnt/g/projet_DU_DSD/results/07_all_peaks_MI/dup_pool/E13_a_P0_all_peaks_dup_pool.bed
```

#### Merge overlapping regions¶

**bedtools merge** was used to merge overlapping regions of the previous file.

**Selected parameters :**

- -header : print a header line
- -i : input file
- -d 0 : maximum distance allowed between features for regions to be merged. Default :0, that is, overlapping and book-ended features are merged.

In [ ]:

```
%%bash

#---------------------------
# 1.Files have to be sorted
#---------------------------

cd /mnt/g/projet_DU_DSD/results/07_all_peaks_MI/dup_pool

sed '1d' E13_a_P0_all_peaks_dup_pool.bed > nh_E13_a_P0_all_peaks_dup_pool.bed

sort -k1,1 -k2,2n nh_E13_a_P0_all_peaks_dup_pool.bed > nh_E13_a_P0_all_peaks_dup_pool_sorted.bed

#---------------------------
# 2.Merge overlapping regions
#---------------------------

mkdir -p /mnt/g/projet_DU_DSD/results/08_all_peaks_merge/dup_pool

bedtools merge -header -i nh_E13_a_P0_all_peaks_dup_pool_sorted.bed -d 0 > /mnt/g/projet_DU_DSD/results/08_all_peaks_merge/dup_pool/E13_a_P0_all_peaks_dup_pool_merge.bed

wc -l  /mnt/g/projet_DU_DSD/results/08_all_peaks_merge/dup_pool/E13_a_P0_all_peaks_dup_pool_merge.bed
# ---> 115752 /mnt/g/projet_DU_DSD/results/08_all_peaks_merge/dup_pool/E13_a_P0_all_peaks_dup_pool_merge.bed <----
```

#### Obtaining a count table of merged regions¶

**htseq-count** was used to obtain reads count of regions for each sample. In order to respect the format of input file, a 'pseudo' .gff files, was created containing regions information.

*nb : line ends encoding of the .gff file must be in Unix format, otherwise htseq-count does not work correctly. It was changed using **Notepad ++** software.*

To avoid biases from paired-end reads overlaps, files must be sorted by name, to allow htseq-count to count paired-end reads features only once.

In [ ]:

```
# Load file containing merged regions
dataset=read.delim("G:/projet_DU_DSD/results/08_all_peaks_merge/dup_pool/nh_E13_a_P0_all_peaks_dup_pool_merge_sorted.bed", header=FALSE, sep="\t")

colnames(dataset)=c("chr","start","end")

head(dataset)

# Creation of specific columns to obtain a .gff formatted file : 
dataset$source = "MACS2" #annotation source
dataset$type = "peaks" #element type
dataset$score ="."
dataset$strand = "."
dataset$phase = "."
dataset$group = paste("peak_id ","\"Peak",1:nrow(dataset),"\"",sep="") #peak_id

head(dataset)

# data reorganization
dataset.gff=dataset[,c("chr","source","type","start","end","score","strand","phase","group")]

head(dataset.gff)

# Save
write.table(dataset.gff, "G:/projet_DU_DSD/results/08_all_peaks_merge/dup_pool/nh_E13_a_P0_all_peaks_dup_pool_merge_sorted.gff", quote=FALSE,row.names = FALSE,col.names=FALSE, sep="\t")
write.table(dataset.gff, "G:/projet_DU_DSD/results/08_all_peaks_merge/dup_pool/E13_a_P0_all_peaks_dup_pool_merge_sorted.gff", quote=FALSE,row.names = FALSE,col.names=TRUE, sep="\t")
```

In [ ]:

```
%%bash 

cd /mnt/g/projet_DU_DSD/results/05_BAM_wt_mt/

# Samples name
sample="E13_A E13_B E14_A E14_B E15_A E15_B E16_A E16_B P0_A P0_B"

for ech in ${sample}
do
    echo "----------------------------------"
    echo "samtools sort - sample : ${ech}"
    echo "----------------------------------"
    samtools sort -n -o ${ech}_ss_mito_sorted.bam ${ech}_ss_mito.bam

    echo "----------------------------------"
    echo "htseq-count - sample : ${ech}"
    echo "-----------------------------------"
    htseq-count --stranded=no --format bam --order name -a=10 --type=peaks --idattr=peak_id --mode=union --nonunique=none ${ech}_ss_mito_sorted.bam /mnt/g/projet_DU_DSD/results/08_all_peaks_merge/dup_pool/nh_E13_a_P0_all_peaks_dup_pool_merge_sorted_Unix.gff > /mnt/g/projet_DU_DSD/results/09_all_peaks_count_table/dup_pool/count_${ech}_dup_pool.txt
done
```

**Selected parameters :**

- --stranded=no : to specify that library is non-strand specific
- --format bam : type of alignment file
- --order name : to specify the sorting order. For paired-end data, the alignment have to be sorted either by read name or by alignment position.
- -a 10 : skip all reads with alignment quality lower than the given minimum value. Default :10
- --type=peaks : element type of the .gff file to be considered, other element types are ignored
- --idattr=peak\_id : attribute from .gff file to be used as element ID.
- --mode=union : mode to handle reads overlapping more than one feature. With union mode, a read is count if there is a overlap with a given region, even if the overlap is partial, as long as there is no ambiguous case (such as overlap with two distinct regions)
- --nonunique=none : reads that mapped several genes is not considered

#### Statistical analysis¶

To identify regions where chromatin accessibility significantly changes during development, pairwise comparisons of successive developmental stages were performed using **edgeR**, an R software and Bioconductor package.

##### Filtering very-low count regions¶

Since regions with very low count across all samples could interfere with the statistical approximations of edgeR, these regions were removed using SARTools filter, *i.e.* by selecting regions which contain at least **minReplicates** (smallest number of replicates = **2**) with at least **counts per million cutoff** (**cpmCutoff = 1**).

##### Normalization to avoid regions composition biases¶

To be able to compare read counts between samples, normalization was carried out according to the edgeR package. Library of each sample was normalised for regions composition using **calcNormFactors()** function, based on a trimmed mean of M-values (TMM) between each pair of samples.
  
Boxplots of raw- and normalised counts distribution were compared to verify the quality of the normalization process. Normalization is supposed to stabilize distributions across samples.

##### Statistical test for differential chromatin accessibility of regions between two successive developmental stages¶

###### Statistical test¶

Differential analysis was carried out according to the edgeR model. **edgeR** aims at fitting one linear model per region. The edgeR model assumes that the count data follow a negative binomial distribution which is a suitable way to analyse the data when variance is higher than the mean. First step of the statistical procedure is to estimate the dispersion of the data.
  
After estimation of dispersions - *i.e.* common (unique value), trended (estimated with splines) and tagwise (estimated from feature counts) dispersions - fitting to a generalized linear model (glm) can be done using **glmFit()** function. Then, statistical test was performed for all pairwise comparisons of successive developmental stages, using a likehood ratio test (lrt method).

###### Differential analysis plots¶

###### MA-plot¶

For each comparison, **MA-plot** of the data was done. It represents the log ratio of differential expression as a function of the mean intensity for each region.

###### Volcano plot¶

**Volcano plots** for the comparisons were performed. It represents the log of the adjusted pvalue as a function of the log ratio of differential expression.

##### Description plots to estimate variability within the experiment¶

###### Scatterplot¶

A **pairwise scatterplot** was produced to verify that intra-group (replicates) similarities are higher than inter-group (samples from distinct developmental stage) ones . This plot is obtained using **log2(counts+1)**, instead of **raw count** values. This pairwise scatterplot is associated with a **SERE** statistic, that was used as a similarity index between ATAC-seq samples.

###### Hierarchical clustering and dendrogram¶

**Hierarchical clustering** and **dendrogram** were performed to estimate variability between samples, *i.e.* determine if samples from distinct developmental stages are separated, while replicates are close to each other. Hierarchical clustering was done after a transformation of the count data as **moderated log-counts-per-million**.
  
To plot dendrogram, an euclidean distance is computed between samples. **Dendrogram** is then obtained from CPM data and built upon the **Ward criterion**.

###### multidimensional scaling plot¶

First two dimensions of a multidimensional scaling plot were plotted to visualise experiment variability. If biological variability is the main source of variance in the data, first dimension is expected to separate samples from the different developmental stages.

##### Adaptations of SARTools functions¶

Following parameters were modified or added from the original Sartools script, to obtain additional graphs or informations:

- addition of a filter to remove uncovered reads. counts\_nf --> counts. In theory, this step is not necessary because uncovered regions should be removed during the filter proposed by SARTools, which deletes regions with very low-counts. However, it allows to have a double control of this step.
- addition of a unormalised library size histogram graph plot (ggplot2).
- change of the raw pvalues histogram design (ggplot2) : summarizeResults.edgeR() --> summarizeResults.edgeR.AD()
- addition of supplementary graphs in run.edgeR() function
- export of supplementary informations (logFC, FDR)
- addition of a filter using logFC and padj thresholds to obtain a restricted list of differentially open or closed regions, in addition to the list given by SARTools script (based on padj threshold only).

In [ ]:

```
# 1. Creation of a file called target.txt, which contains samples information:
target_cible = matrix(0,ncol=4, nrow=10)
colnames(target_cible)=c("label","files","group","day")
target_cible[,"label"]= c("E13.5_A","E13.5_B","E14.5_A", "E14.5_B","E15.5_A","E15.5_B","E16.5_A", "E16.5_B","P0_A","P0_B")

target_cible[,"files"]=c("count_E13_A_dup_pool.txt", "count_E13_B_dup_pool.txt", "count_E14_A_dup_pool.txt","count_E14_B_dup_pool.txt",
                         "count_E15_A_dup_pool.txt", "count_E15_B_dup_pool.txt", "count_E16_A_dup_pool.txt","count_E16_B_dup_pool.txt",
                         "count_P0_A_dup_pool.txt", "count_P0_B_dup_pool.txt")

target_cible[,"group"]=c(rep("E13",2),rep("E14",2),rep("E15",2), rep("E16",2),rep("PO",2))
target_cible[,"day"]="d1"

head(target_cible)
#     label     files                      group day 
# [1,] "E13.5_A" "count_E13_A_dup_pool.txt" "E13" "d1"
# [2,] "E13.5_B" "count_E13_B_dup_pool.txt" "E13" "d1"
# [3,] "E14.5_A" "count_E14_A_dup_pool.txt" "E14" "d1"
# [4,] "E14.5_B" "count_E14_B_dup_pool.txt" "E14" "d1"
# [5,] "E15.5_A" "count_E15_A_dup_pool.txt" "E15" "d1"
# [6,] "E15.5_B" "count_E15_B_dup_pool.txt" "E15" "d1"

write.table(target_cible, "G:/projet_DU_DSD/results/09_all_peaks_count_table/dup_pool/raw_data/target.txt", quote=FALSE, col.names=TRUE,row.names=FALSE, sep="\t")


# This file is then splitted to obtain pairwise successive developmental stages.


# 2. Identification of differentially open or closed chromatin regions during brain development.
# edgeR analysis using modified SARTools script

# ---------
# ----------------------
# --------------------------------------------
# E13 vs E14 developmental stages comparison
# --------------------------------------------
# ----------------------
# ---------

dir.create("G:/projet_DU_DSD/results/09_all_peaks_count_table/dup_pool/SARTools/Analyse_L/E13.14", recursive=TRUE)

# -------------------
# Parameters setting
# -------------------

rm(list=ls())                                        

workDir <- "G:/projet_DU_DSD/results/09_all_peaks_count_table/dup_pool/SARTools/Analyse_L/E13.14"      # working directory for the R session

projectName <- "Sartools-edgeR-E13.14"               # name of the project
author <- "Agathe D."                                # author of the statistical analysis/report

targetFile <- "G:/projet_DU_DSD/results/09_all_peaks_count_table/dup_pool/raw_data/targetE13.14.txt"                           # path to the design/target file
rawDir <- "G:/projet_DU_DSD/results/09_all_peaks_count_table/dup_pool/raw_data/"                                      # path to the directory containing raw counts files
featuresToRemove <- c("alignment_not_unique",        # names of the features to be removed
                      "ambiguous", "no_feature",     # (specific HTSeq-count information and rRNA for example)
                      "not_aligned", "too_low_aQual")# NULL if no feature to remove

varInt <- "group"                                    # factor of interest
condRef <- "E13"                                     # reference biological condition
batch <- NULL                                        # blocking factor: NULL (default) or "batch" for example

alpha <- 0.05                                        # threshold of statistical significance
pAdjustMethod <- "BH"                                # p-value adjustment method: "BH" (default) or "BY"

cpmCutoff <- 1                                       # counts-per-million cut-off to filter low counts
gene.selection <- "pairwise"                         # selection of the features in MDSPlot
normalizationMethod <- "TMM"                         # normalization method: "TMM" (default), "RLE" (DESeq) or "upperquartile"

colors <- c("darkblue","brown3")               

forceCairoGraph <- FALSE

nsamples = 4                                        # nb of samples 
nstage = 2                                          # nb of developemental stages
nrep = 2                                            # nb of replicates per developmental stages
couleur=c("cyan4","brown3","darkgreen","darkblue","plum4","lightgoldenrod3","pink")


# -------------------
# Load packages
# -------------------

setwd(workDir)
library(SARTools)
library(reshape2)
library(ggplot2) 
library(GGally)

#--------------------
# Running script
# checking parameters (unchanged SARTools function)
#--------------------

if (forceCairoGraph) options(bitmapType="cairo")

checkParameters.edgeR(projectName=projectName,author=author,targetFile=targetFile,
                      rawDir=rawDir,featuresToRemove=featuresToRemove,varInt=varInt,
                      condRef=condRef,batch=batch,alpha=alpha,pAdjustMethod=pAdjustMethod,
                      cpmCutoff=cpmCutoff,gene.selection=gene.selection,
                      normalizationMethod=normalizationMethod,colors=colors)

#--------------------
# loading target file (unchanged SARTools function)
#--------------------
target <- loadTargetFile(targetFile=targetFile, varInt=varInt, condRef=condRef, batch=batch)

#-------------------
# loading counts 
#-------------------
counts_nf <- loadCountData(target=target, rawDir=rawDir, featuresToRemove=featuresToRemove)

# Delete rows without counts :
counts <- counts_nf[-(which(rowSums(counts_nf) < 1)),]

#------------------
# description plots 
#------------------
# (unchanged SARTools function)
majSequences <- descriptionPlots(counts=counts, group=target[,varInt], col=colors)

# Supplementary graphs
# Print library size (unormalised data) : 
counts_reshape = melt(counts, id.vars = colnames(counts), variable.name='samples_name')
head(counts_reshape)
dim(counts_reshape)
colnames(counts_reshape) = c("peak_name","samples_name","filter_count")
counts_reshape$stages = with(counts_reshape, rep(substr(colnames(counts), 1, 3), each = nrow(counts)))

lib_size_unorm = ggplot(counts_reshape, aes(x=samples_name,y=filter_count, fill = stages))
lib_size_unorm + geom_bar(stat="identity") + scale_fill_manual(values=c("skyblue","royalblue", "pink","tomato2","darkgreen")) +
  ggtitle("library size (unormalised dataset)")+ theme_light() + 
  scale_x_discrete(name ="samples_name") + scale_y_continuous("total number of reads") +
  theme(plot.title = element_text(hjust = 0.5))
ggsave(path ="figures/", filename = "01 library size - unormalised dataset.pdf")
ggsave(path ="figures/", filename = "01 library size - unormalised dataset.png")

#---------------
# edgeR analysis (adapted from SARTools function)
#---------------

run.edgeR <- function(counts, target, varInt, condRef, batch=NULL, cpmCutoff=1, 
                      normalizationMethod="TMM", pAdjustMethod="BH", ...){
  
  # filtering very low-count regions : select features which contain at least minReplicates (smallest number of replicates) with at least cpmCutoff counts per million
  minReplicates <- min(table(target[,varInt]))
  fcounts <- counts[rowSums(cpm(counts) >= cpmCutoff) >= minReplicates,]
  cat("Number of features discarded by the filtering:\n")
  cat(nrow(counts)-nrow(fcounts),"\n")
  
  # building dge object
  design <- formula(paste("~", ifelse(!is.null(batch), paste(batch,"+"), ""), varInt))
  dge <- DGEList(counts=fcounts, remove.zeros=TRUE)
  dge$design <- model.matrix(design, data=target)
  cat("\nDesign of the statistical model:\n")
  cat(paste(as.character(design),collapse=" "),"\n")
  
  ggsave(path ="figures/", filename ="02 - correlation between samples - filter - unnorm.pdf",ggpairs(as.data.frame(dge$counts[,1:nsamples])))
  ggsave(path ="figures/", filename ="02 - correlation between samples - filter - unnorm.png",ggpairs(as.data.frame(dge$counts[,1:nsamples])))
  
  ggsave(path ="figures/", filename ="03 - correlation between log samples - filter - unnorm.pdf",ggpairs(as.data.frame(log(dge$counts[,1:nsamples]))))
  ggsave(path ="figures/", filename ="03 - correlation between log samples - filter - unnorm.png",ggpairs(as.data.frame(log(dge$counts[,1:nsamples]))))
  
  # normalization for regions composition bias
  dge <- calcNormFactors(dge, method=normalizationMethod)
  cat("\nNormalization factors:\n")
  print(dge$samples$norm.factors)
  
  # estimating dispersions
  dge <- estimateGLMCommonDisp(dge, dge$design)
  dge <- estimateGLMTrendedDisp(dge, dge$design)
  dge <- estimateGLMTagwiseDisp(dge, dge$design)
  
  # graphical representation 
  # BCV based on average log CPM (dispersion representation) : 
  pdf("figures/04 dispersion plot - glmFit method .pdf")
  plotBCV(dge) 
  dev.off()
  
  png("figures/04 dispersion plot - glmFit method .png")
  plotBCV(dge) 
  dev.off()
  
  # MDS plot : 
  pdf("figures/05 MDS plot - glmFit method - bcv.pdf")  
  mds = plotMDS(dge,  method="bcv",col=rep(couleur[1:nstage],each=nrep), pch=1, cex=1, xlim=c(-0.55,1.0),ylim=c(-0.3,0.4))
  text(mds$x, mds$y, labels=rownames(target), col=rep(couleur[1:nstage],each=nrep), pos=3) 
  dev.off()
  
  png("figures/05 MDS plot - glmFit method - bcv.png")  
  mds = plotMDS(dge,  method="bcv",col=rep(couleur[1:nstage],each=nrep), pch=1, cex=1, xlim=c(-0.55,1.0),ylim=c(-0.3,0.4))
  text(mds$x, mds$y, labels=rownames(target), col=rep(couleur[1:nstage],each=nrep), pos=3) 
  dev.off()
  
  pdf("figures/06 MDS plot - glmFit method - logFC.pdf")  
  mds = plotMDS(dge,  method="logFC",col=rep(couleur[1:nstage],each=nrep), pch=1, cex=1, xlim=c(-0.55,1.0),ylim=c(-0.3,0.4))
  text(mds$x, mds$y, labels=rownames(target), col=rep(couleur[1:nstage],each=nrep), pos=3) 
  dev.off()
  
  png("figures/06 MDS plot - glmFit method - logFC.png")  
  mds = plotMDS(dge,  method="logFC",col=rep(couleur[1:nstage],each=nrep), pch=1, cex=1, xlim=c(-0.55,1.0),ylim=c(-0.3,0.4))
  text(mds$x, mds$y, labels=rownames(target), col=rep(couleur[1:nstage],each=nrep), pos=3) 
  dev.off()
  
  # statistical testing: perform all the comparisons between the levels of varInt
  fit <- glmFit(dge, dge$design, ...)
  cat(paste("Coefficients of the model:",paste(colnames(fit$design),collapse="  ")),"\n")
  colsToTest <- grep(varInt,colnames(fit$design))
  namesToTest <- paste0(gsub(varInt,"",colnames(fit$design)[colsToTest]),"_vs_",condRef)
  results <- list()
  
  # testing coefficients individually (tests againts the reference level)
  for (i in 1:length(colsToTest)){
    cat(paste0("Comparison ",gsub("_"," ",namesToTest[i]),": testing coefficient ",colnames(fit$design)[colsToTest[i]]),"\n")
    lrt <- glmLRT(fit, coef=colsToTest[i])
    results[[namesToTest[i]]] <- topTags(lrt,n=nrow(dge$counts),adjust.method=pAdjustMethod,sort.by="none")$table
  }
  # defining contrasts for the other comparisons (if applicable)
  if (length(colsToTest)>=2){
    colnames <- gsub(varInt,"",colnames(fit$design))
    for (comp in combn(length(colsToTest),2,simplify=FALSE)){ 
      contrast <- numeric(ncol(dge$design))
      contrast[colsToTest[comp[1:2]]] <- c(-1,1)
      namecomp <- paste0(colnames[colsToTest[comp[2]]],"_vs_",colnames[colsToTest[comp[1]]])
      cat(paste0("Comparison ",gsub("_"," ",namecomp),": testing contrast (",paste(contrast,collapse=", "),")"),"\n")
      lrt <- glmLRT(fit, contrast=contrast)
      results[[namecomp]] <- topTags(lrt,n=nrow(dge$counts),adjust.method=pAdjustMethod,sort.by="none")$table
    }
  }
  
  return(list(dge=dge,results=results,lrt=lrt))
}


out.edgeR <- run.edgeR(counts=counts, target=target, varInt=varInt, condRef=condRef,
                       batch=batch, cpmCutoff=cpmCutoff, normalizationMethod=normalizationMethod,
                       pAdjustMethod=pAdjustMethod)

#--------------------
# MDS + clustering (unchanged SARTools function)
#--------------------
exploreCounts(object=out.edgeR$dge, group=target[,varInt], gene.selection=gene.selection, col=colors)

#--------------------
# exporting results of the differential analysis (adapted from SARTools function)
#--------------------

rawpHist_AD <- function(complete_AD, outfile=TRUE){
  ncol <- ifelse(length(complete_AD)<=4, ceiling(sqrt(length(complete_AD))), 3)
  nrow <- ceiling(length(complete_AD)/ncol)
  
  par(mfrow=c(nrow,ncol))
  for (name in names(complete_AD)){
    ggplot(as.data.frame(complete_AD[[name]]), aes(x=pvalue, fill="tomato2")) + geom_histogram(binwidth = 0.025, color="black") + scale_fill_discrete(name = "", labels = "pvalues") + labs(title = "Raw pvalues histogram") + theme(plot.margin = margin(2,.8,2,.8, "cm")) 
  }
  if (outfile) ggsave(path= "figures/", filename="rawpHist_ggplot.pdf")
  if (outfile) ggsave(path= "figures/", filename="rawpHist.png")
  if (outfile) dev.off()
}


exportResults.edgeR <- function(out.edgeR, group, counts, alpha=0.05, export=TRUE){
  
  dge <- out.edgeR$dge
  res <- out.edgeR$results
  
  # raw count, normalised count and baseMean
  tmm <- dge$samples$norm.factors
  N <- colSums(dge$counts)
  f <- tmm * N/mean(tmm * N)
  normCounts <- round(scale(dge$counts, center=FALSE, scale=f))
  base <- data.frame(Id=rownames(counts), counts)
  names(base) <- c("Id", colnames(counts))
  norm.bm <- data.frame(Id=rownames(normCounts),normCounts)
  names(norm.bm) <- c("Id", paste0("norm.",colnames(normCounts)))
  norm.bm$baseMean <- round(apply(scale(dge$counts, center=FALSE, scale=f),1,mean),2)
  for (cond in levels(group)){
    norm.bm[,cond] <- round(apply(as.data.frame(normCounts[,group==cond]),1,mean),0)
  }
  base <- merge(base,norm.bm,by="Id",all=TRUE)
  
  complete <- list()
  for (name in names(res)){
    complete.name <- base
    
    # add info from res 
    res.name <- data.frame(Id=rownames(res[[name]]),FC=round(2^(res[[name]][,"logFC"]),3),
                           log2FoldChange=round(res[[name]][,"logFC"],3),pvalue=res[[name]][,"PValue"],
                           padj=res[[name]][,ifelse("FDR" %in% names(res[[name]]), "FDR", "FWER")])
    complete.name <- merge(complete.name, res.name, by="Id", all=TRUE)
    # add info from dge
    dge.add <- data.frame(Id=rownames(dge$counts),tagwise.dispersion=round(dge$tagwise.dispersion,4),
                          trended.dispersion=round(dge$trended.dispersion,4))
    complete.name <- merge(complete.name, dge.add, by="Id", all=TRUE)
    complete[[name]] <- complete.name
    
    if (export){
      # obtain sartools and restricted lists of differentially open and closed regions durong brain development
      up.name <- complete.name[which(complete.name$padj <= alpha & complete.name$log2FoldChange>=0),]
      up.name <- up.name[order(up.name$padj),]
      down.name <- complete.name[which(complete.name$padj <= alpha & complete.name$log2FoldChange<=0),]
      down.name <- down.name[order(down.name$padj),]		
      up.name.strict <- complete.name[which(complete.name$padj <= alpha & complete.name$log2FoldChange>=1),]
      up.name.strict <- up.name.strict[order(up.name.strict$padj),]
      down.name.strict <- complete.name[which(complete.name$padj <= alpha & complete.name$log2FoldChange<=-1),]
      down.name.strict <- down.name.strict[order(down.name.strict$padj),]	
      for (i in list("up.name", "down.name","up.name.strict","down.name.strict")){
        dimension = nrow(get(i))
        print(paste("number of rows :", i, "=", dimension,sep=" "))
      }
      
      # exports
      name <- gsub("_","",name)
      write.table(complete.name, file=paste0("tables/",name,".complete.txt"), sep="\t", row.names=FALSE, dec=".", quote=FALSE)
      write.table(up.name, file=paste0("tables/", name,".up.txt"), row.names=FALSE, sep="\t", dec=".", quote=FALSE)
      write.table(down.name, file=paste0("tables/", name,".down.txt"), row.names=FALSE, sep="\t", dec=".", quote=FALSE)
      write.table(up.name.strict, file=paste0("tables/", name,".up.strict.txt"), row.names=FALSE, sep="\t", dec=".", quote=FALSE)
      write.table(down.name.strict, file=paste0("tables/", name,".down.strict.txt"), row.names=FALSE, sep="\t", dec=".", quote=FALSE)
    }
  }
  
  return(complete)
}


# summary of the analysis (boxplots, dispersions, export table, nDiffTotal, histograms, MA plot)
summarizeResults.edgeR.AD = function (out.edgeR, group, counts, alpha = 0.05, col = c("lightblue", 
                                                                                      "orange", "MediumVioletRed", "SpringGreen"), log2FClim = NULL, 
                                      padjlim = NULL) 
{
  if (!I("figures" %in% dir())) 
    dir.create("figures", showWarnings = FALSE)
  if (!I("tables" %in% dir())) 
    dir.create("tables", showWarnings = FALSE)
  countsBoxplots(out.edgeR$dge, group = group, col = col)
  BCVPlot(dge = out.edgeR$dge)
  complete <- exportResults.edgeR(out.edgeR = out.edgeR, group = group, 
                                  counts = counts, alpha = alpha)
  nDiffTotal <- nDiffTotal(complete = complete, alpha = alpha)
  cat("Number of features down/up and total:\n")
  print(nDiffTotal, quote = FALSE)
  rawpHist_AD(complete_AD = complete, outfile = TRUE)
  MAPlot(complete = complete, alpha = alpha, log2FClim = log2FClim)
  volcanoPlot(complete = complete, alpha = alpha, padjlim = padjlim)
  return(list(complete = complete, nDiffTotal = nDiffTotal))
}


summaryResults.AD <- summarizeResults.edgeR.AD(out.edgeR, group=target[,varInt], counts=counts, alpha=alpha, col=colors)

#--------------------
# save image of the R session (unchanged SARTools script)
#--------------------
save.image(file=paste0(projectName, ".RData"))

#--------------------
# generating HTML report (adapted from SARTools function)
#--------------------
writeReport.edgeR(target=target, counts=counts, out.edgeR=out.edgeR, summaryResults=summaryResults.AD,
                  majSequences=majSequences, workDir=workDir, projectName=projectName, author=author,
                  targetFile=targetFile, rawDir=rawDir, featuresToRemove=featuresToRemove, varInt=varInt,
                  condRef=condRef, batch=batch, alpha=alpha, pAdjustMethod=pAdjustMethod, cpmCutoff=cpmCutoff,
                  colors=colors, gene.selection=gene.selection, normalizationMethod=normalizationMethod)

#--------------------
# save FDR and logCPM information (new in the script)
#--------------------

if (length(names(out.edgeR$results))){
  write.table(out.edgeR$results, file="tables/results_FDR_logCPM_all_filter.txt", sep="\t", row.names=FALSE, dec=".", quote=FALSE)
}else{
  print("Certaines données ne seront pas sauvegardées")
}


# ---------
# ----------------------
# --------------------------------------------
# Same code for E14 vs E15 developmental stages comparison with following parameters setting
# --------------------------------------------
# ----------------------
# ---------

dir.create("G:/projet_DU_DSD/results/09_all_peaks_count_table/dup_pool/SARTools/Analyse_L/E14.15", recursive=TRUE)

rm(list=ls())                                        

workDir <- "G:/projet_DU_DSD/results/09_all_peaks_count_table/dup_pool/SARTools/Analyse_L/E14.15"      # working directory for the R session

projectName <- "Sartools-edgeR-E14.15"               # name of the project
author <- "Agathe D."                                # author of the statistical analysis/report

targetFile <- "G:/projet_DU_DSD/results/09_all_peaks_count_table/dup_pool/raw_data/targetE14.15.txt"                           # path to the design/target file
rawDir <- "G:/projet_DU_DSD/results/09_all_peaks_count_table/dup_pool/raw_data/"                                      # path to the directory containing raw counts files
featuresToRemove <- c("alignment_not_unique",        # names of the features to be removed
                      "ambiguous", "no_feature",     # (specific HTSeq-count information and rRNA for example)
                      "not_aligned", "too_low_aQual")# NULL if no feature to remove

varInt <- "group"                                    # factor of interest
condRef <- "E14"                                     # reference biological condition
batch <- NULL                                        # blocking factor: NULL (default) or "batch" for example

alpha <- 0.05                                        # threshold of statistical significance
pAdjustMethod <- "BH"                                # p-value adjustment method: "BH" (default) or "BY"

cpmCutoff <- 1                                       # counts-per-million cut-off to filter low counts
gene.selection <- "pairwise"                         # selection of the features in MDSPlot
normalizationMethod <- "TMM"                         # normalization method: "TMM" (default), "RLE" (DESeq) or "upperquartile"

colors <- c("darkblue","brown3")               

forceCairoGraph <- FALSE

nsamples = 4                                        # nb of samples 
nstage = 2                                          # nb of developemental stages
nrep = 2                                            # nb of replicates per developmental stages
couleur=c("cyan4","brown3","darkgreen","darkblue","plum4","lightgoldenrod3","pink")


# ---------
# ----------------------
# --------------------------------------------
# Same code for E15 vs E16 developmental stages comparison with following parameters setting
# --------------------------------------------
# ----------------------
# ---------

dir.create("G:/projet_DU_DSD/results/09_all_peaks_count_table/dup_pool/SARTools/Analyse_L/E15.16", recursive=TRUE)

rm(list=ls())                                        

workDir <- "G:/projet_DU_DSD/results/09_all_peaks_count_table/dup_pool/SARTools/Analyse_L/E15.16"      # working directory for the R session

projectName <- "Sartools-edgeR-E15.16"               # name of the project
author <- "Agathe D."                                # author of the statistical analysis/report

targetFile <- "G:/projet_DU_DSD/results/09_all_peaks_count_table/dup_pool/raw_data/targetE15.16.txt"                           # path to the design/target file
rawDir <- "G:/projet_DU_DSD/results/09_all_peaks_count_table/dup_pool/raw_data/"                                      # path to the directory containing raw counts files
featuresToRemove <- c("alignment_not_unique",        # names of the features to be removed
                      "ambiguous", "no_feature",     # (specific HTSeq-count information and rRNA for example)
                      "not_aligned", "too_low_aQual")# NULL if no feature to remove

varInt <- "group"                                    # factor of interest
condRef <- "E15"                                     # reference biological condition
batch <- NULL                                        # blocking factor: NULL (default) or "batch" for example

alpha <- 0.05                                        # threshold of statistical significance
pAdjustMethod <- "BH"                                # p-value adjustment method: "BH" (default) or "BY"

cpmCutoff <- 1                                       # counts-per-million cut-off to filter low counts
gene.selection <- "pairwise"                         # selection of the features in MDSPlot
normalizationMethod <- "TMM"                         # normalization method: "TMM" (default), "RLE" (DESeq) or "upperquartile"

colors <- c("darkblue","brown3")               

forceCairoGraph <- FALSE

nsamples = 4                                        # nb of samples 
nstage = 2                                          # nb of developemental stages
nrep = 2                                            # nb of replicates per developmental stages
couleur=c("cyan4","brown3","darkgreen","darkblue","plum4","lightgoldenrod3","pink")


# ---------
# ----------------------
# --------------------------------------------
# Same code for E16 vs P0 developmental stages comparison with following parameters setting
# --------------------------------------------
# ----------------------
# ---------

dir.create("G:/projet_DU_DSD/results/09_all_peaks_count_table/dup_pool/SARTools/Analyse_L/E16.P0", recursive=TRUE)

rm(list=ls())                                        

workDir <- "G:/projet_DU_DSD/results/09_all_peaks_count_table/dup_pool/SARTools/Analyse_L/E16.P0"      # working directory for the R session

projectName <- "Sartools-edgeR-E16.P0"               # name of the project
author <- "Agathe D."                                # author of the statistical analysis/report

targetFile <- "G:/projet_DU_DSD/results/09_all_peaks_count_table/dup_pool/raw_data/targetE16.P0.txt"                           # path to the design/target file
rawDir <- "G:/projet_DU_DSD/results/09_all_peaks_count_table/dup_pool/raw_data/"                                      # path to the directory containing raw counts files
featuresToRemove <- c("alignment_not_unique",        # names of the features to be removed
                      "ambiguous", "no_feature",     # (specific HTSeq-count information and rRNA for example)
                      "not_aligned", "too_low_aQual")# NULL if no feature to remove

varInt <- "group"                                    # factor of interest
condRef <- "E16"                                     # reference biological condition
batch <- NULL                                        # blocking factor: NULL (default) or "batch" for example

alpha <- 0.05                                        # threshold of statistical significance
pAdjustMethod <- "BH"                                # p-value adjustment method: "BH" (default) or "BY"

cpmCutoff <- 1                                       # counts-per-million cut-off to filter low counts
gene.selection <- "pairwise"                         # selection of the features in MDSPlot
normalizationMethod <- "TMM"                         # normalization method: "TMM" (default), "RLE" (DESeq) or "upperquartile"

colors <- c("darkblue","brown3")               

forceCairoGraph <- FALSE

nsamples = 4                                        # nb of samples 
nstage = 2                                          # nb of developemental stages
nrep = 2                                            # nb of replicates per developmental stages
couleur=c("cyan4","brown3","darkgreen","darkblue","plum4","lightgoldenrod3","pink")
```

#### Obtention of chromosomal coordinates of each region¶

To associate a chromosomal position at each peak name of region dectected as differentially open or closed regions during brain development, combination of different files is necessary :

In [ ]:

```
%%bash 

# 1. Obtain Region ID
# Keep only ENSEMBL ID of regions dectected as differentially open or closed
cd /mnt/g/projet_DU_DSD/results/09_all_peaks_count_table/dup_pool/SARTools/Analyse_L/E13.14/tables
sed '1d' E14vsE13.down.txt | cut -d$'\t' -f1 > Peak_ID_down_DOCR_E13.14_mm9.txt
sed '1d' E14vsE13.up.txt | cut -d$'\t' -f1 > Peak_ID_up_DOCR_E13.14_mm9.txt

cd /mnt/g/projet_DU_DSD/results/09_all_peaks_count_table/dup_pool/SARTools/Analyse_L/E14.15/tables
sed '1d' E15vsE14.down.txt | cut -d$'\t' -f1 > Peak_ID_down_DOCR_E14.15_mm9.txt
sed '1d' E15vsE14.up.txt | cut -d$'\t' -f1 > Peak_ID_up_DOCR_E14.15_mm9.txt

cd /mnt/g/projet_DU_DSD/results/09_all_peaks_count_table/dup_pool/SARTools/Analyse_L/E15.16/tables
sed '1d' E16vsE15.down.txt | cut -d$'\t' -f1 > Peak_ID_down_DOCR_15.E16_mm9.txt
sed '1d' E16vsE15.up.txt | cut -d$'\t' -f1 > Peak_ID_up_DOCR_E15.16_mm9.txt

cd /mnt/g/projet_DU_DSD/results/09_all_peaks_count_table/dup_pool/SARTools/Analyse_L/E14.16/tables
sed '1d' E16vsE14.down.txt | cut -d$'\t' -f1 > Peak_ID_down_DOCR_14.E16_mm9.txt
sed '1d' E16vsE14.up.txt | cut -d$'\t' -f1 > Peak_ID_up_DOCR_E14.16_mm9.txt

cd /mnt/g/projet_DU_DSD/results/09_all_peaks_count_table/dup_pool/SARTools/Analyse_L/E16.P0/tables
sed '1d' POvsE16.down.txt | cut -d$'\t' -f1 > Peak_ID_down_DOCR_E16.P0_mm9.txt
sed '1d' POvsE16.up.txt | cut -d$'\t' -f1 > Peak_ID_up_DOCR_E16.P0_mm9.txt


# 2. Combine ENSEMBL ID wih .gff file containing information of regions/peaks detected by MACS2

# 2.1 dataset 1 (= edgeR output) formatting
# Remove header 
cd  /mnt/g/projet_DU_DSD/results/09_all_peaks_count_table/dup_pool/SARTools/Analyse_L/E13.14/tables
sed '1d' E14vsE13.down.txt > nh_down_DOCR_E13.14_mm9.txt
sed '1d' E14vsE13.up.txt > nh_up_DOCR_E13.14_mm9.txt

cd  /mnt/g/projet_DU_DSD/results/09_all_peaks_count_table/dup_pool/SARTools/Analyse_L/E14.15/tables
sed '1d' E15vsE14.down.txt > nh_down_DOCR_E14.15_mm9.txt
sed '1d' E15vsE14.up.txt > nh_up_DOCR_E14.15_mm9.txt

cd  /mnt/g/projet_DU_DSD/results/09_all_peaks_count_table/dup_pool/SARTools/Analyse_L/E15.16/tables
sed '1d' E16vsE15.down.txt > nh_down_DOCR_E15.16_mm9.txt
sed '1d' E16vsE15.up.txt > nh_up_DOCR_E15.16_mm9.txt

cd  /mnt/g/projet_DU_DSD/results/09_all_peaks_count_table/dup_pool/SARTools/Analyse_L/E14.16/tables
sed '1d' E16vsE14.down.txt > nh_down_DOCR_E14.16_mm9.txt
sed '1d' E16vsE14.up.txt > nh_up_DOCR_E14.16_mm9.txt

cd  /mnt/g/projet_DU_DSD/results/09_all_peaks_count_table/dup_pool/SARTools/Analyse_L/E16.P0/tables
sed '1d' POvsE16.down.txt > nh_down_DOCR_E16.P0_mm9.txt
sed '1d' POvsE16.up.txt > nh_up_DOCR_E16.P0_mm9.txt

#copy and paste files in /mnt/g/projet_DU_DSD/results/10_annotations/Sartools/Biomart/raw_data folder

mkdir -p /mnt/g/projet_DU_DSD/results/10_annotations/Sartools/Biomart/raw_data
```

In [ ]:

```
# 2.2 Load dataset 1 in R environment 

# List containing "Peaks_ID" files
list_files = list.files("G:/projet_DU_DSD/results/10_annotations/Sartools/Biomart/raw_data/")
list_files_Peak = list_files[which(grepl("nh",list_files))]

# 2.3 Load dataset 2 (= .gff file containing informations of regions/peaks detected by MACS2) in R environment 

my_gff = read.table("G:/projet_DU_DSD/results/08_all_peaks_merge/dup_pool/nh_E13_a_P0_all_peaks_dup_pool_merge_sorted_Unix_tab.gff",
                    sep = "\t", header = FALSE)

# Verifications 
dim(my_gff)
head(my_gff)
colnames(my_gff) = c("chr","source","element_type","start","end","V6","strand","V8","V9", "ID")

# 2.4 Combine informations from datasets 1 and 2

for(i in 1:10){
  ID_sd = list_files_Peak[i]
  sartools_file = read.table(file = paste("G:/projet_DU_DSD/results/10_annotations/Sartools/Biomart/raw_data/",ID_sd,sep="/"), header = FALSE, sep = "\t", quote="", dec=".")
  
  colnames(sartools_file) = c("ID",	"cond1_rep1",	"cond1_rep2",	"cond2_rep1", "cond2_rep2",	"norm.cond1_rep1",	"norm.cond1_rep2",	"norm.cond2_rep1",	"norm.cond2_rep2",	"baseMean",	"cond1",	"cond2",	"FC",	"log2FoldChange",	"pvalue",	"padj",	"tagwise.dispersion",	"trended.dispersion")
  merge_dataset = merge(sartools_file,my_gff, by="ID")
  print(dim(sartools_file))
  print(dim(merge_dataset))
  
  write.table(merge_dataset, file=paste("G:/projet_DU_DSD/results/10_annotations/Sartools/Biomart/gff_ID",ID_sd, ".txt",sep="_"), quote=FALSE, sep="\t", row.names = FALSE, col.names = TRUE)
}
```

#### Conversion of regions coordinates : mm9 to mm10¶

In order to combine ATAC-seq data with results of others analyses, mm9 coordinates of differentially open or closed regions were converted into mm10 coordinates, using **[LiftOver]**(https://genome.ucsc.edu/cgi-bin/hgLiftOver) tool proposed by **UCSC**. For that, only chromosomal coordinates (chromosome-start-end) are kept to generate a .bed file.

In [ ]:

```
%%bash

mkdir /mnt/g/projet_DU_DSD/results/10_annotations/Sartools/Biomart/mm10

cd /mnt/g/projet_DU_DSD/results/10_annotations/Sartools/Biomart/

# Samples name
sample="E13.14 E14.15 E15.16 E16.P0 E14.16"

# keep only bed information (chr - start - end) and sort the .bed file
for file in $sample
do
echo "${file}"
sed '1d' gff_ID_nh_down_DOCR_${file}_mm9.txt | awk '{ print $19"\t"$22"\t"$23 }' | sort -k1,1 -k2,2n > bed_gff_ID_down_DOCR_${file}_mm9.sorted.txt

sed '1d' gff_ID_nh_up_DOCR_${file}_mm9.txt | awk '{ print $19"\t"$22"\t"$23 }' | sort -k1,1 -k2,2n > bed_gff_ID_up_DOCR_${file}_mm9.sorted.txt

echo "bed_gff_ID_down_DOCR_${file}_mm9.sorted.txt"
head bed_gff_ID_down_DOCR_${file}_mm9.sorted.txt
echo "bed_gff_ID_up_DOCR_${file}_mm9.sorted.txt"
head bed_gff_ID_up_DOCR_${file}_mm9.sorted.txt

done
```

Default settings of **LiftOver** tool have been retained (*i.e*. at least 0.95 as the minimum ratio of bases that must remap). Some regions can't be converted for distinct reasons:

In [ ]:

```
# For E14.15 up : 
#Partially deleted in new (Sequence insufficiently intersects one chain)
chr9	100999669	101000211

# For E16.P0 up (6 failed) :
#Deleted in new (Sequence intersects no chains)
chr18	11406894	11407504
#Partially deleted in new (Sequence insufficiently intersects one chain)
chr4	130185092	130185630
#Partially deleted in new (Sequence insufficiently intersects one chain)
chrUn_random	126830	127059
#Split in new (Sequence insufficiently intersects multiple chains)
chrUn_random	3318185	3318608
#Split in new (Sequence insufficiently intersects multiple chains)
chrUn_random	3756019	3756421
#Deleted in new (Sequence intersects no chains)
chrUn_random	4062708	4064266
```

Files containing mm10 coordinates have been renamed (*e.g.* bed\_gff\_ID\_down\_DOCR\_E15.16\_mm10.bed), then sorted and associated to mm9 coordinates to keep both annotations.

In [ ]:

```
%%bash 

cd /mnt/g/projet_DU_DSD/results/10_annotations/Sartools/Biomart/mm10

# 1. Sort mm10 .bed files
for file in bed*
do  sort -k1,1 -k2,2n $file > ${file/.bed/.sorted.bed}
echo $file
done

# 2. Manually remove from mm9 regions, those that are missing in mm10 annotation
# For E14.15 up --> bed_gff_ID_up_DOCR_E14.15_mm9_sans_reg_perdu_mm10.sorted.txt
# For E16.P0 up --> bed_gff_ID_up_DOCR_E16.P0_mm9_sans_reg_perdu_mm10.sorted.txt

# 3. Combine mm9 and mm10 coordinates : first coordinate = mm9, second = mm10
cd /mnt/g/projet_DU_DSD/results/10_annotations/Sartools/Biomart/mm10

sample="down_DOCR_E13.14 down_DOCR_E14.15 down_DOCR_E14.16 down_DOCR_E15.16 down_DOCR_E16.P0 up_DOCR_E13.14 up_DOCR_E14.16 up_DOCR_E15.16"

for file in $sample
do
    echo "$file"
    paste /mnt/g/projet_DU_DSD/results/10_annotations/Sartools/Biomart/bed_gff_ID_${file}_mm9.sorted.txt bed_gff_ID_${file}_mm10.sorted.bed > mm9_mm10_correspondance_${file}.txt
done 

sample_bis="up_DOCR_E14.15 up_DOCR_E16.P0"

for file in $sample_bis
do
    echo "$file"
    paste /mnt/g/projet_DU_DSD/results/10_annotations/Sartools/Biomart/bed_gff_ID_${file}_mm9_sans_reg_perdu_mm10.sorted.txt bed_gff_ID_${file}_mm10.sorted.bed > mm9_mm10_correspondance_${file}.txt
done
```

#### Annotation of differentially open or closed regions during brain development¶

Syntaxic annotation of regions of interest was performed using **bedtools intersect** with an annotation file (biomart\_mm10.txt) obtained with **BiomaRt** R package. This file contains :

- ensembl\_gene\_id : gene name
- chromosome\_name : chromosome/scaffold number
- strand
- start\_position : gene start (bp)
- end\_position : gene end (bp)
- entrezgene\_id :NCBI gene ID
- gene\_biotype : gene type
- mgi\_symbol : MGI symbol
- entrezgene\_accession : NCBI gene accession
- entrezgene\_description : NCBI gene description
- uniprot\_gn\_symbol : UniProtKB gene name symbol

In [ ]:

```
#--------------------------------------
# 1. Obtention of biomart_mm10.txt file
#--------------------------------------

# Load BiomaRt package : 
library("biomaRt")

# 1. Select database and reference genome 
mm10 = useMart("ensembl", dataset="mmusculus_gene_ensembl")
# mm10 Mus musculus version used:  Ensembl 97 Jul 2019 http://jul2019.archive.ensembl.org 


# 2. Create the dataset with information of interest
# It's necessary to do it in two steps, because there is a limitation in the number of attributes that can be collected at the same time.
annot_mm10_part1<-getBM(attributes=c("ensembl_gene_id","chromosome_name","strand", "start_position","end_position","entrezgene_id","gene_biotype","mgi_symbol","entrezgene_accession"), mart=mm10)
annot_mm10_part2<-getBM(attributes=c("ensembl_gene_id","entrezgene_description","uniprot_gn_symbol"), mart=mm10)
annot_mm10 = merge(annot_mm10_part1, annot_mm10_part2, by="ensembl_gene_id", all=TRUE)

# Verifications
dim(annot_mm10)
head(annot_mm10)

# Backup
write.table(annot_mm10,"G:/Chip_pilote/data/mm10_genome/biomart_mm10.txt", quote= FALSE, sep="\t",row.names=FALSE)


#--------------------------------------
# 2. Combine differentially open or closed regions with biomart annotation file 
#--------------------------------------
# 2.1 Datasets formatting 
setwd("G:/projet_DU_DSD/results/")
biomart_mm10 =read.table("G:/projet_DU_DSD/results/10_annotations/Sartools/Biomart/mm10/biomart_mm10_sorted.txt",sep="\t", na.strings = "NA", fill=TRUE, quote="", header=TRUE)

head(biomart_mm10)

# Change "-1" and "1" strand encoding by "-" and "+"
for (i in 1:nrow(biomart_mm10)){
  if(biomart_mm10[i,"strand"]==1){
    biomart_mm10[i,"strand"]="+"
  }else if (biomart_mm10[i,"strand"]==-1){
    biomart_mm10[i,"strand"]="-"
  }else{
  }
}

# Change chromosome format  
biomart_mm10$chr = with(biomart_mm10,paste("chr",biomart_mm10[,"chromosome_name"],sep=""))

head(biomart_mm10)

# To combine informations, we will use bedtools intersect. It seems to work with a .bed containing exactly 9 columns. 
# uniprot_genename and entrezgene_description informations are removed, because bedtools intersect doesn't work if these columns are kept.
# (bedtools intersect is not able to detect the file format if these columns are kept). 

biomart_mm10_bis=biomart_mm10[,c("chr","start_position","end_position","strand","entrezgene_accession", "gene_biotype", "mgi_symbol","ensembl_gene_id", "entrezgene_id")] #it works

write.table(biomart_mm10_bis,"G:/projet_DU_DSD/results/10_annotations/Sartools/Biomart/mm10/biomart_mm10_all_info.bed", sep = "\t", col.names=TRUE, row.names=FALSE, quote=FALSE)
```

In [ ]:

```
%%bash

# 2.2 Sort files
# Sort Biomart annotation file
cd /mnt/g/projet_DU_DSD/results/10_annotations/Sartools/Biomart/mm10/

sed '1d' biomart_mm10_all_info.bed | sort -k1,1 -k2,2n > biomart_mm10_all_info_sorted.bed

head biomart_mm10_all_info_sorted.bed


# Sort mm10 .bed files containing differentially open or closed regions coordinates
# Already done, I copy and paste files into  /mnt/g/projet_DU_DSD/results/10_annotations/Sartools/Biomart/mm10/raw_data


#----------------------------------
# 3. Combine information using bedtools intersect
#----------------------------------

cd /mnt/g/projet_DU_DSD/results/10_annotations/Sartools/Biomart/mm10/raw_data

sample="down_DOCR_E13.14 down_DOCR_E14.15 down_DOCR_E14.16 down_DOCR_E15.16 down_DOCR_E16.P0 up_DOCR_E13.14  up_DOCR_E14.15 up_DOCR_E14.16 up_DOCR_E15.16 up_DOCR_E16.P0"

for file in $sample
do 
    echo "$file"
    bedtools intersect -wao -a bed_gff_ID_${file}_mm10.sorted.bed -b /mnt/g/projet_DU_DSD/results/10_annotations/Sartools/Biomart/mm10/biomart_mm10_all_info_sorted.bed > /mnt/g/projet_DU_DSD/results/10_annotations/Sartools/Biomart/mm10/${file}_annot_syntaxique_mm10_biomart.bed

    echo "fichier : $file annoté"
    wc -l /mnt/g/projet_DU_DSD/results/10_annotations/Sartools/Biomart/mm10/${file}_annot_syntaxique_mm10_biomart.bed
    head /mnt/g/projet_DU_DSD/results/10_annotations/Sartools/Biomart/mm10/${file}_annot_syntaxique_mm10_biomart.bed
done
```

#### Combination of annotation and statistical informations¶

In order to obtain a file where each region of interest is represented by a single line, the annotation elements of a given region that are on separated lines have been grouped together. Then, all data (statistical informations, mm9 and mm10 coordinates regions, annotation informations) are combined in a single file (one file per pairwise comparison).

In [ ]:

```
# Load file : List file containing "AnnotBM" or "ID_sd" patterns
list_files = list.files("G:/projet_DU_DSD/results/10_annotations/Sartools/Biomart/mm10/")
list_files_Annot = list_files[which(grepl("annot_syntaxique",list_files))]
list_files_corresp = list_files[which(grepl("correspondance",list_files))]

list_files_bis = list.files("G:/projet_DU_DSD/results/10_annotations/Sartools/Biomart/")
list_files_ID_sd = list_files_bis[which(grepl("gff_ID",list_files_bis))]


for(i in 1:10){
  Annot = list_files_Annot[i]
  DOCR_annot = read.table(file = paste("G:/projet_DU_DSD/results/10_annotations/Sartools/Biomart/mm10",Annot,sep="/"), header = FALSE, sep = "\t", quote="", fill=TRUE)
  
  #------------------------------------------------------------------------
  # A. Concatenate annotation informations : one row = one region of interest
  # 1. Create the ID = chr-start-end
  colnames(DOCR_annot) = c("chr", "start_DOCR","end_DOCR","chr_gene", "start_gene", "end_gene", "strand","entrezgene_accession",
                           "gene_biotype","mgi_symbol", "ensembl_gene_id", "entrezgene_id", "overlap_length")

  DOCR_annot$ID = with(DOCR_annot, paste(DOCR_annot[,"chr"], DOCR_annot[,"start_DOCR"],DOCR_annot[,"end_DOCR"],sep="-"))
  
  # 2. Concatenate informations 
  library(dplyr)
  
  # Liste of unique ID  
  liste_DOCR_unique <- unique(DOCR_annot$ID)
  
  # empty matrice which will be filled during the loop
  # Nb of rows = nb of unique ID | Nb of columns = nb of variables
  treatment_file <- data.frame(matrix(NA,nrow=length(liste_DOCR_unique),ncol=ncol(DOCR_annot)))
  colnames(treatment_file) <- colnames(DOCR_annot)
  
  treatment_file$ID <- liste_DOCR_unique
  for(k in 1:length(liste_DOCR_unique)){
    # Filter according to ID k 
    DOCR_filt <- DOCR_annot %>%
      filter(ID == liste_DOCR_unique[k])
    
    # Collapsing of distincts elements of a variables for ID k 
    # If variable is composed of only one element ==> we keep this element only once 
    # If variable is composed of m distinct elements ==> even if elements can be repeated, we keep all the elements (with repetitions)
    for(j in 1:length(colnames(treatment_file))){
      if(length(unique(DOCR_filt[,j])) == 1){
        treatment_file[k,j] <- paste(unique(DOCR_filt[,j]), collapse=';')
      } else {
        treatment_file[k,j] <- paste(DOCR_filt[,j], collapse=';')  
      }
    }
  }
  print(paste("nb of uniq ID treatment file : ", nrow(treatment_file), sep=""))
  
  
  
  #------------------------------------------------------------------------
  # B. Merge treatment_file with mm9 and mm10 regions coordinates 
  
  ID_corresp = list_files_corresp[i]
  corresp_file = read.table(file = paste("G:/projet_DU_DSD/results/10_annotations/Sartools/Biomart/mm10",ID_corresp,sep="/"), header = FALSE, sep = "\t", quote="", dec=".")
  
  colnames(corresp_file) = c("mm9_chr", "mm9_start", "mm9_end", "mm10_chr", "mm10_start", "mm10_end")
  corresp_file$ID = with(corresp_file, paste(corresp_file[,"mm10_chr"], corresp_file[,"mm10_start"], corresp_file[,"mm10_end"], sep="-"))
  corresp_file$mm9_ID = with(corresp_file, paste(corresp_file[,"mm9_chr"], corresp_file[,"mm9_start"], corresp_file[,"mm9_end"], sep="-"))

  ATAC_corresp = merge(corresp_file,treatment_file, by="ID", all = TRUE)
  
  size_Annot = nrow(treatment_file)
  size_ATAC_corresp = nrow(ATAC_corresp)
  
  if(size_Annot != size_ATAC_corresp){
    print(paste("Les deux fichiers", Annot, "et", ID_corresp, "n'ont pas la meme taille !", sep =" "))
  }else{
  }
  
  #------------------------------------------------------------------------
  # C. Merge ATAC_corresp with statistical informations obtained with SARTools-edgeR for every regions
  
  ID_sd = list_files_ID_sd[i]
  sartools_file = read.table(file = paste("G:/projet_DU_DSD/results/10_annotations/Sartools/Biomart",ID_sd,sep="/"), header = TRUE, sep = "\t", quote="", dec=".")
  
  colnames(sartools_file) = c("mm9_ID",	"cond1_rep1",	"cond1_rep2",	"cond2_rep1", "cond2_rep2",	
                              "norm.cond1_rep1",	"norm.cond1_rep2",	"norm.cond2_rep1",	"norm.cond2_rep2",	"baseMean",	
                              "cond1",	"cond2",	"FC",	"log2FoldChange",	"pvalue",	"padj",	"tagwise.dispersion",	"trended.dispersion", 
                              "mm9_chr", "source", "element_type", "mm9_start", "mm9_end", "V6", "strand", "V8", "V9")
  sartools_file$mm9_ID = with(sartools_file, paste(sartools_file[,"mm9_chr"], sartools_file[,"mm9_start"], sartools_file[,"mm9_end"], sep="-"))
  
  Annot_DOCR = merge(ATAC_corresp,sartools_file, by="mm9_ID", all = TRUE)
  Annot_DOCR_clean = Annot_DOCR[,c("mm9_chr.x", "mm9_start.x", "mm9_end.x", "mm9_ID", "mm10_chr", "mm10_start", "mm10_end", "ID", 
                                "chr_gene", "start_gene", "end_gene", "strand.x", "entrezgene_accession", "gene_biotype", "mgi_symbol", 
                              "ensembl_gene_id", "entrezgene_id",  "overlap_length", "cond1_rep1", "cond1_rep2", "cond2_rep1", "cond2_rep2",
                              "norm.cond1_rep1", "norm.cond1_rep2", "norm.cond2_rep1", "norm.cond2_rep2", "baseMean", "cond1", "cond2",  
                              "FC", "log2FoldChange", "pvalue", "padj", "tagwise.dispersion", "trended.dispersion")]
  
  
  colnames(Annot_DOCR_clean) = c("mm9_chr", "mm9_start", "mm9_end", "mm9_ID", "mm10_chr", "mm10_start", "mm10_end", "mm10_ID", 
                                 "chr_gene", "start_gene", "end_gene", "strand", "entrezgene_accession", "gene_biotype", "mgi_symbol", 
                                 "ensembl_gene_id", "entrezgene_id",  "overlap_length", "cond1_rep1", "cond1_rep2", "cond2_rep1", "cond2_rep2",
                                 "norm.cond1_rep1", "norm.cond1_rep2", "norm.cond2_rep1", "norm.cond2_rep2", "baseMean", "cond1", "cond2",  
                                 "FC", "log2FoldChange", "pvalue", "padj", "tagwise.dispersion", "trended.dispersion")
  
  write.table(Annot_DOCR_clean, file=paste("G:/projet_DU_DSD/results/10_annotations/Sartools/Biomart/mm10/stats_mm9_mm10",Annot,sep="_"), quote=FALSE, sep="\t", row.names = FALSE, col.names = TRUE)
  
}
```
