## Supplementary material for "Immediate perturbations of DNA methylation and transcriptome upon acute prenatal alcohol exposure in the mouse developing brain cortex": Supp data Notebook 3 RNAseq data mining: Supp. data Notebook 3 - RNA-seq workflow - Duchateau et al..html

Supp. data 3 - RNA-seq workflow


### Supplementary data Notebook 3 - RNA-seq workflow: analysis of physiological modifications of gene expression levels throughout brain development in mice¶

##### *Agathe Duchateau*¶

#### Download ENCODE data¶

To study modifications of physiological gene expression levels troughout brain developement, we used available RNA-seq ENCODE data. Experiments were done using embryonic and newborn forebrains of mixed sex C57BL/6N mice (expriments of B. Wold, Caltech laboratory).
Developmental stages that were studied in replicates are :

- embryonic day **E13.5** (Encode accession : ENCFF235DNM,ENCFF959PSX)
- embryonic day **E14.5** (Encode accession : ENCFF270GKY + ENCFF460TCF, ENCFF126IRS + ENCFF748SRJ)
- embryonic day **E15.5** (Encode accession : ENCFF179JEC, ENCFF891HIX)
- embryonic day **E16.5** (Encode accession : ENCFF931IVO, ENCFF114DRT)
- and postnatal day 0 (**P0**, Encode accession : ENCFF037JQC + ENCFF358MFI, ENCFF447EXU + ENCFF458NWF).

These data were obtained with a **non strand specificity poly RNA-seq, from oligo-dT primed total RNA**. In brief, tissues were lysed and RNA was extracted using **Ambion mirVana method**. Fragmentation was performed by **tagmentation (Nextera)** and size selection was done using **SPRI beads**. For each cDNA sample, fragments of **100** nucleotides were single-ended sequenced on a Illumina HiSeq 2500 platform. For more explanations on experimental procedures, please see *Experiment Summary* on ENCODE database, pages available with ENCODE accession of samples cited above.

In [ ]:

```
%%bash 

# Download ENCODE .fastq.gz files 

cd /data/omics-school/aduchateau/DU_projet/data

# For E16.5 first replicate
wget https://www.encodeproject.org/files/ENCFF931IVO/@@download/ENCFF931IVO.fastq.gz 
# For E16.5 second replicate
wget https://www.encodeproject.org/files/ENCFF114DRT/@@download/ENCFF114DRT.fastq.gz 
# For E15.5 first replicate
wget https://www.encodeproject.org/files/ENCFF179JEC/@@download/ENCFF179JEC.fastq.gz 
# For E15.5 second replicate
wget https://www.encodeproject.org/files/ENCFF891HIX/@@download/ENCFF891HIX.fastq.gz 
# For E14.5 first replicate
wget https://www.encodeproject.org/files/ENCFF270GKY/@@download/ENCFF270GKY.fastq.gz 
# For E14.5 first replicate
wget https://www.encodeproject.org/files/ENCFF460TCF/@@download/ENCFF460TCF.fastq.gz
# For E14.5 second replicate
wget https://www.encodeproject.org/files/ENCFF126IRS/@@download/ENCFF126IRS.fastq.gz 
# For E14.5 second replicate
wget https://www.encodeproject.org/files/ENCFF748SRJ/@@download/ENCFF748SRJ.fastq.gz 
# For E13.5 first replicate
wget https://www.encodeproject.org/files/ENCFF235DNM/@@download/ENCFF235DNM.fastq.gz 
# For E13.5 second replicate
wget https://www.encodeproject.org/files/ENCFF959PSX/@@download/ENCFF959PSX.fastq.gz 
# For P0 first replicate
wget https://www.encodeproject.org/files/ENCFF358MFI/@@download/ENCFF358MFI.fastq.gz 
# For P0 first replicate
wget https://www.encodeproject.org/files/ENCFF037JQC/@@download/ENCFF037JQC.fastq.gz 
# For P0 first replicate
wget https://www.encodeproject.org/files/ENCFF458NWF/@@download/ENCFF458NWF.fastq.gz 
# For P0 second replicate
wget https://www.encodeproject.org/files/ENCFF447EXU/@@download/ENCFF447EXU.fastq.gz 

# MD5sum check 
# File containing MD5sum of .fastq.gz files
cat md5sum.txt : 
# f12d3f32677ecc0208756bf9e05f990b	ENCFF931IVO.fastq.gz
# e9b9aae34a242deb0e8ec4b39979fe8c	ENCFF114DRT.fastq.gz
# befdcab4ff5b2bbdc9717064c55c02ef	ENCFF179JEC.fastq.gz
# f2a836c3ca6b515f3ed60488be900244	ENCFF891HIX.fastq.gz
# 1f6d6da942e693019de92fe5ada06f2f	ENCFF270GKY.fastq.gz
# 9781073211592e4f141eaac7f3396a06	ENCFF460TCF.fastq.gz
# ba5e9274ddcf96218d5c152624c73cb3	ENCFF126IRS.fastq.gz
# e8071390598286da2a5c273629a24789	ENCFF748SRJ.fastq.gz
# 66a89656a63f96311f64c7a28cf9102a	ENCFF235DNM.fastq.gz
# 1707d6213d80ba3ea18e323dca07eb8d	ENCFF959PSX.fastq.gz
# e5f5ef9f88ef582526cf1a54023f5ad0	ENCFF037JQC.fastq.gz
# 1d708f7b64c8b98bb9855453d4e6f6d4	ENCFF358MFI.fastq.gz
# 5a07748fe5a29c6b4fad1ddc850df0a5	ENCFF447EXU.fastq.gz
# 35807543b590dc9a98135aea49a20145	ENCFF458NWF.fastq.gz

md5sum -c md5sum.txt
# --> OK for all samples
```

#### Setting up of bioinformatic tools in a conda environment¶

Following tools were installed in a conda environment to perform analysis of RNA-seq from embryonic and newborn mice forebrains, obtained in physiological conditions. Version of each tool is indicated. Since .fasta files are very large, indexation of the reference genome and mapping were done on a distant server (cluster/High Performance Computer, from Institut Français de Bioinformatique - IFB). R packages which are cited below were also used to analyse the data.

In [ ]:

```
# Setting up of conda environment on a distant server (Ubuntu 16.04.5 LTS (GNU/Linux 4.4.0-131-generic x86_64))
conda create -n rnaseq
conda --version # version 4.5.8

conda activate rnaseq

conda install -y fastqc  #version 0.11.7 
conda install -y trim-galore #version 0.5.0
conda install -y cutadapt #version 1.16 

# Mapping on a cluster from Institut Français de Bioinformatique (IFB)
STAR  #version 2.6.1d

# Setting up of another conda environment on local computer
conda create -n DU_projet
conda --version # version 4.6.7
conda install -y htseq #version 0.11.2 
conda install -y samtools #version 1.9

# R version 3.5.2
# R package used for analysis : 
library(edgeR) # version 3.24.3
library(ggplot2) # version 3.2.1
library(reshape2) # version1.4.3
library(GGally) # version 1.4.0
library(SARTools) # version 1.6.9
library(BiomaRt) #version 2.38.0
library(dplyr) #version 0.8.3
library(data.table) # version 1.12.2
library(stringr) # version 1.4.0
```

#### Quality control of the raw data¶

After simplifying .fastq.gz file names, **FASTQC** was used to verify the quality of the sequencing raw data.

In [ ]:

```
%%bash

# To rename files 
cd /data/omics-school/aduchateau/DU_projet/data
cp ENCFF931IVO.fastq.gz ../results/01_raw_data/E16.5_01.fastq.gz
cp ENCFF114DRT.fastq.gz ../results/01_raw_data/E16.5_02.fastq.gz
cp ENCFF179JEC.fastq.gz ../results/01_raw_data/E15.5_01.fastq.gz
cp ENCFF891HIX.fastq.gz ../results/01_raw_data/E15.5_02.fastq.gz
cp ENCFF270GKY.fastq.gz ../results/01_raw_data/E14.5_01.1.fastq.gz
cp ENCFF460TCF.fastq.gz ../results/01_raw_data/E14.5_01.2.fastq.gz
cp ENCFF126IRS.fastq.gz ../results/01_raw_data/E14.5_02.1.fastq.gz
cp ENCFF748SRJ.fastq.gz ../results/01_raw_data/E14.5_02.2.fastq.gz
cp ENCFF235DNM.fastq.gz ../results/01_raw_data/E13.5_01.fastq.gz
cp ENCFF959PSX.fastq.gz ../results/01_raw_data/E13.5_02.fastq.gz
cp ENCFF037JQC.fastq.gz ../results/01_raw_data/E00.5_01.1.fastq.gz
cp ENCFF358MFI.fastq.gz ../results/01_raw_data/E00.5_01.2fastq.gz
cp ENCFF447EXU.fastq.gz ../results/01_raw_data/E00.5_02.1.fastq.gz
cp ENCFF458NWF.fastq.gz ../results/01_raw_data/E00.5_02.2fastq.gz


# Quality control of the raw data 

mkdir 02_FASTQC
cd ~/DU_projet/results/02_FASTQC

conda activate rnaseq

for file in ../01_raw_data/E*
do
fastqc $file
done

# To move files into appropriate folder
cd ~/DU_projet/results/01_raw_data

for file in *html
do
mv $file ../02_FASTQC/$file
done

for file in *zip
do
mv $file ../02_FASTQC/$file
done
```

#### Trimming¶

According to FASTQC reports, we decided to filter the data to remove remaining Nextera Transposase adapters, by using **Trim-galore**. The 15 first bases of all reads were also removed, because of their poor qualities. This trimming was sufficient for the majority of the samples. However some samples (first replicate of E14.5 stage + second replicate of P0 stage), having an over-representation of polyT or primers sequences (Clontech Universal Primer Mix Long), were trimmed again to remove these sequences.  
  
For E14 and P0 replicates, the two files for a given replicate were concatenated before trimming, using **cat unix command**. FASTQC controls were also performed on these concatenate files but reports were similar than those of separated files.
  
  
**Details of parameters used for the trimming:**

- --nextera : to remove Nextera adapters sequences
- -q 20 : to trim low-quality ends from reads in addition to adapter removal. Default value : 20
- --clip\_R1 15 : to remove 15 bases in 5' position
- --stringency 5 : overlap with adapter sequence required to trim a sequence. Default value : 1
- --length 20 : to remove reads that are shorter than 20 bases. Default value : 1
- -fastqc : to run a FASTQC report after the trimming

In [ ]:

```
%%bash

# ---------------------------------------------------
# To concatenate E14.5 and P0 files of a given replicate
# ---------------------------------------------------

cd ~/DU_projet/results/01_raw_data

cat E00.5_01.1.fastq.gz E00.5_01.2fastq.gz > E00.5_01.fastq.gz
cat E00.5_02.1.fastq.gz E00.5_02.2fastq.gz > E00.5_02.fastq.gz
cat E14.5_01.1.fastq.gz E14.5_01.2.fastq.gz > E14.5_01.fastq.gz
cat E14.5_02.1.fastq.gz E14.5_02.2.fastq.gz > E14.5_02.fastq.gz


# To run FASTQC on concatenate files
for file in E00.5_01.fastq.gz E00.5_02.fastq.gz E14.5_01.fastq.gz E14.5_02.fastq.gz
do
    fastqc $file --outdir ~/DU_projet/results/02_FASTQC/
done

# ---------------------------------------------------
# Trimming using trimming.bash script 
# First Round of trimming
# ---------------------------------------------------

# Script that was runned for trimming:
# Code will stop running if any problem is detected
# (first error, undefined variable, pipe error)
set -euo pipefail

# number of samples to analyse
sample="00.5_01 00.5_02 13.5_01 13.5_02 14.5_01 14.5_02 15.5_01 15.5_02 16.5_01 16.5_02"

for sample in ${sample}
do
    echo "=============================================================="
    echo "sample number : ${sample}"
    echo "=============================================================="

# Trimming 
    echo "=============================================================="
    echo "Trimming de l'échantillon ${sample}"
    echo "=============================================================="

# Use of Trim-galore to remove first bases, reads having bad quality score, 
# adapters sequences, too short reads... 

    trim_galore --nextera -q 20 --phred33 --clip_R1 15 --stringency 5 --length 20 
    -fastqc --output_dir ~/DU_projet/results/03_trimming ~/DU_projet/results/01_raw_data/E${sample}.fastq.gz 
    > tg_E${sample}.fastq.gz

done

# ---------------------------------------------------
# Second round of trimming for E14.5 (first replicate) P0 (second replicate)
# ---------------------------------------------------

mkdir -p ~/DU_projet/results/03_trimming/pass1/
for file in E14.5_01*
do
mv $file ~/Du_projet/results/03_trimming/pass1/
done

for file in E00.5_02*
do
mv $file ~/Du_projet/results/03_trimming/pass1/
done

# To remove polyT and Clontech Universal Primer Mix Long sequences
trim_galore -a TTTTTTTTTTTTTTTTTTTTTTTTTTTTTTTTTTTTTTTTTTTTTTTTTT -q 20 --phred33 
--stringency 40 --length 20 -fastqc 
--output_dir ~/DU_projet/results/03_trimming ~/DU_projet/results/03_trimming/pass1/E00.5_02_trimmed.fq.gz

trim_galore -a GTATCAACGCAGAGTACGGGAAGCAGTGGTATCAACGCAGAGTACGGGAA  -q 20 --phred33 
--stringency 40 --length 20 -fastqc 
--output_dir ~/DU_projet/results/03_trimming ~/DU_projet/results/03_trimming/pass1/E14.5_01_trimmed.fq.gz

# --------------------
# To save files
# --------------------

for file in *.gz
do 
    cp $file ~/DU_projet/results/04_mapping/input/tg_$file 
done

cd ~/DU_projet/results/04_mapping/input/
cp ~/DU_projet/results/03_trimming/E00.5_02_trimmed_trimmed.fq.gz ~/DU_projet/results/04_mapping/input/tg_E00.5_02_trimmed.fq.gz
cp ~/DU_projet/results/03_trimming/E14.5_01_trimmed_trimmed.fq.gz ~/DU_projet/results/04_mapping/input/tg_E14.5_01_trimmed.fq.gz
```

#### Mapping¶

Since fasta files are very large, indexation of the reference genome and mapping were done on a distant server (cluster/High Performance Computer, from Institut Français de Bioinformatique - IFB), with the help of O. Kirsh (Paris University, CNRS Epigenetics and Cell Fate, Paris, France).

##### Indexation of mm10 reference genome¶

Mapping was done using **STAR**, on mm10 Mus musculus reference genome, which was already available on the IFB cluster. When mm10 reference genome was indexed, we specified an annotation file (gencode.vM21.annotation.gtf), to improve accuracy of the mapping, as recommended in STAR user manual. This annotation file was downloaded on Gencode database (in Release M21 (GRCm38.p6) - Comprehensive gene annotation - CHR Regions, containing the comprehensive gene annotation on the reference chromosomes only).

**Details of parameters used for the indexation:**

- --runMode genomeGenerate : mode to index reference genome
- --runThreadN : number of threads to be used for genome indexation
- --genomeDir : to specify path to the directory where the genome indices are stored. The file system needs to have at least 100GB of disk.
- --genomeFastaFiles : to specify fasta file with the genome reference sequences.
- --sjdbGTFfile : to specify the path to the file with annotation in the standard GTF format. STAR will extract splice junctions from this file and use them to greatly improve accuracy of the mapping. While this is optional, and STAR can be run without annotations, using annotations is highly recommended whenever they are available.
- --sjdbGTFfeatureExon : to specify feature type in .gtf file that have to be used as exons for building transcripts
- --sjdbOverhang : to specify the length of the genomic sequence around the annotated junction to be used in constructing the splice junctions database. Ideally, this length should be equal to the ReadLength-1, where ReadLength is the length of the reads.

In [ ]:

```
%%bash

cd /Volumes/Maxtor/projet_DU_AD/data/mm10_genome/

# -------------------------------------
# Obtention of Gencode annotation file
# -------------------------------------

# Download Gencode Annotation file
wget ftp://ftp.ebi.ac.uk/pub/databases/gencode/Gencode_mouse/release_M21/gencode.vM21.annotation.gtf.gz

md5sum gencode.vM21.annotation.gtf.gz
#d08f66b2746d0ae66594fda6ea0c9939   gencode.vM21.annotation.gtf.gz
md5sum gencode.vM21.annotation.gtf
#699f4d30a09157711a3e71df018e52f5   gencode.vM21.annotation.gtf
```

In [ ]:

```
# Copy of this annotation file on IFB cluster

# -------------------------------------
# Genome indexation
# -------------------------------------
# Script that was runned for genome indexation:

#!/bin/bash
#
#SBATCH -o slurm.%N.%j.out
#SBATCH -e slurm.%N.%j.err
#SBATCH --mail-type END
#SBATCH --mail-user
#SBATCH --partition long
#SBATCH --cpus-per-task 12
#SBATCH --mem 40GB

module load conda

source  activate star-2.6

STAR --runMode genomeGenerate --runThreadN $SLURM_CPUS_PER_TASK 
--genomeDir /shared/home/okirsh/mapping/mm10_indexation 
--genomeFastaFiles /shared/bank/mus_musculus/mm10/fasta/mm10.fa 
--sjdbGTFfile /shared/home/okirsh/mapping/mm10_genome/gencode.vM21.annotation.gtf 
--sjdbGTFfeatureExon exon --sjdbOverhang 84

source activate
```

##### Mapping¶

Mapping of each sample was performed using **STAR** with the following parameters (see below). Mapping was relatively stringent, since each read can map at only 1 loci, otherwise it is not kept for the analysis (see --outFilterMultimapNmax option).
  
  
nb: For the first replicate of embryonic stage E14.5, we lost informations. Indeed, STAR didn't detect all the reads of this sample (47 515 325 reads were detected during the mapping step, whereas 50 697 328 reads were in the sample, according to FASTQC report). To verify if the file was corrupted during data transfert, we checked MD5sum of this file, but it was right. Because fasta file of this sample - which contains reads - are unsorted, we supposed that lost of reads was unbiased, because reads that were lost were randomly 'selected'. Thus, we decided to keep this sample for the rest of the analysis.
  
  
**Detail of parameters used for mapping:**

- --runMode alignReads : mode to map sample
- --runThreadN 6 : number of threads to run STAR
- --genomeDir : path containing indexed reference genome
- --genomeLoad NoSharedMemory : do not use shared memory, each job will have its own private copy of the genome
- --readFilesType Fastx : to notify that input file is in FASTA format
- --readFilesIn : path to input file
- --readFilesCommand gunzip -c : command line to execute for each of the input file. Here, we uncompressed input file using gunzip without deleting unzip file.
- --outSAMunmapped Within : to obtain a file containing unmapped reads
- --outReadsUnmapped Fastx : output of unmapped and partially mapped (*i.e.* mapped only one mate of a paired end read) reads in separate fasta/fastq file(s), called Unmapped.out.mate1/2.
- --outSAMtype BAM Unsorted : ouput file will be an unsorted .bam file
- --outSAMattributes Standard : string of desired SAM attributes in the order desired for the output. Standard =Standard NH HI AS nM.

***nb: it's impossible to use --outSAMattributes All option (All = NH HI AS nM NM MD jM jI) because file processing fails using htseq-count afterthat.***

- --outFilterType BySJout : type of filtering. Using BYSJout, we keep only those reads that contain junctions that passed filtering into SJ.out.tab
- --outFilterMultimapNmax 1 : maximum number of loci the read is allowed to map to. Alignments (all of them) will be output only if the read maps to no more loci than this value. Otherwise no alignments will be output, and the read will be counted as "mapped to too many loci" in the Log.final.out. Default: 10
- --outFilterMismatchNmax 10 : alignment will be output only if it has no more mismatches than this value. Default: 10
- --outFilterMismatchNoverLmax 0.3 : alignment will be output only if its ratio of mismatches to mapped length is less than or equal to this value. Default: 0.3
- --quantMode TranscriptomeSAM GeneCounts : types of quantification requested. Using TranscriptomeSAM : output SAM/BAM alignments to transcriptome into a separate file called Aligned.toTranscriptome.out.bam. Using GeneCounts : count reads per gene. A read is counted if it overlaps (1nt or more) one and only one gene. Both ends of the paired-end read are checked for overlaps. The counts coincide with those produced by htseq-count with default parameters. Both options can be used together.
- --twopassMode None : 1-pass mapping
- --outFileNamePrefix : to specify prefix name of output file.

In [ ]:

```
#--------------------------------------
#B. Mapping (using strict parameters)
#--------------------------------------

#Script that was runned for mapping: 
#!/bin/bash

for file in *.gz
do
    STAR --runMode alignReads --runThreadN 6 
    --genomeDir /home/olivier/Bureau/agathe/mm10_indexation --genomeLoad NoSharedMemory 
    --readFilesType Fastx --readFilesIn /home/olivier/Bureau/agathe/input/${file} 
    --readFilesCommand gunzip -c --outReadsUnmapped Fastx --outSAMtype BAM Unsorted  
    --outSAMattributes Standard --outSAMunmapped Within --outFilterType BySJout 
    --outFilterMultimapNmax 1 --outFilterMismatchNmax 10 
    --outFilterMismatchNoverLmax 0.3 --quantMode TranscriptomeSAM GeneCounts 
    --twopassMode None 
    --outFileNamePrefix /home/olivier/Bureau/agathe/outputlocal/strict_mm10_Std_${file%%.fq.gz}_
done

#------------------ End of script ----------------------------------


# Mapping output files were copied on the local computer in the following folder: 
/Volumes/Maxtor/projet_DU_AD/results/04_mapping/output_strict_Std_mm10/


# To display mapping report (containing statistical informations): 
cd /Volumes/Maxtor/projet_DU_AD/results/04_mapping/output_strict_Std_mm10/

for file in *Log.final.out
do
    echo $file
    cat $file
done
```

#### Creation of count tables¶

**htseq-count** was used to generate tables containing read counts of each sample, for each gene (see corresponding code after *Mapping visualization section*).
  
  
**Parameter Details:**

- --stranded=no : to specify that RNA-seq data has not been made with a strand-specific protocol.
- --format bam : to specify input file format
- -a 10 : to skip all reads with alignment quality lower than the given minimum value. Default: 10
- --type=exon : to specify feature type (*i.e*. 3rd column of .gtf file) that will be used, all features of other type are ignored. Default: exon
- --idattr=gene\_id : to specify the .gff attribute that will be used as feature ID. Several .gff lines with the same feature ID will be considered as parts of the same feature. The feature ID is used to identity the counts in the output table. Default: gene\_id.
- --mode=union : to specify mode that will be used to handle reads overlapping more than one feature. With union mode, a read is count if there is an overlap with a given region, even if the overlap is partial, as long as there is no ambiguous case (such as overlap with two distinct regions). Default: union.
- --nonunique=none : reads that mapped several genes are skipped.

#### Mapping vizualization¶

To visualize mapping data with **IGV**, mapping output files (*i.e.* .bam files) were sorted and indexed using **samtools sort** and **samtools index**.

In [ ]:

```
%%bash 

# On local computer
conda activate DU_projet
 
cd /mnt/c/Users/Agathe/Desktop/manip_en_cours/projet_DU_remapping

bash script_RNAseq_AD_mm10_htseq_indexIGV.bash

# Script that was runned to generate count tables + .bam indexation (for visualization):

# Code will stop running if any problem is detected
# (first error, undefined variable, pipe error)
set -euo pipefail

# number of samples to analyse
ech="00.5_01 00.5_02 13.5_01 13.5_02 14.5_01 14.5_02 15.5_01 15.5_02 16.5_01 16.5_02"

#-----------------------------------------------------
# Obtention of count tables using htseq-count
#-----------------------------------------------------

mkdir -p /mnt/g/projet_DU_AD/results/05_count_table/output_strict_mm10_Std/

for sample in ${ech}
do
    echo "========================================================================"
    echo "Obtention de la table de comptage de l'échantillon ${sample}-analyse stricte"
    echo "========================================================================"

    htseq-count --stranded=no --format bam -a=10 --type=exon --idattr=gene_id 
    --mode=union --nonunique=none 
    /mnt/g/projet_DU_AD/results/04_mapping/output_strict_Std_mm10/strict_mm10_Std_tg_E${sample}_trimmed_Aligned.out.bam 
    /mnt/g/projet_DU_AD/data/mm10_genome/gencode.vM21.annotation.gtf > 
    /mnt/g/projet_DU_AD/results/05_count_table/output_strict_mm10_Std/c_strict_Std_tg_E${sample}_trimmed_mm10.sorted.bam
done 

#------------------------------------------------
# .bam files indexation for visualization
#------------------------------------------------

cd /mnt/g/projet_DU_AD/results/04_mapping/output_strict_Std_mm10/

echo "=============================================================="
echo "Indexation of sorted bam files for visualisation"
echo "=============================================================="

for sample in ${ech}
do
    echo "=============================================================="
    echo "sample number : ${sample}"
    echo "=============================================================="
    samtools sort strict_mm10_Std_tg_E${sample}_trimmed_Aligned.out.bam > 
    sorted_strict_mm10_Std_tg_E${sample}_trimmed_Aligned.out.bam
    
    samtools index sorted_strict_mm10_Std_tg_E${sample}_trimmed_Aligned.out.bam > sorted_strict_mm10_Std_tg_E${sample}_trimmed_Aligned.out.bai
done
```

#### Identification of differentially expressed genes (DEG) during physiologigal cortex development¶

To identify differentially expressed genes (**DEG**) during brain development, pairwise comparisons of successive developmental stages were performed using **edgeR**, an **R** software and **Bioconductor** package.
  
  
We used (with adaptations) **Sartools** R package, which implementes some **edgeR** functions, and its script template template\_script\_edgeR.r. Main steps of the statistical analysis are described below, followed by the executing code.

##### Filtering very-low counts regions¶

Since regions with very low counts across all samples could interfere with statistical approximations of **edgeR**, these regions were removed using SARTools filter, *i.e.* by selecting regions which contain at least minReplicates (smallest number of replicates = 2) with at least counts per million cutoff of 1 (cpmCutoff = 1).

##### Normalization to avoid regions composition biases¶

To be able to compare read counts between samples, normalization was carried out according to the **edgeR** package methodology. Library of each sample was normalised for regions composition using **calcNormFactors()** function, based on a trimmed mean of M-values (TMM) between each pair of samples.
  
Boxplots of raw- and normalised counts distribution were compared to verify the quality of the normalization process. Normalization is supposed to stabilize distributions across samples.

##### Statistical tests to identify DEG between two successive developmental stages¶

###### Statistical tests¶

Differential analysis was carried out according to the **edgeR** model. edgeR aims at fitting one linear model per region. The edgeR model assumes that the count data follow a negative binomial distribution which is a suitable way to analyse the data when variance is higher than the mean. First step of the statistical procedure is to estimate the dispersion of the data.
After estimation of dispersions - *i.e.* **common** (unique value), **trended** (estimated with splines) and **tagwise** (estimated from feature counts) dispersions - fitting to a **generalized linear model** (glm) can be done using **glmFit()** function. Then, statistical tests was performed for all pairwise comparisons of successive developmental stages, using a **likehood ratio test** (lrt method).
  
Histogram of raw pvalues from the statistical test was plotted to verify the shape of the distribution : it is expected to follow an uniform distribution, with a peak around 0.
Benjamini-Hochberg method was applied to adjust pvalues computed by the statistical test in order to take into account multiple testing and control the false positive rate (FDR). Threshold of statistical significance was set to 0.05.

###### Differential analysis plots¶

###### MA-plot¶

For each comparison, MA-plot of the data was done. It represents the log ratio of differential expression as a function of the mean intensity for each region.

###### Volcano plots¶

Volcano plots were performed for each comparison. It represents the log of the adjusted pvalue as a function of the log ratio of differential expression.

##### Description plots to estimate variability within the experiment¶

###### Scatterplot¶

A pairwise scatterplot was produced to verify that intra-group (replicates) similarities are higher than inter-group (samples from distinct developmental stage) ones. This plot is obtained using **log2(counts+1)**, instead of **raw count values**. This pairwise scatterplot is associated with a **SERE** statistic, that was used as a similarity index between RNA-seq samples.

###### Hierarchical clustering and dendrogram¶

Hierarchical clustering and dendrogram were performed to estimate variability between samples, *i.e.* determine if samples from distinct developmental stages are separated, while replicates are close to each other. Hierarchical clustering was done after a transformation of the count data as **moderated log-counts-per-million**.
  
To plot dendrogram, an euclidean distance is computed between samples. Dendrogram is then obtained from CPM data and built upon the **Ward criterion**.

###### Multidimensional scaling plot (MDS)¶

First two dimensions of a multidimensional scaling plot were plotted to visualise experiment variability. If biological variability is the main source of variance in the data, first dimension is expected to separate samples from the different developmental stages.

##### Adaptations of SARTools functions¶

Following parameters were modified or added from the original Sartools script, to obtain additional graphs or informations:

- addition of a filter to remove uncovered reads. counts\_nf --> counts. In theory, this step is not necessary because uncovered regions should be removed during the filter initially performed by SARTools (it deletes regions with very low-counts). However, it allows to have a double control of this step.
- addition of an unormalised library size histogram graph plot (using ggplot2).
- change of the raw pvalues histogram design (using ggplot2): modifications of summarizeResults.edgeR() function --> summarizeResults.edgeR.AD()
- addition of supplementary graphs in run.edgeR() function
- export of supplementary informations (logFC, FDR)
- addition of a filter using logFC and padj thresholds to obtain a restricted list of DEG, in addition to the list given by SARTools script (which is based on padj threshold only).

In [ ]:

In [ ]:

```
# ----------------------------------------------------------
# 1. Creation of files called targetE13.E14.txt, 
# targetE14.E15.txt, targetE15.E16.txt and targetE16.P0.txt, 
# which contain samples informations for loading count tables:
# -----------------------------------------------------------

#---------------
#For E13 vs E14
#---------------

target_cible = matrix(0,ncol=4, nrow=4)
colnames(target_cible)=c("label","files","group","day")

# Name of samples 
target_cible[,"label"]= c("E13.5_A","E13.5_B","E14.5_A", "E14.5_B")

# Name of files containing count tables
target_cible[,"files"]=c("c_strict_Std_tg_E13.5_01_trimmed_mm10.sorted.txt", 
                         "c_strict_Std_tg_E13.5_02_trimmed_mm10.sorted.txt", 
                         "c_strict_Std_tg_E14.5_01_trimmed_mm10.sorted.txt",
                         "c_strict_Std_tg_E14.5_02_trimmed_mm10.sorted.txt")

# Embryonic days informations
target_cible[,"group"]=c(rep("E13",2), rep("E14",2))

# Day of experiment 
# to correct eventual batch effect.
# Since we don't know when samples where prepared and if they were prepared at the same day, "d1" is written by default, for all samples.
target_cible[,"day"]="d1"

write.table(target_cible, "G:/projet_DU_AD/results/07_DESEq2_mm10/targetE13.14.txt",
            quote=FALSE, col.names=TRUE,row.names=FALSE, sep="\t")


#---------------
#For E14 vs E15
#---------------

target_cible = matrix(0,ncol=4, nrow=4)
colnames(target_cible)=c("label","files","group","day")

# Name of samples 
target_cible[,"label"]= c("E14.5_A","E14.5_B","E15.5_A", "E15.5_B")


# Name of files containing count tables
target_cible[,"files"]=c("c_strict_Std_tg_E14.5_01_trimmed_mm10.sorted.txt",
                         "c_strict_Std_tg_E14.5_02_trimmed_mm10.sorted.txt", 
                         "c_strict_Std_tg_E15.5_01_trimmed_mm10.sorted.txt",
                         "c_strict_Std_tg_E15.5_02_trimmed_mm10.sorted.txt")


# Embryonic days informations
target_cible[,"group"]=c(rep("E14",2), rep("E15",2))

# Day of experiment 
# to correct eventual batch effect.
# Since we don't know when samples where prepared and if they were prepared at the same day, "d1" is written by default, for all samples.
target_cible[,"day"]="d1"

write.table(target_cible, "G:/projet_DU_AD/results/07_DESEq2_mm10/targetE14.15.txt", 
            quote=FALSE, col.names=TRUE,row.names=FALSE, sep="\t")


#---------------
#For E15 vs E16
#---------------

target_cible = matrix(0,ncol=4, nrow=4)
colnames(target_cible)=c("label","files","group","day")

# Name of samples 
target_cible[,"label"]= c("E15.5_A","E15.5_B","E16.5_A", "E16.5_B")

# Name of files containing count tables
target_cible[,"files"]=c("c_strict_Std_tg_E15.5_01_trimmed_mm10.sorted.txt", 
                         "c_strict_Std_tg_E15.5_02_trimmed_mm10.sorted.txt", 
                         "c_strict_Std_tg_E16.5_01_trimmed_mm10.sorted.txt",
                         "c_strict_Std_tg_E16.5_02_trimmed_mm10.sorted.txt")

# Embryonic days informations
target_cible[,"group"]=c(rep("E15",2), rep("E16",2))

# Day of experiment 
# to correct eventual batch effect.
# Since we don't know when samples where prepared and if they were prepared at the same day, "d1" is written by default, for all samples.
target_cible[,"day"]="d1"

write.table(target_cible, "G:/projet_DU_AD/results/07_DESEq2_mm10/targetE15.16.txt", 
            quote=FALSE, col.names=TRUE,row.names=FALSE, sep="\t")


#---------------
#For E16 vs P0
#---------------

target_cible = matrix(0,ncol=4, nrow=4)
colnames(target_cible)=c("label","files","group","day")

# Name of samples 
target_cible[,"label"]= c("E16.5_A","E16.5_B","P0_A", "P0_B")

# Name of files containing count tables
target_cible[,"files"]=c("c_strict_Std_tg_E16.5_01_trimmed_mm10.sorted.txt", 
                         "c_strict_Std_tg_E16.5_02_trimmed_mm10.sorted.txt", 
                         "c_strict_Std_tg_E00.5_01_trimmed_mm10.sorted.txt",
                         "c_strict_Std_tg_E00.5_02_trimmed_mm10.sorted.txt")

# Embryonic days informations
target_cible[,"group"]=c(rep("E16",2), rep("P0",2))

# Day of experiment 
# to correct eventual batch effect.
# Since we don't know when samples where prepared and if they were prepared at the same day, "d1" is written by default, for all samples.
target_cible[,"day"]="d1"

write.table(target_cible, "G:/projet_DU_AD/results/07_DESEq2_mm10/targetE16.P0.txt",
            quote=FALSE, col.names=TRUE,row.names=FALSE, sep="\t")


# ----------------------------------------------------------------------
# 2. Identification of DEG between two developmental stages, 
# in physiological development.
# edgeR analysis using modified SARTools script
# ----------------------------------------------------------------------


# ----------------------
# --------------------------------------------
# E13 vs E14 developmental stages comparison
# --------------------------------------------
# ----------------------

dir.create("G:/projet_DU_AD/results/07_DESEq2_mm10/SARTools/Analyse_L/E13.14", recursive=TRUE)

# -------------------
# Parameters setting
# -------------------

rm(list=ls())                                        

workDir <- "G:/projet_DU_AD/results/07_DESEq2_mm10/SARTools/Analyse_L/E13.14"      
# working directory for the R session

projectName <- "Sartools-edgeR-RNAseq-E13.14"        # name of the project
author <- "Agathe D."                                # author of the statistical analysis/report

targetFile <- "G:/projet_DU_AD/results/07_DESEq2_mm10/targetE13.14.txt"                           
# path to the design/target file
rawDir <- "G:/projet_DU_AD/results/07_DESEq2_mm10/"                                      
# path to the directory containing raw counts files
featuresToRemove <- c("alignment_not_unique",        # names of the features to be removed
                      "ambiguous", "no_feature",     # (specific HTSeq-count information and rRNA for example)
                      "not_aligned", "too_low_aQual")# NULL if no feature to remove

varInt <- "group"                                    # factor of interest
condRef <- "E13"                                     # reference biological condition
batch <- NULL                                        # blocking factor: NULL (default) or "batch" for example

alpha <- 0.05                                        # threshold of statistical significance
pAdjustMethod <- "BH"                                # p-value adjustment method: "BH" (default) or "BY"

cpmCutoff <- 1                                       # counts-per-million cut-off to filter low counts
gene.selection <- "pairwise"                         # selection of the features in MDSPlot
normalizationMethod <- "TMM"                         # normalization method: "TMM" (default), "RLE" (DESeq) or "upperquartile"

colors <- c("darkblue","brown3")               

forceCairoGraph <- FALSE

nsamples = 4                                         # nb of samples
nstage = 2                                           # nb of developemental stages
nrep = 2                                             # nb of replicates per developmental stages
couleur=c("cyan4","brown3","darkgreen","darkblue","plum4","lightgoldenrod3","pink")


# -------------------
# Load packages
# -------------------
setwd(workDir)
library(SARTools)
library(reshape2)
library(ggplot2) 
library(GGally)

#--------------------
# Running script
# checking parameters (unchanged SARTools function)
#--------------------

if (forceCairoGraph) options(bitmapType="cairo")

checkParameters.edgeR(projectName=projectName,author=author,targetFile=targetFile,
                      rawDir=rawDir,featuresToRemove=featuresToRemove,varInt=varInt,
                      condRef=condRef,batch=batch,alpha=alpha,
                      pAdjustMethod=pAdjustMethod,
                      cpmCutoff=cpmCutoff,gene.selection=gene.selection,
                      normalizationMethod=normalizationMethod,colors=colors)

#--------------------
# loading target file (unchanged SARTools function)
#--------------------
target <- loadTargetFile(targetFile=targetFile, varInt=varInt, condRef=condRef, 
                         batch=batch)

#-------------------
# loading count-table
#-------------------
counts_nf <- loadCountData(target=target, rawDir=rawDir, 
                           featuresToRemove=featuresToRemove)

# Delete rows without counts :
counts <- counts_nf[-(which(rowSums(counts_nf) < 1)),]

#------------------
# description plots 
#------------------
# (unchanged SARTools function)
majSequences <- descriptionPlots(counts=counts, group=target[,varInt], col=colors)

# Supplementary graphs
# Print library size (unormalised data) : 
counts_reshape = melt(counts, id.vars = colnames(counts), variable.name='samples_name')
head(counts_reshape)
dim(counts_reshape)
colnames(counts_reshape) = c("peak_name","samples_name","filter_count")
counts_reshape$stages = with(counts_reshape, rep(substr(colnames(counts), 1, 3), 
                                                 each = nrow(counts)))

lib_size_unorm = ggplot(counts_reshape, aes(x=samples_name,y=filter_count, 
                                            fill = stages))

lib_size_unorm + geom_bar(stat="identity") + 
    scale_fill_manual(values=c("skyblue","royalblue", "pink","tomato2","darkgreen")) +
  ggtitle("library size (unormalised dataset)")+ theme_light() + 
  scale_x_discrete(name ="samples_name") + scale_y_continuous("total number of reads") +
  theme(plot.title = element_text(hjust = 0.5))

ggsave(path ="figures/", filename = "01 library size - unormalised dataset.pdf")
ggsave(path ="figures/", filename = "01 library size - unormalised dataset.png")


#---------------
# edgeR analysis (adapted from SARTools function)
#---------------
run.edgeR <- function(counts, target, varInt, condRef, batch=NULL, cpmCutoff=1, 
                      normalizationMethod="TMM", pAdjustMethod="BH", ...){
  # filtering very low-count regions : 
  # select features which contain at least minReplicates 
  # (smallest number of replicates) with at least cpmCutoff counts per million
  minReplicates <- min(table(target[,varInt]))
  fcounts <- counts[rowSums(cpm(counts) >= cpmCutoff) >= minReplicates,]
  cat("Number of features discarded by the filtering:\n")
  cat(nrow(counts)-nrow(fcounts),"\n")
  
  # building dge object
  design <- formula(paste("~", ifelse(!is.null(batch), paste(batch,"+"), ""), varInt))
  dge <- DGEList(counts=fcounts, remove.zeros=TRUE)
  dge$design <- model.matrix(design, data=target)
  cat("\nDesign of the statistical model:\n")
  cat(paste(as.character(design),collapse=" "),"\n")	
  
  ggsave(path ="figures/", filename ="02 - correlation between samples - filter - unnorm.pdf",ggpairs(as.data.frame(dge$counts[,1:nsamples])))
  ggsave(path ="figures/", filename ="02 - correlation between samples - filter - unnorm.png",ggpairs(as.data.frame(dge$counts[,1:nsamples])))
  
  ggsave(path ="figures/", filename ="03 - correlation between log samples - filter - unnorm.pdf",ggpairs(as.data.frame(log(dge$counts[,1:nsamples]))))
  ggsave(path ="figures/", filename ="03 - correlation between log samples - filter - unnorm.png",ggpairs(as.data.frame(log(dge$counts[,1:nsamples]))))
  
  # normalization of regions composition bias
  dge <- calcNormFactors(dge, method=normalizationMethod)
  cat("\nNormalization factors:\n")
  print(dge$samples$norm.factors)
    
  # estimating dispersions
  dge <- estimateGLMCommonDisp(dge, dge$design)
  dge <- estimateGLMTrendedDisp(dge, dge$design)
  dge <- estimateGLMTagwiseDisp(dge, dge$design)
  
  # graphical representation 
  # BCV based on average log CPM (dispersion representation):  
  pdf("figures/04 dispersion plot - glmFit method .pdf")
  plotBCV(dge) 
  dev.off()
  
  png("figures/04 dispersion plot - glmFit method .png")
  plotBCV(dge) 
  dev.off()
  
  # MDS plot: 
  pdf("figures/05 MDS plot - glmFit method - bcv.pdf")  
  mds = plotMDS(dge,  method="bcv",col=rep(couleur[1:nstage],each=nrep), 
                pch=1, cex=1, xlim=c(-0.55,1.0),ylim=c(-0.3,0.4))
  text(mds$x, mds$y, labels=rownames(target), col=rep(couleur[1:nstage],
                                                      each=nrep), pos=3) 
  dev.off()
  
  png("figures/05 MDS plot - glmFit method - bcv.png")  
  mds = plotMDS(dge,  method="bcv",col=rep(couleur[1:nstage],each=nrep), 
                pch=1, cex=1, xlim=c(-0.55,1.0),ylim=c(-0.3,0.4))
  text(mds$x, mds$y, labels=rownames(target), col=rep(couleur[1:nstage],
                                                      each=nrep), pos=3) 
  dev.off()
  
  pdf("figures/06 MDS plot - glmFit method - logFC.pdf")  
  mds = plotMDS(dge,  method="logFC",col=rep(couleur[1:nstage],each=nrep), 
                pch=1, cex=1, xlim=c(-0.55,1.0),ylim=c(-0.3,0.4))
  text(mds$x, mds$y, labels=rownames(target), col=rep(couleur[1:nstage],
                                                      each=nrep), pos=3) 
  dev.off()
  
  png("figures/06 MDS plot - glmFit method - logFC.png")  
  mds = plotMDS(dge,  method="logFC",col=rep(couleur[1:nstage],each=nrep),
                pch=1, cex=1, xlim=c(-0.55,1.0),ylim=c(-0.3,0.4))
  text(mds$x, mds$y, labels=rownames(target), col=rep(couleur[1:nstage],
                                                      each=nrep), pos=3) 
  dev.off()
  
 # statistical testing: 
 # perform all the comparisons between the levels of varInt
  fit <- glmFit(dge, dge$design, ...)
  cat(paste("Coefficients of the model:",paste(colnames(fit$design),collapse="  ")),"\n")
  colsToTest <- grep(varInt,colnames(fit$design))
  namesToTest <- paste0(gsub(varInt,"",colnames(fit$design)[colsToTest]),"_vs_",condRef)
  results <- list()
  # testing coefficients individually (tests against the reference level)
  for (i in 1:length(colsToTest)){
    cat(paste0("Comparison ",gsub("_"," ",namesToTest[i]),": testing coefficient ",
               colnames(fit$design)[colsToTest[i]]),"\n")
      
    lrt <- glmLRT(fit, coef=colsToTest[i])
    results[[namesToTest[i]]] <- topTags(lrt,n=nrow(dge$counts),
                        adjust.method=pAdjustMethod,sort.by="none")$table
  }
  # defining contrasts for the other comparisons (if applicable)
  if (length(colsToTest)>=2){
    colnames <- gsub(varInt,"",colnames(fit$design))
    for (comp in combn(length(colsToTest),2,simplify=FALSE)){ 
      contrast <- numeric(ncol(dge$design))
      contrast[colsToTest[comp[1:2]]] <- c(-1,1)
        
      namecomp <- paste0(colnames[colsToTest[comp[2]]],"_vs_",
                         colnames[colsToTest[comp[1]]])
        
      cat(paste0("Comparison ",gsub("_"," ",namecomp),": testing contrast (",
                 paste(contrast,collapse=", "),")"),"\n")
        
      lrt <- glmLRT(fit, contrast=contrast)
      results[[namecomp]] <- topTags(lrt,n=nrow(dge$counts),
                    adjust.method=pAdjustMethod,sort.by="none")$table
    }
  }
  
  return(list(dge=dge,results=results,lrt=lrt))
}


out.edgeR <- run.edgeR(counts=counts, target=target, varInt=varInt, 
                       condRef=condRef, batch=batch, cpmCutoff=cpmCutoff, 
                       normalizationMethod=normalizationMethod,
                       pAdjustMethod=pAdjustMethod)

#--------------------
# MDS + clustering (unchanged SARTools function)
#--------------------
exploreCounts(object=out.edgeR$dge, group=target[,varInt], 
              gene.selection=gene.selection, col=colors)

#--------------------
# exporting results of the differential analysis (adapted from SARTools function)
#--------------------

rawpHist_AD <- function(complete_AD, outfile=TRUE){
  ncol <- ifelse(length(complete_AD)<=4, ceiling(sqrt(length(complete_AD))), 3)
  nrow <- ceiling(length(complete_AD)/ncol)
  
  par(mfrow=c(nrow,ncol))
  for (name in names(complete_AD)){
    ggplot(as.data.frame(complete_AD[[name]]), aes(x=pvalue, fill="tomato2")) + 
      geom_histogram(binwidth = 0.025, color="black") + 
      scale_fill_discrete(name = "", labels = "pvalues") + 
      labs(title = "Raw pvalues histogram") + 
      theme(plot.margin = margin(2,.8,2,.8, "cm")) 
  }
  if (outfile) ggsave(path= "figures/", filename="rawpHist_ggplot.pdf")
  if (outfile) ggsave(path= "figures/", filename="rawpHist.png")
  if (outfile) dev.off()
}

exportResults.edgeR <- function(out.edgeR, group, counts, alpha=0.05, export=TRUE){
  
  dge <- out.edgeR$dge
  res <- out.edgeR$results
  
  # raw count, normalised count and baseMean
  tmm <- dge$samples$norm.factors
  N <- colSums(dge$counts)
  f <- tmm * N/mean(tmm * N)
  normCounts <- round(scale(dge$counts, center=FALSE, scale=f))
  base <- data.frame(Id=rownames(counts), counts)
  names(base) <- c("Id", colnames(counts))
  norm.bm <- data.frame(Id=rownames(normCounts),normCounts)
  names(norm.bm) <- c("Id", paste0("norm.",colnames(normCounts)))
  norm.bm$baseMean <- round(apply(scale(dge$counts, center=FALSE, scale=f),1,mean),2)
  for (cond in levels(group)){
    norm.bm[,cond] <- round(apply(as.data.frame(normCounts[,group==cond]),1,mean),0)
  }
  base <- merge(base,norm.bm,by="Id",all=TRUE)
  
  complete <- list()
  for (name in names(res)){
    complete.name <- base
    
    # To add info from res
    res.name <- data.frame(Id=rownames(res[[name]]),
                           FC=round(2^(res[[name]][,"logFC"]),3),
                           log2FoldChange=round(res[[name]][,"logFC"],3),
                           pvalue=res[[name]][,"PValue"],
                           padj=res[[name]][,ifelse("FDR" %in% names(res[[name]]), "FDR", "FWER")])
    complete.name <- merge(complete.name, res.name, by="Id", all=TRUE)
    
    # To add info from dge
    dge.add <- data.frame(Id=rownames(dge$counts),
                          tagwise.dispersion=round(dge$tagwise.dispersion,4),
                          trended.dispersion=round(dge$trended.dispersion,4))
    complete.name <- merge(complete.name, dge.add, by="Id", all=TRUE)
    complete[[name]] <- complete.name
    
    if (export){
      # To obtain 'sartools' and restricted lists of DEG during brain development
      up.name <- complete.name[which(complete.name$padj <= alpha & complete.name$log2FoldChange>=0),]
      up.name <- up.name[order(up.name$padj),]
      down.name <- complete.name[which(complete.name$padj <= alpha & complete.name$log2FoldChange<=0),]
      down.name <- down.name[order(down.name$padj),]
      up.name.strict <- complete.name[which(complete.name$padj <= alpha & complete.name$log2FoldChange>=1),]
      up.name.strict <- up.name.strict[order(up.name.strict$padj),]
      down.name.strict <- complete.name[which(complete.name$padj <= alpha & complete.name$log2FoldChange<=-1),]
      down.name.strict <- down.name.strict[order(down.name.strict$padj),]
      for (i in list("up.name", "down.name","up.name.strict","down.name.strict")){
        dimension = nrow(get(i))
        print(paste("number of rows :", i, "=", dimension,sep=" "))
      }
      
      # To save the data
      name <- gsub("_","",name)
      write.table(complete.name, file=paste0("tables/",name,".complete.txt"), 
                  sep="\t", row.names=FALSE, dec=".", quote=FALSE)
        
      write.table(up.name, file=paste0("tables/", name,".up.txt"), row.names=FALSE, 
                  sep="\t", dec=".", quote=FALSE)
        
      write.table(down.name, file=paste0("tables/", name,".down.txt"), row.names=FALSE, 
                  sep="\t", dec=".", quote=FALSE)
        
      write.table(up.name.strict, file=paste0("tables/", name,".up.strict.txt"), 
                  row.names=FALSE, sep="\t", dec=".", quote=FALSE)
        
      write.table(down.name.strict, file=paste0("tables/", name,".down.strict.txt"), 
                  row.names=FALSE, sep="\t", dec=".", quote=FALSE)
    }
  }
  
  return(complete)
}


# Summary of the analysis 
# (boxplots, dispersions, export table, nDiffTotal, histograms, MA plot)
summarizeResults.edgeR.AD = function (out.edgeR, group, counts, alpha = 0.05, 
                        col = c("lightblue","orange", "MediumVioletRed", "SpringGreen"), 
                        log2FClim = NULL, padjlim = NULL) 
{
  if (!I("figures" %in% dir())) 
    dir.create("figures", showWarnings = FALSE)
  if (!I("tables" %in% dir())) 
    dir.create("tables", showWarnings = FALSE)
  countsBoxplots(out.edgeR$dge, group = group, col = col)
  BCVPlot(dge = out.edgeR$dge)
  complete <- exportResults.edgeR(out.edgeR = out.edgeR, group = group, 
                                  counts = counts, alpha = alpha)
  nDiffTotal <- nDiffTotal(complete = complete, alpha = alpha)
  cat("Number of features down/up and total:\n")
  print(nDiffTotal, quote = FALSE)
  rawpHist_AD(complete_AD = complete, outfile = TRUE)
  MAPlot(complete = complete, alpha = alpha, log2FClim = log2FClim)
  volcanoPlot(complete = complete, alpha = alpha, padjlim = padjlim)
  return(list(complete = complete, nDiffTotal = nDiffTotal))
}


summaryResults.AD <- summarizeResults.edgeR.AD(out.edgeR, group=target[,varInt], 
                                        counts=counts, alpha=alpha, col=colors)


#--------------------
# save image of the R session (unchanged SARTools script)
#--------------------
save.image(file=paste0(projectName, ".RData"))


#--------------------
# generating HTML report (adapted from SARTools function)
#--------------------
writeReport.edgeR(target=target, counts=counts, out.edgeR=out.edgeR, 
                  summaryResults=summaryResults.AD, majSequences=majSequences, 
                  workDir=workDir, projectName=projectName, author=author,
                  targetFile=targetFile, rawDir=rawDir, 
                  featuresToRemove=featuresToRemove, varInt=varInt,
                  condRef=condRef, batch=batch, alpha=alpha, 
                  pAdjustMethod=pAdjustMethod, cpmCutoff=cpmCutoff,
                  colors=colors, gene.selection=gene.selection, 
                  normalizationMethod=normalizationMethod)

#--------------------
# save FDR and logCPM information (new in the script)
#--------------------

if (length(names(out.edgeR$results))){
  write.table(out.edgeR$results, file="tables/results_FDR_logCPM_all_filter.txt", sep="\t", row.names=FALSE, dec=".", quote=FALSE)
}else{
  print("Certaines données ne seront pas sauvegardées")
}


# ----------------------
# --------------------------------------------
# Same code for  E14 vs E15 developmental stages comparison 
# with following parameter settings
# --------------------------------------------
# ----------------------


dir.create("G:/projet_DU_AD/results/07_DESEq2_mm10/SARTools/Analyse_L/E14.15", recursive=TRUE)


rm(list=ls())                                       

workDir <- "G:/projet_DU_AD/results/07_DESEq2_mm10/SARTools/Analyse_L/E14.15"      
# working directory for the R session

projectName <- "Sartools-edgeR-RNAseq-E14.15"       # name of the project
author <- "Agathe D."                                # author of the statistical analysis/report

targetFile <- "G:/projet_DU_AD/results/07_DESEq2_mm10/targetE14.15.txt"                           
# path to the design/target file
rawDir <- "G:/projet_DU_AD/results/07_DESEq2_mm10/"                                      
# path to the directory containing raw counts files
featuresToRemove <- c("alignment_not_unique",        # names of the features to be removed
                      "ambiguous", "no_feature",     # (specific HTSeq-count information and rRNA for example)
                      "not_aligned", "too_low_aQual")# NULL if no feature to remove

varInt <- "group"                                    # factor of interest
condRef <- "E14"                                     # reference biological condition
batch <- NULL                                        # blocking factor: NULL (default) or "batch" for example

alpha <- 0.05                                        # threshold of statistical significance
pAdjustMethod <- "BH"                                # p-value adjustment method: "BH" (default) or "BY"

cpmCutoff <- 1                                       # counts-per-million cut-off to filter low counts
gene.selection <- "pairwise"                         # selection of the features in MDSPlot
normalizationMethod <- "TMM"                         # normalization method: "TMM" (default), "RLE" (DESeq) or "upperquartile"

colors <- c("darkblue","brown3")             

forceCairoGraph <- FALSE

nsamples = 4                                         # nb of samples
nstage = 2                                           # nb of developemental stages
nrep = 2                                             # nb of replicates per developmental stages
couleur=c("cyan4","brown3","darkgreen","darkblue","plum4","lightgoldenrod3","pink")


# ----------------------
# --------------------------------------------
# Same code for E15 vs E16 developmental stages comparison
# with following parameter settings
# --------------------------------------------
# ----------------------

dir.create("G:/projet_DU_AD/results/07_DESEq2_mm10/SARTools/Analyse_L/E15.16", recursive=TRUE)

rm(list=ls())                                        

workDir <- "G:/projet_DU_AD/results/07_DESEq2_mm10/SARTools/Analyse_L/E15.16"      # working directory for the R session

projectName <- "Sartools-edgeR-RNAseq-E15.16"        # name of the project
author <- "Agathe D."                                # author of the statistical analysis/report

targetFile <- "G:/projet_DU_AD/results/07_DESEq2_mm10/targetE15.16.txt"                           
# path to the design/target file
rawDir <- "G:/projet_DU_AD/results/07_DESEq2_mm10/"                                      
# path to the directory containing raw counts files
featuresToRemove <- c("alignment_not_unique",        # names of the features to be removed
                      "ambiguous", "no_feature",     # (specific HTSeq-count information and rRNA for example)
                      "not_aligned", "too_low_aQual")# NULL if no feature to remove

varInt <- "group"                                    # factor of interest
condRef <- "E15"                                     # reference biological condition
batch <- NULL                                        # blocking factor: NULL (default) or "batch" for example

alpha <- 0.05                                        # threshold of statistical significance
pAdjustMethod <- "BH"                                # p-value adjustment method: "BH" (default) or "BY"

cpmCutoff <- 1                                       # counts-per-million cut-off to filter low counts
gene.selection <- "pairwise"                         # selection of the features in MDSPlot
normalizationMethod <- "TMM"                         # normalization method: "TMM" (default), "RLE" (DESeq) or "upperquartile"

colors <- c("darkblue","brown3")                    

forceCairoGraph <- FALSE

nsamples = 4                                         # nb of samples
nstage = 2                                           # nb of developemental stages
nrep = 2                                             # nb of replicates per developmental stages
couleur=c("cyan4","brown3","darkgreen","darkblue","plum4","lightgoldenrod3","pink")


# ----------------------
# --------------------------------------------
# Same code for E16 vs P0 developmental stages comparison 
# with following parameter settings
# --------------------------------------------
# ----------------------

dir.create("G:/projet_DU_AD/results/07_DESEq2_mm10/SARTools/Analyse_L/E16.P0", recursive=TRUE)

rm(list=ls())                                        

workDir <- "G:/projet_DU_AD/results/07_DESEq2_mm10/SARTools/Analyse_L/E16.P0"      # working directory for the R session

projectName <- "Sartools-edgeR-RNAseq-E16.P0"                         # name of the project
author <- "Agathe D."                                # author of the statistical analysis/report

targetFile <- "G:/projet_DU_AD/results/07_DESEq2_mm10/targetE16.P0.txt"                           
# path to the design/target file
rawDir <- "G:/projet_DU_AD/results/07_DESEq2_mm10/"                                      
# path to the directory containing raw counts files
featuresToRemove <- c("alignment_not_unique",        # names of the features to be removed
                      "ambiguous", "no_feature",     # (specific HTSeq-count information and rRNA for example)
                      "not_aligned", "too_low_aQual")# NULL if no feature to remove

varInt <- "group"                                    # factor of interest
condRef <- "E16"                                     # reference biological condition
batch <- NULL                                        # blocking factor: NULL (default) or "batch" for example

alpha <- 0.05                                        # threshold of statistical significance
pAdjustMethod <- "BH"                                # p-value adjustment method: "BH" (default) or "BY"

cpmCutoff <- 1                                       # counts-per-million cut-off to filter low counts
gene.selection <- "pairwise"                         # selection of the features in MDSPlot
normalizationMethod <- "TMM"                         # normalization method: "TMM" (default), "RLE" (DESeq) or "upperquartile"

colors <- c("darkblue","brown3")               

forceCairoGraph <- FALSE

nsamples = 4                                         # nb of samples
nstage = 2                                           # nb of developemental stages
nrep = 2                                             # nb of replicates per developmental stages
couleur=c("cyan4","brown3","darkgreen","darkblue","plum4","lightgoldenrod3","pink")
```

#### Annotation of DEG¶

After edgeR running, we obtained lists of DEG with each gene identified by its ENSEMBL ID.

To annotate DEG, a file (biomart\_mm10.txt) containing several annotation data, was generated with **BiomaRt** R package. DEG were annotated using this file and **join unix command**. The file obtained with Biomart contains :

- ensembl\_gene\_id: gene name
- chromosome\_name: chromosome/scaffold number
- strand
- start\_position: gene start (bp)
- end\_position: gene end (bp)
- entrezgene\_id:NCBI gene ID
- gene\_biotype: gene type
- mgi\_symbol: MGI symbol
- entrezgene\_accession: NCBI gene accession
- entrezgene\_description: NCBI gene description
- uniprot\_gn\_symbol: UniProtKB gene name symbol

In [ ]:

```
%%bash

#--------------------------------------
# 1. Obtention of DEG ENSEMBL_ID, without decimal 
# to obtain same ID than those contained in annotation file
#--------------------------------------

cd /mnt/g/projet_DU_AD/results/07_DESEq2_mm10/SARTools/Analyse_L/E13.14/tables
sed '1d' E14vsE13.down.txt | cut -d . -f1 > ENS_ID_down_DEG_E13.14_mm10.txt
sed '1d' E14vsE13.up.txt | cut -d . -f1 > ENS_ID_up_DEG_E13.14_mm10.txt

cd /mnt/g/projet_DU_AD/results/07_DESEq2_mm10/SARTools/Analyse_L/E14.15/tables
sed '1d' E15vsE14.down.txt | cut -d . -f1 > ENS_ID_down_DEG_E14.15_mm10.txt
sed '1d' E15vsE14.up.txt | cut -d . -f1 > ENS_ID_up_DEG_E14.15_mm10.txt

cd /mnt/g/projet_DU_AD/results/07_DESEq2_mm10/SARTools/Analyse_L/E15.16/tables
sed '1d' E16vsE15.down.txt | cut -d . -f1 > ENS_ID_down_DEG_15.E16_mm10.txt
sed '1d' E16vsE15.up.txt | cut -d . -f1 > ENS_ID_up_DEG_E15.16_mm10.txt

cd /mnt/g/projet_DU_AD/results/07_DESEq2_mm10/SARTools/Analyse_L/E14.16/tables
sed '1d' E16vsE14.down.txt | cut -d . -f1 > ENS_ID_down_DEG_14.E16_mm10.txt
sed '1d' E16vsE14.up.txt | cut -d . -f1 > ENS_ID_up_DEG_E14.16_mm10.txt

cd /mnt/g/projet_DU_AD/results/07_DESEq2_mm10/SARTools/Analyse_L/E16.P0/tables
sed '1d' P0vsE16.down.txt | cut -d . -f1 > ENS_ID_down_DEG_16.P0_mm10.txt
sed '1d' P0vsE16.up.txt | cut -d . -f1 > ENS_ID_up_DEG_16.P0_mm10.txt
```

In [ ]:

```
#--------------------------------------
#  2.Obtention of biomart_mm10.txt file
#--------------------------------------

# Loading BiomaRt package :  
library("biomaRt")

# Selection of database and reference genome 
mm10 = useMart("ensembl", dataset="mmusculus_gene_ensembl")
# mm10 Mus musculus version used:  
# Ensembl 97 Jul 2019 http://jul2019.archive.ensembl.org 

# Creating the dataset with information of interest
# It's necessary to do it in two steps, because there is a limitation in the number of attributes that can be collected at the same time.
annot_mm10_part1<-getBM(attributes=c("ensembl_gene_id","chromosome_name","strand", 
                                     "start_position","end_position","entrezgene_id",
                                     "gene_biotype","mgi_symbol",
                                     "entrezgene_accession"), mart=mm10)
annot_mm10_part2<-getBM(attributes=c("ensembl_gene_id","entrezgene_description",
                                     "uniprot_gn_symbol"), mart=mm10)
annot_mm10 = merge(annot_mm10_part1, annot_mm10_part2, by="ensembl_gene_id", all=TRUE)

# Verifications
dim(annot_mm10)
head(annot_mm10)

# Saving
write.table(annot_mm10,
            "E:/projet_DU_AD/results/annotation_mm10/Biomart/biomart_mm10.txt", 
            quote= FALSE, sep="\t",row.names=FALSE)
```

In [ ]:

```
%%bash

#--------------------------------------
# 3.Combine DEG files with biomart annotation file 
#--------------------------------------

# Copy and paste files containing ENSEMBL ID (of up or down DEG)
# in folder /mnt/g/projet_DU_AD/results/09_annotation/Sartools/Biomart/raw_data

# Datasets formatting:
# To put ENSEMBL ID in the first column: 
cd /mnt/g/projet_DU_AD/results/annotation_mm10/Biomart

sed '1d' biomart_mm10.txt | sort > biomart_mm10_sorted.txt

mkdir -p /mnt/g/projet_DU_AD/results/09_annotation/Sartools/Biomart

cp /mnt/g/projet_DU_AD/results/annotation_mm10/Biomart/biomart_mm10_sorted.txt 
    /mnt/g/projet_DU_AD/results/09_annotation/Sartools/Biomart

cd /mnt/g/projet_DU_AD/results/09_annotation/Sartools/Biomart/

sample="E13.14 E14.15 E15.16 E14.16 E16.P0"

for ech in ${sample}
do
    echo "=============================================="
    echo "file name : ${ech}"
    echo "Annotation"
    echo "=============================================="

    sort -k1,1 raw_data/ENS_ID_up_DEG_${ech}_mm10.txt 
    > raw_data/ENS_ID_up_DEG_${ech}_mm10_sorted.txt
    
    sort -k1,1 raw_data/ENS_ID_down_DEG_${ech}_mm10.txt 
    > raw_data/ENS_ID_down_DEG_${ech}_mm10_sorted.txt

# To combine informations
    join -1 1 -2 1 raw_data/ENS_ID_up_DEG_${ech}_mm10_sorted.txt 
    biomart_mm10_sorted.txt -t $'\t' > AnnotBM_up_${ech}_mm10.txt 
    
    join -1 1 -2 1 raw_data/ENS_ID_down_DEG_${ech}_mm10_sorted.txt 
    biomart_mm10_sorted.txt -t $'\t' > AnnotBM_down_${ech}_mm10.txt 

    echo "AnnotBM_up_${ech}_mm10.txt"
    wc -l AnnotBM_up_${ech}_mm10.txt
    head AnnotBM_up_${ech}_mm10.txt

    echo "AnnotBM_down_${ech}_mm10.txt"
    wc -l AnnotBM_down_${ech}_mm10.txt
    head AnnotBM_down_${ech}_mm10.txt 

done
```

#### Combination of annotations and statistical informations¶

In order to obtain a file where each region of interest is represented by a single row, the annotation elements of a given region that are on distinct lines have been grouped together. Then, all data (statistical informations from edgeR, mm10 coordinates regions, annotation informations) are combined in an single file (one file per pairwise comparison).

In [ ]:

```
%%bash 

# ----------------------------------------------------------------------
# Dataset n° 1 (files containing statistical informations) formatting
# ----------------------------------------------------------------------

# To remove header
cd  /mnt/g/projet_DU_AD/results/07_DESEq2_mm10/SARTools/Analyse_L/E13.14/tables
sed '1d' E14vsE13.down.txt > nh_down_DEG_E13.14_mm10.txt
sed '1d' E14vsE13.up.txt > nh_up_DEG_E13.14_mm10.txt

cd  /mnt/g/projet_DU_AD/results/07_DESEq2_mm10/SARTools/Analyse_L/E14.15/tables
sed '1d' E15vsE14.down.txt > nh_down_DEG_E14.15_mm10.txt
sed '1d' E15vsE14.up.txt > nh_up_DEG_E14.15_mm10.txt

cd  /mnt/g/projet_DU_AD/results/07_DESEq2_mm10/SARTools/Analyse_L/E15.16/tables
sed '1d' E16vsE15.down.txt > nh_down_DEG_E15.16_mm10.txt
sed '1d' E16vsE15.up.txt > nh_up_DEG_E15.16_mm10.txt

cd  /mnt/g/projet_DU_AD/results/07_DESEq2_mm10/SARTools/Analyse_L/E14.16/tables
sed '1d' E16vsE14.down.txt > nh_down_DEG_E14.16_mm10.txt
sed '1d' E16vsE14.up.txt > nh_up_DEG_E14.16_mm10.txt

cd  /mnt/g/projet_DU_AD/results/07_DESEq2_mm10/SARTools/Analyse_L/E16.P0/tables
sed '1d' P0vsE16.down.txt > nh_down_DEG_E16.P0_mm10.txt
sed '1d' P0vsE16.up.txt > nh_up_DEG_E16.P0_mm10.txt

# To combine files, we need a common ID = ENSEMBL_ID (without decimal)
# To replace ENSEMBL_ID (with decimal) to ENSEMBL_ID (without) in identified DEG files, containing statistical informations
sample="E13.14 E14.15 E15.16 E14.16 E16.P0"

for ech in ${sample}
do
    cd /mnt/g/projet_DU_AD/results/07_DESEq2_mm10/SARTools/Analyse_L/${ech}/tables
    
    echo "------------------------------------"
    echo "${ech}"
    echo "------------------------------------"

    paste ENS_ID_down_DEG_${ech}_mm10.txt nh_down_DEG_${ech}_mm10.txt > 
    ID_double_nh_down_DEG_${ech}_mm10.txt
    
    paste ENS_ID_up_DEG_${ech}_mm10.txt nh_up_DEG_${ech}_mm10.txt > 
    ID_double_nh_up_DEG_${ech}_mm10.txt
    
    awk '{ print $1"\t"$3"\t"$4"\t"$5"\t"$6"\t"$7"\t"$8"\t"$9"\t"$10"\t"$11"\t"$12"\t"$13"\t"$14"\t"$15"\t"$16"\t"$17"\t"$18"\t"$19}' ID_double_nh_down_DEG_${ech}_mm10.txt > ID_sd_nh_down_${ech}_mm10.txt
    awk '{ print $1"\t"$3"\t"$4"\t"$5"\t"$6"\t"$7"\t"$8"\t"$9"\t"$10"\t"$11"\t"$12"\t"$13"\t"$14"\t"$15"\t"$16"\t"$17"\t"$18"\t"$19 }' ID_double_nh_up_DEG_${ech}_mm10.txt > ID_sd_nh_up_${ech}_mm10.txt

    echo "ID_sd_nh_down${ech}_mm10.txt"
    wc -l ID_sd_nh_down_${ech}_mm10.txt
    head ID_sd_nh_down_${ech}_mm10.txt

    echo "ID_sd_nh_up${ech}_mm10.txt"
    wc -l ID_sd_nh_up_${ech}_mm10.txt
    head ID_sd_nh_up_${ech}_mm10.txt
done
```

In [ ]:

```
# ----------------------------------------------------------------------
# To concatenate informations
# ----------------------------------------------------------------------

#------------------------------------------------------------------------
# 1. To Concatenate annotation informations : one row = one region of interest / DEG
# Load files : List files containing "AnnotBM" or "ID_sd" patterns
list_files = list.files("G:/projet_DU_AD/results/09_annotation/Sartools/Biomart")
list_files_Annot = list_files[which(grepl("AnnotBM",list_files))]
list_files_ID_sd = list_files[which(grepl("ID_sd",list_files))]

for(i in 1:10){
  Annot = list_files_Annot[i]
  DEG_annot = read.table(file = paste("G:/projet_DU_AD/results/09_annotation/Sartools/Biomart",Annot,sep="/"), header = FALSE, sep = "\t", quote="", fill=TRUE)
  
  colnames(DEG_annot) = c("ID", "chr", "strand","start_gene","end_gene","entrezgene_id",
                          "gene_biotype","mgi_symbol","entrezgene_accession",
                          "entrezgene_description", "uniprot_gn_symbol")
  
  # To concatenate informations 
  library(dplyr)
  
  # List of unique ID  
  liste_DEG_unique <- unique(DEG_annot$ID)
  
  # empty matrice which will be filled during the loop
  # Number of rows = nb of unique ID | Nb of columns = nb of variables
  treatment_file <- data.frame(matrix(NA,nrow=length(liste_DEG_unique),ncol=ncol(DEG_annot)))
  colnames(treatment_file) <- colnames(DEG_annot)
  
  treatment_file$ID <- liste_DEG_unique
  for(k in 1:length(liste_DEG_unique)){
    # Filter according to ID k 
    DEG_filt <- DEG_annot %>%
      filter(ID == liste_DEG_unique[k])
    
    # Collapsing of distincts elements of a variables for ID k 
    # If variable is composed of only one element ==> we keep this element only once 
    # If variable is composed of m distinct elements ==> even if elements is repeated, 
      # we keep all the elements (with repetitions)
    for(j in 2:length(colnames(treatment_file))){
      if(length(unique(DEG_filt[,j])) == 1){
        treatment_file[k,j] <- paste(unique(DEG_filt[,j]), collapse=';')
      } else {
        treatment_file[k,j] <- paste(DEG_filt[,j], collapse=';')  
      }
    }
  }
  print(paste("nb of uniq ID treatment file : ", nrow(treatment_file), sep=""))
  
  #------------------------------------------------------------------------
  # 2. To merge DEG annotations with statistical informations obtained with SARTools-edgeR for every regions
  
  ID_sd = list_files_ID_sd[i]
  sartools_file = read.table(file = paste("G:/projet_DU_AD/results/09_annotation/Sartools/Biomart",ID_sd,sep="/"), header = FALSE, sep = "\t", quote="", dec=".")
  
  colnames(sartools_file) = c("ID","cond1_rep1","cond1_rep2","cond2_rep1", 
                              "cond2_rep2","norm.cond1_rep1","norm.cond1_rep2",
                              "norm.cond2_rep1","norm.cond2_rep2","baseMean",
                              "cond1","cond2","FC","log2FoldChange",
                              "pvalue","padj","tagwise.dispersion","trended.dispersion")
    
  Annot_DESeq = merge(sartools_file,treatment_file, by="ID", all = TRUE)
    
  write.table(Annot_DESeq, file=paste("G:/projet_DU_AD/results/09_annotation/Sartools/Biomart/edgeR",Annot,sep="_"), quote=FALSE, sep="\t", row.names = FALSE, col.names = TRUE)
  
  size_Annot = nrow(treatment_file)
  size_ID_sd = nrow(sartools_file)
  
  if(size_Annot != size_ID_sd){
    print(paste("The two files", Annot, "and", ID_sd, 
                "don't have the same size !", sep =" "))
  }else{
  }
}
```

Because Gencode and Biomart annotation files not always contain exactly the same informations, due to differences between available informations in each database, few DEG were not associated to a gene name (in 5 of 8 files), using Biomart annotation file.

**Number of DEG without annotation:**

- 3 downregulated genes between E13 and E14
- 1 downregulated gene between E14 and E15
- 0 downregulated gene between E15 and E16
- 2 downregulated genes between E16 and P0
- 1 upregulated gene between E13 and E14
- 2 upregulated genes between E14 and E15
- 0 upregulated gene between E15 and E16
- 0 upregulated gene between E16 and P0

The DEG lacking annotation information were manually annotated, based on Gencode annotation contained in the file called gencode.vM21.annotation.gtf.

In [ ]:

```
# Automatisation to annotate DEG that lack annotation information, using Gencode annotation database.

library(data.table)
library(stringr)

# Name of samples
nom_sample = list("down_E13.14", "down_E14.15", "down_E14.16", "down_E15.16", 
                  "down_E16.P0", "up_E13.14", "up_E14.15", "up_E14.16", 
                  "up_E15.16", "up_E16.P0" )
list_sample <- list()

# Load samples files : 
for(i in 1:length(nom_sample)){
  list_sample[[i]] <- read.table(paste("G:/projet_DU_AD/results/09_annotation/Sartools/Biomart/edgeR_AnnotBM_", 
            nom_sample[i] , "_mm10.txt", sep=""), sep = "\t", header= TRUE, 
            dec=".", quote="", stringsAsFactors = FALSE)
    
  print(paste("head_ech",nom_sample[i], sep=""))
  print(head(list_sample[[i]]))
  print(paste("dimension du dataset ech", nom_sample[i], sep = "_"))
  print(dim(list_sample[[i]]))
}
names(list_sample) <- nom_sample
```

In [ ]:

```
%%bash 

# Gencode file formatting
# To remove 5 first lines of the file (corresponding to the header)

cd /mnt/g/projet_DU_AD/data/mm10_genome
sed -e '1,5d' gencode.vM21.annotation.gtf > nh_gencode.vM21.annotation.gtf
```

In [ ]:

```
gencode <- read.table("G:/projet_DU_AD/data/mm10_genome/nh_gencode.vM21.annotation.gtf", sep = "\t", header= FALSE)

colnames(gencode) <- c("chr","source","element_type","start","end","V6","strand","V8","infos")

head(gencode)
dim(gencode)

for(i in 1:length(nom_sample)){
  # To find rows where annotation informations are missing
  na_values = list_sample[[i]][is.na(list_sample[[i]][,"strand"])==TRUE,]
  
  if(nrow(na_values)>0){
    print(nom_sample[[i]])
    print(na_values)
    # to replace missing annotation informations by data from Gencode database
    na_values_id <- as.character(na_values$ID)
    
    for(j in 1:length(na_values_id)){
      info_to_get <- gencode[(gencode$infos %like% na_values_id[j]) & gencode$element_type =="gene",]
      list_sample[[i]][rownames(na_values)[j],"chr"] <- info_to_get$chr
      list_sample[[i]][rownames(na_values)[j],"strand"] <- info_to_get$strand
      list_sample[[i]][rownames(na_values)[j],"start_gene"] <- info_to_get$start
      list_sample[[i]][rownames(na_values)[j],"end_gene"] <- info_to_get$end
      
      list_sample[[i]][rownames(na_values)[j],"entrezgene_id"] <- "." 
      
      list_sample[[i]][rownames(na_values)[j],"gene_biotype"] <- 
        str_split(str_split(info_to_get$infos,pattern = "; ")[[1]][2], 
                  pattern="gene_type ")[[1]][2]
        
      list_sample[[i]][rownames(na_values)[j],"mgi_symbol"] <- 
        str_split(str_split(info_to_get$infos,pattern = "; ")[[1]][3], 
                  pattern="gene_name ")[[1]][2]
        
      list_sample[[i]][rownames(na_values)[j],"entrezgene_accession"] <- 
        str_split(str_split(info_to_get$infos,pattern = "; ")[[1]][3], 
                  pattern="gene_name ")[[1]][2]
        
      list_sample[[i]][rownames(na_values)[j],"entrezgene_description"] <- "."
      list_sample[[i]][rownames(na_values)[j],"uniprot_gn_symbol"] <- "." 
      
    }
    print(list_sample[[i]][rownames(na_values),])
    
  } else {
     write.table(list_sample[[i]],
                file=paste("G:/projet_DU_AD/results/09_annotation/Sartools/Biomart/edgeR_AnnotBM_et_Man_",nom_sample[i],"_mm10.txt",sep=""),
               quote=FALSE,sep="\t",row.names=FALSE,col.names=TRUE)
  }
  
  # Save data
   write.table(list_sample[[i]],
              file=paste("G:/projet_DU_AD/results/09_annotation/Sartools/Biomart/edgeR_AnnotBM_et_Man_",nom_sample[i],"_mm10.txt",sep=""),
              quote=FALSE,sep="\t",row.names=FALSE,col.names=TRUE)
  
}
```

#### Generate .bed files for data visualization¶

To visualize DEG coordinates using **IGV**, .bed files (containing chromosome, start and end informations of each DEG) were generated from files containing DEG informations.

In [ ]:

```
nom_sample = list=c("down_E13.14", "down_E14.15", "down_E14.16", "down_E15.16", 
                    "down_E16.P0", "up_E13.14", "up_E14.15", "up_E14.16", 
                    "up_E15.16", "up_E16.P0" )

# Load files: 
for(name in nom_sample){
  assign(x=paste("ech",name, sep = "_"), 
         value = read.table(paste("G:/projet_DU_AD/results/09_annotation/Sartools/Biomart/edgeR_AnnotBM_et_Man_", 
         name , "_mm10.txt", sep=""), sep = "\t", header= TRUE, dec=".", quote=""))
  print(paste("head_ech",name, sep=""))
  print(head(eval(parse(text = paste("ech",name, sep="_")))))
  print(paste("dimension du dataset ech",name, sep = "_"))
  print(dim(eval(parse(text = paste("ech",name, sep = "_")))))
}


setwd("G:/projet_DU_AD/results/09_annotation/Sartools/Biomart/")

mes_fichiers = list("ech_up_E13.14", "ech_up_E14.15", "ech_up_E14.16", 
                    "ech_up_E15.16", "ech_up_E16.P0", "ech_down_E13.14", 
                    "ech_down_E14.15", "ech_down_E14.16", "ech_down_E15.16", 
                    "ech_down_E16.P0")

for (i in mes_fichiers){
  dataset = get(i)
  dataset$chromosome = with(dataset, paste("chr",dataset[,"chr"], sep=""))
  bed_file = dataset[,c("chromosome", "start_gene", "end_gene")]
  print(paste("dimension du fichier bed de ",i, ":", nrow(bed_file), sep =" "))
  print(head(bed_file))
  write.table(bed_file, file=paste("G:/projet_DU_AD/results/09_annotation/Sartools/Biomart/bed_edge_AnnotBM_et_Man_",i,"_mm10.bed", sep=""), quote=FALSE, row.names=FALSE, col.names=FALSE, sep="\t")
}
```

In [ ]:

```
%%bash

# To sort .bed files
cd /mnt/g/projet_DU_AD/results/09_annotation/Sartools/Biomart

for file in bed*
do  sort -k1,1 -k2,2n $file > ${file/.bed/.sorted.bed}
echo $file
done
```

#### Gene ontology¶

To perform a Gene ontology (GO) analysis, Gene names of DEG were isolated from files containing DEG informations.
GO analysis was done using **WEB-based GEne SeT AnaLysis Toolkit** (GESTALT) for DEG that were found between E14.5 and E15.5 and between E15.5 and E16.5.

In [ ]:

```
setwd("E:/projet_DU_AD/results/09_annotation/Sartools/Biomart")

DEG_u14_15 = read.table("edgeR_AnnotBM_et_Man_up_E14.15_mm10.txt", 
                        header = TRUE, fill=TRUE, quote="", sep="\t", 
                        na.strings = "", stringsAsFactors = FALSE)

DEG_u15_16 = read.table("edgeR_AnnotBM_et_Man_up_E15.16_mm10.txt", 
                        header = TRUE, fill=TRUE, quote="", sep="\t", 
                        na.strings = "", stringsAsFactors = FALSE)

DEG_d14_15 = read.table("edgeR_AnnotBM_et_Man_down_E14.15_mm10.txt",
                        header = TRUE, fill=TRUE, quote="", sep="\t", 
                        na.strings = "", stringsAsFactors = FALSE)

DEG_d15_16 = read.table("edgeR_AnnotBM_et_Man_down_E15.16_mm10.txt",
                        header = TRUE, fill=TRUE, quote="", sep="\t", 
                        na.strings = "", stringsAsFactors = FALSE)

head(DEG_u14_15)
head(DEG_u15_16)
head(DEG_d14_15)
head(DEG_d15_16)

dim(DEG_u14_15)
dim(DEG_u15_16)
dim(DEG_d14_15)
dim(DEG_d15_16)

# Function to extract ID
unlist_gene = function(treatment_file, gene_ID, file_name, folder){
  # To extract ID from mm10 Biomart annotation files: 
  # Obtention of a matrix with only one column, containing ID
  gene_name_ID = as.character(treatment_file[,gene_ID])
  
  # Obtention of a list with ID which are deconcatenate
  list_gene_splitted <- strsplit(gene_name_ID,split=";")
  vec_gene_splitted <- unique(unlist(list_gene_splitted))
  
  # Remove missing data = "." : 
  vec_gene_splitted_filter = vec_gene_splitted[vec_gene_splitted!="."]
  
  # Obtention of data in a table:
  table_VGS_filter = as.data.frame(vec_gene_splitted_filter)
  
  write.table(table_VGS_filter, paste(folder,gene_ID,"_",file_name,".txt",sep=""), 
              quote=FALSE, col.names=FALSE, row.names=FALSE)
}


# Extraction of ENSEMBL ID : ID fileswere renamed to precise that ID correspond to ENSEMBL ID.
unlist_gene(treatment_file=DEG_u14_15, gene_ID="ID", file_name="DEG_up_14_15",
            folder="E:/projet_DU_AD/results/09_annotation/Sartools/Biomart/ID/")

unlist_gene(treatment_file=DEG_u15_16, gene_ID="ID", file_name="DEG_up_15_16",
            folder="E:/projet_DU_AD/results/09_annotation/Sartools/Biomart/ID/")

unlist_gene(treatment_file=DEG_d14_15, gene_ID="ID", file_name="DEG_down_14_15",
            folder="E:/projet_DU_AD/results/09_annotation/Sartools/Biomart/ID/")

unlist_gene(treatment_file=DEG_u15_16, gene_ID="ID", file_name="DEG_down_15_16",
            folder="E:/projet_DU_AD/results/09_annotation/Sartools/Biomart/ID/")

# Extraction of Entrez accession ID
unlist_gene(treatment_file=DEG_u14_15, gene_ID="entrezgene_accession", 
            file_name="DEG_up_14_15",
            folder="E:/projet_DU_AD/results/09_annotation/Sartools/Biomart/ID/")

unlist_gene(treatment_file=DEG_u15_16, gene_ID="entrezgene_accession", 
            file_name="DEG_up_15_16",
            folder="E:/projet_DU_AD/results/09_annotation/Sartools/Biomart/ID/")

unlist_gene(treatment_file=DEG_d14_15, gene_ID="entrezgene_accession", 
            file_name="DEG_down_14_15",
            folder="E:/projet_DU_AD/results/09_annotation/Sartools/Biomart/ID/")

unlist_gene(treatment_file=DEG_u15_16, gene_ID="entrezgene_accession", 
            file_name="DEG_down_15_16",
            folder="E:/projet_DU_AD/results/09_annotation/Sartools/Biomart/ID/")

#Extraction of mgi ID
unlist_gene(treatment_file=DEG_u14_15, gene_ID="mgi_symbol", 
            file_name="DEG_up_14_15",
            folder="E:/projet_DU_AD/results/09_annotation/Sartools/Biomart/ID/")

unlist_gene(treatment_file=DEG_u15_16, gene_ID="mgi_symbol", 
            file_name="DEG_up_15_16",
            folder="E:/projet_DU_AD/results/09_annotation/Sartools/Biomart/ID/")

unlist_gene(treatment_file=DEG_d14_15, gene_ID="mgi_symbol", 
            file_name="DEG_down_14_15",
            folder="E:/projet_DU_AD/results/09_annotation/Sartools/Biomart/ID/")

unlist_gene(treatment_file=DEG_u15_16, gene_ID="mgi_symbol", 
            file_name="DEG_down_15_16",
            folder="E:/projet_DU_AD/results/09_annotation/Sartools/Biomart/ID/")


#Extraction of Entrez_gene_ID
unlist_gene(treatment_file=DEG_u14_15, gene_ID="entrezgene_id", 
            file_name="DEG_up_14_15",
            folder="E:/projet_DU_AD/results/09_annotation/Sartools/Biomart/ID/")

unlist_gene(treatment_file=DEG_u15_16, gene_ID="entrezgene_id", 
            file_name="DEG_up_15_16",
            folder="E:/projet_DU_AD/results/09_annotation/Sartools/Biomart/ID/")

unlist_gene(treatment_file=DEG_d14_15, gene_ID="entrezgene_id", 
            file_name="DEG_down_14_15",
            folder="E:/projet_DU_AD/results/09_annotation/Sartools/Biomart/ID/")

unlist_gene(treatment_file=DEG_u15_16, gene_ID="entrezgene_id", 
            file_name="DEG_down_15_16",
            folder="E:/projet_DU_AD/results/09_annotation/Sartools/Biomart/ID/")
```
