## Supplementary material for "Immediate perturbations of DNA methylation and transcriptome upon acute prenatal alcohol exposure in the mouse developing brain cortex": Supp data Notebook 4 Integration data: Supp. data Notebook 4 - integration dataset _ Duchateau et al..html

Supp. data 4 integration\_DMR\_and\_other\_data


### Supplementary data Notebook 4 - Bioinformatic workflow: combination of data from distinct analyses¶

##### *Agathe Duchateau*¶

#### Integration of DMR with DOCR¶

To determine whether DMRs are located within genes that are differential opened or closed (DOCR) during physiological brain development, DMRs and DOCR regions were compared using our own R function called ***find\_overlaps\_AD\_table()***. Both overlaping regions and regions less than 1000 bases apart were investigated, by setting ***tolerance*** argument to 0 or 1000. The obtained regions were then annotated, using annotations that were already achieved in the two complete datasets.

In [ ]:

```
%%R

####################################################################
# Integration of DMR and DOCR to find potential overlap
# (mm10 chromosomal coordinates)
####################################################################

# 1.To load DMR432 data - mm10 
DMR432 = read.table("H:/methylome/fusion/post_methylkit/DMR432_pval0.07/annotations/Biomart/mm10/DMR432_unique_annot_mm10_BMT_meth_cgi.bed", sep="\t", header = FALSE)
head(DMR432)

# To extract DMR432 chromosomic coordinates
DMR432_bed = DMR432[,c("V1","V2","V3")]
head(DMR432_bed)
dim(DMR432_bed)


# 2. To load DOCR data - mm10 (from edgeR analysis):
# ATAC-seq SORTED files - obtained with SARTools edgeR :
nom_sample = list=c("down_DOCR_E13.14", "down_DOCR_E14.15", "down_DOCR_E14.16", 
                    "down_DOCR_E15.16", "down_DOCR_E16.P0", "up_DOCR_E13.14", 
                    "up_DOCR_E14.15", "up_DOCR_E14.16", "up_DOCR_E15.16", 
                    "up_DOCR_E16.P0" )

# /!\ /!\ Dataset must be in a matrix format, otherwise, function doesn't work /!\/!\
for(name in nom_sample){
  assign(x=name, value = as.matrix(read.table(paste("G:/projet_DU_DSD/results/10_annotations/Sartools/Biomart/mm10/bed_gff_ID_", name, "_mm10.sorted.bed",  
              sep=""), sep = "\t", header= FALSE, dec=".", quote="")))
  print(paste("head_",name, sep=""))
  print(head(eval(parse(text = name))))
  print(paste("dimension du dataset ",name, sep = ""))
  print(dim(eval(parse(text = name))))
}

# Creation of a list containing each DOCR file (up or down for each developmental stage) 
ATAC = list(down_DOCR_E13.14 = down_DOCR_E13.14, down_DOCR_E14.15 = down_DOCR_E14.15, 
            down_DOCR_E15.16 = down_DOCR_E15.16, down_DOCR_E14.16 = down_DOCR_E14.16, 
            down_DOCR_E16.P0 = down_DOCR_E16.P0,up_DOCR_E13.14 = up_DOCR_E13.14, 
            up_DOCR_E14.15 = up_DOCR_E14.15 ,up_DOCR_E15.16 = up_DOCR_E15.16, 
            up_DOCR_E14.16 = up_DOCR_E14.16, up_DOCR_E16.P0 = up_DOCR_E16.P0)


for (i in 1:length(ATAC)){
  print(names(ATAC)[i])
  print(head(ATAC[[i]]))
  print(dim(ATAC[[i]]))
} 


# 3. Function to find overlap or close regions (depending on value given for tolerance argument)
find_overlaps_AD_table <- function (regions_1,regions_2,tolerance=0)
{
  common_region_table = NULL
  number_of_regions_1 <- nrow(regions_1)
  number_of_regions_2 <- nrow(regions_2)
  
  overlapped_is <- c()
  overlapped_js <- c()
  
  for (i in 1:number_of_regions_1) {
    
    for (j in 1:number_of_regions_2) {
      
      if (as.character(regions_1[i,1])==as.character(regions_2[j,1])) {
        start_1 <- as.numeric(regions_1[i,2])
        end_1 <- as.numeric(regions_1[i,3])
        expanded_start_1 <- start_1 - tolerance
        expanded_end_1 <- end_1 + tolerance
        start_2 <- as.numeric(regions_2[j,2])
        end_2 <- as.numeric(regions_2[j,3])
        
        if (expanded_end_1>=start_2) {
          
          if (end_2>=expanded_start_1) {
            
            overlapped_is <- c(overlapped_is,i)
            overlapped_js <- c(overlapped_js,j)
            
            first_region <- paste(regions_1[i,1],start_1,end_1,sep="_")
            second_region <- paste(regions_2[j,1],start_2,end_2,sep="_")
            overlap_size <- min(end_1,end_2)-max(start_1,start_2)+1
            common_region_table = rbind(common_region_table,c(as.character(regions_1[i,1]),start_1,end_1, 
                                                              as.character(regions_2[j,1]),start_2,end_2, overlap_size))
            
            if (overlap_size>0) {
              print(paste(first_region,second_region,sep=" "))
            } else {
              print(paste(first_region,second_region, sep=" "))
            }
          }
        } 
      }
    }
  }
  print(paste("Number matched in first data set:",length(unique(overlapped_is))))
  print(paste("Number matched in second data set:",length(unique(overlapped_js))))
  if (length(common_region_table) > 0){
    colnames(common_region_table) = c("chr_reg1", "start_reg1", "end_reg1", "chr_reg2", "start_reg2", "end_reg2","overlap_size")
  }else{
  }
  return(as.data.frame(common_region_table))
}


# 4. To search for common regions between DMR and DOCR 
# overlap
for (i in 1:length(ATAC)) {
  print(names(ATAC)[i])
  DMR_ATAC = find_overlaps_AD_table(DMR432_bed, ATAC[[i]], tolerance=0)
  write.table(DMR_ATAC,paste("H:/methylome/fusion/post_methylkit/DMR432_pval0.07/croisement_autre_data/mm10_DMR432", names(ATAC)[i], "dist0.txt",sep="_"), sep = "\t", quote=FALSE, row.names = FALSE, col.names=FALSE)
}


# 5. To search for common regions between DMR and DOCR 
# maximum distance between 2 regions = 1000bases: 
for (i in 1:length(ATAC)) {
  print(names(ATAC)[i])
  DMR_ATAC = find_overlaps_AD_table(DMR432_bed, ATAC[[i]], tolerance=1000)
  write.table(DMR_ATAC,paste("H:/methylome/fusion/post_methylkit/DMR432_pval0.07/croisement_autre_data/mm10_DMR432", names(ATAC)[i], "dist1000.txt",sep="_"), sep = "\t", quote=FALSE, row.names = FALSE, col.names=FALSE)
}


###################
# Annotation
###################

# To load files or list of files
# To load DMR+DOCR regions
list_DMR_DOCR = list.files("H:/methylome/fusion/post_methylkit/DMR432_pval0.07/croisement_autre_data/DMR_ATACseq/")
list_DMR_DOCR_dist0 = list_DMR_DOCR[which(grepl("dist0",list_DMR_DOCR))]
list_DMR_DOCR_dist1000 = list_DMR_DOCR[which(grepl("dist1000",list_DMR_DOCR))]

# List of DOCR informations 
# Since some files were empty (no overlap), we need to specify the list of files we want to annotate.
list_DOCR_Annot=list("stats_mm9_mm10_down_DOCR_E14.15_annot_syntaxique_mm10_biomart.bed",
        "stats_mm9_mm10_down_DOCR_E14.16_annot_syntaxique_mm10_biomart.bed",
        "stats_mm9_mm10_down_DOCR_E16.P0_annot_syntaxique_mm10_biomart.bed", 
        "stats_mm9_mm10_up_DOCR_E14.15_annot_syntaxique_mm10_biomart.bed",
        "stats_mm9_mm10_up_DOCR_E14.16_annot_syntaxique_mm10_biomart.bed", 
        "stats_mm9_mm10_up_DOCR_E15.16_annot_syntaxique_mm10_biomart.bed",
        "stats_mm9_mm10_up_DOCR_E16.P0_annot_syntaxique_mm10_biomart.bed")

# To load DMR432 annotation
DMR_Annot = read.table("H:/methylome/fusion/post_methylkit/DMR432_pval0.07/annotations/Biomart/mm10/DMR432_unique_annot_mm10_BMT_meth_cgi.bed", header=FALSE, sep="\t", quote="")
head(DMR_Annot)
dim(DMR_Annot)
colnames(DMR_Annot) = c("mm10_chr","DMR_start","DMR_end","DMR_ID","chr_annot",
                        "start_annot","end_annot","strand","entrezgene_accession",
                        "gene_biotype","mgi_symbol","ENSEMBL_ID","Entrez_ID",
                        "overlap_annot_length","chr_mm9","start_mm9","end_mm9",
                        "chr_mm10","start_mm10","end_mm10",	"CpG_nb","pvalue",
                        "qvalue","meth.diff","CGi_chr",  "CGi_start", "CGi_end", 
                        "CGi_name","CGi_length", "CGi_cpgNum", "CGi_overlap")

DMR_Annot_clean = DMR_Annot[,c("mm10_chr","DMR_start","DMR_end","DMR_ID",
                               "chr_annot","start_annot","end_annot","strand",
                               "entrezgene_accession","gene_biotype","mgi_symbol",
                               "ENSEMBL_ID","Entrez_ID","overlap_annot_length",
                               "chr_mm9","start_mm9","end_mm9","CpG_nb","pvalue",
                               "qvalue","meth.diff","CGi_chr",  "CGi_start", 
                               "CGi_end", "CGi_name","CGi_length", "CGi_cpgNum", 
                               "CGi_overlap")]


# for distance=0
for(i in 1:7){
  
  DMR_DOCR_sample = list_DMR_DOCR_dist0[i]
  Annot_DMR_DOCR = read.table(file = paste("H:/methylome/fusion/post_methylkit/DMR432_pval0.07/croisement_autre_data/DMR_ATACseq/",DMR_DOCR_sample,sep="/"), 
                              header = FALSE, sep = "\t", quote="", fill=TRUE)
  
  # To rename columns
  colnames(Annot_DMR_DOCR) <- c("DMR_chr","DMR_start", "DMR_end", "DOCR_chr", "DOCR_start", "DOCR_end", "DMR_DOCR_overlap")
  
  # To create 2 ID : DMR_ID et DOCR_ID
  Annot_DMR_DOCR$DMR_ID = paste(Annot_DMR_DOCR$DMR_chr, Annot_DMR_DOCR$DMR_start, Annot_DMR_DOCR$DMR_end, sep=";")
  Annot_DMR_DOCR$DOCR_ID = paste(Annot_DMR_DOCR$DOCR_chr, Annot_DMR_DOCR$DOCR_start, Annot_DMR_DOCR$DOCR_end, sep=";")
  
  # To Load Annotation informations of all DOCR
  DMR_sample <- DMR_Annot_clean
  
  Annot_DOCR_sample = list_DOCR_Annot[i]
  Annot_DOCR = read.table(file = paste("G:/projet_DU_DSD/results/10_annotations/Sartools/Biomart/mm10",Annot_DOCR_sample,sep="/"), header = TRUE, sep = "\t", quote="", fill=TRUE)
  
  colnames(Annot_DOCR) = c("mm9_chr", "mm9_start", "mm9_end", "mm9_ID", "mm10_chr", 
                           "mm10_start", "mm10_end", "mm10_ID", "chr_gene", 
                           "ensembl_gene_id", "entrezgene_id", "overlap_length", 
                           "cond1_rep1", "cond1_rep2", "cond2_rep1", "cond2_rep2", 
                           "norm.cond1_rep1", "norm.cond1_rep2", "norm.cond2_rep1", 
                           "norm.cond2_rep2", "baseMean", "cond1", "cond2", "FC", 
                           "log2FoldChange", "pvalue", "padj", "tagwise.dispersion", 
                           "trended.dispersion")
    
  Annot_DOCR$DOCR_ID=paste(Annot_DOCR$mm10_chr, Annot_DOCR$mm10_start, Annot_DOCR$mm10_end,sep=";")
  
  Annot_DMR_DOCR=merge(Annot_DMR_DOCR, DMR_sample, by="DMR_ID")
  Annot_DMR_DOCR=merge(Annot_DMR_DOCR, Annot_DOCR, by="DOCR_ID")
  
  # To remove duplicate columns
  Annot_DMR_DOCR_clean = Annot_DMR_DOCR[,c("DMR_chr", "DMR_start.x", "DMR_end.x", "DMR_ID","chr_annot" , "start_annot", "end_annot", "strand.x",
                                           "entrezgene_accession.x", "gene_biotype.x", "mgi_symbol.x", "ENSEMBL_ID", "Entrez_ID",
                                           "overlap_annot_length", "chr_mm9" , "start_mm9", "end_mm9" , "CpG_nb", "pvalue.x" ,  "qvalue" , "meth.diff",
                                           "CGi_chr" ,"CGi_start" , "CGi_end" ,"CGi_name", "CGi_length" ,"CGi_cpgNum", "CGi_overlap",
                                           "DMR_DOCR_overlap", "DOCR_chr", "DOCR_start", "DOCR_end","DOCR_ID" ,
                                           "mm9_chr", "mm9_start", "mm9_end", "mm9_ID","cond1_rep1", "cond1_rep2", "cond2_rep1" , "cond2_rep2", 
                                           "norm.cond1_rep1", "norm.cond1_rep2",  "norm.cond2_rep1" , "norm.cond2_rep2",  "baseMean",  "cond1" , "cond2" ,  "FC",
                                           "log2FoldChange", "pvalue.y",  "padj", "tagwise.dispersion", "trended.dispersion",
                                           "strand.y", "chr_gene", "start_gene", "end_gene", "entrezgene_accession.y", "gene_biotype.y", 
                                           "mgi_symbol.y" , "ensembl_gene_id", "entrezgene_id", "overlap_length")]
  
       
  colnames(Annot_DMR_DOCR_clean) = c("DMR_mm10_chr", "DMR_mm10_start", "DMR_mm10_end", "DMR_mm10_ID","mm10_chr_annot_DMR" , "mm10_start_annot_DMR", "mm10_end_annot_DMR", 
                                     "mm_10_strand_annot_DMR",
                                     "entrezgene_accession_annot_DMR", "gene_biotype_annot_DMR", "mgi_symbol_annot_DMR", "ENSEMBL_ID_annot_DMR", "Entrez_ID_annot_DMR",
                                     "overlap_annot_DMR_length", "DMR_mm9_chr" , "DMR_mm9_start", "DMR_mm9_end" , "DMR_CpG_nb", "DMR_pvalue" ,  "DMR_qvalue" , "DMR_meth.diff" ,
                                     "CGi_chr" ,"CGi_start" , "CGi_end" ,"CGi_name", "CGi_length" ,"CGi_cpgNum", "CGi_overlap",
                                     "DMR_DOCR_overlap", "DOCR_mm10_chr", "DOCR_mm10_start", "DOCR_mm10_end","DOCR_mm10_ID",
                                     "DOCR_mm9_chr", "DOCR_mm9_start", "DOCR_mm9_end", "DOCR_mm9_ID","DOCR_cond1_rep1", "DOCR_cond1_rep2", 
                                     "DOCR_cond2_rep1" , "DOCR_cond2_rep2",
                                     "DOCR_norm.cond1_rep1", "DOCR_norm.cond1_rep2",  "DOCR_norm.cond2_rep1" , "DOCR_norm.cond2_rep2",  "DOCR_baseMean",  "DOCR_cond1" , "DOCR_cond2" ,
                                     "DOCR_FC",
                                     "DOCR_log2FoldChange", "DOCR_pvalue",  "DOCR_padj", "DOCR_tagwise.dispersion", "DOCR_trended.dispersion", 
                                     "strand_annot_DOCR", "mm10_chr_gene_annot_DOCR", "mm10_start_gene_annot_DOCR","mm10_end_gene_annot_DOCR", 
                                     "entrezgene_accession_annot_DOCR", "gene_biotype_annot_DOCR", 
                                     "mgi_symbol_annot_DOCR" , "ensembl_gene_id_annot_DOCR", "entrezgene_id_annot_DOCR", "overlap_length_annot_DOCR")                                   
  
  
  write.table(Annot_DMR_DOCR_clean, file=paste("H:/methylome/fusion/post_methylkit/DMR432_pval0.07/croisement_autre_data/DMR_ATACseq/clean_annot",DMR_DOCR_sample,sep="_"), quote=FALSE, sep="\t", row.names = FALSE, col.names = TRUE)
  write.table(Annot_DMR_DOCR, file=paste("H:/methylome/fusion/post_methylkit/DMR432_pval0.07/croisement_autre_data/DMR_ATACseq/raw_annot",DMR_DOCR_sample,sep="_"), quote=FALSE, sep="\t", row.names = FALSE, col.names = TRUE)
  
  
  # To keep only gene names (entrezgene_accession_annot_DMR)
  # annotation of DMR - mm10 BIOMART : 
  
  # To uncollapse data 
  decondat_Annot_DMR_DOCR_clean = as.character(Annot_DMR_DOCR_clean$entrezgene_accession_annot_DMR)
  print(DMR_DOCR_sample)
  print(head(decondat_Annot_DMR_DOCR_clean)) 
  
  # To obtain list of gene names
  list_gene_splitted <- strsplit(as.character(Annot_DMR_DOCR_clean$entrezgene_accession_annot_DMR),split=";")
  vec_gene_splitted <- unlist(list_gene_splitted)
  
  # To remove "." : 
  vec_gene_splitted_filter = vec_gene_splitted[vec_gene_splitted!="."]
  
  # To save data in a table
  table_VGS_filter = as.data.frame(unique(vec_gene_splitted_filter))
  
  write.table(table_VGS_filter, paste("H:/methylome/fusion/post_methylkit/DMR432_pval0.07/croisement_autre_data/DMR_ATACseq/list_gene",DMR_DOCR_sample, sep="_"), quote=FALSE, col.names=FALSE, row.names=FALSE)
  
}


rm(list = ls())

#-------------------
# Idem for distance=1000
# To load files or list of files
# To load DMR+DOCR regions
list_DMR_DOCR = list.files("H:/methylome/fusion/post_methylkit/DMR432_pval0.07/croisement_autre_data/DMR_ATACseq/")
list_DMR_DOCR_dist0 = list_DMR_DOCR[which(grepl("dist0",list_DMR_DOCR))]
list_DMR_DOCR_dist1000 = list_DMR_DOCR[which(grepl("dist1000",list_DMR_DOCR))]

# List of DOCR informations 
# Since some files were empty (no overlap), we need to specify, in a list, which file we want to annotate.
list_DOCR_Annot=list("stats_mm9_mm10_down_DOCR_E14.15_annot_syntaxique_mm10_biomart.bed","stats_mm9_mm10_down_DOCR_E14.16_annot_syntaxique_mm10_biomart.bed",
                     "stats_mm9_mm10_down_DOCR_E16.P0_annot_syntaxique_mm10_biomart.bed", "stats_mm9_mm10_up_DOCR_E14.15_annot_syntaxique_mm10_biomart.bed",
                     "stats_mm9_mm10_up_DOCR_E14.16_annot_syntaxique_mm10_biomart.bed", "stats_mm9_mm10_up_DOCR_E15.16_annot_syntaxique_mm10_biomart.bed",
                     "stats_mm9_mm10_up_DOCR_E16.P0_annot_syntaxique_mm10_biomart.bed")


# To load DMR432 annotation
DMR_Annot = read.table("H:/methylome/fusion/post_methylkit/DMR432_pval0.07/annotations/Biomart/mm10/DMR432_unique_annot_mm10_BMT_meth_cgi.bed", header=FALSE, sep="\t", quote="")
head(DMR_Annot)
dim(DMR_Annot)
colnames(DMR_Annot) = c("mm10_chr",	"DMR_start",	"DMR_end",	"DMR_ID",	"chr_annot",	"start_annot",	"end_annot",	"strand",
                        "entrezgene_accession",	"gene_biotype",	"mgi_symbol",	"ENSEMBL_ID",	"Entrez_ID",	"overlap_annot_length",
                        "chr_mm9",	"start_mm9",	"end_mm9",	"chr_mm10",	"start_mm10",	"end_mm10",	"CpG_nb",	"pvalue",	"qvalue",	"meth.diff",
                        "CGi_chr",  "CGi_start", "CGi_end", "CGi_name","CGi_length", "CGi_cpgNum", "CGi_overlap")

DMR_Annot_clean = DMR_Annot[,c("mm10_chr",	"DMR_start",	"DMR_end",	"DMR_ID","chr_annot",	"start_annot",	"end_annot",	"strand",
                               "entrezgene_accession",	"gene_biotype",	"mgi_symbol",	"ENSEMBL_ID",	"Entrez_ID",	"overlap_annot_length",
                               "chr_mm9",	"start_mm9",	"end_mm9",	"CpG_nb",	"pvalue",	"qvalue",	"meth.diff",
                               "CGi_chr",  "CGi_start", "CGi_end", "CGi_name","CGi_length", "CGi_cpgNum", "CGi_overlap")]


# for distance = 1000
for(i in 1:7){
  
  DMR_DOCR_sample = list_DMR_DOCR_dist1000[i]
  Annot_DMR_DOCR = read.table(file = paste("H:/methylome/fusion/post_methylkit/DMR432_pval0.07/croisement_autre_data/DMR_ATACseq/",DMR_DOCR_sample,sep="/"), 
                              header = FALSE, sep = "\t", quote="", fill=TRUE)
  
  # To rename columns
  colnames(Annot_DMR_DOCR) <- c("DMR_chr","DMR_start", "DMR_end", "DOCR_chr", "DOCR_start", "DOCR_end", "DMR_DOCR_overlap")
  
  # To create 2 ID : DMR_ID et DOCR_ID
  Annot_DMR_DOCR$DMR_ID = paste(Annot_DMR_DOCR$DMR_chr, Annot_DMR_DOCR$DMR_start, Annot_DMR_DOCR$DMR_end, sep=";")
  Annot_DMR_DOCR$DOCR_ID = paste(Annot_DMR_DOCR$DOCR_chr, Annot_DMR_DOCR$DOCR_start, Annot_DMR_DOCR$DOCR_end, sep=";")
  
  # To load annotation informations of all DOCR
  DMR_sample <- DMR_Annot_clean
  
  Annot_DOCR_sample = list_DOCR_Annot[i]
  Annot_DOCR = read.table(file = paste("G:/projet_DU_DSD/results/10_annotations/Sartools/Biomart/mm10",Annot_DOCR_sample,sep="/"), header = TRUE, sep = "\t", quote="", fill=TRUE)
  
  colnames(Annot_DOCR) = c("mm9_chr", "mm9_start", "mm9_end", "mm9_ID", "mm10_chr", "mm10_start", "mm10_end", "mm10_ID", 
                           "chr_gene", "start_gene", "end_gene", "strand", "entrezgene_accession", "gene_biotype", "mgi_symbol", 
                           "ensembl_gene_id", "entrezgene_id", "overlap_length", "cond1_rep1", "cond1_rep2", "cond2_rep1", "cond2_rep2", 
                           "norm.cond1_rep1", "norm.cond1_rep2", "norm.cond2_rep1", "norm.cond2_rep2", "baseMean", "cond1", "cond2", "FC", 
                           "log2FoldChange", "pvalue", "padj", "tagwise.dispersion", "trended.dispersion")
  
  
  Annot_DOCR$DOCR_ID=paste(Annot_DOCR$mm10_chr, Annot_DOCR$mm10_start, Annot_DOCR$mm10_end,sep=";")
  
  Annot_DMR_DOCR=merge(Annot_DMR_DOCR, DMR_sample, by="DMR_ID")
  Annot_DMR_DOCR=merge(Annot_DMR_DOCR, Annot_DOCR, by="DOCR_ID")
  
  # To remove duplicate columns
  Annot_DMR_DOCR_clean = Annot_DMR_DOCR[,c("DMR_chr", "DMR_start.x", "DMR_end.x", "DMR_ID","chr_annot" , "start_annot", "end_annot", "strand.x",
                                           "entrezgene_accession.x", "gene_biotype.x", "mgi_symbol.x", "ENSEMBL_ID", "Entrez_ID",
                                           "overlap_annot_length", "chr_mm9" , "start_mm9", "end_mm9" , "CpG_nb", "pvalue.x" ,  "qvalue" , "meth.diff",
                                           "CGi_chr" ,"CGi_start" , "CGi_end" ,"CGi_name", "CGi_length" ,"CGi_cpgNum", "CGi_overlap",
                                           "DMR_DOCR_overlap", "DOCR_chr", "DOCR_start", "DOCR_end","DOCR_ID" ,
                                           "mm9_chr", "mm9_start", "mm9_end", "mm9_ID","cond1_rep1", "cond1_rep2", "cond2_rep1" , "cond2_rep2", 
                                           "norm.cond1_rep1", "norm.cond1_rep2",  "norm.cond2_rep1" , "norm.cond2_rep2",  "baseMean",  "cond1" , "cond2" ,  "FC",
                                           "log2FoldChange", "pvalue.y",  "padj", "tagwise.dispersion", "trended.dispersion",
                                           "strand.y", "chr_gene", "start_gene", "end_gene", "entrezgene_accession.y", "gene_biotype.y", 
                                           "mgi_symbol.y" , "ensembl_gene_id", "entrezgene_id", "overlap_length")]
  
  
  
  
  
  colnames(Annot_DMR_DOCR_clean) = c("DMR_mm10_chr", "DMR_mm10_start", "DMR_mm10_end", "DMR_mm10_ID","mm10_chr_annot_DMR" , "mm10_start_annot_DMR", "mm10_end_annot_DMR", 
                                     "mm_10_strand_annot_DMR",
                                     "entrezgene_accession_annot_DMR", "gene_biotype_annot_DMR", "mgi_symbol_annot_DMR", "ENSEMBL_ID_annot_DMR", "Entrez_ID_annot_DMR",
                                     "overlap_annot_DMR_length", "DMR_mm9_chr" , "DMR_mm9_start", "DMR_mm9_end" , "DMR_CpG_nb", "DMR_pvalue" ,  "DMR_qvalue" , "DMR_meth.diff" ,
                                     "CGi_chr" ,"CGi_start" , "CGi_end" ,"CGi_name", "CGi_length" ,"CGi_cpgNum", "CGi_overlap",
                                     "DMR_DOCR_overlap", "DOCR_mm10_chr", "DOCR_mm10_start", "DOCR_mm10_end","DOCR_mm10_ID",
                                     "DOCR_mm9_chr", "DOCR_mm9_start", "DOCR_mm9_end", "DOCR_mm9_ID","DOCR_cond1_rep1", "DOCR_cond1_rep2", 
                                     "DOCR_cond2_rep1" , "DOCR_cond2_rep2",
                                     "DOCR_norm.cond1_rep1", "DOCR_norm.cond1_rep2",  "DOCR_norm.cond2_rep1" , "DOCR_norm.cond2_rep2",  "DOCR_baseMean",  "DOCR_cond1" , "DOCR_cond2" ,
                                     "DOCR_FC",
                                     "DOCR_log2FoldChange", "DOCR_pvalue",  "DOCR_padj", "DOCR_tagwise.dispersion", "DOCR_trended.dispersion", 
                                     "strand_annot_DOCR", "mm10_chr_gene_annot_DOCR", "mm10_start_gene_annot_DOCR","mm10_end_gene_annot_DOCR", 
                                     "entrezgene_accession_annot_DOCR", "gene_biotype_annot_DOCR", 
                                     "mgi_symbol_annot_DOCR" , "ensembl_gene_id_annot_DOCR", "entrezgene_id_annot_DOCR", "overlap_length_annot_DOCR")                                   
  
  
  write.table(Annot_DMR_DOCR_clean, file=paste("H:/methylome/fusion/post_methylkit/DMR432_pval0.07/croisement_autre_data/DMR_ATACseq/clean_annot",DMR_DOCR_sample,sep="_"), quote=FALSE, sep="\t", row.names = FALSE, col.names = TRUE)
  write.table(Annot_DMR_DOCR, file=paste("H:/methylome/fusion/post_methylkit/DMR432_pval0.07/croisement_autre_data/DMR_ATACseq/raw_annot",DMR_DOCR_sample,sep="_"), quote=FALSE, sep="\t", row.names = FALSE, col.names = TRUE)
  
  
  # To keep only gene names (entrezgene_accession_annot_DMR) - annotation of DMR mm10 BIOMART : 
  
  # To uncollapse data 
  decondat_Annot_DMR_DOCR_clean = as.character(Annot_DMR_DOCR_clean$entrezgene_accession_annot_DMR)
  print(DMR_DOCR_sample)
  print(head(decondat_Annot_DMR_DOCR_clean)) 
  
  # To obtain list of genes
  list_gene_splitted <- strsplit(as.character(Annot_DMR_DOCR_clean$entrezgene_accession_annot_DMR),split=";")
  vec_gene_splitted <- unlist(list_gene_splitted)
  
  # To remove "." : 
  vec_gene_splitted_filter = vec_gene_splitted[vec_gene_splitted!="."]
  
  # To save data in a table
  table_VGS_filter = as.data.frame(unique(vec_gene_splitted_filter))
  
  write.table(table_VGS_filter, paste("H:/methylome/fusion/post_methylkit/DMR432_pval0.07/croisement_autre_data/DMR_ATACseq/list_gene",DMR_DOCR_sample, sep="_"), quote=FALSE, col.names=FALSE, row.names=FALSE)
  
}
```

#### Integration of DMR with DEG¶

To determine whether DMRs are located within genes that are differentially expressed (DEG) during physiological brain development, DMRs and DEG regions were compared using our own R function called ***find\_overlaps\_AD\_table()***. Both overlaping regions and regions less than 1000 bases apart were investigated, by setting ***tolerance*** argument to 0 or 1000. The obtained regions were then annotated, using annotations that were already achieved in the two complete datasets.

In [ ]:

```
%%R

####################################################################
# Integration of DMR and DEG to find potential overlap
# (using mm10 chromosomal coordinates)
####################################################################

# 1.Load DMR432 data - mm10 
DMR432 = read.table("H:/methylome/fusion/post_methylkit/DMR432_pval0.07/annotations/Biomart/mm10/DMR432_unique_annot_mm10_BMT_meth_cgi.bed", sep="\t", header = FALSE)
head(DMR432)

# To extract DMR432 chromosomic coordinates
DMR432_bed = DMR432[,c("V1","V2","V3")]
head(DMR432_bed)
dim(DMR432_bed)


# 2. Load DEG data - mm10 (from edgeR analysis):
nom_sample = list=c("down_E13.14", "down_E14.15", "down_E14.16", "down_E15.16", 
                    "down_E16.P0", "up_E13.14", "up_E14.15", "up_E14.16", 
                    "up_E15.16", "up_E16.P0" )

# /!\ /!\ Dataset must be in a matrix format, otherwise, function doesn't work /!\/!\
for(name in nom_sample){
  assign(x=name, value = as.matrix(read.table(paste("G:/projet_DU_AD/results/09_annotation/Sartools/Biomart/bed_edge_AnnotBM_et_Man_ech_",  
                                          name , "_mm10.sorted.bed", sep=""), sep = "\t", header= FALSE, dec=".", quote="")))
  print(paste("head_",name, sep=""))
  print(head(eval(parse(text = name))))
  print(paste("dimension du dataset ",name, sep = ""))
  print(dim(eval(parse(text = name))))
}

# Creation of a list containing each DEG file (up or down, each developmental stage) 
RNA = list(downE13.14 = down_E13.14, downE14.15 = down_E14.15, downE15.16 = down_E15.16, downE14.16 = down_E14.16, downE16.P0 = down_E16.P0,
           upE13.14 = up_E13.14, upE14.15 = up_E14.15 ,upE15.16 = up_E15.16, upE14.16 = up_E14.16, upE16.P0 = up_E16.P0)

for (i in 1:length(RNA)){
  print(names(RNA)[i])
  print(head(RNA[[i]]))
  print(dim(RNA[[i]]))
} 


# 3. Function to find overlap or close regions (depending on value given for tolerance parameter)
find_overlaps_AD_table <- function (regions_1,regions_2,tolerance=0)
{
  common_region_table = NULL
  number_of_regions_1 <- nrow(regions_1)
  number_of_regions_2 <- nrow(regions_2)
  
  overlapped_is <- c()
  overlapped_js <- c()
  
  for (i in 1:number_of_regions_1) {
    
    for (j in 1:number_of_regions_2) {
      
      if (as.character(regions_1[i,1])==as.character(regions_2[j,1])) {
        start_1 <- as.numeric(regions_1[i,2])
        end_1 <- as.numeric(regions_1[i,3])
        expanded_start_1 <- start_1 - tolerance
        expanded_end_1 <- end_1 + tolerance
        start_2 <- as.numeric(regions_2[j,2])
        end_2 <- as.numeric(regions_2[j,3])
        
        if (expanded_end_1>=start_2) {
          
          if (end_2>=expanded_start_1) {
            
            overlapped_is <- c(overlapped_is,i)
            overlapped_js <- c(overlapped_js,j)
            
            first_region <- paste(regions_1[i,1],start_1,end_1,sep="_")
            second_region <- paste(regions_2[j,1],start_2,end_2,sep="_")
            overlap_size <- min(end_1,end_2)-max(start_1,start_2)+1
            common_region_table = rbind(common_region_table,c(as.character(regions_1[i,1]),start_1,end_1, 
                                                              as.character(regions_2[j,1]),start_2,end_2, overlap_size))
            
            if (overlap_size>0) {
              print(paste(first_region,second_region,sep=" "))
            } else {
              print(paste(first_region,second_region, sep=" "))
            }
          }
        } 
      }
    }
  }
  print(paste("Number matched in first data set:",length(unique(overlapped_is))))
  print(paste("Number matched in second data set:",length(unique(overlapped_js))))
  if (length(common_region_table) > 0){
    colnames(common_region_table) = c("chr_reg1", "start_reg1", "end_reg1", "chr_reg2", "start_reg2", "end_reg2","overlap_size")
  }else{
  }
  return(as.data.frame(common_region_table))
}


# 4. Search for common regions between DMR and DEG - overlap
for (i in 1:length(RNA)) {
  print(names(RNA)[i])
  DMR_RNA = find_overlaps_AD_table(DMR432_bed, RNA[[i]], tolerance=0)
  write.table(DMR_RNA,paste("H:/methylome/fusion/post_methylkit/DMR432_pval0.07/croisement_autre_data/mm10_DMR432", names(RNA)[i], "dist0.txt",sep="_"), sep = "\t", quote=FALSE, row.names = FALSE, col.names=FALSE)
}

# 5. Search for common regions between DMR and DEG - maximum distance between 2 regions = 1000bases: 
for (i in 1:length(RNA)) {
  print(names(RNA)[i])
  DMR_RNA = find_overlaps_AD_table(DMR432_bed, RNA[[i]], tolerance=1000)
  write.table(DMR_RNA,paste("H:/methylome/fusion/post_methylkit/DMR432_pval0.07/croisement_autre_data/mm10_DMR432", names(RNA)[i], "dist1000.txt",sep="_"), sep = "\t", quote=FALSE, row.names = FALSE, col.names=FALSE)
}


###################
# Annotation
###################

# Load files or list of files
# Load DMR+DEG regions
list_DMR_DEG = list.files("H:/methylome/fusion/post_methylkit/DMR432_pval0.07/croisement_autre_data/DMR_RNAseq/")
list_DMR_DEG_dist0 = list_DMR_DEG[which(grepl("dist0",list_DMR_DEG))]
list_DMR_DEG_dist1000 = list_DMR_DEG[which(grepl("dist1000",list_DMR_DEG))]

# Load DEG annotation
list_files_DEG = list.files("G:/projet_DU_AD/results/09_annotation/Sartools/Biomart/")
list_DEG_Annot = list_files_DEG[which(grepl("edgeR_AnnotBM_et_Man",list_files_DEG))]

# Load DMR432 annotation
DMR_Annot = read.table("H:/methylome/fusion/post_methylkit/DMR432_pval0.07/annotations/Biomart/mm10/DMR432_unique_annot_mm10_BMT_meth_cgi.bed", header=FALSE, sep="\t", quote="")
head(DMR_Annot)
dim(DMR_Annot)
colnames(DMR_Annot) = c("mm10_chr",	"DMR_start",	"DMR_end",	"DMR_ID",	"chr_annot",	"start_annot",	"end_annot",	"strand",
                        "entrezgene_accession",	"gene_biotype",	"mgi_symbol",	"ENSEMBL_ID",	"Entrez_ID",	"overlap_annot_length",
                        "chr_mm9",	"start_mm9",	"end_mm9",	"chr_mm10",	"start_mm10",	"end_mm10",	"CpG_nb",	"pvalue",	"qvalue",	"meth.diff",
                        "CGi_chr",  "CGi_start", "CGi_end", "CGi_name","CGi_length", "CGi_cpgNum", "CGi_overlap")

DMR_Annot_clean = DMR_Annot[,c("mm10_chr",	"DMR_start",	"DMR_end",	"DMR_ID","chr_annot",	"start_annot",	"end_annot",	"strand",
                               "entrezgene_accession",	"gene_biotype",	"mgi_symbol",	"ENSEMBL_ID",	"Entrez_ID",	"overlap_annot_length",
                               "chr_mm9",	"start_mm9",	"end_mm9",	"CpG_nb",	"pvalue",	"qvalue",	"meth.diff",
                               "CGi_chr",  "CGi_start", "CGi_end", "CGi_name","CGi_length", "CGi_cpgNum", "CGi_overlap")]


# for distance=0
for(i in 1:10){
  
  DMR_DEG_sample = list_DMR_DEG_dist0[i]
  Annot_DMR_DEG = read.table(file = paste("H:/methylome/fusion/post_methylkit/DMR432_pval0.07/croisement_autre_data/DMR_RNAseq/",DMR_DEG_sample,sep="/"), 
                             header = FALSE, sep = "\t", quote="", fill=TRUE)
  
  # Change column names
  colnames(Annot_DMR_DEG) <- c("DMR_chr","DMR_start", "DMR_end", "DEG_chr", "DEG_start", "DEG_end", "DMR_DEG_overlap")
  
  # Create ID : DMR_ID et DEG_ID
  Annot_DMR_DEG$DMR_ID = paste(Annot_DMR_DEG$DMR_chr, Annot_DMR_DEG$DMR_start, Annot_DMR_DEG$DMR_end, sep=";")
  Annot_DMR_DEG$DEG_ID = paste(Annot_DMR_DEG$DEG_chr, Annot_DMR_DEG$DEG_start, Annot_DMR_DEG$DEG_end, sep=";")
  
  # Load Annot of DEG
  DMR_sample <- DMR_Annot_clean
  DEG_sample <- list_DEG_Annot[i]
  Annot_DEG = read.table(file = paste("G:/projet_DU_AD/results/09_annotation/Sartools/Biomart",DEG_sample,sep="/"), header = TRUE, sep = "\t", quote="", fill=TRUE)
  
  colnames(Annot_DEG)= c("ENS_ID", "cond1_rep1", "cond1_rep2", "cond2_rep1", "cond2_rep2", 
                         "norm.cond1_rep1", "norm.cond1_rep2", "norm.cond2_rep1", "norm.cond2_rep2", 
                         "baseMean", "cond1", "cond2", "FC", "log2FoldChange", "pvalue", "padj", "tagwise.dispersion", 
                         "trended.dispersion", "mm10_chr", "strand", "mm10_start_gene", "mm10_end_gene", "entrezgene_id", "gene_biotype", 
                         "mgi_symbol", "entrezgene_accession", "entrezgene_description", "uniprot_gn_symbol")
  
  Annot_DEG$DEG_ID=paste(paste("chr",Annot_DEG$mm10_chr,sep=""), Annot_DEG$mm10_start_gene, Annot_DEG$mm10_end_gene,sep=";")
  
  Annot_DMR_DEG=merge(Annot_DMR_DEG, DMR_sample, by="DMR_ID")
  Annot_DMR_DEG=merge(Annot_DMR_DEG, Annot_DEG, by="DEG_ID")
  
  # Remove duplicate columns
  Annot_DMR_DEG_clean = Annot_DMR_DEG[,c("DMR_chr", "DMR_start.x", "DMR_end.x", "DMR_ID", "chr_annot" , "start_annot", "end_annot", "strand.x",
                                         "entrezgene_accession.x", "gene_biotype.x", "mgi_symbol.x", "ENSEMBL_ID", "Entrez_ID",
                                         "overlap_annot_length", "chr_mm9" , "start_mm9", "end_mm9" , "CpG_nb", "pvalue.x" ,  "qvalue" , "meth.diff" ,
                                         "CGi_chr" ,"CGi_start" , "CGi_end" ,"CGi_name", "CGi_length" ,"CGi_cpgNum", "CGi_overlap",
                                         "DMR_DEG_overlap", "DEG_chr", "DEG_start", "DEG_end","DEG_ID" ,   "ENS_ID" , "cond1_rep1", "cond1_rep2", "cond2_rep1" , "cond2_rep2",
                                         "norm.cond1_rep1", "norm.cond1_rep2",  "norm.cond2_rep1" , "norm.cond2_rep2",  "baseMean",  "cond1" , "cond2" ,  "FC"  ,
                                          "log2FoldChange", "pvalue.y",  "padj", "tagwise.dispersion", "trended.dispersion", "strand.y" ,              
                                          "mm10_start_gene","mm10_end_gene", "entrezgene_id", "gene_biotype.y" , "mgi_symbol.y" , "entrezgene_accession.y", "entrezgene_description",
                                        "uniprot_gn_symbol")]

  colnames(Annot_DMR_DEG_clean) = c("DMR_mm10_chr", "DMR_mm10_start", "DMR_mm10_end", "DMR_mm10_ID", "mm10_chr_gene_annot_DMR" , "mm10_start_gene_annot_DMR", "mm10_end_gene_annot_DMR", 
                                   "strand_gene_annot_DMR", "entrezgene_accession_annot_DMR", "gene_biotype_annot_DMR", "mgi_symbol_annot_DMR", "ENSEMBL_ID_annot_DMR", "Entrez_ID_annot_DMR",
                                         "overlap_annot_DMR_length", "DMR_mm9_chr" , "DMR_mm9_start", "DMR_mm9_end" , "DMR_CpG_nb", "DMR_pvalue" ,  "DMR_qvalue" , "DMR_meth.diff" ,
                                         "CGi_chr" ,"CGi_start" , "CGi_end" ,"CGi_name", "CGi_length" ,"CGi_cpgNum", "CGi_overlap",
                                         "DMR_DEG_overlap", "DEG_mm10_chr", "DEG_mm10_start", "DEG_mm10_end","DEG_mm10_ID" ,   "DEG_ENS_ID" , "DEG_cond1_rep1", "DEG_cond1_rep2", 
                                   "DEG_cond2_rep1" , "DEG_cond2_rep2",
                                         "DEG_norm.cond1_rep1", "DEG_norm.cond1_rep2",  "DEG_norm.cond2_rep1" , "DEG_norm.cond2_rep2",  "DEG_baseMean",  "DEG_cond1" , "DEG_cond2" , 
                                   "DEG_FC",
                                         "DEG_log2FoldChange", "DEG_pvalue",  "DEG_padj", "DEG_tagwise.dispersion", "DEG_trended.dispersion", "DEG_strand" ,              
                                         "mm10_start_gene_annot_DEG","mm10_end_gene_annot_DEG", "entrezgene_id_annot_DEG", "gene_biotype_annot_DEG" , "mgi_symbol_annot_DEG" , 
                                   "entrezgene_accession_annot_DEG", "entrezgene_description_annot_DEG",
                                         "uniprot_gn_symbol_annot_DEG")                                   
                                        
                       

  write.table(Annot_DMR_DEG_clean, file=paste("H:/methylome/fusion/post_methylkit/DMR432_pval0.07/croisement_autre_data/DMR_RNAseq/clean_annot",DMR_DEG_sample,sep="_"), quote=FALSE, sep="\t", row.names = FALSE, col.names = TRUE)
  write.table(Annot_DMR_DEG, file=paste("H:/methylome/fusion/post_methylkit/DMR432_pval0.07/croisement_autre_data/DMR_RNAseq/raw_annot",DMR_DEG_sample,sep="_"), quote=FALSE, sep="\t", row.names = FALSE, col.names = TRUE)
  
  
  # To keep only gene names (entrezgene_accession_annot_DMR) - annotation of DMR - mm10 - BIOMART  : 
  
  # To uncollapse data
  decondat_Annot_DMR_DEG_clean = as.character(Annot_DMR_DEG_clean$entrezgene_accession_annot_DMR)
  print(DMR_DEG_sample)
  print(head(decondat_Annot_DMR_DEG_clean)) 
  
  # To split gene names, when necessary
  list_gene_splitted <- strsplit(as.character(Annot_DMR_DEG_clean$entrezgene_accession_annot_DMR),split=";")
  vec_gene_splitted <- unlist(list_gene_splitted)
  
  # To remove "." : 
  vec_gene_splitted_filter = vec_gene_splitted[vec_gene_splitted!="."]
  
  #To save data in a table
  table_VGS_filter = as.data.frame(unique(vec_gene_splitted_filter))
  
  write.table(table_VGS_filter, paste("H:/methylome/fusion/post_methylkit/DMR432_pval0.07/croisement_autre_data/DMR_RNAseq/list_gene",DMR_DEG_sample, sep="_"), quote=FALSE, col.names=FALSE, row.names=FALSE)
  
}


rm(list = ls())


#--------------------------
# Idem for distance=1000
# Load files or list of files
# Load DMR+DEG regions
list_DMR_DEG = list.files("H:/methylome/fusion/post_methylkit/DMR432_pval0.07/croisement_autre_data/DMR_RNAseq/")
list_DMR_DEG_dist0 = list_DMR_DEG[which(grepl("dist0",list_DMR_DEG))]
list_DMR_DEG_dist1000 = list_DMR_DEG[which(grepl("dist1000",list_DMR_DEG))]

# Load DEG annotation
list_files_DEG = list.files("G:/projet_DU_AD/results/09_annotation/Sartools/Biomart/")
list_DEG_Annot = list_files_DEG[which(grepl("edgeR_AnnotBM_et_Man",list_files_DEG))]

# Load DMR432 annotation
DMR_Annot = read.table("H:/methylome/fusion/post_methylkit/DMR432_pval0.07/annotations/Biomart/mm10/DMR432_unique_annot_mm10_BMT_meth_cgi.bed", header=FALSE, sep="\t", quote="")
head(DMR_Annot)
dim(DMR_Annot)
colnames(DMR_Annot) = c("mm10_chr",	"DMR_start",	"DMR_end",	"DMR_ID",	"chr_annot",	"start_annot",	"end_annot",	"strand",
                        "entrezgene_accession",	"gene_biotype",	"mgi_symbol",	"ENSEMBL_ID",	"Entrez_ID",	"overlap_annot_length",
                        "chr_mm9",	"start_mm9",	"end_mm9",	"chr_mm10",	"start_mm10",	"end_mm10",	"CpG_nb",	"pvalue",	"qvalue",	"meth.diff",
                        "CGi_chr",  "CGi_start", "CGi_end", "CGi_name","CGi_length", "CGi_cpgNum", "CGi_overlap")

DMR_Annot_clean = DMR_Annot[,c("mm10_chr",	"DMR_start",	"DMR_end",	"DMR_ID","chr_annot",	"start_annot",	"end_annot",	"strand",
                               "entrezgene_accession",	"gene_biotype",	"mgi_symbol",	"ENSEMBL_ID",	"Entrez_ID",	"overlap_annot_length",
                               "chr_mm9",	"start_mm9",	"end_mm9",	"CpG_nb",	"pvalue",	"qvalue",	"meth.diff",
                               "CGi_chr",  "CGi_start", "CGi_end", "CGi_name","CGi_length", "CGi_cpgNum", "CGi_overlap")]


for(i in 1:10){
  
  DMR_DEG_sample = list_DMR_DEG_dist1000[i]
  Annot_DMR_DEG = read.table(file = paste("H:/methylome/fusion/post_methylkit/DMR432_pval0.07/croisement_autre_data/DMR_RNAseq/",DMR_DEG_sample,sep="/"), 
                             header = FALSE, sep = "\t", quote="", fill=TRUE)
  
  # To rename columns
  colnames(Annot_DMR_DEG) <- c("DMR_chr","DMR_start", "DMR_end", "DEG_chr", "DEG_start", "DEG_end", "DMR_DEG_overlap")
  
  # To create 2 ID : DMR_ID et DEG_ID
  Annot_DMR_DEG$DMR_ID = paste(Annot_DMR_DEG$DMR_chr, Annot_DMR_DEG$DMR_start, Annot_DMR_DEG$DMR_end, sep=";")
  Annot_DMR_DEG$DEG_ID = paste(Annot_DMR_DEG$DEG_chr, Annot_DMR_DEG$DEG_start, Annot_DMR_DEG$DEG_end, sep=";")
  
  # Load Annotation for DEG
  DMR_sample <- DMR_Annot_clean
  DEG_sample <- list_DEG_Annot[i]
  Annot_DEG = read.table(file = paste("G:/projet_DU_AD/results/09_annotation/Sartools/Biomart",DEG_sample,sep="/"), header = TRUE, sep = "\t", quote="", fill=TRUE)
  
  colnames(Annot_DEG)= c("ENS_ID", "cond1_rep1", "cond1_rep2", "cond2_rep1", "cond2_rep2", 
                         "norm.cond1_rep1", "norm.cond1_rep2", "norm.cond2_rep1", "norm.cond2_rep2", 
                         "baseMean", "cond1", "cond2", "FC", "log2FoldChange", "pvalue", "padj", "tagwise.dispersion", 
                         "trended.dispersion", "mm10_chr", "strand", "mm10_start_gene", "mm10_end_gene", "entrezgene_id", "gene_biotype", 
                         "mgi_symbol", "entrezgene_accession", "entrezgene_description", "uniprot_gn_symbol")
  
  Annot_DEG$DEG_ID=paste(paste("chr",Annot_DEG$mm10_chr,sep=""), Annot_DEG$mm10_start_gene, Annot_DEG$mm10_end_gene,sep=";")
  
  Annot_DMR_DEG=merge(Annot_DMR_DEG, DMR_sample, by="DMR_ID")
  Annot_DMR_DEG=merge(Annot_DMR_DEG, Annot_DEG, by="DEG_ID")
  
  # To remove duplicate columns
  Annot_DMR_DEG_clean = Annot_DMR_DEG[,c("DMR_chr", "DMR_start.x", "DMR_end.x", "DMR_ID", "chr_annot" , "start_annot", "end_annot", "strand.x",
                                         "entrezgene_accession.x", "gene_biotype.x", "mgi_symbol.x", "ENSEMBL_ID", "Entrez_ID",
                                         "overlap_annot_length", "chr_mm9" , "start_mm9", "end_mm9" , "CpG_nb", "pvalue.x" ,  "qvalue" , "meth.diff" ,
                                         "CGi_chr" ,"CGi_start" , "CGi_end" ,"CGi_name", "CGi_length" ,"CGi_cpgNum", "CGi_overlap",
                                         "DMR_DEG_overlap", "DEG_chr", "DEG_start", "DEG_end","DEG_ID" ,   "ENS_ID" , "cond1_rep1", "cond1_rep2", "cond2_rep1" , "cond2_rep2",
                                         "norm.cond1_rep1", "norm.cond1_rep2",  "norm.cond2_rep1" , "norm.cond2_rep2",  "baseMean",  "cond1" , "cond2" ,  "FC"  ,
                                         "log2FoldChange", "pvalue.y",  "padj", "tagwise.dispersion", "trended.dispersion", "strand.y" ,              
                                         "mm10_start_gene","mm10_end_gene", "entrezgene_id", "gene_biotype.y" , "mgi_symbol.y" , "entrezgene_accession.y", "entrezgene_description",
                                         "uniprot_gn_symbol")]
  
  colnames(Annot_DMR_DEG_clean) = c("DMR_mm10_chr", "DMR_mm10_start.x", "DMR_mm10_end.x", "DMR_mm10_ID", "mm10_chr_gene_annot_DMR" , "mm10_start_gene_annot_DMR", "mm10_end_gene_annot_DMR", 
                                    "strand_gene_annot_DMR", "entrezgene_accession_annot_DMR", "gene_biotype_annot_DMR", "mgi_symbol_annot_DMR", "ENSEMBL_ID_annot_DMR", "Entrez_ID_annot_DMR",
                                    "overlap_annot_DMR_length", "DMR_mm9_chr" , "DMR_mm9_start", "DMR_mm9_end" , "DMR_CpG_nb", "DMR_pvalue" ,  "DMR_qvalue" , "DMR_meth.diff" ,
                                    "CGi_chr" ,"CGi_start" , "CGi_end" ,"CGi_name", "CGi_length" ,"CGi_cpgNum", "CGi_overlap",
                                    "DMR_DEG_overlap", "DEG_mm10_chr", "DEG_mm10_start", "DEG_mm10_end","DEG_mm10_ID" ,   "DEG_ENS_ID" , "DEG_cond1_rep1", "DEG_cond1_rep2", 
                                    "DEG_cond2_rep1" , "DEG_cond2_rep2",
                                    "DEG_norm.cond1_rep1", "DEG_norm.cond1_rep2",  "DEG_norm.cond2_rep1" , "DEG_norm.cond2_rep2",  "DEG_baseMean",  "DEG_cond1" , "DEG_cond2" , 
                                    "DEG_FC",
                                    "DEG_log2FoldChange", "DEG_pvalue",  "DEG_padj", "DEG_tagwise.dispersion", "DEG_trended.dispersion", "DEG_strand" ,              
                                    "mm10_start_gene_annot_DEG","mm10_end_gene_annot_DEG", "entrezgene_id_annot_DEG", "gene_biotype_annot_DEG" , "mgi_symbol_annot_DEG" , 
                                    "entrezgene_accession_annot_DEG", "entrezgene_description_annot_DEG",
                                    "uniprot_gn_symbol_annot_DEG")                                   
  
  
  
  write.table(Annot_DMR_DEG_clean, file=paste("H:/methylome/fusion/post_methylkit/DMR432_pval0.07/croisement_autre_data/DMR_RNAseq/clean_annot",DMR_DEG_sample,sep="_"), quote=FALSE, sep="\t", row.names = FALSE, col.names = TRUE)
  write.table(Annot_DMR_DEG, file=paste("H:/methylome/fusion/post_methylkit/DMR432_pval0.07/croisement_autre_data/DMR_RNAseq/raw_annot",DMR_DEG_sample,sep="_"), quote=FALSE, sep="\t", row.names = FALSE, col.names = TRUE)
  
  # To keep only gene names (entrezgene_accession_annot_DMR) - annotation of DMR - mm10 BIOMART : 
  
  # To uncollapse data
  decondat_Annot_DMR_DEG_clean = as.character(Annot_DMR_DEG_clean$entrezgene_accession_annot_DMR)
  print(DMR_DEG_sample)
  print(head(decondat_Annot_DMR_DEG_clean)) 
  
  # To split gene names, when necessary
  list_gene_splitted <- strsplit(as.character(Annot_DMR_DEG_clean$entrezgene_accession_annot_DMR),split=";")
  vec_gene_splitted <- unlist(list_gene_splitted)
  
  # To remove "." : 
  vec_gene_splitted_filter = vec_gene_splitted[vec_gene_splitted!="."]
  
  # To save data in a table 
   table_VGS_filter = as.data.frame(unique(vec_gene_splitted_filter))
  
  write.table(table_VGS_filter, paste("H:/methylome/fusion/post_methylkit/DMR432_pval0.07/croisement_autre_data/DMR_RNAseq/list_gene",DMR_DEG_sample, sep="_"), quote=FALSE, col.names=FALSE, row.names=FALSE)
  
}
```

#### Integration of DEG with DOCR¶

To determine whether DEG are also DOCR during physiological brain development or not, DEG and DOCR regions were compared using our own R function called ***find\_overlaps\_AD\_table()***. Both overlaping regions and regions less than 1000 bases apart were investigated, by setting ***tolerance*** argument to 0 or 1000. The obtained regions were then annotated, using annotations that were already achieved in the two complete datasets.

In [ ]:

```
%%R

####################################################################
# Integration of DEG and DOCR to find potential overlap
# (mm10 chromosomal coordinates)
####################################################################

# 1. Function to find overlap or close regions (depending on value given for tolerance parameter)
find_overlaps_AD_table <- function (regions_1,regions_2,tolerance=0)
{
  common_region_table = NULL
  number_of_regions_1 <- nrow(regions_1)
  number_of_regions_2 <- nrow(regions_2)
  
  overlapped_is <- c()
  overlapped_js <- c()
  
  for (i in 1:number_of_regions_1) {
    
    for (j in 1:number_of_regions_2) {
      
      if (as.character(regions_1[i,1])==as.character(regions_2[j,1])) {
        start_1 <- as.numeric(regions_1[i,2])
        end_1 <- as.numeric(regions_1[i,3])
        expanded_start_1 <- start_1 - tolerance
        expanded_end_1 <- end_1 + tolerance
        start_2 <- as.numeric(regions_2[j,2])
        end_2 <- as.numeric(regions_2[j,3])
        
        if (expanded_end_1>=start_2) {
          
          if (end_2>=expanded_start_1) {
            
            overlapped_is <- c(overlapped_is,i)
            overlapped_js <- c(overlapped_js,j)
            
            first_region <- paste(regions_1[i,1],start_1,end_1,sep="_")
            second_region <- paste(regions_2[j,1],start_2,end_2,sep="_")
            overlap_size <- min(end_1,end_2)-max(start_1,start_2)+1
            common_region_table = rbind(common_region_table,c(as.character(regions_1[i,1]),start_1,end_1, 
                                                              as.character(regions_2[j,1]),start_2,end_2, overlap_size))
            
            if (overlap_size>0) {
              print(paste(first_region,second_region,sep=" "))
            } else {
              print(paste(first_region,second_region, sep=" "))
            }
          }
        } 
      }
    }
  }
  print(paste("Number matched in first data set:",length(unique(overlapped_is))))
  print(paste("Number matched in second data set:",length(unique(overlapped_js))))
  if (length(common_region_table) > 0){
    colnames(common_region_table) = c("chr_reg1", "start_reg1", "end_reg1", "chr_reg2", "start_reg2", "end_reg2","overlap_size")
  }else{
  }
  return(as.data.frame(common_region_table))
}


# 2. Load DOCR and DEG data - mm10 (from edgeR analysis):
nom_sample_ATAC = list("down_DOCR_E13.14", "down_DOCR_E14.15", "down_DOCR_E14.16", "down_DOCR_E15.16", "down_DOCR_E16.P0", 
                    "up_DOCR_E13.14", "up_DOCR_E14.15", "up_DOCR_E14.16", "up_DOCR_E15.16", "up_DOCR_E16.P0" )
nom_sample_RNA = list("down_E13.14", "down_E14.15", "down_E14.16", "down_E15.16", "down_E16.P0", "up_E13.14", "up_E14.15", 
                      "up_E14.16", "up_E15.16", "up_E16.P0" )

# /!\ /!\ Dataset must be in a matrix format, otherwise function doesn't work /!\/!\
for(i in 1:length(nom_sample_ATAC)){
  assign(x=nom_sample_ATAC[[i]], value = as.matrix(read.table(paste("G:/projet_DU_DSD/results/10_annotations/Sartools/Biomart/mm10/bed_gff_ID_", nom_sample_ATAC[[i]], "_mm10.sorted.bed",  
                                                    sep=""), sep = "\t", header= FALSE, dec=".", quote="")))
  assign(x=nom_sample_RNA[[i]], value = as.matrix(read.table(paste("G:/projet_DU_AD/results/09_annotation/Sartools/Biomart/bed_edge_AnnotBM_et_Man_ech_",  
                                                                   nom_sample_RNA[[i]] , "_mm10.sorted.bed", sep=""), sep = "\t", header= FALSE, dec=".", quote="")))
  
  
  print(paste("dimension du dataset ",nom_sample_ATAC[[i]], sep = ""))
  print(dim(eval(parse(text = nom_sample_ATAC[[i]]))))
  
  print(paste("dimension du dataset ",nom_sample_RNA[[i]], sep = ""))
  print(dim(eval(parse(text = nom_sample_RNA[[i]]))))
  
  # 3. Search for common regions between DEG and DOCR - overlap
  print("distance = 0")
  recoupement = find_overlaps_AD_table(get(nom_sample_RNA[[i]]), get(nom_sample_ATAC[[i]]), tolerance=0)
  write.table(recoupement,paste("H:/methylome/fusion/post_methylkit/DMR432_pval0.07/croisement_autre_data/mm10_RNA_ATAC", 
                                nom_sample_RNA[[i]], "dist0.txt",sep="_"), sep = "\t", quote=FALSE, row.names = FALSE, col.names=FALSE)
  
  # 4. Search for common regions between DEG and DOCR - maximum distance between 2 regions = 1000bases: 
  print("distance = 1000")
  recoupement_large = find_overlaps_AD_table(get(nom_sample_RNA[[i]]), get(nom_sample_ATAC[[i]]), tolerance=1000)
  write.table(recoupement_large,paste("H:/methylome/fusion/post_methylkit/DMR432_pval0.07/croisement_autre_data/mm10_RNA_ATAC", 
                                      nom_sample_RNA[[i]], "dist1000.txt",sep="_"), sep = "\t", quote=FALSE, row.names = FALSE, col.names=FALSE)
    
}


###################
# Annotation 
###################

# Load files containing DEG+DOCR regions
list_DEG_DOCR = list.files("H:/methylome/fusion/post_methylkit/DMR432_pval0.07/croisement_autre_data/RNA_ATAC/")
list_DEG_DOCR_dist0 = list_DEG_DOCR[which(grepl("dist0",list_DEG_DOCR))]
list_DEG_DOCR_dist1000 = list_DEG_DOCR[which(grepl("dist1000",list_DEG_DOCR))]

# List of files containing DOCR informations 
# Since some files were empty (no overlap), we need to specify, in a list, which file we want to annotate.
list_DOCR_Annot=list("stats_mm9_mm10_down_DOCR_E14.15_annot_syntaxique_mm10_biomart.bed","stats_mm9_mm10_down_DOCR_E14.16_annot_syntaxique_mm10_biomart.bed",
                     "stats_mm9_mm10_down_DOCR_E15.16_annot_syntaxique_mm10_biomart.bed",
                     "stats_mm9_mm10_down_DOCR_E16.P0_annot_syntaxique_mm10_biomart.bed", "stats_mm9_mm10_up_DOCR_E14.15_annot_syntaxique_mm10_biomart.bed",
                     "stats_mm9_mm10_up_DOCR_E14.16_annot_syntaxique_mm10_biomart.bed", "stats_mm9_mm10_up_DOCR_E15.16_annot_syntaxique_mm10_biomart.bed",
                     "stats_mm9_mm10_up_DOCR_E16.P0_annot_syntaxique_mm10_biomart.bed")


# Load DEG annotation files
# Since some files were empty (no overlap), we need to specify, in a list, which file we want to annotate.
list_DEG_Annot = list("edgeR_AnnotBM_et_Man_down_E14.15_mm10.txt", "edgeR_AnnotBM_et_Man_down_E14.16_mm10.txt" ,
                      "edgeR_AnnotBM_et_Man_down_E15.16_mm10.txt", 
                      "edgeR_AnnotBM_et_Man_down_E16.P0_mm10.txt", "edgeR_AnnotBM_et_Man_up_E14.15_mm10.txt",
                      "edgeR_AnnotBM_et_Man_up_E14.16_mm10.txt", "edgeR_AnnotBM_et_Man_up_E15.16_mm10.txt",
                      "edgeR_AnnotBM_et_Man_up_E16.P0_mm10.txt")

# for distance=0
for(i in 1:8){
  
  DEG_DOCR_sample = list_DEG_DOCR_dist0[i]
  Annot_DEG_DOCR = read.table(file = paste("H:/methylome/fusion/post_methylkit/DMR432_pval0.07/croisement_autre_data/RNA_ATAC/",DEG_DOCR_sample,sep="/"), 
                              header = FALSE, sep = "\t", quote="", fill=TRUE)
  
  # To rename columns
  colnames(Annot_DEG_DOCR) <- c("DEG_chr","DEG_start", "DEG_end", "DOCR_chr", "DOCR_start", "DOCR_end", "DEG_DOCR_overlap")
  
  # To create two ID : DEG_ID et DOCR_ID
  Annot_DEG_DOCR$DEG_ID = paste(Annot_DEG_DOCR$DEG_chr, Annot_DEG_DOCR$DEG_start, Annot_DEG_DOCR$DEG_end, sep=";")
  Annot_DEG_DOCR$DOCR_ID = paste(Annot_DEG_DOCR$DOCR_chr, Annot_DEG_DOCR$DOCR_start, Annot_DEG_DOCR$DOCR_end, sep=";")
  
  # Load annotation of all DEG regions
  DEG_sample <- list_DEG_Annot[i]
  Annot_DEG = read.table(file = paste("G:/projet_DU_AD/results/09_annotation/Sartools/Biomart",DEG_sample,sep="/"), header = TRUE, sep = "\t", quote="", fill=TRUE)
  
  colnames(Annot_DEG)= c("ENS_ID", "cond1_rep1", "cond1_rep2", "cond2_rep1", "cond2_rep2", 
                         "norm.cond1_rep1", "norm.cond1_rep2", "norm.cond2_rep1", "norm.cond2_rep2", 
                         "baseMean", "cond1", "cond2", "FC", "log2FoldChange", "pvalue", "padj", "tagwise.dispersion", 
                         "trended.dispersion", "mm10_chr", "strand", "mm10_start_gene", "mm10_end_gene", "entrezgene_id", "gene_biotype", 
                         "mgi_symbol", "entrezgene_accession", "entrezgene_description", "uniprot_gn_symbol")
  
  Annot_DEG$DEG_ID=paste(paste("chr",Annot_DEG$mm10_chr,sep=""), Annot_DEG$mm10_start_gene, Annot_DEG$mm10_end_gene,sep=";")
  
  
  # Load annoation of all DOCR
  Annot_DOCR_sample = list_DOCR_Annot[i]
  Annot_DOCR = read.table(file = paste("G:/projet_DU_DSD/results/10_annotations/Sartools/Biomart/mm10",Annot_DOCR_sample,sep="/"), header = TRUE, sep = "\t", quote="", fill=TRUE)
  
  colnames(Annot_DOCR) = c("mm9_chr", "mm9_start", "mm9_end", "mm9_ID", "mm10_chr", "mm10_start", "mm10_end", "mm10_ID", 
                           "chr_gene", "start_gene", "end_gene", "strand", "entrezgene_accession", "gene_biotype", "mgi_symbol", 
                           "ensembl_gene_id", "entrezgene_id", "overlap_length", "cond1_rep1", "cond1_rep2", "cond2_rep1", "cond2_rep2", 
                           "norm.cond1_rep1", "norm.cond1_rep2", "norm.cond2_rep1", "norm.cond2_rep2", "baseMean", "cond1", "cond2", "FC", 
                           "log2FoldChange", "pvalue", "padj", "tagwise.dispersion", "trended.dispersion")
  
  
  Annot_DOCR$DOCR_ID=paste(Annot_DOCR$mm10_chr, Annot_DOCR$mm10_start, Annot_DOCR$mm10_end,sep=";")
  
  Annot_DEG_DOCR=merge(Annot_DEG_DOCR, Annot_DEG, by="DEG_ID")
  Annot_DEG_DOCR=merge(Annot_DEG_DOCR, Annot_DOCR, by="DOCR_ID")
  
  # To remove duplicate columns 
  Annot_DEG_DOCR_clean = Annot_DEG_DOCR[,c("DEG_chr", "DEG_start", "DEG_end","DEG_ID" , "ENS_ID" , 
                                           "cond1_rep1.x", "cond1_rep2.x", "cond2_rep1.x" , "cond2_rep2.x",
                                           "norm.cond1_rep1.x", "norm.cond1_rep2.x",  "norm.cond2_rep1.x" , "norm.cond2_rep2.x","baseMean.x", "cond1.x","cond2.x","FC.x",
                                           "log2FoldChange.x", "pvalue.x",  "padj.x", "tagwise.dispersion.x", "trended.dispersion.x", "strand.x",
                                           "mm10_chr.x","mm10_start_gene","mm10_end_gene", "entrezgene_id.x", "gene_biotype.x" , "mgi_symbol.x" , "entrezgene_accession.x", "entrezgene_description",
                                           "uniprot_gn_symbol",
                                           "DEG_DOCR_overlap", "DOCR_chr", "DOCR_start", "DOCR_end","DOCR_ID" ,
                                           "mm9_chr", "mm9_start", "mm9_end", "mm9_ID","cond1_rep1.y", "cond1_rep2.y", "cond2_rep1.y" , "cond2_rep2.y", 
                                           "norm.cond1_rep1.y", "norm.cond1_rep2.y",  "norm.cond2_rep1.y" , "norm.cond2_rep2.y",  "baseMean.y",  "cond1.y" , "cond2.y" ,  "FC.y",
                                           "log2FoldChange.y", "pvalue.y",  "padj.y", "tagwise.dispersion.y", "trended.dispersion.y",
                                           "strand.y", "chr_gene", "start_gene", "end_gene", "entrezgene_accession.y", "gene_biotype.y", 
                                           "mgi_symbol.y" , "ensembl_gene_id", "entrezgene_id.y", "overlap_length")]
  
  
  colnames(Annot_DEG_DOCR_clean)=c("DEG_mm10_chr", "DEG_mm10_start", "DEG_mm10_end","DEG_mm10_ID" ,"DEG_ENS_ID" , 
                                   "DEG_cond1_rep1", "DEG_cond1_rep2", "DEG_cond2_rep1" , "DEG_cond2_rep2",
                                   "DEG_norm.cond1_rep1", "DEG_norm.cond1_rep2",  "DEG_norm.cond2_rep1" , "DEG_norm.cond2_rep2", 
                                   "DEG_baseMean",  "DEG_cond1" , "DEG_cond2" , "DEG_FC",
                                   "DEG_log2FoldChange", "DEG_pvalue",  "DEG_padj", "DEG_tagwise.dispersion", "DEG_trended.dispersion", "DEG_strand", "mm10_chr_gene_annot_DEG",             
                                   "mm10_start_gene_annot_DEG","mm10_end_gene_annot_DEG", "entrezgene_id_annot_DEG", "gene_biotype_annot_DEG" , "mgi_symbol_annot_DEG" , 
                                   "entrezgene_accession_annot_DEG", "entrezgene_description_annot_DEG","uniprot_gn_symbol_annot_DEG",
                                   "DEG_DOCR_overlap",
                                   "DOCR_mm10_chr", "DOCR_mm10_start", "DOCR_mm10_end","DOCR_mm10_ID",
                                   "DOCR_mm9_chr", "DOCR_mm9_start", "DOCR_mm9_end", "DOCR_mm9_ID","DOCR_cond1_rep1", "DOCR_cond1_rep2", 
                                   "DOCR_cond2_rep1" , "DOCR_cond2_rep2",
                                   "DOCR_norm.cond1_rep1", "DOCR_norm.cond1_rep2",  "DOCR_norm.cond2_rep1" , "DOCR_norm.cond2_rep2",  "DOCR_baseMean",  "DOCR_cond1" , "DOCR_cond2" ,
                                   "DOCR_FC",
                                   "DOCR_log2FoldChange", "DOCR_pvalue",  "DOCR_padj", "DOCR_tagwise.dispersion", "DOCR_trended.dispersion", 
                                   "strand_annot_DOCR", "mm10_chr_gene_annot_DOCR", "mm10_start_gene_annot_DOCR","mm10_end_gene_annot_DOCR", 
                                   "entrezgene_accession_annot_DOCR", "gene_biotype_annot_DOCR", 
                                   "mgi_symbol_annot_DOCR" , "ensembl_gene_id_annot_DOCR", "entrezgene_id_annot_DOCR", "overlap_length_annot_DOCR")
  
  
  
  write.table(Annot_DEG_DOCR_clean, file=paste("H:/methylome/fusion/post_methylkit/DMR432_pval0.07/croisement_autre_data/RNA_ATAC/clean_annot",DEG_DOCR_sample,sep="_"), quote=FALSE, sep="\t", row.names = FALSE, col.names = TRUE)
  write.table(Annot_DEG_DOCR, file=paste("H:/methylome/fusion/post_methylkit/DMR432_pval0.07/croisement_autre_data/RNA_ATAC/raw_annot",DEG_DOCR_sample,sep="_"), quote=FALSE, sep="\t", row.names = FALSE, col.names = TRUE)
  
  
  # To keep only gene names (entrezgene_accession_annot_DEG) - annotation of DEG - mm10 BIOMART : 
  # To uncollapse data
  decondat_Annot_DEG_DOCR_clean = as.character(Annot_DEG_DOCR_clean$entrezgene_accession_annot_DEG)
  print(DEG_DOCR_sample)
  print(head(decondat_Annot_DEG_DOCR_clean)) 
  
  # To obtain list of genes 
  list_gene_splitted <- strsplit(as.character(Annot_DEG_DOCR_clean$entrezgene_accession_annot_DEG),split=";")
  vec_gene_splitted <- unlist(list_gene_splitted)
  
  # To remove "." : 
  vec_gene_splitted_filter = vec_gene_splitted[vec_gene_splitted!="."]
  
  # To save data on a table
  table_VGS_filter = as.data.frame(unique(vec_gene_splitted_filter))
  
  write.table(table_VGS_filter, paste("H:/methylome/fusion/post_methylkit/DMR432_pval0.07/croisement_autre_data/RNA_ATAC/list_gene",DEG_DOCR_sample, sep="_"), quote=FALSE, col.names=FALSE, row.names=FALSE)
  
}


rm(list = ls())

#-----------------------------------
# Idem for distance=1000
# Load files containing DEG+DOCR regions
list_DEG_DOCR = list.files("H:/methylome/fusion/post_methylkit/DMR432_pval0.07/croisement_autre_data/RNA_ATAC/")
list_DEG_DOCR_dist0 = list_DEG_DOCR[which(grepl("dist0",list_DEG_DOCR))]
list_DEG_DOCR_dist1000 = list_DEG_DOCR[which(grepl("dist1000",list_DEG_DOCR))]

# List of files containing DOCR informations
# Since some files were empty (no overlap), we need to specify, in a list, which file we want to annotate.
list_DOCR_Annot=list("stats_mm9_mm10_down_DOCR_E14.15_annot_syntaxique_mm10_biomart.bed","stats_mm9_mm10_down_DOCR_E14.16_annot_syntaxique_mm10_biomart.bed",
                     "stats_mm9_mm10_down_DOCR_E15.16_annot_syntaxique_mm10_biomart.bed",
                     "stats_mm9_mm10_down_DOCR_E16.P0_annot_syntaxique_mm10_biomart.bed", "stats_mm9_mm10_up_DOCR_E14.15_annot_syntaxique_mm10_biomart.bed",
                     "stats_mm9_mm10_up_DOCR_E14.16_annot_syntaxique_mm10_biomart.bed", "stats_mm9_mm10_up_DOCR_E15.16_annot_syntaxique_mm10_biomart.bed",
                     "stats_mm9_mm10_up_DOCR_E16.P0_annot_syntaxique_mm10_biomart.bed")


# Load DEG annotation files
# Since some files were empty (no overlap), we need to specify, in a list, which file we want to annotate.
list_DEG_Annot = list("edgeR_AnnotBM_et_Man_down_E14.15_mm10.txt", "edgeR_AnnotBM_et_Man_down_E14.16_mm10.txt" ,
                      "edgeR_AnnotBM_et_Man_down_E15.16_mm10.txt", 
                      "edgeR_AnnotBM_et_Man_down_E16.P0_mm10.txt", "edgeR_AnnotBM_et_Man_up_E14.15_mm10.txt",
                      "edgeR_AnnotBM_et_Man_up_E14.16_mm10.txt", "edgeR_AnnotBM_et_Man_up_E15.16_mm10.txt",
                      "edgeR_AnnotBM_et_Man_up_E16.P0_mm10.txt")


# for distance=1000
for(i in 1:8){
  
  DEG_DOCR_sample = list_DEG_DOCR_dist1000[i]
  Annot_DEG_DOCR = read.table(file = paste("H:/methylome/fusion/post_methylkit/DMR432_pval0.07/croisement_autre_data/RNA_ATAC/",DEG_DOCR_sample,sep="/"), 
                              header = FALSE, sep = "\t", quote="", fill=TRUE)
  
  # To rename columns
  colnames(Annot_DEG_DOCR) <- c("DEG_chr","DEG_start", "DEG_end", "DOCR_chr", "DOCR_start", "DOCR_end", "DEG_DOCR_overlap")
  
  #To create two ID : DEG_ID et DOCR_ID
  Annot_DEG_DOCR$DEG_ID = paste(Annot_DEG_DOCR$DEG_chr, Annot_DEG_DOCR$DEG_start, Annot_DEG_DOCR$DEG_end, sep=";")
  Annot_DEG_DOCR$DOCR_ID = paste(Annot_DEG_DOCR$DOCR_chr, Annot_DEG_DOCR$DOCR_start, Annot_DEG_DOCR$DOCR_end, sep=";")
  
  # Load annotation of all DEG regions
  DEG_sample <- list_DEG_Annot[i]
  Annot_DEG = read.table(file = paste("G:/projet_DU_AD/results/09_annotation/Sartools/Biomart",DEG_sample,sep="/"), header = TRUE, sep = "\t", quote="", fill=TRUE)
  
  colnames(Annot_DEG)= c("ENS_ID", "cond1_rep1", "cond1_rep2", "cond2_rep1", "cond2_rep2", 
                         "norm.cond1_rep1", "norm.cond1_rep2", "norm.cond2_rep1", "norm.cond2_rep2", 
                         "baseMean", "cond1", "cond2", "FC", "log2FoldChange", "pvalue", "padj", "tagwise.dispersion", 
                         "trended.dispersion", "mm10_chr", "strand", "mm10_start_gene", "mm10_end_gene", "entrezgene_id", "gene_biotype", 
                         "mgi_symbol", "entrezgene_accession", "entrezgene_description", "uniprot_gn_symbol")
  
  Annot_DEG$DEG_ID=paste(paste("chr",Annot_DEG$mm10_chr,sep=""), Annot_DEG$mm10_start_gene, Annot_DEG$mm10_end_gene,sep=";")
  
    # Load annoation of all DOCR
  Annot_DOCR_sample = list_DOCR_Annot[i]
  Annot_DOCR = read.table(file = paste("G:/projet_DU_DSD/results/10_annotations/Sartools/Biomart/mm10",Annot_DOCR_sample,sep="/"), header = TRUE, sep = "\t", quote="", fill=TRUE)
  
  colnames(Annot_DOCR) = c("mm9_chr", "mm9_start", "mm9_end", "mm9_ID", "mm10_chr", "mm10_start", "mm10_end", "mm10_ID", 
                           "chr_gene", "start_gene", "end_gene", "strand", "entrezgene_accession", "gene_biotype", "mgi_symbol", 
                           "ensembl_gene_id", "entrezgene_id", "overlap_length", "cond1_rep1", "cond1_rep2", "cond2_rep1", "cond2_rep2", 
                           "norm.cond1_rep1", "norm.cond1_rep2", "norm.cond2_rep1", "norm.cond2_rep2", "baseMean", "cond1", "cond2", "FC", 
                           "log2FoldChange", "pvalue", "padj", "tagwise.dispersion", "trended.dispersion")
  
  
  Annot_DOCR$DOCR_ID=paste(Annot_DOCR$mm10_chr, Annot_DOCR$mm10_start, Annot_DOCR$mm10_end,sep=";")
  
  Annot_DEG_DOCR=merge(Annot_DEG_DOCR, Annot_DEG, by="DEG_ID")
  Annot_DEG_DOCR=merge(Annot_DEG_DOCR, Annot_DOCR, by="DOCR_ID")
  
   
  # To remove duplicate columns 
  Annot_DEG_DOCR_clean = Annot_DEG_DOCR[,c("DEG_chr", "DEG_start", "DEG_end","DEG_ID" , "ENS_ID" , 
                                           "cond1_rep1.x", "cond1_rep2.x", "cond2_rep1.x" , "cond2_rep2.x",
                                           "norm.cond1_rep1.x", "norm.cond1_rep2.x",  "norm.cond2_rep1.x" , "norm.cond2_rep2.x","baseMean.x", "cond1.x","cond2.x","FC.x",
                                           "log2FoldChange.x", "pvalue.x",  "padj.x", "tagwise.dispersion.x", "trended.dispersion.x", "strand.x",
                                           "mm10_chr.x","mm10_start_gene","mm10_end_gene", "entrezgene_id.x", "gene_biotype.x" , "mgi_symbol.x" , "entrezgene_accession.x", "entrezgene_description",
                                           "uniprot_gn_symbol",
                                           "DEG_DOCR_overlap", "DOCR_chr", "DOCR_start", "DOCR_end","DOCR_ID" ,
                                           "mm9_chr", "mm9_start", "mm9_end", "mm9_ID","cond1_rep1.y", "cond1_rep2.y", "cond2_rep1.y" , "cond2_rep2.y", 
                                           "norm.cond1_rep1.y", "norm.cond1_rep2.y",  "norm.cond2_rep1.y" , "norm.cond2_rep2.y",  "baseMean.y",  "cond1.y" , "cond2.y" ,  "FC.y",
                                           "log2FoldChange.y", "pvalue.y",  "padj.y", "tagwise.dispersion.y", "trended.dispersion.y",
                                           "strand.y", "chr_gene", "start_gene", "end_gene", "entrezgene_accession.y", "gene_biotype.y", 
                                           "mgi_symbol.y" , "ensembl_gene_id", "entrezgene_id.y", "overlap_length")]
  
  
  colnames(Annot_DEG_DOCR_clean)=c("DEG_mm10_chr", "DEG_mm10_start", "DEG_mm10_end","DEG_mm10_ID" ,"DEG_ENS_ID" , 
                                   "DEG_cond1_rep1", "DEG_cond1_rep2", "DEG_cond2_rep1" , "DEG_cond2_rep2",
                                   "DEG_norm.cond1_rep1", "DEG_norm.cond1_rep2",  "DEG_norm.cond2_rep1" , "DEG_norm.cond2_rep2", 
                                   "DEG_baseMean",  "DEG_cond1" , "DEG_cond2" , "DEG_FC",
                                   "DEG_log2FoldChange", "DEG_pvalue",  "DEG_padj", "DEG_tagwise.dispersion", "DEG_trended.dispersion", "DEG_strand", "mm10_chr_gene_annot_DEG",             
                                   "mm10_start_gene_annot_DEG","mm10_end_gene_annot_DEG", "entrezgene_id_annot_DEG", "gene_biotype_annot_DEG" , "mgi_symbol_annot_DEG" , 
                                   "entrezgene_accession_annot_DEG", "entrezgene_description_annot_DEG","uniprot_gn_symbol_annot_DEG",
                                   "DEG_DOCR_overlap",
                                   "DOCR_mm10_chr", "DOCR_mm10_start", "DOCR_mm10_end","DOCR_mm10_ID",
                                   "DOCR_mm9_chr", "DOCR_mm9_start", "DOCR_mm9_end", "DOCR_mm9_ID","DOCR_cond1_rep1", "DOCR_cond1_rep2", 
                                   "DOCR_cond2_rep1" , "DOCR_cond2_rep2",
                                   "DOCR_norm.cond1_rep1", "DOCR_norm.cond1_rep2",  "DOCR_norm.cond2_rep1" , "DOCR_norm.cond2_rep2",  "DOCR_baseMean",  "DOCR_cond1" , "DOCR_cond2" ,
                                   "DOCR_FC",
                                   "DOCR_log2FoldChange", "DOCR_pvalue",  "DOCR_padj", "DOCR_tagwise.dispersion", "DOCR_trended.dispersion", 
                                   "strand_annot_DOCR", "mm10_chr_gene_annot_DOCR", "mm10_start_gene_annot_DOCR","mm10_end_gene_annot_DOCR", 
                                   "entrezgene_accession_annot_DOCR", "gene_biotype_annot_DOCR", 
                                   "mgi_symbol_annot_DOCR" , "ensembl_gene_id_annot_DOCR", "entrezgene_id_annot_DOCR", "overlap_length_annot_DOCR")
  
  
  
  write.table(Annot_DEG_DOCR_clean, file=paste("H:/methylome/fusion/post_methylkit/DMR432_pval0.07/croisement_autre_data/RNA_ATAC/clean_annot",DEG_DOCR_sample,sep="_"), quote=FALSE, sep="\t", row.names = FALSE, col.names = TRUE)
  write.table(Annot_DEG_DOCR, file=paste("H:/methylome/fusion/post_methylkit/DMR432_pval0.07/croisement_autre_data/RNA_ATAC/raw_annot",DEG_DOCR_sample,sep="_"), quote=FALSE, sep="\t", row.names = FALSE, col.names = TRUE)
  
  
  # To keep only gene names (entrezgene_accession_annot_DEG) - annotation of DEG - mm10 BIOMART : 
  # To uncollapse data
  decondat_Annot_DEG_DOCR_clean = as.character(Annot_DEG_DOCR_clean$entrezgene_accession_annot_DEG)
  print(DEG_DOCR_sample)
  print(head(decondat_Annot_DEG_DOCR_clean)) 
  
  # To obtain list of genes 
  list_gene_splitted <- strsplit(as.character(Annot_DEG_DOCR_clean$entrezgene_accession_annot_DEG),split=";")
  vec_gene_splitted <- unlist(list_gene_splitted)
  
  # To remove "." 
  vec_gene_splitted_filter = vec_gene_splitted[vec_gene_splitted!="."]
  
  # To save data on a table
  table_VGS_filter = as.data.frame(unique(vec_gene_splitted_filter))
  
  write.table(table_VGS_filter, paste("H:/methylome/fusion/post_methylkit/DMR432_pval0.07/croisement_autre_data/RNA_ATAC/list_gene",DEG_DOCR_sample, sep="_"), quote=FALSE, col.names=FALSE, row.names=FALSE)
  
}
```

#### Estimation of the proportion of DOCR included in methylome capture¶

To observe overlap between DOCR and DMR, sequences of DOCR must be included in methylome capture, otherwise, it will not be possible to detect a DMR, even if it exists. To determine proportion of DOCR regions that are in methylome capture, methylome capture regions and DOCR ones were compared using our own R function called ***find\_overlaps\_AD\_table()***. Both overlaping regions and regions les than 1000 bases apart were investigated by setting ***tolerance*** argument to 0 or 1000. For these comparisons, mm9 coordinates were used.

In [ ]:

```
%%R

# Load DOCR files
setwd("E:/projet_DU_DSD/results/10_annotations/Sartools/Biomart/mm10")

DOCR_u14_15 = read.table("stats_mm9_mm10_up_DOCR_E14.15_annot_syntaxique_mm10_biomart.bed", header = TRUE, fill=TRUE, quote="", sep="\t", na.strings = "", stringsAsFactors = FALSE)
DOCR_u15_16 = read.table("stats_mm9_mm10_up_DOCR_E15.16_annot_syntaxique_mm10_biomart.bed", header = TRUE, fill=TRUE, quote="", sep="\t", na.strings = "", stringsAsFactors = FALSE)
DOCR_d14_15 = read.table("stats_mm9_mm10_down_DOCR_E14.15_annot_syntaxique_mm10_biomart.bed", header = TRUE, fill=TRUE, quote="", sep="\t", na.strings = "", stringsAsFactors = FALSE)
DOCR_d15_16 = read.table("stats_mm9_mm10_down_DOCR_E15.16_annot_syntaxique_mm10_biomart.bed", header = TRUE, fill=TRUE, quote="", sep="\t", na.strings = "", stringsAsFactors = FALSE)

head(DOCR_u14_15)
head(DOCR_u15_16)
head(DOCR_d14_15)
head(DOCR_d15_16)

dim(DOCR_u14_15)
dim(DOCR_u15_16)
dim(DOCR_d14_15)
dim(DOCR_d15_16)

# Load capture files containing regions included in the methylome capture 
setwd("H:/methylome/fusion/capture/capture_regions/")

capture = read.table("Capture_76800_et150seq.merge.sorted.bed", header = FALSE, fill=TRUE, quote="", sep="\t", na.strings = "", stringsAsFactors = FALSE)

dim(capture)
head(capture)

# Function to find overlap (or close) regions between two datasets
find_overlaps_AD_table <- function (regions_1,regions_2,tolerance=0)
{
  common_region_table = NULL
  number_of_regions_1 <- nrow(regions_1)
  number_of_regions_2 <- nrow(regions_2)
  
  overlapped_is <- c()
  overlapped_js <- c()
  
  for (i in 1:number_of_regions_1) {
    
    for (j in 1:number_of_regions_2) {
      
      if (as.character(regions_1[i,1])==as.character(regions_2[j,1])) {
        start_1 <- as.numeric(regions_1[i,2])
        end_1 <- as.numeric(regions_1[i,3])
        expanded_start_1 <- start_1 - tolerance
        expanded_end_1 <- end_1 + tolerance
        start_2 <- as.numeric(regions_2[j,2])
        end_2 <- as.numeric(regions_2[j,3])
        
        if (expanded_end_1>=start_2) {
          
          if (end_2>=expanded_start_1) {
            
            overlapped_is <- c(overlapped_is,i)
            overlapped_js <- c(overlapped_js,j)
            
            first_region <- paste(regions_1[i,1],start_1,end_1,sep="_")
            second_region <- paste(regions_2[j,1],start_2,end_2,sep="_")
            overlap_size <- min(end_1,end_2)-max(start_1,start_2)+1
            common_region_table = rbind(common_region_table,c(as.character(regions_1[i,1]),start_1,end_1, 
                                                              as.character(regions_2[j,1]),start_2,end_2, overlap_size))
            
            if (overlap_size>0) {
              print(paste(first_region,second_region,sep=" "))
            } else {
              print(paste(first_region,second_region, sep=" "))
            }
          }
        } 
      }
    }
  }
  print(paste("Number matched in first data set:",length(unique(overlapped_is))))
  print(paste("Number matched in second data set:",length(unique(overlapped_js))))
  if (length(common_region_table) > 0){
    colnames(common_region_table) = c("chr_reg1", "start_reg1", "end_reg1", "chr_reg2", "start_reg2", "end_reg2","overlap_size")
  }else{
  }
  return(as.data.frame(common_region_table))
}


# Comparison of DOCR files and capture - dist0
capt_DOCR_u14_15 = find_overlaps_AD_table(DOCR_u14_15,capture,tolerance=0)
dim(capt_DOCR_u14_15)
write.table(capt_DOCR_u14_15,
            "H:/methylome/fusion/post_methylkit/DMR432_pval0.07/croisement_autre_data/ATACseq_capture/capt_DOCR_up_14_15_dist0.txt", 
            sep = "\t", quote=FALSE, row.names = FALSE, col.names=FALSE)

            
capt_DOCR_u15_16 = find_overlaps_AD_table(DOCR_u15_16,capture,tolerance=0)
dim(capt_DOCR_u15_16)
write.table(capt_DOCR_u15_16,
            "H:/methylome/fusion/post_methylkit/DMR432_pval0.07/croisement_autre_data/ATACseq_capture/capt_DOCR_up_15_16_dist0.txt", 
            sep = "\t", quote=FALSE, row.names = FALSE, col.names=FALSE)

            
capt_DOCR_d14_15 = find_overlaps_AD_table(DOCR_d14_15,capture,tolerance=0)
dim(capt_DOCR_d14_15)
write.table(capt_DOCR_d14_15,
            "H:/methylome/fusion/post_methylkit/DMR432_pval0.07/croisement_autre_data/ATACseq_capture/capt_DOCR_down_14_15_dist0.txt", 
            sep = "\t", quote=FALSE, row.names = FALSE, col.names=FALSE)

            
capt_DOCR_d15_16 = find_overlaps_AD_table(DOCR_d15_16,capture,tolerance=0)
dim(capt_DOCR_d15_16)
write.table(capt_DOCR_d15_16,
            "H:/methylome/fusion/post_methylkit/DMR432_pval0.07/croisement_autre_data/ATACseq_capture/capt_DOCR_down_15_16_dist0.txt", 
            sep = "\t", quote=FALSE, row.names = FALSE, col.names=FALSE)

    


# Comparison of DOCR files and capture - dist1000
large_capt_DOCR_u14_15 = find_overlaps_AD_table(DOCR_u14_15,capture,tolerance=1000)
dim(large_capt_DOCR_u14_15)
write.table(large_capt_DOCR_u14_15,
            "H:/methylome/fusion/post_methylkit/DMR432_pval0.07/croisement_autre_data/ATACseq_capture/large_capt_DOCR_up_14_15_dist1000.txt", 
            sep = "\t", quote=FALSE, row.names = FALSE, col.names=FALSE)


large_capt_DOCR_u15_16 = find_overlaps_AD_table(DOCR_u15_16,capture,tolerance=1000)
dim(large_capt_DOCR_u15_16)
write.table(large_capt_DOCR_u15_16,
            "H:/methylome/fusion/post_methylkit/DMR432_pval0.07/croisement_autre_data/ATACseq_capture/large_capt_DOCR_up_15_16_dist1000.txt", 
            sep = "\t", quote=FALSE, row.names = FALSE, col.names=FALSE)


large_capt_DOCR_d14_15 = find_overlaps_AD_table(DOCR_d14_15,capture,tolerance=1000)
dim(large_capt_DOCR_d14_15)
write.table(large_capt_DOCR_d14_15,
            "H:/methylome/fusion/post_methylkit/DMR432_pval0.07/croisement_autre_data/ATACseq_capture/large_capt_DOCR_down_14_15_dist1000.txt", 
            sep = "\t", quote=FALSE, row.names = FALSE, col.names=FALSE)


large_capt_DOCR_d15_16 = find_overlaps_AD_table(DOCR_d15_16,capture,tolerance=1000)
dim(large_capt_DOCR_d15_16)
write.table(large_capt_DOCR_d15_16,
            "H:/methylome/fusion/post_methylkit/DMR432_pval0.07/croisement_autre_data/ATACseq_capture/large_capt_DOCR_down_15_16_dist1000.txt", 
            sep = "\t", quote=FALSE, row.names = FALSE, col.names=FALSE)
```

#### Estimation of the proportion of DEG included in methylome capture¶

To observe overlap between DEG and DMR, sequences of DEG must be in methylome capture, otherwise, it will not be possible to detect a DMR, even if it exists. To determine proportion of DEG regions that are included in methylome capture, methylome capture regions and DEG were compared using our own R function called ***find\_overlaps\_AD\_table()***. Both overlaping regions and regions less than 1000 bases apart were investigated by setting ***tolerance*** argument to 0 or 1000. To be able to compare the two datasets, DEG regions with mm10 coordinates were converted to mm9 coordinates using **LiftOver**, before comparisons (capture regions have mm9 coordinates). Default settings of LiftOver tool have been retained (*i.e.* at least 0.95 as the minimum ratio of bases that must remap). Some regions can't be converted for distinct reasons (see below).

**Explanations of conversion problem:**

- Deleted in new: Sequence intersects no chains
- Partially deleted in new: Sequence insufficiently intersects one chain
- Split in new: Sequence insufficiently intersects multiple chains
- Duplicated in new: Sequence sufficiently intersects multiple chains
- Boundary problem: Missing start or end base in an exon

In [ ]:

```
%%R 

#-----------------------------------------------------------------
# Conversion of mm9 coordinates --> mm10 coordinates with LiftOver

# Files that were converted:
# E:\projet_DU_AD\results\09_annotation\Sartools\Biomart\bed\bed_edge_AnnotBM_et_Man_ech_down_E14.15_mm10.sorted.bed 
# + bed_edge_AnnotBM_et_Man_ech_down_E15.16_mm10.sorted.bed
# + bed_edge_AnnotBM_et_Man_ech_up_E14.15_mm10.sorted.bed
# + bed_edge_AnnotBM_et_Man_ech_up_E15.16_mm10.sorted.bed

#---
# For DEG_E14.15_down : 
Successfully converted 2500 records
Conversion failed on 12 records. 

#Partially deleted in new
chr15	8967949	9067335
#Partially deleted in new
chr4	99829198	99912788
#Partially deleted in new
chr6	30747554	30896794
#Partially deleted in new
chr8	39005845	39165114
#Split in new
chr9	101074101	101104800
#Deleted in new
chrMT	70	1024
#Deleted in new
chrMT	2751	3707
#Deleted in new
chrMT	3845	3913
#Deleted in new
chrMT	10167	11544
#Deleted in new
chrMT	11742	13565
#Deleted in new
chrMT	14145	15288
#Partially deleted in new
chrX	140956907	141164270


#---
# For DEG_down_E15.16
Successfully converted 324 records
Conversion failed on 3 records. 

#Split in new
chr5	142724661	142817662
#Deleted in new
chrMT	3772	3842
#Deleted in new
chrMT	3845	3913


#---
#pour DEG_up_E14.15 : 
Successfully converted 3414 records
Conversion failed on 12 records.

#Partially deleted in new
chr1	22286251	22805994
#Split in new
chr12	18648214	18649998
#Partially deleted in new
chr2	58821070	59160683
#Split in new
chr4	156235919	156247616
#Partially deleted in new
chr5	113490333	113589725
#Split in new
chr5	142724661	142817662
#Partially deleted in new
chr7	6343759	6355956
#Partially deleted in new
chr7	138940730	139083977
#Split in new
chr9	124422622	124476898
#Deleted in new
chrMT	3772	3842
#Deleted in new
chrX	170009659	170019281
#Split in new
chrY	90784738	90816465


#---
#Pour DEG_up_E15.16
Successfully converted 196 records


# Data were saved on following files: 
#E:\projet_DU_AD\results\09_annotation\Sartools\Biomart\bed\mm9\mm9_bed_DEG_E14.15_down.bed
# + mm9_bed_DEG_E15.16_down.bed
# + mm9_bed_DEG_E14.15_up.bed
# + mm9_bed_DEG_E15.16_up.bed


#-----------------------------------------------------------------
# Comparison of DEG and capture regions
# Load DEG (mm9) files
setwd("E:/projet_DU_AD/results/09_annotation/Sartools/Biomart/bed/mm9/")

DEG_u14_15 = read.table("mm9_bed_DEG_E14.15_up.bed", header = FALSE, fill=TRUE, quote="", sep="\t", na.strings = "", stringsAsFactors = FALSE)
DEG_u15_16 = read.table("mm9_bed_DEG_E15.16_up.bed", header = FALSE, fill=TRUE, quote="", sep="\t", na.strings = "", stringsAsFactors = FALSE)
DEG_d14_15 = read.table("mm9_bed_DEG_E14.15_down.bed", header = FALSE, fill=TRUE, quote="", sep="\t", na.strings = "", stringsAsFactors = FALSE)
DEG_d15_16 = read.table("mm9_bed_DEG_E15.16_down.bed", header = FALSE, fill=TRUE, quote="", sep="\t", na.strings = "", stringsAsFactors = FALSE)

head(DEG_u14_15)
head(DEG_u15_16)
head(DEG_d14_15)
head(DEG_d15_16)

dim(DEG_u14_15)
dim(DEG_u15_16)
dim(DEG_d14_15)
dim(DEG_d15_16)


# Load capture file - mm9
setwd("H:/methylome/fusion/capture/capture_regions/")

capture = read.table("Capture_76800_et150seq.merge.sorted.bed", header = FALSE, fill=TRUE, quote="", sep="\t", na.strings = "", stringsAsFactors = FALSE)

dim(capture)
head(capture)


# Function to compare the two datasets
find_overlaps_AD_table <- function (regions_1,regions_2,tolerance=0)
{
  common_region_table = NULL
  number_of_regions_1 <- nrow(regions_1)
  number_of_regions_2 <- nrow(regions_2)
  
  overlapped_is <- c()
  overlapped_js <- c()
  
  for (i in 1:number_of_regions_1) {
    
    for (j in 1:number_of_regions_2) {
      
      if (as.character(regions_1[i,1])==as.character(regions_2[j,1])) {
        start_1 <- as.numeric(regions_1[i,2])
        end_1 <- as.numeric(regions_1[i,3])
        expanded_start_1 <- start_1 - tolerance
        expanded_end_1 <- end_1 + tolerance
        start_2 <- as.numeric(regions_2[j,2])
        end_2 <- as.numeric(regions_2[j,3])
        
        if (expanded_end_1>=start_2) {
          
          if (end_2>=expanded_start_1) {
            
            overlapped_is <- c(overlapped_is,i)
            overlapped_js <- c(overlapped_js,j)
            
            first_region <- paste(regions_1[i,1],start_1,end_1,sep="_")
            second_region <- paste(regions_2[j,1],start_2,end_2,sep="_")
            overlap_size <- min(end_1,end_2)-max(start_1,start_2)+1
            common_region_table = rbind(common_region_table,c(as.character(regions_1[i,1]),start_1,end_1, 
                                                              as.character(regions_2[j,1]),start_2,end_2, overlap_size))
            
            if (overlap_size>0) {
              print(paste(first_region,second_region,sep=" "))
            } else {
              print(paste(first_region,second_region, sep=" "))
            }
          }
        } 
      }
    }
  }
  print(paste("Number matched in first data set:",length(unique(overlapped_is))))
  print(paste("Number matched in second data set:",length(unique(overlapped_js))))
  if (length(common_region_table) > 0){
    colnames(common_region_table) = c("chr_reg1", "start_reg1", "end_reg1", "chr_reg2", "start_reg2", "end_reg2","overlap_size")
  }else{
  }
  return(as.data.frame(common_region_table))
}


          
# Comparisons of DEG and capture regions - dist0
capt_DEG_u14_15 = find_overlaps_AD_table(DEG_u14_15,capture,tolerance=0)
dim(capt_DEG_u14_15)
write.table(capt_DEG_u14_15,
            "H:/methylome/fusion/post_methylkit/DMR432_pval0.07/croisement_autre_data/RNAseq_capture/capt_DEG_up_14_15_dist0.txt", 
            sep = "\t", quote=FALSE, row.names = FALSE, col.names=FALSE)


capt_DEG_u15_16 = find_overlaps_AD_table(DEG_u15_16,capture,tolerance=0)
dim(capt_DEG_u15_16)
write.table(capt_DEG_u15_16,
            "H:/methylome/fusion/post_methylkit/DMR432_pval0.07/croisement_autre_data/RNAseq_capture/capt_DEG_up_15_16_dist0.txt", 
            sep = "\t", quote=FALSE, row.names = FALSE, col.names=FALSE)


capt_DEG_d14_15 = find_overlaps_AD_table(DEG_d14_15,capture,tolerance=0)
dim(capt_DEG_d14_15)
write.table(capt_DEG_d14_15,
            "H:/methylome/fusion/post_methylkit/DMR432_pval0.07/croisement_autre_data/RNAseq_capture/capt_DEG_down_14_15_dist0.txt", 
            sep = "\t", quote=FALSE, row.names = FALSE, col.names=FALSE)


capt_DEG_d15_16 = find_overlaps_AD_table(DEG_d15_16,capture,tolerance=0)
dim(capt_DEG_d15_16)
write.table(capt_DEG_d15_16,
            "H:/methylome/fusion/post_methylkit/DMR432_pval0.07/croisement_autre_data/RNAseq_capture/capt_DEG_down_15_16_dist0.txt", 
            sep = "\t", quote=FALSE, row.names = FALSE, col.names=FALSE)


# Comparisons of DEG and capture regions - dist1000
large_capt_DEG_u14_15 = find_overlaps_AD_table(DEG_u14_15,capture,tolerance=1000)
dim(large_capt_DEG_u14_15)
write.table(large_capt_DEG_u14_15,
            "H:/methylome/fusion/post_methylkit/DMR432_pval0.07/croisement_autre_data/RNAseq_capture/large_capt_DEG_up_14_15_dist1000.txt", 
            sep = "\t", quote=FALSE, row.names = FALSE, col.names=FALSE)


large_capt_DEG_u15_16 = find_overlaps_AD_table(DEG_u15_16,capture,tolerance=1000)
dim(large_capt_DEG_u15_16)
write.table(large_capt_DEG_u15_16,
            "H:/methylome/fusion/post_methylkit/DMR432_pval0.07/croisement_autre_data/RNAseq_capture/large_capt_DEG_up_15_16_dist1000.txt", 
            sep = "\t", quote=FALSE, row.names = FALSE, col.names=FALSE)


large_capt_DEG_d14_15 = find_overlaps_AD_table(DEG_d14_15,capture,tolerance=1000)
dim(large_capt_DEG_d14_15)
write.table(large_capt_DEG_d14_15,
            "H:/methylome/fusion/post_methylkit/DMR432_pval0.07/croisement_autre_data/RNAseq_capture/large_capt_DEG_down_14_15_dist1000.txt", 
            sep = "\t", quote=FALSE, row.names = FALSE, col.names=FALSE)


large_capt_DEG_d15_16 = find_overlaps_AD_table(DEG_d15_16,capture,tolerance=1000)
dim(large_capt_DEG_d15_16)
write.table(large_capt_DEG_d15_16,
            "H:/methylome/fusion/post_methylkit/DMR432_pval0.07/croisement_autre_data/RNAseq_capture/large_capt_DEG_down_15_16_dist1000.txt", 
            sep = "\t", quote=FALSE, row.names = FALSE, col.names=FALSE)
```

#### Are DEG and DOCR over-represented among DMR ?¶

To estimate whether DMRs are particularly located to DEG or DOCR observed during physiological brain development, hypergeometric tests were done as follow:

In [ ]:

```
%%R

# Hypergeomtric tests on DOCR among DMR :
# We observed: 
# 2 DMR that are also DOCR_down between E14.5 and E15.5 (dist0)
# 0 DMR that are also DOCR_down between E15.5 and E16.5 (dist0)
# 5 DMR that are also DOCR_up between E14.5 and E15.5 (dist0)
# 1 DMR that are also DOCR_up between E15.5 and E16.5 (dist0)

# 3 DMR that are also DOCR_down between E14.5 and E15.5 (dist1000)
# 0 DMR that are also DOCR_down between E15.5 and E16.5 (dist1000)
# 5 DMR that are also DOCR_up between E14.5 and E15.5 (dist1000)
# 1 DMR that are also DOCR_up between E15.5 and E16.5 (dist1000)

# We observed 432 DMR in total
# among 58 611 regions (all capture regions) 
nb_DMR = 432
nb_capt_tot = 58611


# For DMR-DOCR down E14.15_dist0 : 
nb_ov = 2
nb_ATAC_capt = 449
phyper(nb_ov, nb_ATAC_capt, nb_capt_tot - nb_ATAC_capt, nb_DMR, lower.tail = FALSE)
phyper(nb_ov, nb_ATAC_capt, nb_capt_tot - nb_ATAC_capt, nb_DMR, lower.tail = TRUE)

# For DMR-DOCR down E15.16_dist0 : 
nb_ov = 0
nb_ATAC_capt = 79
phyper(nb_ov, nb_ATAC_capt, nb_capt_tot - nb_ATAC_capt, nb_DMR, lower.tail = FALSE)
phyper(nb_ov, nb_ATAC_capt, nb_capt_tot - nb_ATAC_capt, nb_DMR, lower.tail = TRUE)

# For DMR-DOCR up E14.15_dist0 : 
nb_ov = 5
nb_ATAC_capt = 313 
phyper(nb_ov, nb_ATAC_capt, nb_capt_tot - nb_ATAC_capt, nb_DMR, lower.tail = FALSE)
phyper(nb_ov, nb_ATAC_capt, nb_capt_tot - nb_ATAC_capt, nb_DMR, lower.tail = TRUE)

# For DMR-DOCR up E15.16_dist0 : 
nb_ov = 1
nb_ATAC_capt = 75
phyper(nb_ov, nb_ATAC_capt, nb_capt_tot - nb_ATAC_capt, nb_DMR, lower.tail = FALSE)
phyper(nb_ov, nb_ATAC_capt, nb_capt_tot - nb_ATAC_capt, nb_DMR, lower.tail = TRUE)


# For DMR-DOCR down E14.15_dist1000 : 
nb_ov = 3
nb_ATAC_capt = 561
phyper(nb_ov, nb_ATAC_capt, nb_capt_tot - nb_ATAC_capt, nb_DMR, lower.tail = FALSE)
phyper(nb_ov, nb_ATAC_capt, nb_capt_tot - nb_ATAC_capt, nb_DMR, lower.tail = TRUE)

# For DMR-DOCR down E15.16_dist1000 : 
nb_ov = 0
nb_ATAC_capt = 119
phyper(nb_ov, nb_ATAC_capt, nb_capt_tot - nb_ATAC_capt, nb_DMR, lower.tail = FALSE)
phyper(nb_ov, nb_ATAC_capt, nb_capt_tot - nb_ATAC_capt, nb_DMR, lower.tail = TRUE)

# For DMR-DOCR up E14.15_dist1000 : 
nb_ov = 5
nb_ATAC_capt = 377
phyper(nb_ov, nb_ATAC_capt, nb_capt_tot - nb_ATAC_capt, nb_DMR, lower.tail = FALSE)
phyper(nb_ov, nb_ATAC_capt, nb_capt_tot - nb_ATAC_capt, nb_DMR, lower.tail = TRUE)

# For DMR-DOCR up E15.16_dist1000 : 
nb_ov = 1
nb_ATAC_capt = 85
phyper(nb_ov, nb_ATAC_capt, nb_capt_tot - nb_ATAC_capt, nb_DMR, lower.tail = FALSE)
phyper(nb_ov, nb_ATAC_capt, nb_capt_tot - nb_ATAC_capt, nb_DMR, lower.tail = TRUE)


# Hypergeomtric tests on DEG among DMR :
# We observed: 
# 21 DMR that are also DEG_down between E14.5 and E15.5 (dist0)
# 6 DMR that are also DEG_down between E15.5 and E16.5 (dist0)
# 96 DMR that are also DEG_up between E14.5 and E15.5 (dist0), but 73 uniq regions from dataset RNA and 88 from dataset DMR
# 16 DMR that are also DEG_up between E15.5 and E16.5 (dist0), but 13 uniq regions from dataset RNA and 16 from dataset DMR

# 27 DMR that are also DEG_down between E14.5 and E15.5 (dist1000), but 27 uniq regions from dataset RNA and 26 from dataset DMR
# 10 DMR that are also DEG_down between E15.5 and E16.5 (dist1000)
# 106 DMR that are also DEG_up between E14.5 and E15.5 (dist1000), but 83 uniq regions from dataset RNA and 98 from dataset DMR
# 18 DMR that are also DEG_up between E15.5 and E16.5 (dist1000), but 15 uniq regions from dataset RNA and 18 from dataset DMR

# We observed 432 DMR in total
# among 58 611 regions (all capture regions) 
nb_DMR = 432
nb_capt_tot = 58611

# For DEG down E14.15_dist0 : 
nb_ov = 21
nb_RNA_capt = 4678
phyper(nb_ov, nb_RNA_capt, nb_capt_tot - nb_RNA_capt, nb_DMR, lower.tail = FALSE)
phyper(nb_ov, nb_RNA_capt, nb_capt_tot - nb_RNA_capt, nb_DMR, lower.tail = TRUE)

# For DEG down E15.16_dist0 : 
nb_ov = 6
nb_RNA_capt = 826
phyper(nb_ov, nb_RNA_capt, nb_capt_tot - nb_RNA_capt, nb_DMR, lower.tail = FALSE)
phyper(nb_ov, nb_RNA_capt, nb_capt_tot - nb_RNA_capt, nb_DMR, lower.tail = TRUE)

# For DEG up E14.15_dist0 : 
nb_ov = 96
nb_RNA_capt = 10458
  phyper(nb_ov, nb_RNA_capt, nb_capt_tot - nb_RNA_capt, nb_DMR, lower.tail = FALSE)
phyper(nb_ov, nb_RNA_capt, nb_capt_tot - nb_RNA_capt, nb_DMR, lower.tail = TRUE)

# For DEG up E15.16_dist0 : 
nb_ov = 16
nb_RNA_capt = 1465
  phyper(nb_ov, nb_RNA_capt, nb_capt_tot - nb_RNA_capt, nb_DMR, lower.tail = FALSE)
phyper(nb_ov, nb_RNA_capt, nb_capt_tot - nb_RNA_capt, nb_DMR, lower.tail = TRUE)


# For DEG down E14.15_dist1000 : 
nb_ov = 27
nb_RNA_capt = 5250
phyper(nb_ov, nb_RNA_capt, nb_capt_tot - nb_RNA_capt, nb_DMR, lower.tail = FALSE)
phyper(nb_ov, nb_RNA_capt, nb_capt_tot - nb_RNA_capt, nb_DMR, lower.tail = TRUE)

# For DEG down E15.16_dist1000 : 
nb_ov = 10
nb_RNA_capt = 902
phyper(nb_ov, nb_RNA_capt, nb_capt_tot - nb_RNA_capt, nb_DMR, lower.tail = FALSE)
phyper(nb_ov, nb_RNA_capt, nb_capt_tot - nb_RNA_capt, nb_DMR, lower.tail = TRUE)

# For DEG up E14.15_dist1000 : 
nb_ov = 106
nb_RNA_capt = 11141
  phyper(nb_ov, nb_RNA_capt, nb_capt_tot - nb_RNA_capt, nb_DMR, lower.tail = FALSE)
phyper(nb_ov, nb_RNA_capt, nb_capt_tot - nb_RNA_capt, nb_DMR, lower.tail = TRUE)

# For DEG up E15.16_dist1000 : 
nb_ov = 18
nb_RNA_capt = 1508
  phyper(nb_ov, nb_RNA_capt, nb_capt_tot - nb_RNA_capt, nb_DMR, lower.tail = FALSE)
phyper(nb_ov, nb_RNA_capt, nb_capt_tot - nb_RNA_capt, nb_DMR, lower.tail = TRUE)
```
