## Supplementary material for "Immediate perturbations of DNA methylation and transcriptome upon acute prenatal alcohol exposure in the mouse developing brain cortex": Supp data Notebook 5 Statistical report Sartools-edgeR-RNAseq-PBSvsEtOH: Supp. data Notebook 5 Statistical report Sartools-edgeR-RNAseq-PBSvsEtOH.html

Statistical report of project Sartools-edgeR-RNAseq-PBSvsEtOH: pairwise comparison(s) of conditions with edgeR


### Statistical report of project Sartools-edgeR-RNAseq-PBSvsEtOH: pairwise comparison(s) of conditions with edgeR

###### Agathe Duchateau

#### 2022-01-17

The SARTools R package which generated this report has been developped at PF2 - Institut Pasteur by M.-A. Dillies and H. Varet. Thanks to cite H. Varet, L. Brillet-Guéguen, J.-Y. Coppee and M.-A. Dillies, *SARTools: A DESeq2- and EdgeR-Based R Pipeline for Comprehensive Differential Analysis of RNA-Seq Data*, PLoS One, 2016, doi: http://dx.doi.org/10.1371/journal.pone.0157022 when using this tool for any analysis published.

### 1 Introduction

The analyses reported in this document are part of the Sartools-edgeR-RNAseq-PBSvsEtOH project. The aim is to find features that are differentially expressed between PBS and EtOH. The statistical analysis process includes data normalization, graphical exploration of raw and normalized data, test for differential expression for each feature between the conditions, raw p-value adjustment and export of lists of features having a significant differential expression between the conditions.

The analysis is performed using the R software [1], Bioconductor [2] packages including edgeR [3] and the SARTools package developed at PF2 - Institut Pasteur. Normalization and differential analysis are carried out according to the edgeR model and package. This report comes with additional tab-delimited text files that contain lists of differentially expressed features.

For more details about the edgeR methodology, please refer to its related publications [3–6].

### 2 Description of raw data

The count data files and associated biological conditions are listed in the following table.

Table 1: Data files and associated biological conditions.

| label | files | group | day |
| --- | --- | --- | --- |
| PBS1 | mm10\_strict\_Std\_tg\_PBS1.txt | PBS | batch\_n1 |
| PBS2 | mm10\_strict\_Std\_tg\_PBS2.txt | PBS | batch\_n1 |
| PBS3 | mm10\_strict\_Std\_tg\_PBS3.txt | PBS | batch\_n1 |
| PBS4 | mm10\_strict\_Std\_tg\_PBS4.txt | PBS | batch\_n1 |
| PBS5 | mm10\_strict\_Std\_tg\_PBS5.txt | PBS | batch\_n2 |
| PBS6 | mm10\_strict\_Std\_tg\_PBS6.txt | PBS | batch\_n2 |
| PBS7 | mm10\_strict\_Std\_tg\_PBS7.txt | PBS | batch\_n2 |
| PBS8 | mm10\_strict\_Std\_tg\_PBS8.txt | PBS | batch\_n2 |
| EtOH1 | mm10\_strict\_Std\_tg\_EtOH1.txt | EtOH | batch\_n1 |
| EtOH2 | mm10\_strict\_Std\_tg\_EtOH2.txt | EtOH | batch\_n1 |
| EtOH3 | mm10\_strict\_Std\_tg\_EtOH3.txt | EtOH | batch\_n1 |
| EtOH4 | mm10\_strict\_Std\_tg\_EtOH4.txt | EtOH | batch\_n1 |
| EtOH5 | mm10\_strict\_Std\_tg\_EtOH5.txt | EtOH | batch\_n2 |
| EtOH6 | mm10\_strict\_Std\_tg\_EtOH6.txt | EtOH | batch\_n2 |
| EtOH7 | mm10\_strict\_Std\_tg\_EtOH7.txt | EtOH | batch\_n2 |
| EtOH8 | mm10\_strict\_Std\_tg\_EtOH8.txt | EtOH | batch\_n2 |

After loading the data we first have a look at the raw data table itself. The data table contains one row per annotated feature and one column per sequenced sample. Row names of this table are feature IDs (unique identifiers). The table contains raw count values representing the number of reads that map onto the features. For this project, there are 35809 features in the count data table.

Table 2: Partial view of the count data table.

|  | PBS1 | PBS2 | PBS3 | PBS4 | PBS5 | PBS6 | PBS7 | PBS8 | EtOH1 | EtOH2 |
| --- | --- | --- | --- | --- | --- | --- | --- | --- | --- | --- |
| ENSMUSG00000000001.4 | 4127 | 3827 | 4009 | 3955 | 5868 | 3447 | 3891 | 3669 | 4315 | 4168 |
| ENSMUSG00000000028.15 | 327 | 274 | 316 | 315 | 484 | 279 | 335 | 327 | 347 | 268 |
| ENSMUSG00000000031.16 | 606 | 648 | 669 | 865 | 881 | 525 | 797 | 591 | 861 | 599 |
| ENSMUSG00000000037.16 | 134 | 121 | 138 | 154 | 215 | 105 | 127 | 130 | 163 | 171 |
| ENSMUSG00000000049.11 | 1 | 0 | 0 | 1 | 0 | 0 | 0 | 1 | 0 | 0 |
| ENSMUSG00000000056.7 | 3019 | 2944 | 3210 | 3218 | 4748 | 2641 | 3026 | 3141 | 2956 | 2551 |

Looking at the summary of the count table provides a basic description of these raw counts (min and max values, median, etc).

Table 3: Summary of the raw counts.

|  | Min. | 1st Qu. | Median | Mean | 3rd Qu. | Max. |
| --- | --- | --- | --- | --- | --- | --- |
| PBS1 | 0 | 0 | 4 | 583 | 324 | 143871 |
| PBS2 | 0 | 0 | 4 | 551 | 302 | 141769 |
| PBS3 | 0 | 0 | 4 | 567 | 312 | 144113 |
| PBS4 | 0 | 0 | 5 | 591 | 331 | 153016 |
| PBS5 | 0 | 0 | 7 | 878 | 495 | 224174 |
| PBS6 | 0 | 0 | 4 | 499 | 282 | 132898 |
| PBS7 | 0 | 0 | 5 | 570 | 319 | 146235 |
| PBS8 | 0 | 0 | 5 | 561 | 316 | 137571 |
| EtOH1 | 0 | 0 | 5 | 580 | 333 | 137628 |
| EtOH2 | 0 | 0 | 6 | 588 | 339 | 137593 |
| EtOH3 | 0 | 0 | 4 | 552 | 309 | 142602 |
| EtOH4 | 0 | 0 | 3 | 396 | 222 | 106296 |
| EtOH5 | 0 | 0 | 6 | 572 | 333 | 137317 |
| EtOH6 | 0 | 0 | 5 | 556 | 318 | 129085 |
| EtOH7 | 0 | 0 | 7 | 633 | 361 | 151585 |
| EtOH8 | 0 | 0 | 5 | 537 | 311 | 119238 |

Figure 1 shows the total number of mapped and counted reads for each sample. We expect total read counts to be similar within conditions, they may be different across conditions. Total counts sometimes vary widely between replicates. This may happen for several reasons, including:

- different rRNA contamination levels between samples (even between biological replicates);
- slight differences between library concentrations, since they may be difficult to measure with high precision.

Figure 1: Number of mapped reads per sample. Colors refer to the biological condition of the sample.

Figure 2 shows the percentage of features with no read count in each sample. We expect this percentage to be similar within conditions. Features with null read counts in the 16 samples will not be taken into account for the analysis with edgeR. Here, 0 features (0%) are in this situation (dashed line).

Figure 2: Percentage of features with null read counts in each sample.

Figure 3 shows the distribution of read counts for each sample (on a log scale to improve readability). Again we expect replicates to have similar distributions. In addition, this figure shows if read counts are preferably low, medium or high. This depends on the organisms as well as the biological conditions under consideration.

Figure 3: Density distribution of read counts.

It may happen that one or a few features capture a high proportion of reads (up to 20% or more). This phenomenon should not influence the normalization process. The edgeR normalization has proved to be robust to this situation [7]. Anyway, we expect these high count features to be the same across replicates. They are not necessarily the same across conditions. Figure 4 and table 4 illustrate the possible presence of such high count features in the data set.

Figure 4: Percentage of reads associated with the sequence having the highest count (provided in each box on the graph) for each sample.

Table 4: Percentage of reads associated with the sequences having the highest counts.

|  | ENSMUSG00000064351.1 | ENSMUSG00000037742.14 | ENSMUSG00000064370.1 | ENSMUSG00000106106.2 |
| --- | --- | --- | --- | --- |
| PBS1 | 0.69 | 0.52 | 0.47 | 0.09 |
| PBS2 | 0.72 | 0.52 | 0.50 | 0.70 |
| PBS3 | 0.71 | 0.52 | 0.48 | 0.09 |
| PBS4 | 0.72 | 0.51 | 0.54 | 0.07 |
| PBS5 | 0.71 | 0.49 | 0.53 | 0.09 |
| PBS6 | 0.74 | 0.53 | 0.51 | 0.25 |
| PBS7 | 0.72 | 0.51 | 0.53 | 0.09 |
| PBS8 | 0.68 | 0.49 | 0.54 | 0.35 |
| EtOH1 | 0.66 | 0.48 | 0.43 | 0.09 |
| EtOH2 | 0.65 | 0.50 | 0.42 | 0.08 |
| EtOH3 | 0.72 | 0.51 | 0.47 | 0.07 |
| EtOH4 | 0.75 | 0.50 | 0.44 | 0.06 |
| EtOH5 | 0.67 | 0.49 | 0.44 | 0.07 |
| EtOH6 | 0.65 | 0.50 | 0.43 | 0.06 |
| EtOH7 | 0.67 | 0.43 | 0.48 | 0.11 |
| EtOH8 | 0.62 | 0.47 | 0.45 | 0.22 |

We may wish to assess the similarity between samples across conditions. A pairwise scatter plot is produced (figure 5) to show how replicates and samples from different biological conditions are similar or different (using a log scale). Moreover, as the Pearson correlation has been shown not to be relevant to measure the similarity between replicates, the SERE statistic has been proposed as a similarity index between RNA-Seq samples [8]. It measures whether the variability between samples is random Poisson variability or higher. Pairwise SERE values are printed in the lower triangle of the pairwise scatter plot. The value of the SERE statistic is:

- 0 when samples are identical (no variability at all: this may happen in the case of a sample duplication);
- 1 for technical replicates (technical variability follows a Poisson distribution);
- greater than 1 for biological replicates and samples from different biological conditions (biological variability is higher than technical one, data are over-dispersed with respect to Poisson). The higher the SERE value, the lower the similarity. It is expected to be lower between biological replicates than between samples of different biological conditions. Hence, the SERE statistic can be used to detect inversions between samples.

Figure 5: Pairwise comparison of samples (not produced when more than 12 samples).

### 3 Filtering low counts

edgeR suggests to filter features with null or low counts because they do not supply much information. For this project, 21416 features (59.81%) have been removed from the analysis because they did not satisfy the following condition: having at least 1 counts-per-million in at least 8 samples.

### 4 Variability within the experiment: data exploration

The main variability within the experiment is expected to come from biological differences between the samples. This can be checked in two ways. The first one is to perform a hierarchical clustering of the whole sample set. This is performed after a transformation of the count data as moderated log-counts-per-million. Figure 6 shows the dendrogram obtained from CPM data. An euclidean distance is computed between samples, and the dendrogram is built upon the Ward criterion. We expect this dendrogram to group replicates and separate biological conditions.

Figure 6: Sample clustering based on normalized data.

Another way of visualizing the experiment variability is to look at the first two dimensions of a multidimensional scaling plot, as shown on figure 7. On this figure, the first dimension is expected to separate samples from the different biological conditions, meaning that the biological variability is the main source of variance in the data.

Figure 7: Multidimensional scaling plot of the samples.

### 5 Normalization

Normalization aims at correcting systematic technical biases in the data, in order to make read counts comparable across samples. The normalization proposed by edgeR is called Trimmed Mean of M-values (TMM) but it is also possible to use the RLE (DESeq) or upperquartile normalizations. It relies on the hypothesis that most features are not differentially expressed.

edgeR computes a factor for each sample. These normalization factors apply to the total number of counts and cannot be used to normalize read counts in a direct manner. Indeed, normalization factors are used to normalize total counts. These in turn are used to normalize read counts according to a total count normalization: if \(N\_j\) is the total number of reads of the sample \(j\) and \(f\_j\) its normalization factor, \(N'\_j=f\_j \times N\_j\) is the normalized total number of reads. Then, let \(s\_j=N'\_j/\bar{N'}\) with \(\bar{N'}\) the mean of the \(N'\_j\) s. Finally, the normalized counts of the sample \(j\) are defined as \(x'\_{ij}=x\_{ij}/s\_j\) where \(i\) is the gene index.

Table 5: Normalization factors.

|  | PBS1 | PBS2 | PBS3 | PBS4 | PBS5 | PBS6 | PBS7 | PBS8 | EtOH1 | EtOH2 | EtOH3 | EtOH4 | EtOH5 | EtOH6 | EtOH7 | EtOH8 |
| --- | --- | --- | --- | --- | --- | --- | --- | --- | --- | --- | --- | --- | --- | --- | --- | --- |
| TMM normalization factors | 0.99 | 0.98 | 0.99 | 1 | 1 | 1 | 1 | 1 | 1.01 | 1.01 | 1 | 1 | 1.02 | 1.01 | 1 | 1.01 |

Boxplots are often used to assess the quality of the normalization process, as they show how distributions are globally affected during this process. We expect normalization to stabilize distributions across samples. Figure 8 shows boxplots of raw (left) and normalized (right) data respectively.

Figure 8: Boxplots of raw (left) and normalized (right) read counts.

### 6 Differential analysis

#### 6.1 Modelization

edgeR aims at fitting one linear model per feature. For this project, the design used is ~ group and the goal is to estimate the models’ coefficients which can be interpreted as \(\log\_2(\texttt{FC})\). These coefficients will then be tested to get p-values and adjusted p-values.

#### 6.2 Dispersions estimation

The edgeR model assumes that the count data follow a negative binomial distribution which is a robust alternative to the Poisson law when data are over-dispersed (the variance is higher than the mean). The first step of the statistical procedure is to estimate the dispersion of the data.

Figure 9: Dispersion estimates.

Figure 9 shows the result of the dispersion estimation step. The x- and y-axes represent the mean count value and the estimated dispersion respectively. Black dots represent empirical dispersion estimates for each feature (from the observed count values). The blue curve shows the relationship between the means of the counts and the dispersions modeled with splines. The red segment represents the common dispersion.

#### 6.3 Statistical test for differential expression

Once the dispersion estimation and the model fitting have been done, edgeR can perform the statistical testing. Figure 10 shows the distributions of raw p-values computed by the statistical test for the comparison(s) done. This distribution is expected to be a mixture of a uniform distribution on \([0,1]\) and a peak around 0 corresponding to the differentially expressed features.

Figure 10: Distribution(s) of raw p-values.

#### 6.4 Final results

A p-value adjustment is performed to take into account multiple testing and control the false positive rate to a chosen level \(\alpha\). For this analysis, a BH p-value adjustment was performed [9,10] and the level of controlled false positive rate was set to 0.05.

Table 6: Number of up-, down- and total number of differentially expressed features for each comparison.

| Test vs Ref | # down | # up | # total |
| --- | --- | --- | --- |
| EtOH vs PBS | 2210 | 2598 | 4808 |

Figure 11 represents the MA-plot of the data for the comparisons done, where differentially expressed features are highlighted in red. A MA-plot represents the log ratio of differential expression as a function of the mean intensity for each feature. Triangles correspond to features having a too low/high \(\log\_2(\text{FC})\) to be displayed on the plot.

Figure 11: MA-plot(s) of each comparison. Red dots represent significantly differentially expressed features.

Figure 12 shows the volcano plots for the comparisons performed and differentially expressed features are still highlighted in red. A volcano plot represents the log of the adjusted P value as a function of the log ratio of differential expression.

Figure 12: Volcano plot(s) of each comparison. Red dots represent significantly differentially expressed features.

Full results as well as lists of differentially expressed features are provided in the following text files which can be easily read in a spreadsheet. For each comparison:

- TestVsRef.complete.txt contains results for all the features;
- TestVsRef.up.txt contains results for up-regulated features. Features are ordered from the most significant adjusted p-value to the less significant one;
- TestVsRef.down.txt contains results for down-regulated features. Features are ordered from the most significant adjusted p-value to the less significant one.

These files contain the following columns:

- Id: unique feature identifier;
- sampleName: raw counts per sample;
- norm.sampleName: rounded normalized counts per sample;
- baseMean: base mean over all samples;
- PBS and EtOH: means (rounded) of normalized counts of the biological conditions;
- FoldChange: fold change of expression, calculated as \(2^{\log\_2(\text{FC})}\);
- log2FoldChange: \(\log\_2(\text{FC})\) as estimated by the GLM model. It reflects the differential expression between Test and Ref and can be interpreted as \(\log\_2(\frac{\text{Test}}{\text{Ref}})\). If this value is:
  - around 0: the feature expression is similar in both conditions;
  - positive: the feature is up-regulated (\(\text{Test} > \text{Ref}\));
  - negative: the feature is down-regulated (\(\text{Test} < \text{Ref}\));
- pvalue: raw p-value from the statistical test;
- padj: adjusted p-value on which the cut-off \(\alpha\) is applied;
- tagwise.dispersion: dispersion parameter estimated from feature counts (i.e. black dots on figure 9);
- trended.dispersion: dispersion parameter estimated with splines (i.e. blue curve on figure 9).

### 7 R session information and parameters

The versions of the R software and Bioconductor packages used for this analysis are listed below. It is important to save them if one wants to re-perform the analysis in the same conditions.

- R version 4.0.3 (2020-10-10), x86\_64-w64-mingw32
- Locale: LC\_COLLATE=French\_France.1252, LC\_CTYPE=French\_France.1252, LC\_MONETARY=French\_France.1252, LC\_NUMERIC=C, LC\_TIME=French\_France.1252
- Running under: Windows >= 8 x64 (build 9200)
- Matrix products: default
- Base packages: base, datasets, graphics, grDevices, methods, parallel, stats, stats4, utils
- Other packages: Biobase 2.50.0, BiocGenerics 0.36.1, DESeq2 1.30.1, edgeR 3.32.1, GenomeInfoDb 1.26.7, GenomicRanges 1.42.0, GGally 2.1.2, ggplot2 3.3.5, IRanges 2.24.1, kableExtra 1.3.4, limma 3.46.0, MatrixGenerics 1.2.1, matrixStats 0.61.0, reshape2 1.4.4, S4Vectors 0.28.1, SARTools 1.7.3, SummarizedExperiment 1.20.0
- Loaded via a namespace (and not attached): annotate 1.68.0, AnnotationDbi 1.52.0, assertthat 0.2.1, BiocParallel 1.24.1, bit 4.0.4, bit64 4.0.5, bitops 1.0-7, blob 1.2.2, bslib 0.3.1, cachem 1.0.6, colorspace 2.0-2, compiler 4.0.3, crayon 1.4.1, DBI 1.1.1, DelayedArray 0.16.3, digest 0.6.27, dplyr 1.0.7, ellipsis 0.3.2, evaluate 0.14, fansi 0.5.0, farver 2.1.0, fastmap 1.1.0, genefilter 1.72.1, geneplotter 1.68.0, generics 0.1.0, GenomeInfoDbData 1.2.4, ggdendro 0.1.22, ggrepel 0.9.1, glue 1.4.2, grid 4.0.3, gridExtra 2.3, gtable 0.3.0, highr 0.9, hms 1.1.1, htmltools 0.5.2, httr 1.4.2, jquerylib 0.1.4, jsonlite 1.7.2, knitr 1.36, labeling 0.4.2, lattice 0.20-45, lifecycle 1.0.1, locfit 1.5-9.4, magrittr 2.0.1, MASS 7.3-54, Matrix 1.3-4, memoise 2.0.0, munsell 0.5.0, pillar 1.6.3, pkgconfig 2.0.3, plyr 1.8.6, prettyunits 1.1.1, progress 1.2.2, purrr 0.3.4, R6 2.5.1, RColorBrewer 1.1-2, Rcpp 1.0.7, RCurl 1.98-1.5, reshape 0.8.8, rlang 0.4.10, rmarkdown 2.11, RSQLite 2.2.8, rstudioapi 0.13, rvest 1.0.1, sass 0.4.0, scales 1.1.1, splines 4.0.3, stringi 1.7.5, stringr 1.4.0, survival 3.2-13, svglite 2.0.0, systemfonts 1.0.2, tibble 3.1.5, tidyselect 1.1.1, tools 4.0.3, utf8 1.2.2, vctrs 0.3.8, viridisLite 0.4.0, webshot 0.5.2, withr 2.4.2, xfun 0.26, XML 3.99-0.8, xml2 1.3.2, xtable 1.8-4, XVector 0.30.0, yaml 2.2.1, zlibbioc 1.36.0

Parameter values used for this analysis are:

- workDir: G:/RNA\_seq\_AD1/results/06\_DEG\_research/Sartools\_mm10/Analyse\_1/
- projectName: Sartools-edgeR-RNAseq-PBSvsEtOH
- author: Agathe Duchateau
- targetFile: G:/RNA\_seq\_AD1/results/06\_DEG\_research/Sartools\_mm10/target\_PBS\_vs\_EtOH.txt
- rawDir: G:/RNA\_seq\_AD1/results/05\_count\_table/output\_strict\_mm10\_Std/ensembl
- featuresToRemove: alignment\_not\_unique, ambiguous, no\_feature, not\_aligned, too\_low\_aQual
- varInt: group
- condRef: PBS
- batch: NULL
- alpha: 0.05
- pAdjustMethod: BH
- cpmCutoff: 1
- gene.selection: pairwise
- normalizationMethod: TMM
- colors: darkblue, brown3

### Bibliography

1. R Core Team. *R: A language and environment for statistical computing*. Vienna, Austria : R Foundation for Statistical Computing, 2017 :

2. Gentleman RC, Carey VJ, Bates DM *et al.* Bioconductor: Open software development for computational biology and bioinformatics. *Genome Biology* 2004 ; 5: R80.

3. Robinson M, McCarthy D, Smyth G. EdgeR: A bioconductor package for differential expression analysis of digital gene expression data. *Bioinformatics* 2010 ; 26: 139.

4. Robinson M, Smyth G. Moderated statistical tests for assessing differences in tag abundance. *Bioinformatics* 2007 ; 23: 2881.

5. Robinson M, Smyth G. Small-sample estimation of negative binomial dispersion, with applications to sage data. *Biostatistics* 2008 ; 9: 321.

6. McCarthy D, Chen Y, Smyth G. Differential expression analysis of multifactor rna-seq experiments with respect to biological variation. *Nucleic Acids Research* 2012 ; 40: 4288.

7. Dillies M-A, Rau A, Aubert J *et al.* A comprehensive evaluation of normalization methods for illumina high-throughput rna sequencing data analysis. *Briefings in Bioinformatics* 2013 ; 14: 671.

8. Schulze SK, Kanwar R, Gölzenleuchter M *et al.* SERE: Single-parameter quality control and sample comparison for rna-seq. *BMC Genomics* 2012 ; 13: 524.

9. Benjamini Y, Hochberg Y. Controlling the false discovery rate: A practical and powerful approach to multiple testing. *Journal of the Royal Statistical Society. Series B (Methodological)* 1995 ; 57: 289–300.

10. Benjamini Y, Yekutieli D. The control of the false discovery rate in multiple testing under dependency. *The Annals of Statistics* 2001 ; 29: 1165–1188.
